# Steric control of signaling bias in the immunometabolic receptor GPR84

**DOI:** 10.1101/2025.07.30.667614

**Authors:** Pinqi Wang, Xuan Zhang, Abdul-Akim Guseinov, Laura Jenkins, Carl von Hallerstein, Jonathan D. Colburn, Rowan Ives, Vincent B. Luscombe, Sara Marsango, Listiana Oktavia, Arun Raja, David R. Greaves, Philip C. Biggin, Graeme Milligan, Cheng Zhang, Irina G. Tikhonova, Angela J. Russell

## Abstract

Biased signaling in G protein-coupled receptors offers therapeutic promise, yet rational design of biased ligands remains challenging due to limited mechanistic understanding. Here, we report a molecular framework for controlling signaling bias at the immunometabolic receptor GPR84. We identified three structurally-matched ligands (OX04529, OX04954, and OX04539) with varying steric profiles that exhibit comparable G_i_ protein activation but dramatically different β-arrestin recruitment capacities. A high-resolution cryo-EM structure of GPR84-G_i_ in complex with OX04529, complemented by molecular dynamics simulations and targeted mutagenesis, revealed that steric interactions between ligand substituents and Leu336^6.52^ and Phe187^5.47^ indirectly disrupt a critical polar network involving Tyr332^6.48^, Asn104^3.36^ and Asn362^7.45^ essential for β-arrestin recruitment. Based on these insights, we developed a steric-dependent model that enabled rational design of G protein-biased agonists with predictable β-arrestin recruitment profiles. This mechanistic framework provides a blueprint for designing biased agonists with customized signaling profiles at GPR84 and potentially other class A GPCRs.

## Introduction

Heptahelical transmembrane G protein-coupled receptors (GPCRs) constitute the most extensive category of cell surface receptors in humans and regulate a wide range of physiological processes^1^. Activation of GPCRs by endogenous and exogenous agonists can promote intracellular signaling through a range of pathways, including heterotrimeric G proteins^2^, and receptor phosphorylation by G protein-coupled receptor- (GRK)^3^ and second messenger-regulated kinases^4^, as well as various adapter proteins including β-arrestins^5^. Recent advances in structural biology have revealed that GPCR-activating ligands can bind at different (orthosteric or allosteric) sites and may exhibit signaling bias by preferentially activating specific pathways^6–8^ (**Fig. 1a**). Such bias leads to distinct functional outcomes, adding complexity to GPCR-mediated signaling. Despite these observations, the molecular mechanisms responsible for GPCR biased agonism are obscure and clinical implications remain restricted to specific receptors^9,10^. This knowledge gap stems from limited ligand diversity, limitations of static structures, and complex receptor dynamics. Elucidating distinct mechanisms of receptor activation and their effects on signaling is crucial for exploiting their therapeutic potential. Such understanding could revolutionize treatment approaches across numerous diseases and open a pathway to rational structure-based design of biased ligands, which has so far been elusive.

**Fig. 1.**
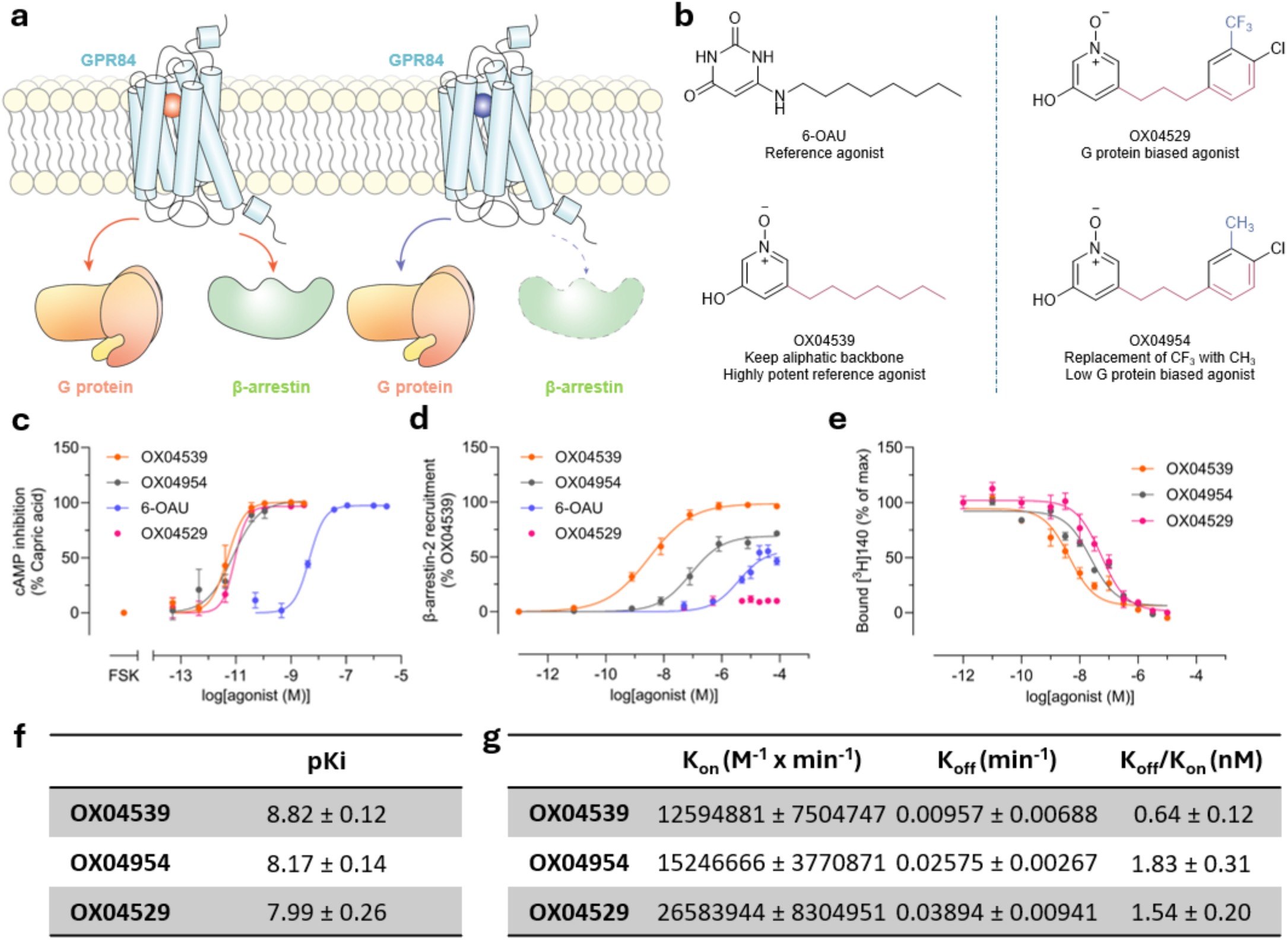
OX04529, OX04954 and OX04539 produce varying levels of β-arrestin-2 recruitment but show comparable G_i_ protein signaling, binding affinity and kinetics. **a,** Occupying the orthosteric binding pocket of GPR84 with different ligands may lead to distinct effector interactions and downstream signaling pathways. b, Structures of reference agonists triggering both G protein and β-arrestin engagement (6-OAU and OX04539) (left) and G protein-biased agonists (OX04529 and OX04954) (right). c, G_i_ activity and (d) β-arrestin-2 recruitment measured by HitHunter cAMP inhibition or PathHunter β-arrestin-2 assays. In cAMP assays data are normalized to the efficacy of capric acid (C10:0) (100 µM). In β-arrestin-2 assays data are normalized to the E_max_ of OX04539. e, Competition binding, (f) calculated inhibitory constants and (g) association/dissociation rates for OX04529, OX04954 and OX04539 at GPR84 were characterized in radioligand binding assays using the orthosteric antagonist [^3^H]140. Data are means ± S.E.M. (n = 3 biological replicates each performed in triplicate).

G protein-coupled receptor 84 (GPR84) is a class A GPCR predominantly expressed by innate immune cells, including monocytes, neutrophils, and macrophages. Upon activation, GPR84 engages multiple downstream effectors, primarily coupling to inhibitory (G_i_) G proteins and, following phosphorylation by GRK2/3,^11^ interacting with β-arrestins (**Fig. 1a**). *In vitro* and *ex vivo* studies have demonstrated that GPR84 activation by agonists can enhance immune cell migration and increase the secretion of cytokines, chemokines, and other inflammatory mediators^12–14^. Reference agonists such as 6-n-octylaminouracil (6-OAU), which promote interactions with both G proteins and β-arrestins, enhance macrophage bacterial adhesion and phagocytosis^15,16^. Notably, 6-OAU can act in synergy with anti-CD47 therapies to promote pro-phagocytic activity against cancer cells^17,18^. While GPR84 activation of G_i_ shows promise as an immunotherapy in cancer, whether interactions with a β-arrestin enhances or limits such pro-phagocytic effects remain unknown. The biased agonist DL-175^16^, while a potent and efficacious activator of G_i_ signaling, fails to induce GPR84 phosphorylation and β-arrestin recruitment^11^ (**Fig. 1a**). Moreover, GPR84 biased agonists have also been shown to enhance macrophage phagocytosis, but in contrast to β-arrestin-recruiting reference ligands, fail to induce chemotaxis in macrophages and fail to induce receptor internalization. Biased agonists may therefore offer therapeutic advantages over β-arrestin-recruiting agonists, but to date there are no models which allow for the rational design and optimization of biased ligands at the receptor to test this hypothesis.

Recent structures of GPR84-G_i_ complexes with the reference synthetic agonists 6-OAU^18^ (**<u>8G05</u>**), LY237^19^ (**<u>8J19</u>**) and a medium chain-length fatty acid^19^ (**<u>8J18</u>**) have provided insights into mechanisms of activation of G_i_ signaling. However, the structural and molecular mechanisms resulting in biased agonism at GPR84 remain unexplored. We recently reported OX04529^20^ as a highly potent, G protein-biased GPR84 agonist that exhibits very limited β-arrestin recruitment. Herein we report detailed structural, dynamic, and functional analyses of the mechanisms by which ligands from this chemical series with varying extents of signaling bias bind to and activate GPR84 to induce G_i_- and β-arrestin-interactions. Our study reveals molecular and atomic-level details of these interactions, providing crucial insights into the mechanisms of activation of an important innate immune receptor. This establishes a structural foundation for the rational design of G protein-biased agonists for GPR84 that display varying levels of β-arrestin engagement, with broad implications for understanding and manipulating biased signaling at other class A GPCRs.

### Minor structural modifications of OX04529 enhance GPR84-β-arrestin interactions

GPR84-β-arrestin-2 complementation studies in CHO-K1 cells expressing GPR84 revealed distinct signaling profiles for OX04529 and two of its structural analogs which retained the polar 3-hydroxy pyridine *N*-oxide head group but varied the non-polar tail (**Fig. 1b**). Whilst each of these displayed similar potency and efficacy and were markedly more potent than the previously characterized GPR84 agonist 6-OAU in both cAMP (**Fig. 1c**) and [^35^S]GTPγS binding assays (**Extended Data Table 1, Extended Data Table 3**), OX04529 failed to promote substantial GPR84-β-arrestin-2 interactions (**Fig. 1d**). By contrast, OX04954, in which CH_3_ replaced the aryl 3-CF_3_ substituent (**Fig. 1b**), gave a moderate, concentration-dependent increase in β-arrestin- 2 recruitment with sub-µM potency (**Fig. 1d**). OX04539, where the aryl ring was replaced with a linear saturated alkyl tail (**Fig. 1b**), similar to endogenous medium-chain fatty acids and synthetic ligands such as 6- OAU and LY237, showed even greater efficacy and potency in promoting GPR84-β-arrestin-2 interactions (**Fig. 1d**). The variation in effectiveness of OX04529, OX04954 and OX04539 in promoting β-arrestin-2 recruitment was not restricted to CHO-K1 cells. Similar compound rank-order of potency and efficacy was also observed for GPR84 expressed in HEK293T cells using a Bioluminescence Resonance Energy Transfer (BRET)-based approach (**Extended Data Fig. 1a**) and was also observed when β-arrestin-2 was replaced by β-arrestin-1 (**Extended Data Fig. 1b**). OX04539 was subsequently used as a high potency structure-matched reference ligand throughout the studies.

OX04529, OX04954 and OX04539 each fully outcompeted specific binding of the high affinity GPR84 orthosteric antagonist [^3^H]140^21^ (**Fig. 1e**), with estimated affinities between 1 and 10 nM (**Fig. 1f**), confirming the orthosteric nature of these ligands. Previous studies have suggested that the kinetics of ligand binding can influence ligand bias, with biased agonists showing comparable association rates but different dissociation rates compared to reference agonists.^22,23^ However, kinetic association and dissociation rates of these ligands were also similar (**Fig. 1g**), with affinity calculated from these rate constants ranging from 1-2 nM, consistent with the competition binding studies. It was hence evident that differences in ligand binding kinetics are unable to explain the observed ligand bias.

### Overall structure and binding mode of the OX04529-GPR84-Gi complex

To elucidate the mechanism of GPR84 biased signaling induced by OX04529, we determined a cryo- EM structure of the GPR84-G_i_ complex with OX04529 at a nominal resolution of 3.03 Å **(Fig. 2a and Extended Data Fig. 2)**. We used the NanoBiT tethering strategy to obtain the complex for structure determination, following the methods we previously used to determine the structure of GPR84-G_i_ complex with 6-OAU (**<u>8G05</u>**)^18^. The clear cryo-EM map allows unambiguous modeling of most of the receptor residues, the Gi heterotrimer, and the ligand (**Fig. 2a).**

**Fig. 2.**
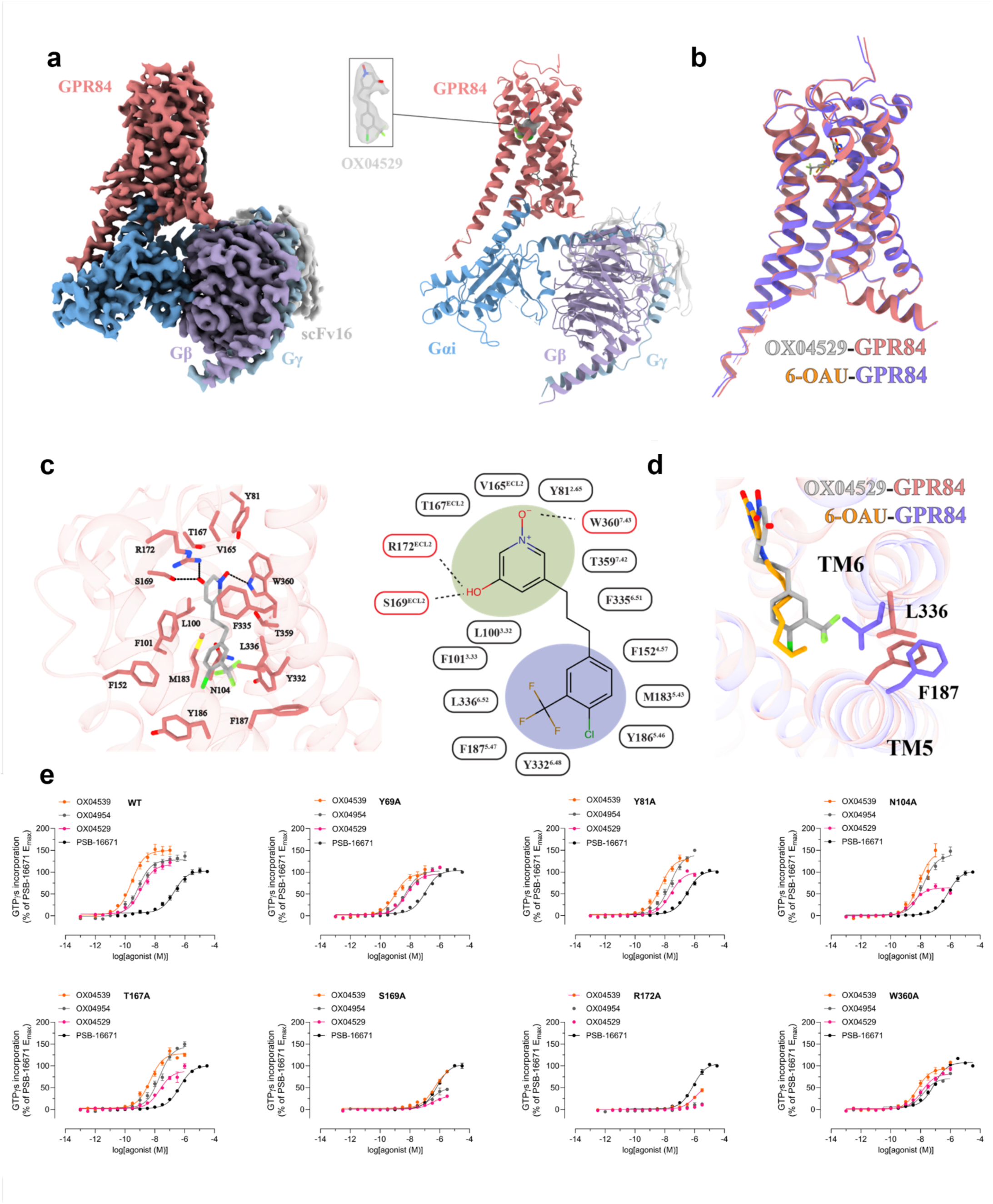
Overall structure and binding mode of the OX04529-GPR84-G_i_ complex. **a**, The left and right panels show the cryo-EM density map and the overall structure of the OX04529-GPR84- G_i_ complex, respectively. The chemical structure of OX04529 and the cryo-EM density of OX04529 contoured are shown in the middle. **b**, Superimposition of the OX04529-GPR84-G_i_ complex with the 6-OAU-GPR84-G_i_ complex. **c**, Details of the interactions between OX04529 and GPR84. **d**, F187^5.47^ and L336^6.52^ of GPR84 assume distinct conformations in these two structures. **e**, Mutagenesis studies to verify key interaction residues for OX04529 and OX04539 using [^35^S]GTPγS incorporation assays. Data represent means ± S.E.M. from at least three independent experiments (see **Extended Data Table 2** for quantitative details).

The transmembrane domain (TMs) and extracellular loops (ECLs) of GPR84 exhibit high structural similarity between the OX04529- and 6-OAU-bound structures, with an RMSD of 0.536 Å for the Cα atoms **(Fig. 2b)**. The two ligands also adopt highly similar binding poses **(Fig. 2b)**. In both structures, the polar head group of the ligand engages in extensive hydrogen-bonding interactions with S169 and R172 in ECL2, as well as and W360^7.43^ (superscripts represent Ballesteros-Weinstein numbering^24^) of GPR84 **(Fig. 2c)**. Although the position of OX04529 in GPR84 could be modelled with a high degree of certainty, the precise orientation of the 3-hydroxypyridine *N*-oxide ring system is ambiguous. The ring can be modelled to engage Arg172 and Ser169 (**Extended Data Fig. 3a**, b) with either its hydroxyl group or its pyridine *N*-oxide oxygen. The measured pKa of the hydroxyl group at 6.26 suggests a tendency for the deprotonated form but this may not be reflective of the binding site environment. However, that the DL-175 analog containing only the pyridine *N*-oxide group^16^ does not display reduced potency at an Arg172Ala mutation of GPR84^11^ suggests this group is most likely to interact with Trp360^7.43^. In addition to the polar interactions, the hydrophobic tail of OX04529 or 6-OAU is positioned within a hydrophobic sub-pocket formed by GPR84 residues F101^3.33^, F152^4.57^, Y186^5.46^, Y332^6.48^, F335^6.51^, and L336^6.52^ **(Fig. 2c)**. Despite the overall structural similarity, notable differences are observed within the binding pocket, particularly at the binding interface of the hydrophobic tail of the ligand. Compared to the 6-OAU-GPR84-G_i_ complex, TM5 and TM6 in the OX04529-bound structure exhibit a slight inward shift. As a result, the side chains of GPR84 residues F187^5.47^ and L336^6.52^ assume distinct conformations in these two structures **(Fig. 2d)**. Notably, overlay of other available GPR84- G_i_ structures bound with the synthetic agonist LY237 (**8J19**) or the endogenous ligand 3-hydroxy-lauric acid (**8J18**) indicated the altered positioning of Phe187^5.47^ and Leu336^6.52^ was only produced by OX04529 (**Extended Data Fig. 3c**). These structural differences may contribute to the biased signaling property associated with OX04529, which will be discussed later. Further inspection of the global structures revealed that key conserved motifs in class A GPCRs involved in G protein activation, including the toggle switch^25^, PIF motif^26^, DRY motif^27,28^, NPxxY motif^29^ and CWxP motif^30^, remained essentially unchanged across the GPR84 structures bound to OX04529, 6-OAU, LY237, and 3-hydroxy-lauric acid (**Extended Data Fig. 4**).

To investigate whether the observed interactions of OX04529 and OX04539 with orthosteric pocket residues in GPR84 were important for inducing potent G_i_ pathway activation, we mutated each of Tyr69^2.53^, Tyr81^2.65^, Asn104^3.36^, Thr167^ECL^^2^, Ser169^ECL^^2^, Arg172^ECL^^2^, and Trp360^7.43^ to Ala in the context of a GPR84- G_i2_α fusion protein^31^ and performed [^35^S]GTPγS binding studies to assess activation of the constructs in membranes of Flp-In T-REx 293 cells transfected to stably express each variant. To define that these mutations did not result in lack of expression or general misfolding of the receptor we performed initial studies using the GPR84 allosteric agonist, PSB-16671^11,31,32^, which binds to a yet to be identified location. PSB-16671 is able to promote activation of G_i_- but is biased and unable to recruit β-arrestins^11^. In the [^35^S]GTPγS binding assays potency of PSB-16671 was unchanged compared to wild type GPR84 at Tyr69^2.53^Ala, and Tyr81^2.65^Ala whilst a small but statistically significant reduction in potency was observed for the Thr167^ECL2^Ala mutant with a similarly small increase in potency at Trp360^7.43^Ala GPR84 (**Fig. 2e**). A 3-5 fold reduction in potency for PSB- 16671 was noted at both the Asn104^3.36^Ala and the Arg172^ECL2^Ala mutants (**Fig. 2e**, **Extended Data Table 1**), potentially reflecting the importance of these residues for the general structure of GPR84. By contrast, in addition to lacking detectable potency at Arg172^ECL2^Ala GPR84 (**Fig. 2e**), as previously reported for traditional GPR84 orthosteric agonists such as 6-OAU,^11^ OX04529 displayed some 10-fold reduced potency at the Ala mutants of Tyr81^2.65^ and Trp360^7.43^, some 300-fold reduced potency at the Ser169^ECL^^2^ Ala mutant and a more modest 5-fold reduction at Asn104^3.36^Ala GPR84 (**Fig. 2e**, **Extended Data Table 1**). Despite not forming an H-bond with Thr167^ECL^^2^ in the cryo-EM structure, OX04529 also displayed a greater than 10-fold reduction in potency at Thr167^ECL2^Ala GPR84 **(Extended Data Table 1**, **Fig. 2e**). Outcomes for OX04539 and also OX04954 were broadly similar, with almost 5000-fold reduction in potency at Arg172^ECL2^Ala, more than 1000-fold reduced potency at Ser169^ECL2^Ala and some 10-fold reduced potency at the Ala mutations of each of Tyr81^2.65^ Asn104^3.36^, Thr167^ECL^^2^ and Trp360^7.43^Ala GPR84 (**Extended Data Table 1**, **Fig. 2e**).

### Distinct ligand-induced conformations at TM5 and TM6 impact interactions between GPR84 and β- arrestins but not G protein

The key structural difference between the highly G protein-biased ligand OX04529 and the partially biased OX04954 lies solely in a single substituent: a CF₃ group versus CH₃ at the 3-position of the aryl ring. Structural comparison of the OX04529-GPR84-G_i_ cryo-EM structure with those containing reference agonists able to promote interactions with both G protein and arrestins (6-OAU, 3-hydroxy-lauric acid, or LY237) revealed that the CF₃ group of OX04529 is oriented toward transmembrane domains TM5 and TM6, causing notable conformational changes in the side-chains of Phe187^5.47^ and Leu336^6.52^ (**Fig. 3a and Extended Data Fig. 3c**). Specifically, the Leu336^6.52^ side-chain rotates toward the exterior of the binding pocket, while the phenyl group of Phe187^5.47^ shifts downward.

**Fig. 3.**
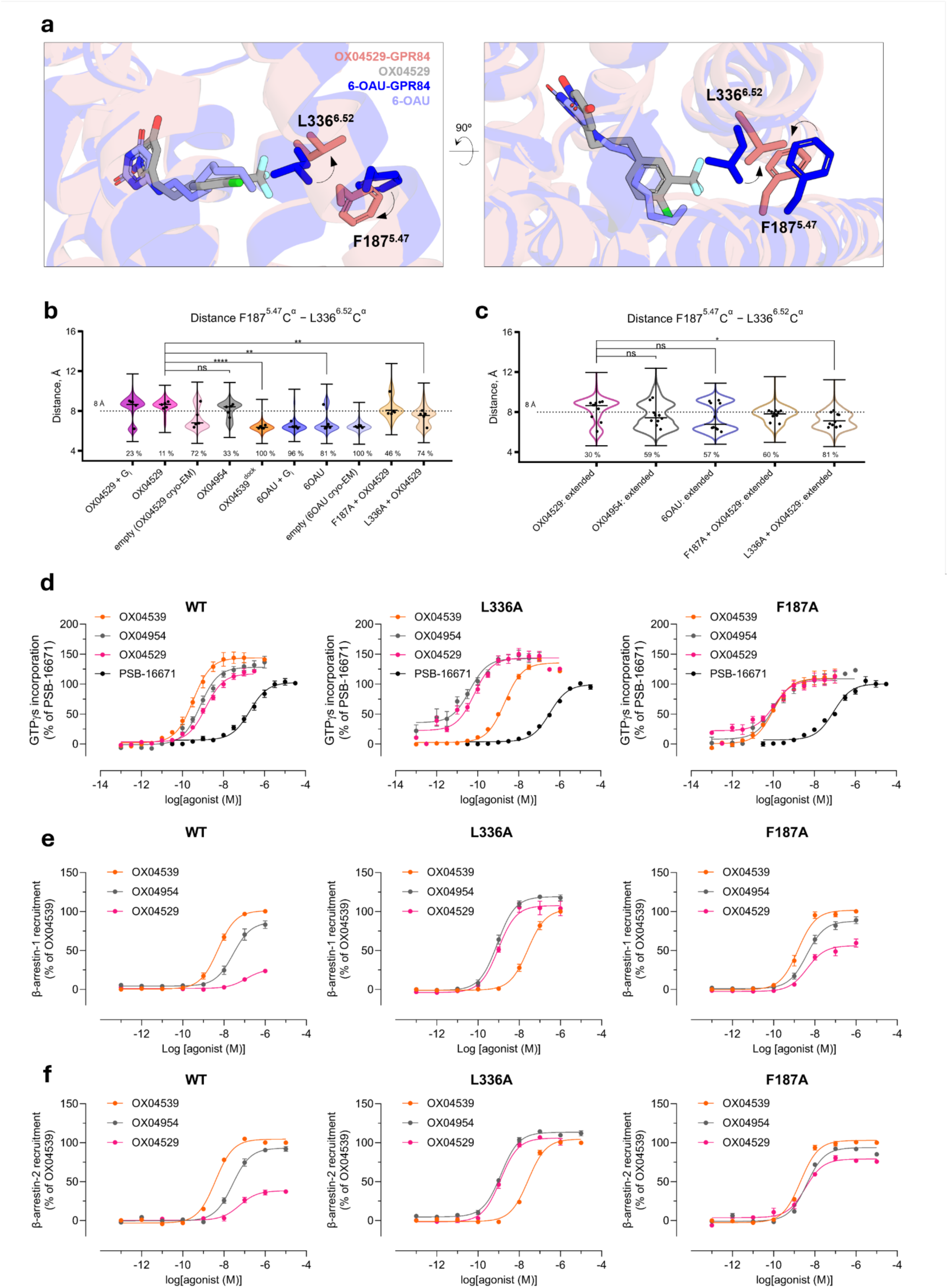
The conformation of Leu336^6.52^and Phe187^5.47^directly impacts β-arrestin recruitment and biased agonism. **a**, Leu336^6.52^and Phe187^5.47^ adopt a different conformation in the OX04529-GPR84-G_i_ complex compared to the 6-OAU-GPR84-G_i_ complex. **b,** Violin plots show the distribution of Cα-Cα distances between Phe187^5.47^ and Leu336^6.52^ measured across five replicate 1-μs MD simulation trajectories. Systems include GPR84 bound to G protein-biased ligand OX04529 (with and without G protein), partially biased OX04954, reference agonists (6-OAU and OX04539 computationally docked into the OX04529 cryo-EM structure, along with 6-OAU native cryo-EM structure), ligand-free receptor states derived from cryo-EM structures, and point mutants (Phe187^5.47^Ala and Leu336^6.52^Ala). The dashed line at 8 Å demarcates the threshold separating compact and extended conformational states. Percentages indicate the proportion of simulation frames with distances below this threshold, demonstrating that OX04529 uniquely stabilizes the extended conformation while β-arrestin recruiting agonists favor the compact state. **c**, Violin plots show the distribution of Cα-Cα distances between Phe187^5.47^ and Leu336^6.52^ measured across ten replicate 2-μs MD simulation trajectories using the CHARMM36m force field. Systems include GPR84 bound to G protein-biased ligand OX04529, partially biased OX04954, β-arrestin recruiting agonists (6- OAU), and point mutants (Phe187^5.47^Ala-OX04529 and Leu336^6.52^Ala-OX04529). The dashed line at 8 Å demarcates the threshold separating compact and extended conformational states. Percentages indicate the proportion of simulation frames with distances below this threshold. Extended sampling confirms that OX04529 uniquely stabilizes the separated conformation (30% below threshold) while β-arrestin recruiting agonists predominantly maintain the compact state (57-60% below threshold). **d**, [^35^S]GTPγS incorporation, (**e**) β-arrestin-1 and (**f**) β-arrestin-2 recruitment to GPR84 wild type (WT), Leu336^6.52^Ala and Phe187^5.47^Ala mutants in response to OX04539, OX04954 and OX04529. In **d** effects of the allosteric agonist PSB-16671 are also shown. As this ligand does not promote β-arrestin recruitment^11,31,32^ it was not used in panels (**e**) or (**f**). Data represent means ± S.E.M. from at least three independent biological replicates.

MD simulations of GPR84 structures with various ligands (OX04529, 6-OAU, OX04539, OX04954), with and without G_i_ protein, confirmed the structural observations. Since the 3-hydroxypyridine *N*-oxide ring orientation remained ambiguous (**Extended Data Fig. 3a**, b), we tested both possible conformations in preliminary simulations (five repeats of 1 μs) (**Fig. 3b, Extended Data Fig. 5**). Extended simulations (ten repeats of 2 μs) using different force fields and protonation states validated these findings (**Fig. 3c**). OX04529 uniquely increased the Phe187^5.47^–Leu336^6.52^ distance (∼8.5 Å vs. ∼6 Å for 6-OAU) and induced outward Leu336^6.52^ sidechain rotation (**Extended Data Fig. 6**). These effects were independent of headgroup orientation and G protein presence. The reference agonist (6-OAU) showed no such conformational change, and computational replacement of OX04529 with OX04539 or OX04954 reversed these effects. Simulations of Phe187^5.47^Ala and Leu336^6.52^Ala mutants similarly abolished the OX04529-induced conformational changes, confirming that CF₃ steric effects on these residues prevent β-arrestin recruitment.

To experimentally validate these computational insights, we conducted a series of experiments using a wild type GPR84-G_i2_α fusion protein or its Leu336^6.52^Ala or Phe187^5.47^Ala variants, once more stably expressed in Flp-In T-REx 293 cells. As for the point mutants described above, we initially assessed response to PSB-16671 in [^35^S]GTPγS binding studies (**Fig. 3d**). Whilst potency for the allosteric agonist was unchanged at Leu336^6.52^Ala, this ligand displayed slightly higher potency at Phe187^5.47^Ala GPR84 (**Fig. 3d**,

**Extended Data Table 3**). Notably OX04529 displayed more than 10-fold greater potency at both these variants than at wild type, with EC_50_ values close to 100 pM (**Fig. 3d**, **Extended Data Table 3**). Whilst both OX04954 and OX04529 also displayed substantially higher potency at Phe187^5.47^Ala GPR84 (**Extended Data Table 3**), OX04539 showed a 10-fold reduction in potency at Leu336^6.52^Ala GPR84 compared to wild type (**Extended Data Table 3**), resulting in a reversal of rank-order of potency for OX04539 relative to OX04529 and OX04954 at wild type GPR84. Notably, the enhanced potency of OX04529 at Leu336^6.52^Ala GPR84 was shown to reflect higher affinity (K_i_) based on its ability to compete with the orthosteric antagonist [^3^H]140 to bind to this variant (**Extended Data Fig. 7**).

Although in [^35^S]GTPγS G_i_-activation assays OX04529, OX4539 and OX04954 were equally efficacious at each of Phe187^5.47^Ala, Leu336^6.52^Ala GPR84 and wild type GPR84 (**Fig. 3d**) this was not the case when interactions with β-arrestins were assessed. Remarkably, although as noted earlier, OX04529 was a weak partial agonist at wild type GPR84, this ligand became a potent (**Extended Data Table 4**), full agonist in promoting interactions with both β-arrestin-1 and β-arrestin-2 in HEK293T cells at Leu336^6.52^Ala GPR84 (**Fig. 3e**, **f**). OX04529 also displayed substantial efficacy at Phe187^5.47^Ala GPR84 (**Fig. 3e**, **f**) with a more pronounced effect when using β-arrestin-2 (**Fig. 3e**, **f**).

### Alterations in OX04529 recruitment of arrestins is mirrored by phosphorylation of residues Thr263^ICL^^3^ and Thr264^ICL^^3^

To extend the observations of differences in GPR84 ligand-induced interactions with β-arrestins, we focused on the phosphorylation of residues Thr263^ICL^^3^ and Thr264^ICL^^3^ that are located in the receptor third intracellular loop. Reference orthosteric agonists promote phosphorylation of these residues, whilst mutation of these residues to Ala greatly diminishes the ability of such agonists to promote recruitment of β-arrestins^11^. Using a 96-well plate magnetic bead-based capture assay and an antiserum that detects these alterations specifically we observed that OX04954 and OX04539 promoted phosphorylation of these residues in wild type GPR84 with similar high potency and efficacy (**Extended Data Fig. 8a**). By contrast, OX04529 displayed weak efficacy (**Extended Data Fig. 8a)**. We extended these studies using immunoblots on lysates of cells stably expressing eYFP-tagged forms of wild type, Leu336^6.52^Ala and Phe187^5.47^Ala GPR84. In such immunoblots phosphorylation of Thr263^ICL^^3^ and Thr264^ICL^^3^ was barely detectable in the absence of a ligand (**Extended Data Fig. 8b)**. However, at 1 μM both OX04954 and OX04539 promoted phosphorylation of wild type GPR84, whilst OX04529 was ineffective (**Extended Data Fig. 8b**). By contrast at Phe187^5.47^Ala GPR84 a partial effect of OX04529 was observed compared to OX04954 and OX04539, whilst at Leu336^6.52^Ala GPR84 OX04529 became a full agonist in this assay (**Extended Data Fig. 8b**).

### OX04529 induces a distinct GPR84 conformation

Our MD simulations revealed that the G protein-biased ligand OX04529 induces distinctive conformational changes in GPR84 compared to reference agonists. The steric hindrance created by the CF_3_ group of OX04529 caused a notable outward shift of the extracellular tip of TM6 relative to OX04539 and 6- OAU (**Fig. 4a, Extended Data Fig. 9**). This movement was particularly pronounced in extended simulations, where TM6 underwent 20-30° rotations compared to the cryo-EM structures in 3 out of 10 simulation replicates (**Fig. 4b, Extended Data Fig. 9**)—a conformational change never observed with arrestin-recruiting ligands.

**Fig. 4.**
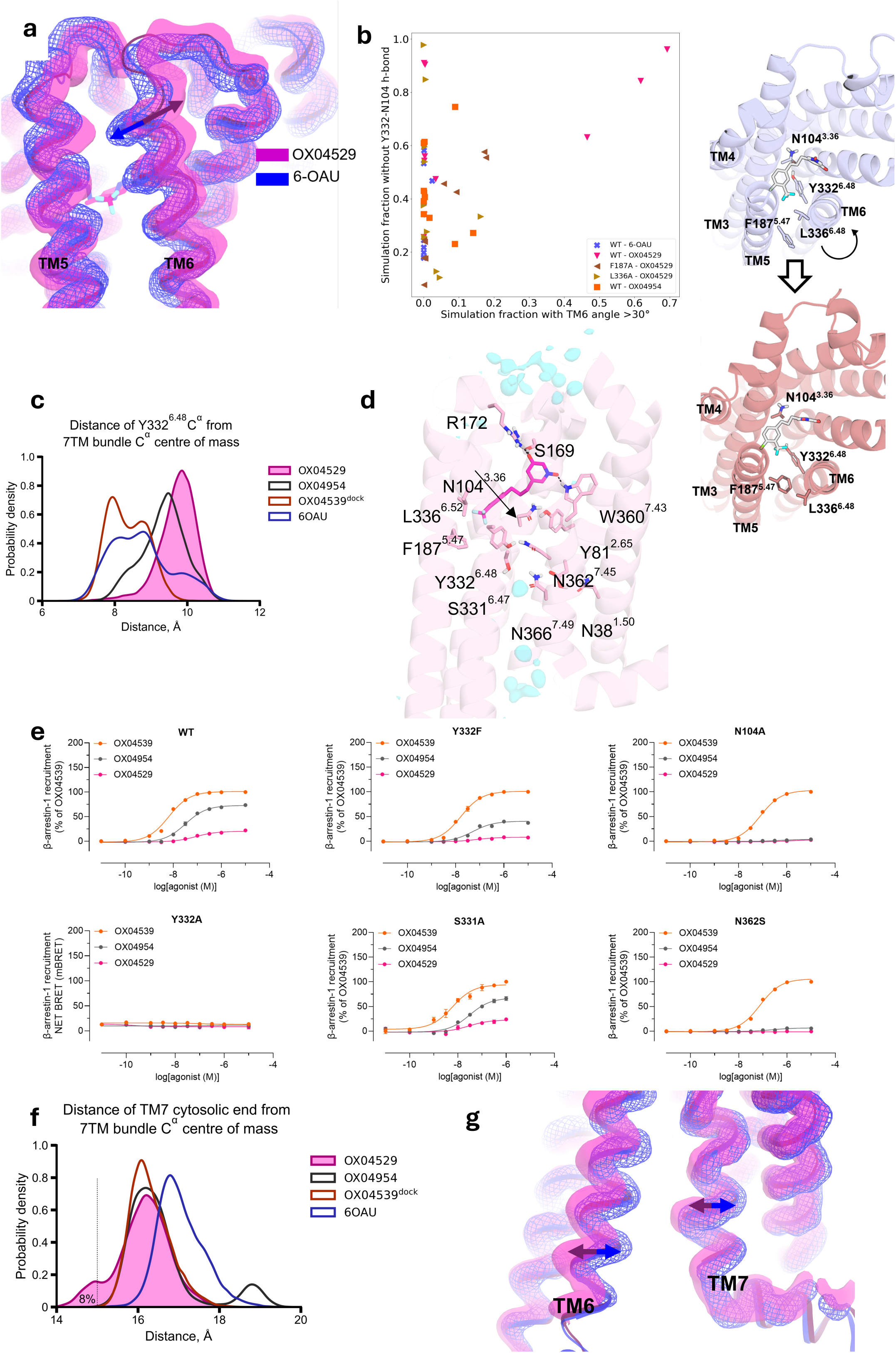
G protein-biased agonist OX04529 favors distinct H-bond connections between TMDs. **a**, The G protein-biased OX04529-bound structure (magenta) exhibits a distinctive outward shift of TM6 compared to the β-arrestin recruiting agonist 6-OAU complex (blue). Cartoon representations of the cryo-EM structures are overlaid with semi-transparent surfaces (solid for OX04529, mesh for 6-OAU) showing the spatial occupancy of protein backbone atoms (N, Cα, C) observed in >75% of molecular dynamics simulation frames. The arrows indicate the direction of TM6 movement induced by each ligand. **b**, Scatter plot illustrating the correlation of the simulation fraction with a TM6 angle rotation greater than 30 degrees (see Methods for definition) versus the loss of the Tyr332-Asn104 hydrogen bond as observed in 10 x 2 us simulations for WT-6-OAU (blue crosses), WT-OX04529 (downward red triangle), Phe187Ala- OX04529 (left brown triangle), Leu336Ala-OX04529 (right tan triangle) and WT-OX04954 (orange square) systems. Top, and separated out, view highlighting the significant rotation of TM6 observed in OX04529 simulations. Note the breaking of the hydrogen bond between Tyr332 and Asn104 in these simulations. **c,** Density distribution showing the distance between the Cα atom of Tyr332^6.48^ and the center of mass of the seven-transmembrane bundle across molecular dynamics simulations. The G protein-biased ligand OX04529 (magenta) stabilizes Tyr332^6.48^ in a significantly outward-displaced position (greater distance from bundle center) compared to the reference agonists OX04539 (orange) and 6-OAU (blue), with OX04954 (black) showing an intermediate distribution. This displacement reflects the disruption of the conserved toggle-switch network critical for β-arrestin recruitment. **d.** Representative snapshots from molecular dynamics simulations showing the binding poses and key interactions of G protein-biased OX04529 (magenta) in the GPR84 orthosteric pocket. Key residues involved in ligand recognition and signal transduction are labeled. Transparent cyan surface shows water density. **e**, β-arrestin-1 recruitment to GPR84 wild type (WT) and mutations (Tyr332^6.48^Phe, Asn104^3.36^Ala, Tyr332^6.48^Ala, Ser331^6.47^Ala, Asn362^7.45^Ala) with OX04539, OX04954 and OX04529. Data represent mean ± S.E.M. from at least three biologically independent experiments. **f,** Density distribution showing the distance between the TM7 cytosolic terminus and the center of mass of the seven-transmembrane bundle across molecular dynamics simulations. The G protein-biased ligand OX04529 (magenta) stabilizes TM7 in a more inward position (shorter distance from bundle center) compared to the reference agonists OX04539 (orange) and 6-OAU (blue), with OX04954 (black) displaying an intermediate distribution. This inward displacement of TM7, coupled with the outward movement of TM6, contributes to the intracellular interface remodeling that favors G protein coupling over β-arrestin recruitment. **g,** Conformational comparison of the intracellular portions of TM6 and TM7 in GPR84 bound to G protein- biased OX04529 (magenta) versus β-arrestin recruiting agonist 6-OAU (blue). Cartoon representations of the structures are overlaid with semi-transparent surfaces (solid for OX04529, mesh for 6-OAU) showing the spatial occupancy of protein backbone atoms (N, Cα, C) observed in >75% of molecular dynamics simulation frames. The arrows indicate the direction of TM6 and TM7 movements induced by each ligand.

The TM5-TM6 separation induced by OX04529 led to distinct conformational changes in Tyr332^6.48^, a critical ’toggle-switch’ residue within the conserved CWxP motif (SY^6.48^xP in GPR84) that is essential for Class A GPCR activation^33^ (**Fig. 4c**). In longer simulations, TM6 movement correlated with the disruption of the hydrogen bond between Asn104^3.36^ and Tyr332^6.48^ (**Fig. 4b, Extended Data Fig. 9**) and outward displacement of the top half of TM7 (**Extended Data Fig. 10**). This displacement caused Tyr332^6.48^ to move outward and downward, forming alternative hydrogen bonds with Asn104^3.36^, Asn362^7.45^, or Asn366^7.49^ of the conserved NP^7.50^xxY motif in different simulation systems (**Fig. 4d, Extended Data Fig. 9**). Additionally, we observed dynamic hydrogen bonding between Asn362^7.45^ and Ser331^6.47^ (**Fig. 4d**).

To test the functional importance of these interactions in ligand bias, we generated the mutations Tyr332^6.48^Ala (and Tyr332^6.48^Phe), Asn104^3.36^Ala, Asn362^7.45^Ser and Ser331^6.47^Ala. Conservative alteration of Tyr332^6.48^ to Phe preserved, and indeed slightly enhanced, G protein activation (**Extended Data Table 5**) while reducing β-arrestin recruitment in response to OX04954 and all but eliminated response to OX04529 compared to OX04539 (**Fig. 4e**, **Extended Data Table 6**). The more radical Tyr332^6.48^Ala alteration proved highly disruptive, severely impairing G protein activation (**Extended Data Table 5)** and eliminating β-arrestin recruitment for all compounds (**Fig. 4e**, **Extended Data Table 6**) meaning that it was impossible to reach conclusions based on this mutation. The Asn104^3.36^Ala mutation, despite reduced potency compared to wild type, preserved G protein activation for both OX04954 and OX04539 and, with reduced efficacy, for OX04529 (**Extended Data Table 1**). However, only the aliphatic-tailed OX04539 maintained β-arrestin recruitment, while the aromatic-tailed compounds, OX04954 and OX04539 completely lost this capacity (**Fig. 4e**, **Extended Data Table 6**). Interestingly, an Asn362^7.45^Ser mutation enhanced G protein activation for all compounds, with efficacies higher than at the wild-type receptor (**Extended Data Table 5)** yet nearly eliminated β-arrestin recruitment for the aromatic-tailed compounds **(Fig. 4e**, **Extended Data Table 6**). In contrast, the Ser331^6.47^Ala alteration minimally impacted the receptor signalling profile (**Fig. 4e**, **Extended Data Table 6**). These results indicate that hydrogen bonding involving Tyr332^6.48^, Asn104^3.36^ and Asn362^7.45^ is critical for β-arrestin recruitment, specifically for compounds with an aromatic ring in the non-polar tail.

Finally, analysis of intracellular interfaces reveals OX04529 distinctly reconfigures transmembrane helix positioning. Distance analysis confirms OX04529 shifts the cytosolic end of TM7 inward (**Fig 4f**) while displacing TM6 outward (**Fig. 4g, Extended Data** Figs. 9 and 10) This coordinated rearrangement of helices remodels the intracellular signaling interface in a manner that favors G protein coupling over β-arrestin recruitment.

### Rational design of a chemical library emphasizes the key criterion of steric demand that determines potency and efficacy for β-arrestin recruitment

To further investigate the role of steric occupancy in controlling ligand bias at GPR84 we conducted a more extensive structure–functional selectivity relationship (SFSR) study (**Fig. 5a**). We focused on the region of the GPR84 agonists interacting with TM5 and TM6 and their potential positioning in relation to Leu336^6.52^ (**Fig. 5b**), which, based on our structural, dynamic simulations and pharmacological insights, appeared to indirectly effect downstream polar contacts. We rationally substituted the aryl 3-position of the ligands with various functional groups beyond CH_3_ (OX04954) and CF_3_ (OX04529) to test the prediction that steric occupancy of the 3-substituent would correlate with β-arrestin recruitment. These included the derivative lacking a substituent at the 3-position (OX04540) as a baseline (**Fig. 5c**), a medium-sized hydrophilic CH_2_OH (OX04955) and halogens ranging from small F (OX04958) to medium-sized Cl (OX04957) and Br (OX04956). We also investigated the successive removal of fluorine atoms in CF_3_ (04529) to give CF_2_H (OX04959) and CH_2_F (OX04960), medium-sized unsaturated substituents (vinyl, OX04962 and ethylene, OX04963), and their saturated counterpart (ethyl, OX04961). Additionally, we explored increasing steric occupancy by increased branching in the substituent at the 3-position including isopropyl (OX04964), cyclopropyl (OX04965) and 1-methyl cyclopropyl (OX04966) (**Fig. 5c**). This diverse set of substituents allowed us to systematically investigate the effects of steric occupancy and van der Waals volume as well as hydrophobicity and electronic properties, on ligand bias.

**Fig. 5.**
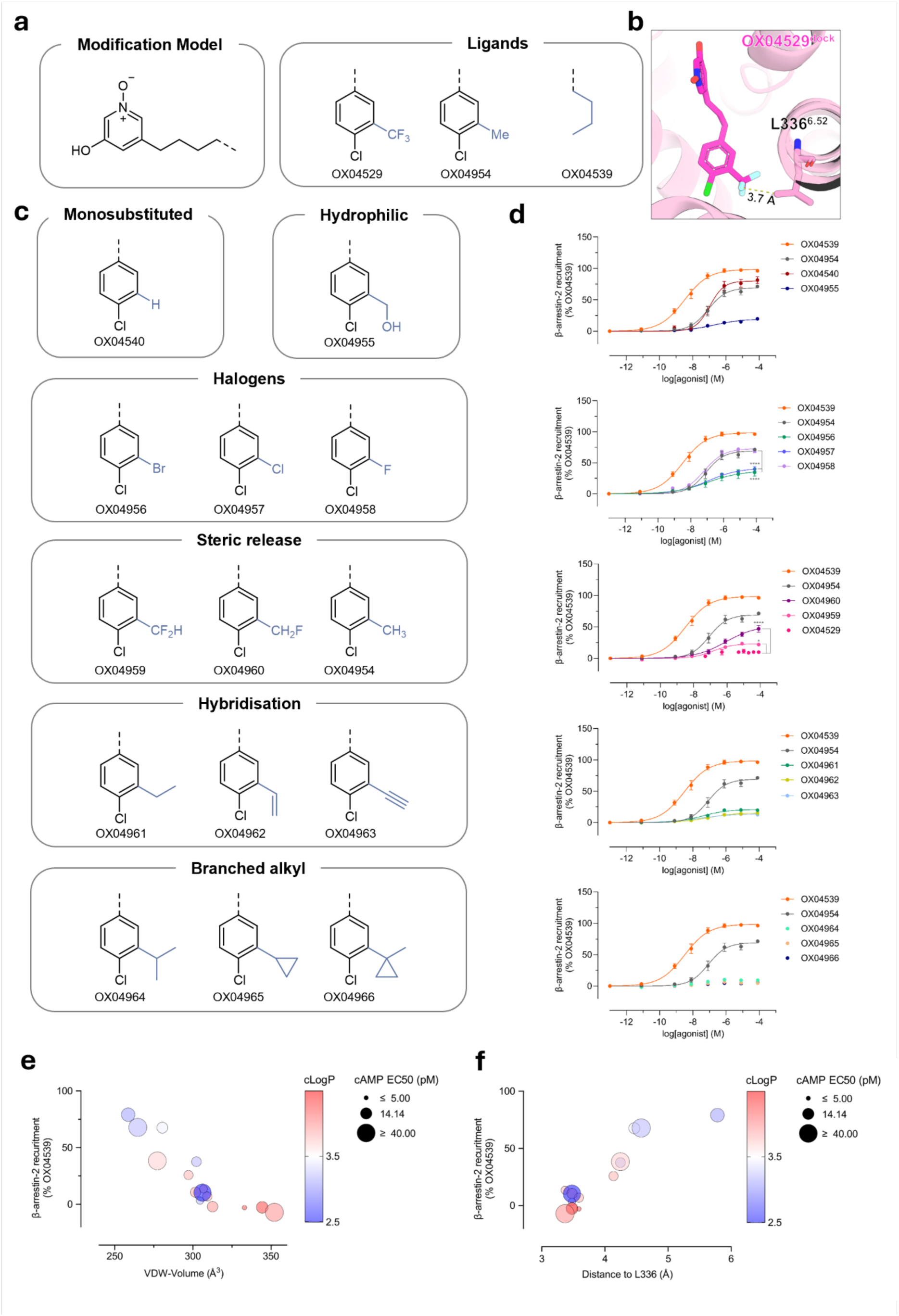
Rational design of a molecular library shows steric-dependent β-arrestin-2 recruitment efficacy. **a**, Chemical structures of OX04529 derivatives (left) and the rhs of the original agonists (right). **b**, Measurement of interatomic distance between the most distal heavy atoms of the ligand aryl 3-substituent and the side chain of Leu336^6.52^ taken from the cryo-EM structure of the OX04529-GPR84 complex. **c**, Design of a small library by replacing the 3-substituted CF_3_ within OX04529. **d**, β-arrestin-2 recruitment of small molecule library on GPR84. Data are means ± S.E.M. from at least three independent biological experiments, each performed in technical triplicate. **Extended Data Table 7** shows values for all panels and full data for all OX04529 analogs. **e**,**f**, Correlation plots of β-arrestin-2 maximum efficacy, Van der Waals (VDW) volume, distance from the most distal heavy (non H) atom of the ligand arene 3-substituent to Leu336, cLogP and cAMP activity. β- arrestin-2 maximum efficacy and VDW-volume are negatively correlated (**e**) while β-arrestin-2 maximum efficacy and the shortest distance between the most distal heavy atoms on the ligand and Leu336^6.52^ are positively correlated (**f**). **Extended Data Fig. 11** provides a full correlation map of ligand properties with β- arrestin efficacy, cAMP potency and efficacy well as ligand identity, docking and interatomic distance measurements.

All ligands tested behaved as full agonists in the cAMP inhibition assay conducted in CHO-K1 cells stably expressing GPR84 (**Extended Data Table 7**). Most ligands showed comparable potency (EC_50_ < 50 pM) in this assay apart from the 1-methyl cyclopropyl derivative OX04966, which showed significantly decreased potency compared to OX04529, suggesting that although the pocket can accommodate bulkier, branched substituents at this position through outward movement of Leu336^6.52^ and Phe187^5.47^, there is an intrinsic steric limitation where bond rotation can no longer sufficiently accommodate the ligand in the orthosteric binding site. By contrast the ligands displayed a diverse range of efficacy in β-arrestin-2 interaction studies (**Fig. 5d**, **Extended Data Table 7**).

Analysis of the correlation between a range of physicochemical parameters and measured cAMP inhibition and β-arrestin-2 recruitment revealed striking patterns (**Fig. 5e**, **f**, **Extended Data Fig. 11**). β-arrestin-2 efficacy showed a strong negative correlation with the calculated van der Waals volume of the ligand but not with ligand lipophilicity (**Fig. 5e**). Extending the carbon chain in the 3-substituent beyond methyl was found to be sufficient to reduce efficacy in β-arrestin-2 recruitment, where ethyl (OX04961), vinyl (OX04962), alkynyl (OX04954) and branched alkyl derivatives OX04964, OX04965 and OX04966 all showed no significant change in efficacy in β-arrestin-2 recruitment compared to OX04929. Notably, there was no apparent correlation with electronegativity or electron withdrawing capacity of the substituents (**Extended Data Fig. 11**). Consistent with these observations the halogen-substituted ligands (OX04956 (Br), OX04957 (Cl) and OX04958 (F)) gave a trend of F > Cl ≈ Br in efficacy in β-arrestin-2 recruitment assays, whereas successive replacement of the fluorine substituents within the trifluoromethyl group of OX04529 to yield OX04959 (CHF_2_), and OX04960 (CH_2_F) increased significantly the efficacy in β-arrestin-2 recruitment assays and showed a trend of CH_3_ > CH_2_F > CHF_2_ > CF_3_ (**Fig. 5e**). Whilst displaying reducing potency in [^35^S]GTPγS binding assays compared with OX04529, OX04959 (CHF_2_), and OX04960 (CH_2_F) (**Extended Data Table 3**) these ligands did not display reduced efficacy in this assay of G_i_-activation.

Furthermore, when measuring the interatomic distance between the most distal heavy atom of the aryl 3-substituent and of the Leu336^6.52^ side chain from the OX04529-GPR84 cryo-EM structure and comparing this with the measured interatomic distance taken from docking models of a selection of ligands, we observed a strong correlation between interatomic distance and β-arrestin-2 recruitment efficacy (**Fig. 5f**). Taken together, this strongly suggests that steric factors, rather than electronic effects or lipophilicity are dominant in determining β-arrestin-2 recruitment and hence the extent of ligand bias.

## Discussion

Functional selectivity has been observed in many GPCRs, including angiotensin^8,33^, adrenergic^34^, dopamine^35^, opioid^6,7,36–39^, cannabinoid^40^ and other receptors^41–44^. However, the multiple distinct binding sites at the receptors and the intrinsic structural flexibility of the receptors have made the interpretation of the molecular mechanism challenging and limited our ability to design ligands with predictable levels of signaling bias. We previously described a highly potent G protein-biased agonist of GPR84 (OX04529). During our SAR analysis to identify OX04529 we synthesized OX04954 and OX04539, which displayed markedly different efficacy in β-arrestin-2 recruitment while maintaining potency and efficacy through G_i_ signaling. Remarkably this was achieved with a relatively minor structural change in the hydrophobic tail of the parent G protein-biased ligand, OX04529. Each ligand exhibited comparable low nanomolar affinity in a radioligand binding assay and, intriguingly, comparable association and dissociation rates, demonstrating that ligand bias in this case was not driven by differences in binding kinetics. We solved a cryo-EM structure of OX04529 in complex with GPR84-G_i_: OX04529 was found to adopt a binding pose very similar to the reference agonist 6-OAU. Mutagenesis studies showed OX04529 activated GPR84 in a manner similar to 6-OAU by forming key hydrogen bonds with Ser169^ECL^^2^, Arg172^ECL^^2^ and Trp360^7.43^. The only observed significant differences from other described structures of ligands bound to GPR84-G_i_ complexes arose from the orientation of the trifluoromethyl group at the 3-position of the arene on OX04529 which was directed toward Leu336^6.52^, causing a conformational change in the side chains of both Leu336^6.52^ and Phe187^5.47^. Mutagenesis studies confirmed the functional importance of these conformational changes, as Leu336^6.52^Ala and Phe187^5.47^Ala mutations restored β-arrestin recruitment for OX04529 while preserving G protein activation.

MD simulations revealed OX04529 stabilizes a unique tilted state with distinctive TM6 twisting at its extracellular tip, separating TM6 from TM5 at the binding site. This rearrangement alters the conformation of Tyr332^6.48^, the ’toggle-switch’ residue of the conserved CWxP motif critical for GPCR activation^45^, disrupting its polar contacts. Mutagenesis studies confirmed these computational findings. Similar toggle-switch disruption has been linked to biased signaling in other GPCRs^8,38,39^. The functional consequence is impaired phosphorylation of Thr263^ICL^^3^ and Thr264^ICL^^3^ in GPR84, a characteristic shared with the G protein-biased agonist DL-175. Leu336^6.52^Ala or Phe187^5.47^Ala mutations restored this phosphorylation with OX04529, consistent with recovered β-arrestin recruitment. As these phosphorylation events are GRK2/3-mediated, biased ligands appear to position ICL3 in a configuration preventing efficient kinase access. This effect is not due to differences in G protein activation, as all ligands showed equivalent G activation efficacy. This is relevant because GRK2/3 recruitment to the plasma membrane is dependent on the availability of β/γ complex released from activated G protein heterotrimers^46^ and thus variation in G protein activation could have offered a different explanation for our findings

Based on these findings, we proposed that steric occupancy of the ligand 3-substituent and its van der Waals volume should inversely correlate with β-arrestin recruitment. We tested this by synthesizing compounds with systematic modifications to the aryl 3-substituent of OX04529, varying hydrophobicity, electronic properties, and steric profiles. All compounds were full agonists in cAMP inhibition assays with high potency (except the bulky OX04966). Consistent with our model, β-arrestin-2 recruitment efficacy showed strong negative correlation with van der Waals volume and positive correlation with the distance between the 3-substituent and Leu336^6.52^, confirming that molecular size and steric occupancy primarily determine ligand bias at GPR84. In summary, we elucidate a molecular mechanism for designing G protein-biased GPR84 agonists through functional, structural, and dynamic analyses. The functional selectivity can be precisely controlled by tuning steric occupancy near Leu336^6.52^, which alters the conformations of Leu336^6.52^ and Phe187^5.47^, disrupts Tyr332^6.48^ polar contacts, and consequently impairs ICL3 phosphorylation and β- arrestin recruitment. These findings provide valuable insights for the design of new ligands targeting GPR84 and may also be applicable to other class A GPCRs.

## Materials and Methods

6-n-Octylaminouracil (6-OAU), 3,3’,5,5’-tetramethylbenzidine (TMB) and horseradish peroxidase (HRP)- sheep anti-rabbit IgG were from Merck. di(5,7-difluoro-1H-indole-3-yl)methane (PSB-16671) was synthesized by DC Chemicals (Shanghai, China). [^35^S]GTPγS, GF/C glassfibre filter-bottom 96-well microplates, and Ultima Gold^TM^ XR were from Revvity. Compound 837 (3-((5,6-bis(4-methoxyphenyl)-1,2,4- triazin-3-yl)methyl)-1H-indole) was synthesized as described previously^21^. [^3^H]140 (3-((5,6-diphenyl-1,2,4- triazin-3-yl)methyl)-1H-indole) (40 Ci/mmol) was produced by Pharmaron (Cardiff, UK) and synthesized as described previously^21^. Cell culture reagents, NuPAGE Novex 4% to 12% Bis-Tris Gels, NuPAGE MOPS SDS running buffer, and bicinchoninic acid (BCA) protein assay kit were from Thermo Fisher Scientific. All molecular biology enzymes and reagents were from Promega. Polyethylenimine (PEI) (linear poly(vinyl alcohol) [MW 25,000]) was from Polysciences (Warrington, PA). PhosSTOP™ and cOmplete™ protease inhibitors tablets were from Roche Diagnostics. The rabbit phospho-site-specific GPR84 antiserum pThr263/pThr264-GPR84 (7TM0120B) was developed in collaboration with 7TMAntibodies GmbH. IRDye 800CW donkey anti-rabbit IgG was from LI-COR Biosciences and the sheep GFP antiserum was generated in-house. GFP-Trap® Agarose and GFP-Trap® Magnetic Agarose beads were from Proteintech.

### Protein complex expression and purification

The wild-type human GPR84 gene was synthesized and cloned into the pFastBac vector (Fisher), incorporating a bovine prolactin signal peptide, an N-terminal Flag-tag, and a C-terminal His₈-tag. To enhance complex stability, the NanoBiT tethering strategy was employed by fusing the LgBiT subunit^47^ to the C-terminus of GPR84 after R387 via a 15-amino acid flexible linker (GSSGGGGSGGGGSSG). A dominant negative human Gα_i1_ (DNGα_i1_) containing four mutations (S47N, G203A, E245A, A326S) was cloned into the pFastBac vector. Human Gβ_1_ was fused with an N-terminal His_6_-tag and a C-terminal HiBiT subunit connected with a 15-amino acid linker, was cloned into pFastBac dual vector^48^ together with human Gγ_2_.

The expression and purification of scFv16 were performed as previously described^49^. Briefly, scFv16 was expressed in High Five cells and purified by nickel affinity chromatography. The protein was further purified by size-exclusion chromatography using a Superdex 200 Increase 10/300 GL column (GE Healthcare). Monomeric peak fractions were pooled, concentrated, and stored at -80 °C until use.

GPR84, DNGα_i1_ and Gβ_1_γ_2_ were co-expressed in Sf9 insect cells using Bac-to-Bac baculovirus expression system. Cells were infected with the three respective viruses at a 1:1:1 ratio. After infection for 48 h at 27 °C, cell pellets were harvested and stored at -80 °C until use. Cell pellets were thawed in lysis buffer containing 20 mM HEPES, pH 7.5, 50 mM NaCl, 10 mM MgCl_2_, 5 mM CaCl_2,_ 2.5 μg/ml leupeptin, 300 μg/ml benzamidine. To facilitate complex formation, 1 μM OX04529, 25 mU/ml Apyrase (NEB), and 100 μM TCEP was added, and the mixture was incubated at room temperature for 2 h. Membranes were isolated by centrifugation at 30,700 × g for 30 min and resuspended in solubilization buffer containing 20 mM HEPES, pH7.5, 100 mM NaCl, 0.5% (w/v) lauryl maltose neopentyl glycol (LMNG, Anatrace), 0.1% (w/v) cholesteryl hemisuccinate (CHS, Anatrace), 10% (v/v) glycerol, 10 mM MgCl_2_, 5 mM CaCl_2_, 12.5 mU/ml Apyrase, 1 µM OX04529, 2.5 μg/ml leupeptin, 300 μg/ml benzamidine, 100 µM TECP for 2h at 4 °C. Insoluble material was removed by centrifugation at 38,900 × g for 45 min. The supernatant was incubated with Ni-affinity resin at 4 °C for 2 h. The resin was washed with buffer A (20 mM HEPES pH 7.5, 100 mM NaCl, 0.05% (w/v) LMNG, 0.01% (w/v) CHS, 20 mM imidazole, 1 μM OX04529, 2.5 μg/ml leupeptin, 300 μg/ml benzamidine, and 100 μM TCEP), and the complex was eluted with buffer A supplemented with 400 mM imidazole. The eluate was supplemented with 2 mM CaCl₂ and incubated overnight at 4 °C with anti-Flag M1 antibody resin. The Flag resin was washed with 10 column volumes of buffer A containing 2 mM CaCl₂. Then the complex was eluted by 3.5 column volumes of buffer A containing 5 mM EDTA and 200 μg/ml FLAG peptide. The eluted complex was concentrated using 100 kDa molecular weight cut-off concentrators (Millipore) and mixed with purified scFv16 at a 1.3:1 molar ratio. The sample was then loaded onto a Superdex 200 Increase 10/300 column (GE Healthcare) pre-equilibrated with buffer containing 20 mM HEPES pH 7.5, 100 mM NaCl, 0.00075% (w/v) LMNG, 0.00025% (w/v) GDN, 0.00015% (w/v) CHS, 1 µM OX04529 and 100 µM TECP.

Peak fractions of the complex were pooled and concentrated to 5 mg/ml for cryo-EM studies.

### Cryo-EM sample preparation and data acquisition

For cryo-EM grid preparation of the OX04529-GPR84-G_i_ complex, 3 μl of protein sample was applied to plasma-cleaned 1.2/1.3 UltrAuFoil 300 mesh grids and plunge-frozen in liquid ethane using a Vitrobot Mark IV (Thermo Fisher Scientific). Cryo-EM data were collected on a Titan Krios electron microscope operating at 300 kV, equipped with a Gatan K3 Summit direct electron detector and an energy filter. Micrographs were acquired in super-resolution mode using SerialEM software at a nominal magnification of 105,000×, yielding a calibrated pixel size of 0.414 Å. The defocus range was set between -1.0 and -1.8 μm. Each image stack was dose-fractionated into 52 frames with a total electron dose of 58 e^-^/Å². A total of 8,671 movies were collected for the OX04529-GPR84-G_i_ complex.

### Data processing, 3D reconstruction and modeling building

Image stacks were initially corrected for patch motion using cryoSPARC. Contrast transfer function (CTF) parameters were determined using the patch CTF estimation tool. For the OX04529-GPR84-G_i_ complex, a total of 6,052,532 particles were auto-picked and subsequently subjected to 2D classification to eliminate poor-quality particles. Following ab initio reconstruction and heterogeneous refinement, 218,137 particles underwent non-uniform refinement and local refinement, resulting in a map with a global resolution of 3.03 Å based on Fourier shell correlation (FSC) at 0.143. Local resolutions of density maps were estimated in cryoSPARC.

The model was built based on our previously reported structure of 6-OAU-GPR84-G_i_-scFv16 complex. The structure of GPR84-G_i_-scFv16 was used as initial model for model rebuilding and refinement against the electron microscopy map. The model was docked into the electron microscopy density map using Chimera^50^ followed by iterative manual adjustment and rebuilding in COOT^51^. Real space refinement was performed using Phenix programs^52^. The model statistics were validated using MolProbity. Structural figures were prepared in Chimera and PyMOL (https://pymol.org/2/). The final refinement statistics are provided in **Extended Data Table 2**.

### Plasmids and mutagenesis

The human GPR84 receptor with an N-terminal FLAG tag and either enhanced yellow fluorescent protein (eYFP), or Gα_i2_-Cys352Ile fused to the C terminus, was cloned into the pcDNA5/FRT/TO expression vector as described previously^53^. Site-directed mutagenesis to generate the point mutations described was performed according to the QuikChange method (Stratagene, Cheshire, UK).

### Cell culture, transfection and generation of cell lines

HEK293T cells were maintained in high-glucose Dulbecco’s modified Eagle’s medium (DMEM) without sodium pyruvate, supplemented with 10% (v/v) fetal bovine serum (FBS), 1% (v/v) penicillin/streptomycin mixture (100 U/ml), and 2 mg/ml normocin (Invivogen), at 37°C in a 5% CO_2_ humidified atmosphere.

Flp-In™ TREx™ 293 cells (Invitrogen) were maintained in high-glucose DMEM without sodium pyruvate, supplemented with 10% (v/v) FBS, 1% penicillin/streptomycin mixture (100 U/ml), 2 mg/ml normocin, and 5 μg/ml blasticidin (Invivogen) at 37°C in a 5% CO_2_ humidified atmosphere.

hGPR84-CHO-cAMP cells were maintained in Ham’s F-12 Nutrient Mix with 10% (v/v) FBS, 19 mM HEPES, 600 µg/mL G418 and 1% (v/v) penicillin-streptomycin mixture, at 37°C in a 5% CO_2_ humidified atmosphere.

hGPR84-CHO-*β*-arrestin-2 cells were maintained in Ham’s F-12 Nutrient Mix with 10% (v/v) FBS, 19 mM HEPES, 600 µg/mL G418, 300 µg/mL hygromycin B and 1% (v/v) penicillin-streptomycin mixture, at 37°C in a 5% CO_2_ humidified atmosphere.

To generate Flp-In™ TREx™ 293 cells able to express the various FLAG-GPR84-eYFP or FLAG-GPR84- Gα_i2_ receptor constructs in an inducible manner, cells were transfected with a mixture containing the desired cDNA in pcDNA5/FRT/TO vector and pOG44 vector (1:9) by using 1mg/ml PEI (MW-25000). Cells were plated until 60 to 80% confluent then transfected with 8 μg of required plasmid DNA and PEI (ratio 1:6 DNA/PEI), diluted in 150 mM NaCl, pH 7.4. After incubation at room temperature for 10 min, the mixture was added to cells. After 48 h, the medium was changed to medium supplemented with 200 μg/ml hygromycin B (Invivogen) to initiate the selection of stably transfected cells. After isolation of resistant cells, expression of the appropriate construct from the Flp-In™ TREx™ locus was induced by adding up to 100 ng/ml doxycycline for 24 h.

### Bioluminescence Resonance Energy Transfer (BRET) β-arrestin-1 and β-arrestin-2 recruitment assays

HEK293T cells were transiently co-transfected with FLAG-GPR84-eYFP constructs and either β-arrestin-1 or β-arrestin-2 fused to nano-luciferase, in a ratio of 100:1. As a control, cells were transfected with 100:1 ratio of empty plasmid and the appropriate arrestin isoform fused to nano-luciferase. After 24 h, cells were seeded at 50,000 cells per well in poly-D-lysine-coated white 96-well plates and incubated at 37°C overnight. After 24 h, cells were washed twice with Hank’s balanced salt solution (HBSS), pH 7.4, and 10 μl/well of the nano-luciferase substrate coelenterazine-h (Nanolight Technologies) was added to a final concentration of 5 µM. Cells were incubated in the dark for 10 min at 37°C, after which 10 μl/well of the agonist compound was added, and the cells were incubated for another 5 minutes. 15 min after coelenterazine-h addition, reading of emission signals were performed on a PHERAstar FS plate reader (BMG Labtech) at 475 and 535 nm, representing nano-luciferase and eYFP emission signals, respectively. The net bioluminescence resonance energy transfer (BRET) ratio (mBRET) was calculated as follows:

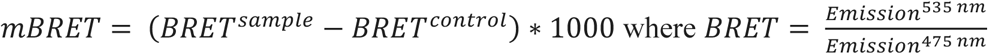

### [^35^S]GTPγS binding assays

Membranes were generated from Flp-In™ T-REx™ 293 cells following 100 ng/ml doxycycline treatment to induce receptor expression. Cells were washed with ice-cold phosphate-buffered saline (PBS), removed from dishes by scraping and centrifuged at 1,800 x *g* for 5 min at 4 °C. Pellets were resuspended in TE buffer (10mM Tris-HCl, 0.1 mM EDTA; pH 7.4) containing a protease inhibitor mixture and homogenized with a 5 ml hand-held homogenizer. Samples were centrifuged at 450 x *g* for 5 min at 4 °C and the supernatant was further centrifuged at 90,000 x *g* for 45 min at 4 °C. The resulting pellet was resuspended in TE buffer and protein content was assessed using a BCA protein assay kit.

Prepared membrane protein (3 µg per well) was incubated in assay buffer (20 mM HEPES, 5 mM MgCl_2_, 160 mM NaCl, 0.05 % fatty-acid-free bovine serum albumin; pH 7.4) containing the indicated ligand concentrations. The reaction was initiated by addition of [^35^S]GTPγS (100 nCi per reaction) with 1 µM GDP, and incubated at 30 °C for 60 min. The reaction was terminated by rapid vacuum filtration through GF/C glassfibre filter-bottom 96-well microplates using a UniFilter FilterMate Harvester (PerkinElmer). Unbound radioligand was removed from filters by three washes with ice-cold PBS. Ultima Gold^TM^ XR was added to dried filters, and [^35^S]GTPγS binding quantified by liquid scintillation spectroscopy.

### Competition binding assays

Competition binding experiments using approximate K_d_ concentrations of [^3^H]140 were conducted in binding buffer (20 mM HEPES, 5 mM MgCl_2_, 160 mM NaCl, 0.05% fatty-acid-free bovine serum albumin; pH 7.4) in a total assay volume of 500 μL in 96 deep-well blocks. Binding was initiated by the addition of 5 μg membrane protein generated from Flp-In^TM^ T-REx^TM^ cells induced to express the receptor construct of interest with 100 ng/ml doxycycline. Nonspecific binding of the radioligand was determined in the presence of 10 μM compound 837. All assays were performed at 25 °C for 1 h before termination by the addition of ice-cold phosphate-buffered saline (PBS) and vacuum filtration through GF/C glassfibre filter-bottom 96-well microplates using a UniFilter FilterMate Harvester (Revvity). Each well was washed three times with ice-cold PBS and filters were allowed to dry for 2–3 h before addition of 50 µl of Ultima Gold^TM^ XR. Radioactivity was quantified by liquid scintillation spectrometry.

### Determination of the “on” and “off” rates of unlabeled ligands through measurement of competition binding kinetics of [^3^H]140

The kinetic binding parameters of unlabeled ligands were determined through assessment of the binding kinetics of [^3^H]140.^21^ Here, binding buffer, [^3^H]140 (at an approximate K_d_ concentration of 1 nM), and three different concentrations of competitor (1-, 3-, and 10-fold the estimated respective K_i_ concentration) were added simultaneously to the wells of 96 deep-well plates. At the indicated time points, membrane protein was added, and tubes were gently agitated. Three different concentrations of competitor were assessed to ensure that the rate parameters calculated were independent of ligand concentration.

### pThr263/pThr264 GPR84 phosphorylation assay

Flp-In™ TREx™ 293 cells engineered to express FLAG-GPR84-eYFP were seeded onto poly-D-lysine-coated clear 96-well plates at a density of 5 x 10^4^ cells/well and grown overnight before treatment with 100 ng/ml doxycycline to induce receptor expression. After 24 h, cells were serum starved for 2 hours then stimulated with the indicated agonists, prepared in serum free media, for 5 min at 37 °C. After washing with ice-cold PBS, cells were lysed in 100 µl/well lysis buffer (150 mM NaCl, 50 mM Tris-HCl, 5 mM EDTA, 1% IGEPAL (Nonidet P-40), 0.5% deoxycholic acid, and 0.1% SDS; pH 7.4) supplemented with protease and phosphatase inhibitor cocktail on an orbital shaker for 1 hour at 4 °C. Lysates were cleared by centrifugation at 3700 × *g* for 20 min at 4 °C and 80 µl of supernatants transferred into 96-U-bottom-well assay plates (Greiner Bio-One). GFP-Trap® magnetic agarose beads (1 µl/well) were washed then diluted in lysis buffer to give a total volume of 20 µl of bead slurry per well before adding to supernatant. Assay plates were incubated on a microplate shaker at 700 rpm for 2 h at 4 °C then washed three times with 100 µl of PBS with 0.1% Tween-20 (PBS-T) under magnetic force using a handheld magnetic separation block (Milliplex, Millipore). After washing, anti-pThr263/pThr264 antiserum was diluted 1:1000 in PBS-T then 60 µl added to assay plate. The plates were then incubated overnight at 4 °C on a microplate shaker. After another wash cycle (3 x with PBS-T), anti-rabbit horseradish peroxide (HRP)-linked secondary antibody was added to PBS-T to a final dilution of 1:10,000 and incubated on a microplate shaker for 2 h at room temperature. After a further wash cycle, 100 µl of TMB substrate solution was added to each well and the plates were incubated for 15 min. The reaction was stopped by the addition of 100 μl of 0.3 M sulphuric acid. The assay plates were then placed on a magnetic separation block and 100 µl of the supernatant was transferred to a transparent 96-well F-bottom plate. The OD at 450 nm was then read on a PHERAstar FS plate reader (BMG Labtech).

### Receptor immunoprecipitation and immunoblotting

eYFP-linked receptor constructs were immunoprecipitated from 200 µl cell lysates using GFP-Trap® agarose beads (Proteintech) according to manufacturer’s instructions. Immunocomplexes were washed three times in washing buffer, resuspended in 100 μl 2 × SDS-PAGE sample buffer and incubated at 60 °C for 10 min. Following centrifugation at 2500*g* for 5 min, 20 μl of immunoprecipitated proteins was resolved by SDS-PAGE on 4% to 12% BisTris gels. After separation, the proteins were transferred electrophoretically onto nitrocellulose membrane, which was then blocked using 5% bovine serum albumin (BSA) in Tris-buffered saline (TBS, 50 mM Tris-Cl, 150 mM NaCl, pH 7.6) for 1 h at room temperature on an orbital shaker. The membrane was then incubated with appropriate primary antibody in 5% BSA in TBS supplemented with 0.1% Tween (TBS-T) overnight at 4 °C. Anti-pThr^263^/pThr^264^ antiserum was diluted 1:2000 and anti-GFP was diluted 1:10,000. Subsequently, the membrane was washed (3 × 10 min with TBS-T) and incubated for 2 h with anti-rabbit secondary antibody diluted 1:10,000 in 5% BSA in TBS-T. After washing (3 × 10 min with TBS-T), proteins were detected using an Odyssey imaging system (Licor) according to the manufacturer’s instructions.

### CHO-K1-Based cAMP Assay

CHO-K1 cells stably expressing human GPR84 (DiscoverX 95-0158C2) were seeded in 384-well plates at 15,000 cells per well in 20 µL of F12 medium supplemented with 10% fetal bovine serum (FBS), 19 mM HEPES, and 1% penicillin-streptomycin. After overnight incubation at 37 °C and 5% CO_2_, the medium was removed and replaced with PBS containing 0.1% BSA. Cells were stimulated for 30 min with forskolin (25 µM) and test compounds prepared in Dulbecco’s PBS (DPBS) with 0.1% BSA. Compounds were dissolved in DMSO and diluted appropriately. cAMP levels were quantified using the HitHunter® cAMP Assay for Small Molecules (DiscoverX 90-0075SM2) according to the manufacturer’s protocol. Luminescence was measured 18 h post-assay using a PHERAstar® FS plate reader (BMG Labtech). EC_50_ values were calculated using a four-parameter logistic model in GraphPad Prism (v9.5.0) and normalised to the response induced by forskolin (25 µM) and and the maximum effect of capric acid (100 μM).

### CHO-K1-Based *β*-arrestin-2 Recruitment Assay

CHO-K1 cells co-expressing ProLink-tagged hGPR84 and enzyme acceptor-tagged β-arrestin-2 (DiscoverX 93-0647C2) were seeded at 5,000 cells per well in 384-well plates and cultured overnight under the same conditions described above. Cells were stimulated with test ligands for 90 min. Compounds were prepared at 5× concentration in DPBS containing 0.1% BSA and 0.125% Tween-80 to minimise aggregation. β-arrestin recruitment was quantified using the PathHunter® β-Arrestin Assay (DiscoverX 93-0001) following the manufacturer’s instructions. Luminescence was recorded after 18 h using a PHERAstar® FS plate reader. EC_50_ values were calculated in GraphPad Prism using a four-parameter logistic model and normalised to the maximal response of the reference agonist OX04539 or 6-OAU.

### Computer Simulations

#### Molecular docking

Molecular docking was done via Maestro 2021-3 ^54^ GLIDE build 137 ^55–58^ in the “Standard Precision” mode with default settings, with a 20 Å inner box and 30 Å outer box centered at the orthosteric ligand geometric center.

#### Molecular docking using Molecular Operating Environment (MOE)

Docking studies were performed using the cryo-EM structure of the GPR84-OX04529 complex as the receptor model. Ligands were protonated, assigned partial atomic charges, and energy-minimised using the MMFF94x force field in MOE (v2019.0102). The protein structure was protonated at physiological pH (7.4) using Protonate3D, and hydrogen bonding networks were optimised. The receptor was treated as rigid during docking. Planar regions of ligands were constrained to retain conformational integrity. Docking was conducted within the orthosteric binding site defined by OX04529 in the cryo-EM structure. For each ligand, 5,000 poses were generated using the London dG scoring function, followed by refinement and rescoring with GBVI/WSA dG. The top 10 poses were retained, and the highest-ranked pose in which the 3-substituent was oriented toward Leu336 was selected. The shortest heavy-atom distance to the side chain of Leu336 was measured in PyMOL (v3.1.3).

#### Classical Molecular Dynamics

To ensure robust and reproducible results, we employed two complementary molecular dynamics simulation protocols with different force fields, equilibration procedures, and simulation lengths. This dual-approach strategy was designed to validate our findings across different computational frameworks and exclude potential artifacts arising from specific methodological choices.

#### Validation Simulations with AMBER Force Field

To validate key findings using an independent computational framework, we performed five replicate 1-μs simulations using AMBER20^59^ with GPU acceleration^60–62^ and ff19SB^63^/lipid21^64^/GAFF2^65^ force fields for protein, lipid, and ligand respectively, with tip3p^66^ water. Systems were prepared via Schrödinger Maestro 2021-3 with missing loops (up to 10 residues) filled using homology modelling protocol. Histidine residues were assigned δ-tautomeric states, with titratable residues protonated according to physiological pH (7.4), except Asp66^2.50^ which was protonated based on PROPKA3^67,68^ prediction (pKₐ = 8.16). The ligand 3- hydroxyl group was deprotonated per Schrödinger Epik^69,70^ prediction.

Membrane systems were built using CHARMM-GUI server^71–78^ with protein oriented via their OPM server^79^ communication tool and POPC bilayers (100×100 Å for receptor-only systems, 130×130 Å for complexes) in 150 mM NaCl solution with at least 22.5 Å thickness at each side. Systems underwent modified CHARMM-GUI equilibration protocol (47.5 ns total): 50,000-step minimization (first 15,000 steps steepest descent) with positional restraints on protein/ligand atoms (10 kcal/mol/Å²) and lipid phosphorus atoms (2.5 kcal/mol/Å²), followed by six equilibration phases with decreasing restraints. Production simulations used Langevin dynamics^80^ (1.0 ps⁻¹ friction coefficient) at 310.15 K and 1 bar with Berendsen barostat^81^ (τcouple 1.0 ps⁻¹) and 9 Å nonbonded cutoffs.

#### Extended Sampling with CHARMM36m Force Field

For comprehensive conformational sampling, we performed extended simulations using ten replicate 2-μs trajectories for key systems (6-OAU, wild-type OX04529, Phe187Ala-OX04529, Leu336Ala-OX04529, and OX04539). Simulations of 6-OAU were initiated from the published crystal structure (PDB: 8G05), while OX04529 simulations used our cryo-EM structure with the ligand positioned such that the pyridine N-oxide interacts with Arg172^ECL^^2^. OX04954 simulations were initiated by computationally replacing the CF_3_ group of OX04529 with CH_3_ using PyMOL.

The PropKa tool identified Asp66^2.50^ as likely protonated, consistent with established GPCR simulation conventions^82^. G-protein binding partners were removed, and structures were membrane-embedded using CHARMM-GUI^64 with a 35:65 CHL1:POPC lipid bilayer^71,74,75^ in rectangular boxes (80×80×120 Å for most systems, 83×82×142 Å for mutants). Systems were solvated and neutralized with 150 mM NaCl using the bilayer builder functionality.

Missing residues (1-5, 309-314, 388-394) were modelled using AlphaFold2^83^ structure alignment, while the intrinsically disordered loop (residues 231-309) was omitted to avoid sampling artifacts. Simulations employed the CHARMM36m force field with TIP3P water using GROMACS 2021.4^84^. Ligand parameters were generated using the “Ligand Reader and Modeler” functionality of CHARMM-GUI^73^.

Systems underwent steepest-descent minimization (5000 steps) with LINCS restraints on backbone, sidechain, lipids and dihedrals (4000, 2000, 1000 and 1000 units respectively), followed by six equilibration rounds with gradually relaxed restraints over 1.875 ns total. Production simulations (2 μs each, 10 replicates per system) were conducted in the NPT ensemble at 310 K using the Bussi-Donadio-Parrinello (“v-rescale”) thermostat^85^ and Parrinello-Rahman barostat^86,87^, with 12 Å nonbonded cutoffs and Fast Smooth Particle-Mesh-Ewald algorithm for long range electrostatics^88^.

### Analysis Methods

Trajectories were analysed using MDAnalysis 2.6.1^89,90^ and AmberTools23 CPPTRAJ^59^. Systems were centred, aligned, and imaged to remove periodic boundary artifacts using GROMACS functionalities and CPPTRAJ by Cα atoms of the 7-transmembrane bundle. Hydrogen bonds were detected using 3.5 Å distance and 150°/30° angle cutoffs for GROMACS/AMBER analyses, respectively. TM6 rotation angles were measured by comparing Leu336^6.52^-Leu337^6.53^ Cα vectors to the reference cryo-EM structure oriented by PPM3^91^ on OPM server^79^. Distance measurements, contact analysis using MDAnalysis, and occupancy mapping (0.5 Å grid, 75% isosurface) were performed. Numeric analyses used Numpy^92^ and Scipy^93^ libraries, with visualizations created using Matplotlib^94^ and Seaborn^95^.

This dual-protocol approach ensured that our key findings—particularly the OX04529-induced TM5-TM6 separation and associated conformational changes—were reproducible across different force fields, equilibration strategies, and simulation engines, thereby strengthening confidence in the mechanistic insights derived from our computational studies.

### Compound synthesis

Detailed synthetic routes for all compounds are provided in the Supplementary Information. All compounds used in biological assays were ≥95% pure, as determined by high-performance liquid chromatography (HPLC) using a Shimadzu SIL-20AC HT instrument.

### pKa measurements

#### pKa Determination by UV-Metric Titration

OX04529 pKa was determined using a UV-metric titration method on a Sirius T3 with a combination Ag/AgCl pH electrode. 5 µL of OX04529 stock solution was added to 25 µL of a neutral linear buffer. The sample was then pre-acidified to pH < 2.0 using 0.5 M HCl and titrated three times until pH > 12.0 using 0.5 M KOH. Titrations were carried out at a constant ionic strength of 0.15 M (adjusted with KCl) and maintained at 25.0 ± 0.5 °C. UV absorbance data across varying wavelengths and pH values were recorded. Molar absorption profiles and species distribution plots were generated in the Sirius T3 software, from which apparent pKa value was calculated.

#### pKa Determination by pH-Metric Titration

OX04529 pKa was measured using pH-metric titration on the Sirius T3 system. The solution was pre-acidified to pH < 3.0 with 0.5 M HCl and titrated three times until pH > 11.0 using 0.5 M KOH. All titrations were conducted at 25.0 ± 0.5 °C under constant ionic strength (0.15 M). pKa value was calculated from potentiometric titration curves fitted into to a theoretical ionization model using Sirius T3 software.

### Calculation of physicochemical properties

The logarithm of the partition coefficient (logP) was calculated using DataWarrior (v5.5.0). To assess lipophilic efficiency (LLE), values were computed as the difference between biological potency and lipophilicity:

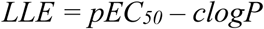

Herein, pEC_50_ values were derived from cAMP accumulation assays, representing the activation of G protein signalling at GPR84.

### Van der Waals volume estimation

Van der Waals molecular volumes were estimated using two approaches: (a) Element-based VDM-V: atomic contributions were summed using standard Van der Waals radii in DataWarrior (v5.5.0). (b) Structure-based VDM-V: 3D ligand conformations from MOE docking were imported into Visual Molecular Dynamics (VMD), and solvent-excluded volumes were calculated using built-in van der Waals volume measurement tools.

### Correlation analysis

To evaluate correlations between physicochemical properties and biological activity, Pearson correlation coefficients were computed to evaluate relationships between physicochemical and biological parameters. Statistical significance was determined using two-tailed p-values, adjusted for multiple comparisons via the Holm–Bonferroni method. All statistical analyses were conducted in R (v4.5.0).

## Acknowledgements

This work was supported by Biotechnology and Biological Sciences Research Council (U.K.) grant BB/T000562/1 (to G.M.) and BB/R007101/1 (to I.G.T.), the Wellcome Trust, Grant Number 218514/Z/19/Z, Merck Sharp and Dohme Corp. and Janssen Pharmaceutica NV (to A.J.R., R.I), European Union’s Horizon 2020 research and innovation program (101026581) through a Marie Skłodowska-Curie Individual Fellowship (to A.R.), Sir William Dunn School of Pathology, Mary Somerville Clarendon Graduate Scholarship, Clarendon Fund (to V.B.L.), and the British Heart Foundation grants RG/15/10/23915 & RG/15/10/31485 and EPA Trust GN05 (to D.R.G.). This project made use of computational time on Kelvin- 2 supported by Engineering and Physical Sciences Research Council (EPSRC) (grant no. <u>EP/T022175/1 and</u> <u>EP/W03204X/1</u>) and ARCHER2 granted via the UK High-End Computing Consortium for Biomolecular Simulation, HECBioSim (https://www.hecbiosim.ac.uk), supported by EPSRC (grant no. EP/X035603/1). The Pittsburgh Center for CryoEM (RRID:SCR_025216) used for data collection in this project was supported, in part, by the University of Pittsburgh, the School of Medicine, the Department of Structural Biology, and the National Institutes of Health (grants S10-OD-019995 and S10-OD-025009). The content is solely the responsibility of the authors and does not necessarily represent the official views of the National Institutes of Health. C.Z. is supported by the National Institutes of Health funding R35GM128641. L.O. is supported by the Indonesia Endowment Fund for Education (S-3917/LPDP.4/2022).

## Author Contributions

A.J.R., G.M., I.G.T. and C.Z. conceived the project and designed the research with P.W. P.W. designed the ligands; P.W., A.R., R.I., and L.O. synthesized the compounds and performed chemical characterization under the supervision of A.J.R. X.Z. performed protein expression and purification studies, screened cryo-EM grids, collected cryo-EM data, and processed the data under the supervision of C.Z. L.J. and S.M. performed pharmacological and mutagenesis studies under the supervision of G.M. A-A.G. performed computational analysis under the supervision of I.G.T.; C.vH. and J.D.C. performed computational analysis under the supervision of P.C.B. P.W., V.B.L., R.I., and L.O. performed initial cAMP and β-arrestin-2 recruitment assays under the supervision of D.R.G. A.J.R., G.M., I.G.T., C.Z. and P.W. wrote the manuscript, with support from D.R.G., X.Z., A.-A.G., L.J. and C.v.H. All authors proofread the manuscript.

## Materials & Correspondence

Correspondence and material requests should be addressed to A.R., C.Z., I.G.T., or G.M.

## Extended Data Tables

**Extended Data Table 1:** Potency and relative efficacy of compounds at orthosteric site mutations in the [^35^S]GTPψS binding assay.

|  | PSB-16671 |  | OX04539 |  | OX04954 |  | OX04529 |  |
| --- | --- | --- | --- | --- | --- | --- | --- | --- |
|  | pEC<sub>50</sub> ± SEM | E<sub>max</sub> ± SEM | pEC<sub>50</sub> ± SEM | E<sub>max</sub> ± SEM | pEC<sub>50</sub> ± SEM | E<sub>max</sub> ± SEM | pEC<sub>50</sub> ± SEM | E<sub>max</sub> ± SEM |
| GPR84-WT | 6.73 ± 0.15 | 100 | 9.61 ± 0.05 | 150 ± 14 | 9.18 ± 0.20 | 129 ± 12 | 8.93 ± 0.03 | 119 ± 13 |
| Y69A | 6.96 ± 0.07 | 100 | 9.04 ± 0.02<sup>**</sup> | 101 ± 14 | 8.29 ± 0.11<sup>***</sup> | 106 ± 4 | 8.34 ± 0.11<sup>**</sup> | 94 ± 9 |
| Y81A | 6.49 ± 0.11 | 100 | 8.27 ± 0.16<sup>****</sup> | 134 ± 5 | 7.79 ± 0.19<sup>****</sup> | 133 ± 18 | 7.67 ± 0.07<sup>****</sup> | 85 ± 4 |
| N104A | 6.14 ± 0.08<sup>****</sup> | 100 | 8.12 ± 0.10<sup>****</sup> | 150 ± 23 | 7.93 ± 0.07<sup>****</sup> | 130 ± 20 | 8.40 ± 0.02<sup>*</sup> | 68 ± 8 |
| T167A | 6.32 ± 0.05<sup>**</sup> | 100 | 8.23 ± 0.06<sup>****</sup> | 139 ± 7 | 7.82 ± 0.10<sup>****</sup> | 129 ± 15 | 7.75 ± 0.10<sup>****</sup> | 82 ± 10 |
| S169A | 6.18 ± 0.04<sup>***</sup> | 100 | 6.32 ± 0.07<sup>****</sup> | 84 ± 4 | 6.42 ± 0.14<sup>****</sup> | 47 ± 11<sup>*</sup> | 6.38 ± 0.04<sup>****</sup> | 30 ± 2<sup>**</sup> |
| R172A | 6.09 ± 0.06<sup>****</sup> | 100 | 5.67 ± 0.05<sup>****</sup> | 42 ± 4 | NR | NR | NR | NR |
| W360A | 7.09 ± 0.02<sup>*</sup> | 100 | 8.23 ± 0.03<sup>****</sup> | 94 ± 13 | 7.87 ± 0.15<sup>****</sup> | 75 ± 8 | 8.06 ± 0.07<sup>****</sup> | 75 ± 10 |

\*p<0.05, \*\*p<0.01, \*\*\*p<0.001 \*\*\*\*p<0.0001. One-way ANOVA followed by Dunnett's multiple comparison test. E<sub>max</sub> calculated as % of PSB-16671 response, NR, no detectable response. Data represent means ± S.E.M. from at least three biologically independent experiments.

**Extended Data Table 2:**
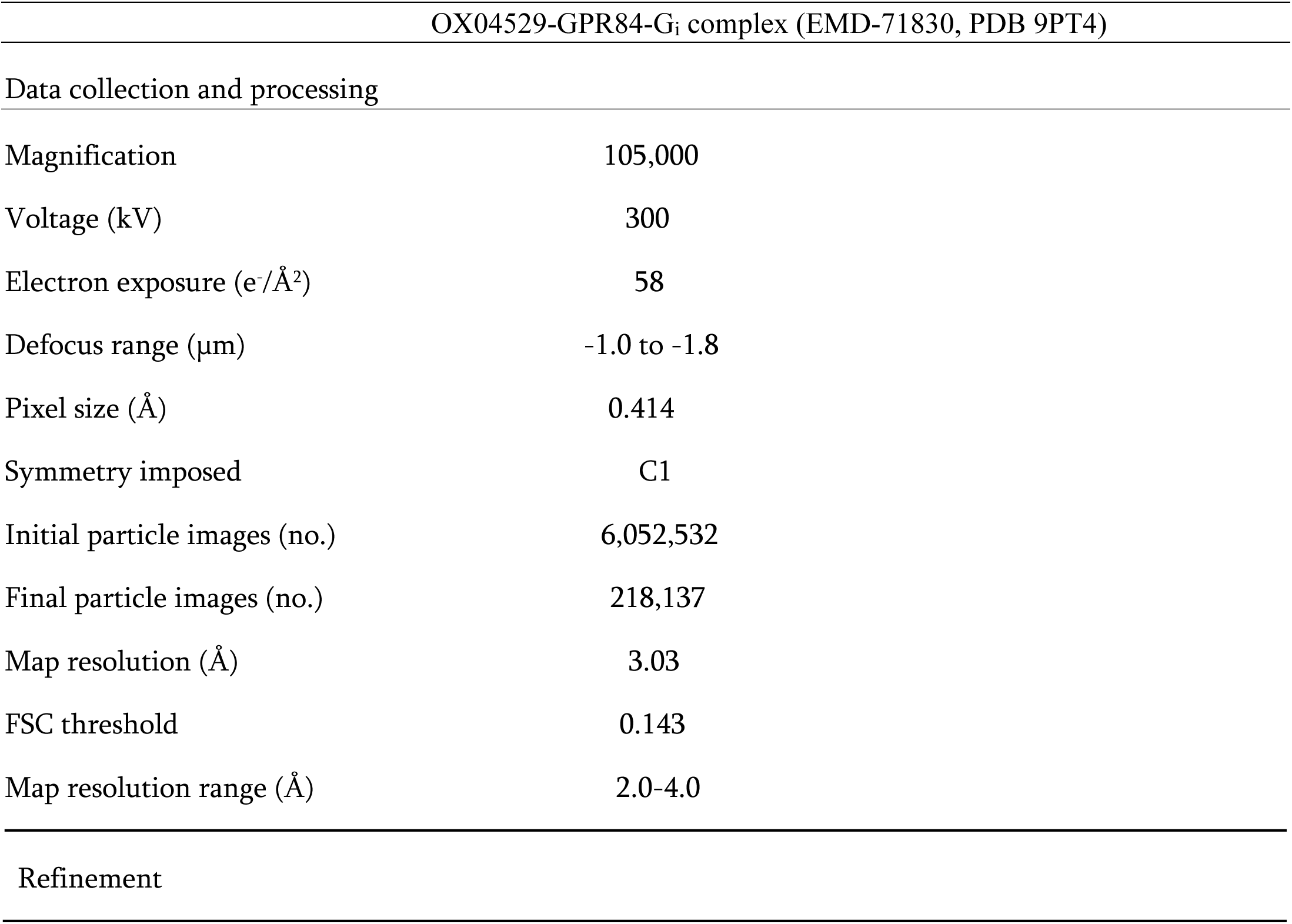
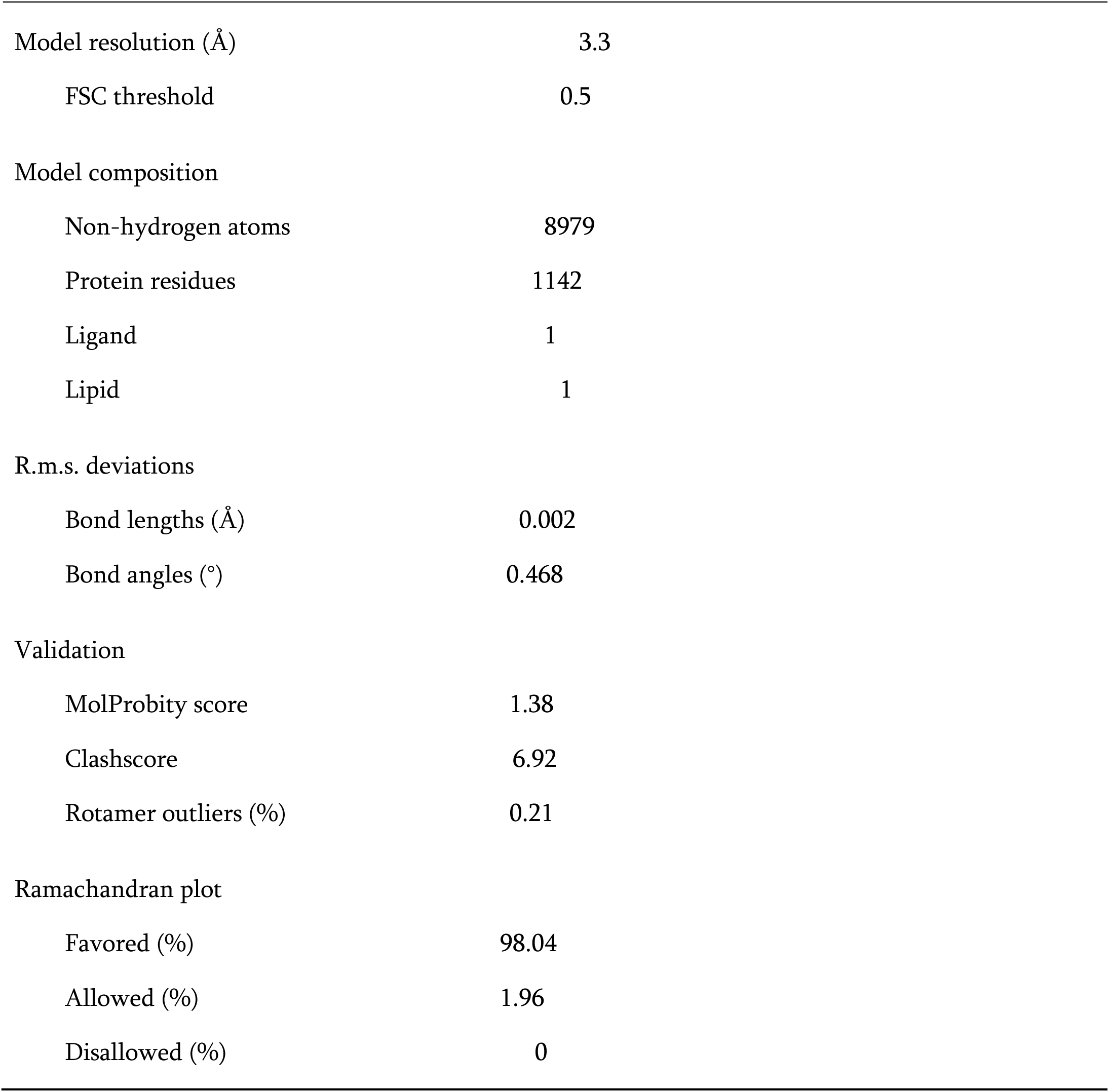
Cryo-EM data collection and refinement statistics.

| OX04529-GPR84-G<sub>i</sub> complex (EMD-71830, PDB 9PT4) |  |
| --- | --- |
| Data collection and processing |  |
| Magnification | 105,000 |
| Voltage (kV) | 300 |
| Electron exposure (e<sup>-</sup>/Å<sup>2</sup>) | 58 |
| Defocus range (μm) | -1.0 to -1.8 |
| Pixel size (Å) | 0.414 |
| Symmetry imposed | C1 |
| Initial particle images (no.) | 6,052,532 |
| Final particle images (no.) | 218,137 |
| Map resolution (Å) | 3.03 |
| FSC threshold | 0.143 |
| Map resolution range (Å) | 2.0-4.0 |
| Refinement |  |
| Model resolution (Å) | 3.3 |
| FSC threshold | 0.5 |
| Model composition |  |
| Non-hydrogen atoms | 8979 |
| Protein residues | 1142 |
| Ligand | 1 |
| Lipid | 1 |
| R.m.s. deviations |  |
| Bond lengths (Å) | 0.002 |
| Bond angles (°) | 0.468 |
| Validation |  |
| MolProbity score | 1.38 |
| Clashscore | 6.92 |
| Rotamer outliers (%) | 0.21 |
| Ramachandran plot |  |
| Favored (%) | 98.04 |
| Allowed (%) | 1.96 |
| Disallowed (%) | 0 |

**Extended Data Table 3:** Potency of compounds in [^35^S]GTPψS binding assay at Phe^187^Ala and Leu^336^Ala GPR84.

|  | OX04539<br>pEC<sub>50</sub> ± SEM | OX04954<br>pEC<sub>50</sub> ± SEM | OX04529<br>pEC<sub>50</sub> ± SEM | OX04959<br>pEC<sub>50</sub> ± SEM | OX04960<br>pEC<sub>50</sub> ± SEM | PSB-16671<br>pEC<sub>50</sub> ± SEM | 6-OAU<br>pEC<sub>50</sub> ± SEM |
| --- | --- | --- | --- | --- | --- | --- | --- |
| GPR84-WT | 9.49 ± 0.12 | 9.18 ± 0.20 | 8.90 ± 0.04 | 8.40 ± 0.05 | 8.20 ± 0.09 | 6.73 ± 0.15 | 6.48 ± 0.03 |
| F187A | 10.08 ± 0.17** | 9.97 ± 0.10** | 9.99 ± 0.01**** | nd | nd | 7.08 ± 0.09* | 7.50 ± 0.18*** |
| L336A | 8.64 ± 0.07**** | 10.37 ± 0.16**** | 10.16 ± 0.09**** | nd | nd | 6.48 ± 0.07 | 6.32 ± 0.10 |

\*p<0.05, \*\*p<0.01, \*\*\*p<0.001 \*\*\*\*p<0.0001. One-way ANOVA followed by Dunnett's multiple comparison test. nd = not determined
Data represent mean ± S.E.M. from at least three biologically independent experiments.

**Extended Data Table 4:** The potency of OX04529 to recruit β-arrestins isoforms is enhanced at Phe^187^Ala and Leu^336^Ala GPR84.

|  | OX04539 |  | OX04954 |  | OX04529 |  |
| --- | --- | --- | --- | --- | --- | --- |
|  | β-arrestin-1<br>pEC<sub>50</sub> ± SEM | β-arrestin-2<br>pEC<sub>50</sub> ± SEM | β-arrestin-1<br>pEC<sub>50</sub> ± SEM | β-arrestin-2<br>pEC<sub>50</sub> ± SEM | β-arrestin-1<br>pEC<sub>50</sub> ± SEM | β-arrestin-2<br>pEC<sub>50</sub> ± SEM |
| GPR84-WT | 8.17 ± 0.05 | 8.42 ± 0.08 | 7.40 ± 0.05 | 7.55 ± 0.16 | 7.31 ± 0.09 | 7.23 ± 0.14 |
| F187A | 8.63 ± 0.08** | 8.69 ± 0.06 | 8.24 ± 0.04**** | 8.39 ± 0.05** | 8.42 ± 0.17*** | 8.51 ± 0.22** |
| L336A | 7.51 ± 0.08*** | 7.56 ± 0.11*** | 8.67 ± 0.05**** | 8.88 ± 0.07*** | 8.73 ± 0.08*** | 8.91 ± 0.06*** |

\*p<0.05, \*\*p<0.01, \*\*\*p<0.001 \*\*\*\*p<0.0001. One-way ANOVA followed by Dunnett's multiple comparison test. nd = not determined
Data represent mean ± S.E.M. from at least three biologically independent experiments.

**Extended Data Table 5:** Potency and efficacy of compounds in [^35^S]GTPψS binding assays at various mutants of GPR84.

|  | PSB-16671 |  | OX04539 |  | OX04954 |  | OX04529 |  |
| --- | --- | --- | --- | --- | --- | --- | --- | --- |
|  | pEC<sub>50</sub> ± SEM | E<sub>max</sub> ± SEM | pEC<sub>50</sub> ± SEM | E<sub>max</sub> ± SEM | pEC<sub>50</sub> ± SEM | E<sub>max</sub> ± SEM | pEC<sub>50</sub> ± SEM | E<sub>max</sub> ± SEM |
| GPR84-WT | 6.73 ± 0.15 | 100 | 9.61 ± 0.05 | 150 ± 14 | 9.18 ± 0.20 | 129 ± 12 | 8.93 ± 0.03 | 119 ± 13 |
| Y332A | 6.23 ± 0.03*** | 100 | 8.14 ± 0.17**** | 57 ± 10 | 9.37 ± 0.21 | 60 ± 11 | 9.46 ± 0.27* | 50 ± 12 |
| Y332F | 6.46 ± 0.07 | 100 | 8.95 ± 0.09*** | 167 ± 2* | 8.38 ± 0.03** | 151 ± 10 | 8.59 ± 0.07 | 134 ± 12 |
| N362S | 6.23 ± 0.08*** | 100 | 7.91 ± 0.09**** | 196 ± 32*** | 7.62 ± 0.02**** | 124 ± 19 | 7.78 ± 0.08**** | 126 ± 30 |

\*p<0.05, \*\*p<0.01, \*\*\*p<0.001 \*\*\*\*p<0.0001. One-way ANOVA followed by Dunnett's multiple comparison test. E<sub>max</sub> calculated as % of PSB-16671 response
Data represent mean ± S.E.M. from at least three biologically independent experiments.

**Extended Data Table 6:** The potency and efficacy of OX04529, OX04539 and OX04954 to recruit β- arrestin-1 to various mutants of GPR84.

|  | **OX04539** |  | **OX04954** |  | **OX04529** |  |
| --- | --- | --- | --- | --- | --- | --- |
|  | **pEC<sub>50</sub> ± SEM** | **E<sub>max</sub> ± SEM** | **pEC<sub>50</sub> ± SEM** | **E<sub>max</sub> ± SEM** | **pEC<sub>50</sub> ± SEM** | **E<sub>max</sub> ± SEM** |
| **GPR84-WT** | 8.25 ± 0.11 | 100 | 7.50 ± 0.11 | 71 ± 4<sup>**</sup> | 7.10 ± 0.11 | 18 ± 4<sup>****</sup> |
| **N104A** | 7.08 ± 0.09<sup>***</sup> | 100 | NR | NR | NR | NR |
| **S331A** | 8.25 ± 0.21 | 100 | 7.51 ± 0.04 | 66 ± 7<sup>**</sup> | 7.54 ± 0.22 | 24 ± 6<sup>****</sup> |
| **Y332A** | NR | NR | NR | NR | NR | NR |
| **Y332F** | 7.83 ± 0.11 | 100 | 7.30 ± 0.06 | 40 ± 4<sup>***</sup> | NR | NR |
| **N362S** | 7.11 ± 0.08<sup>***</sup> | 100 | NR | NR | NR | NR |

<sup>\*\*</sup>p<0.01, <sup>\*\*\*</sup>p<0.001 <sup>\*\*\*\*</sup>p<0.0001. One-way ANOVA followed by Dunnett's multiple comparison test.
E<sub>max</sub> calculated as % of OX04539 response. NR: no response
Data represents mean ± S.E.M. from at least three biologically independent experiments.

**Extended Data Table 7:** Summary of cAMP inhibition and β-arrestin-2 recruitment of OX04529 analogs at wild type GPR84.

|  | **cAMP** | **cAMP** | **β-arrestin-2** | **β-arrestin-2** |
| --- | --- | --- | --- | --- |
|  | **pEC<sub>50</sub> ± SEM** | **%E<sub>max</sub> ± SEM** | **pEC<sub>50</sub> ± SEM** | **%E<sub>max</sub> ± SEM** |
| **OX04529** | 10.92 ± 0.18 | 97 ± 0.2 | ND | 10 ± 1.5 |
| **OX04539** | 11.25 ± 0.10 | 100 ± 0.8 | 8.11 ± 0.25 | 100 |
|  | ns | ns |  | **** |
| **OX04540** | 10.69 ± 0.09 | 97 ± 0.7 | 6.96 ± 0.17 | 81 ± 5.3 |
|  | ns | ns | * | **** |
| **OX04954** | 10.90 ± 0.23 | 98 ± 1.8 | 7.03 ± 0.24 | 71 ± 1.3 |
|  | ns | ns | * | **** |
| **OX04955** | 10.45 ± 0.20 | 101 ± 1.5 | 6.58 ± 0.23 | 21 ± 2.9 |
|  | ns | ns | ** | ns |
| **OX04956** | 11.04 ± 0.12 | 98 ± 2.1 | 6.86 ± 0.39 | 34 ± 6.0 |
|  | ns | ns | * | **** |
| **OX04957** | 10.41 ± 0.11 | 102 ± 3.0 | 6.68 ± 0.23 | 46 ± 5.8 |
|  | ns | ns | ** | **** |
| **OX04958** | 10.39 ± 0.13 | 99 ± 0.1 | 7.15 ± 0.01 | 72 ± 1.3 |
|  | ns | ns | ns | **** |
| **OX04959** | 11.14 ± 0.25 | 95 ± 11.5 | 6.82 ± 0.14 | 24 ± 2.3 |
|  | ns | ns | ** | * |
| **OX04960** | 10.99 ± 0.15 | 107 ± 2.1 | 6.35 ± 0.31 | 45 ± 1.2 |
|  | ns | ns | *** | **** |
| **OX04961** | 10.95 ± 0.26 | 103 ± 3.6 | 7.40 ± 0.04 | 21 ± 0.3 |
|  | ns | ns | ns | ns |
| **OX04962** | 11.03 ± 0.32 | 97 ± 3.4 | 6.75 ± 0.36 | 18 ± 2.2 |
|  | ns | ns | ** | ns |
| **OX04963** | 11.19 ± 0.12 | 106 ± 3.8 | 7.43 ± 0.18 | 15 ± 1.1 |
|  | ns | ns | ns | ns |
| **OX04964** | 10.84 ± 0.31 | 94 ± 7.4 | ND | 9 ± 1.4 |
|  | ns | ns |  | ns |
| **OX04965** | 11.40 ± 0.08 | 103 ± 3.7 | ND | 9 ± 3.0 |
|  | ns | ns |  | ns |
| **OX04966** | 9.77 ± 0.21 | 107 ± 5.6 | ND | 5 ± 1.5 |
|  | ** | ns |  | ns |

\*p<0.05, \*\*p<0.01, \*\*\*p<0.001 \*\*\*\*p<0.0001, ns not significant. One-way ANOVA followed by Dunnett's multiple comparison test. ND = not determinable. All compounds are compared to OX04529, β-arrestin-2 pEC<sub>50</sub> is compared to OX04539.

## Extended Data Figures

**Extended Data Fig. 1:**
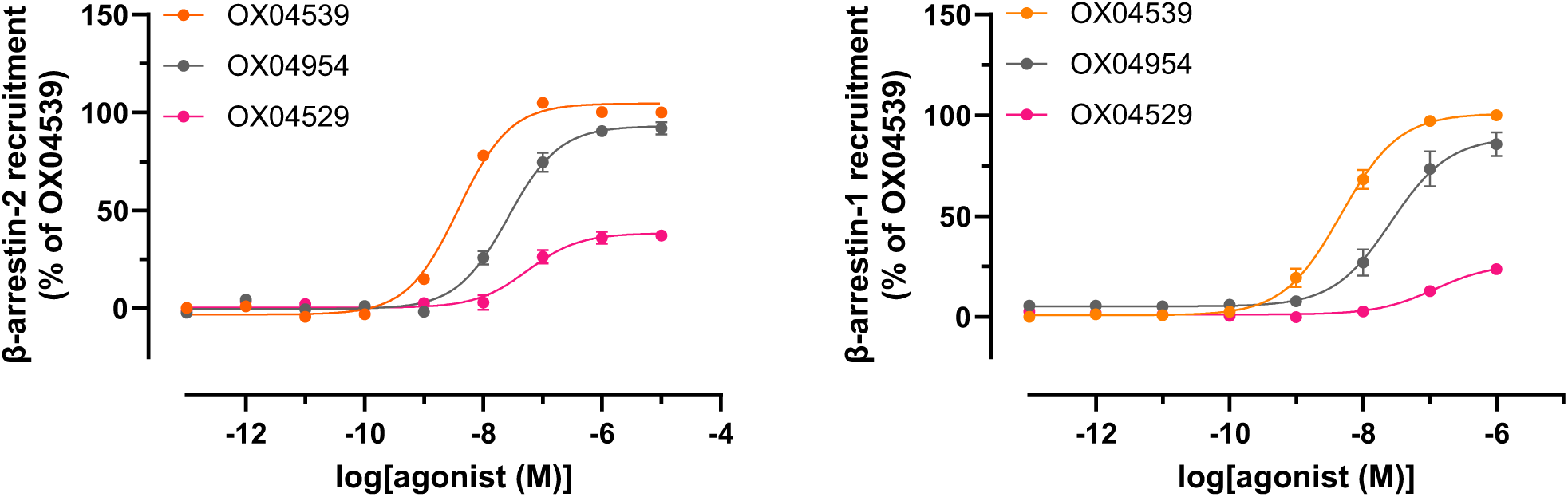
GPR84-arrestin interactions in HEK293T cells. Interactions between GPR84 and either β-arrestin-2 (left) or β-arrestin-1 (right) produced by various concentrations of the indicated ligands are shown. Data represent means ± S.E.M. from at least three biologically independent experiments.

**Extended Data Fig. 2:**
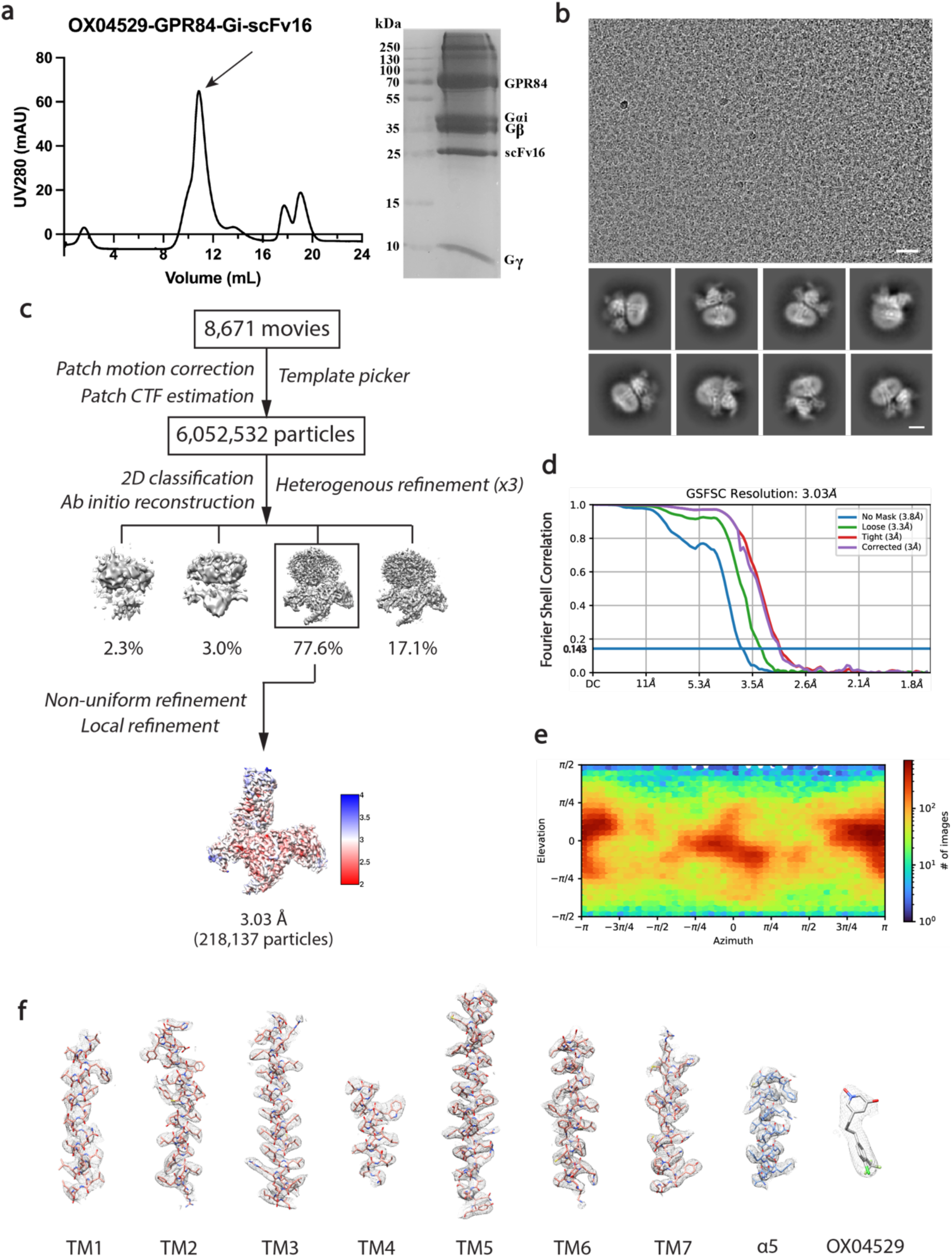
Purification of the OX04529-GPR84-G_i_ complex and cryo-EM data processing. **(a)** Size-exclusion chromatography profile and SDS-PAGE analysis of the purified OX04529-GPR84-G_i_ complex. **(b)** Representative cryo-EM micrograph (scale bar: 50 nm) and 2D class averages (scale bar: 5 nm). **(c)** Cryo-EM image processing workflow for the OX04529-GPR84-G_i_ complex. Density map according to local resolution estimation. **(d)** Gold-standard Fourier shell correlation (FSC) curve showing an overall resolution is 3.03 Å at FSC=0.143. **(e)** Angular distribution of the particles used in the final reconstruction. **(f)** Cryo-EM density maps and models of the seven transmembrane helices (TM1-7), α5 helix of G_i_ and the ligand of OX04529 bound GPR84-G_i_ complex are shown.

**Extended Data Fig. 3:**
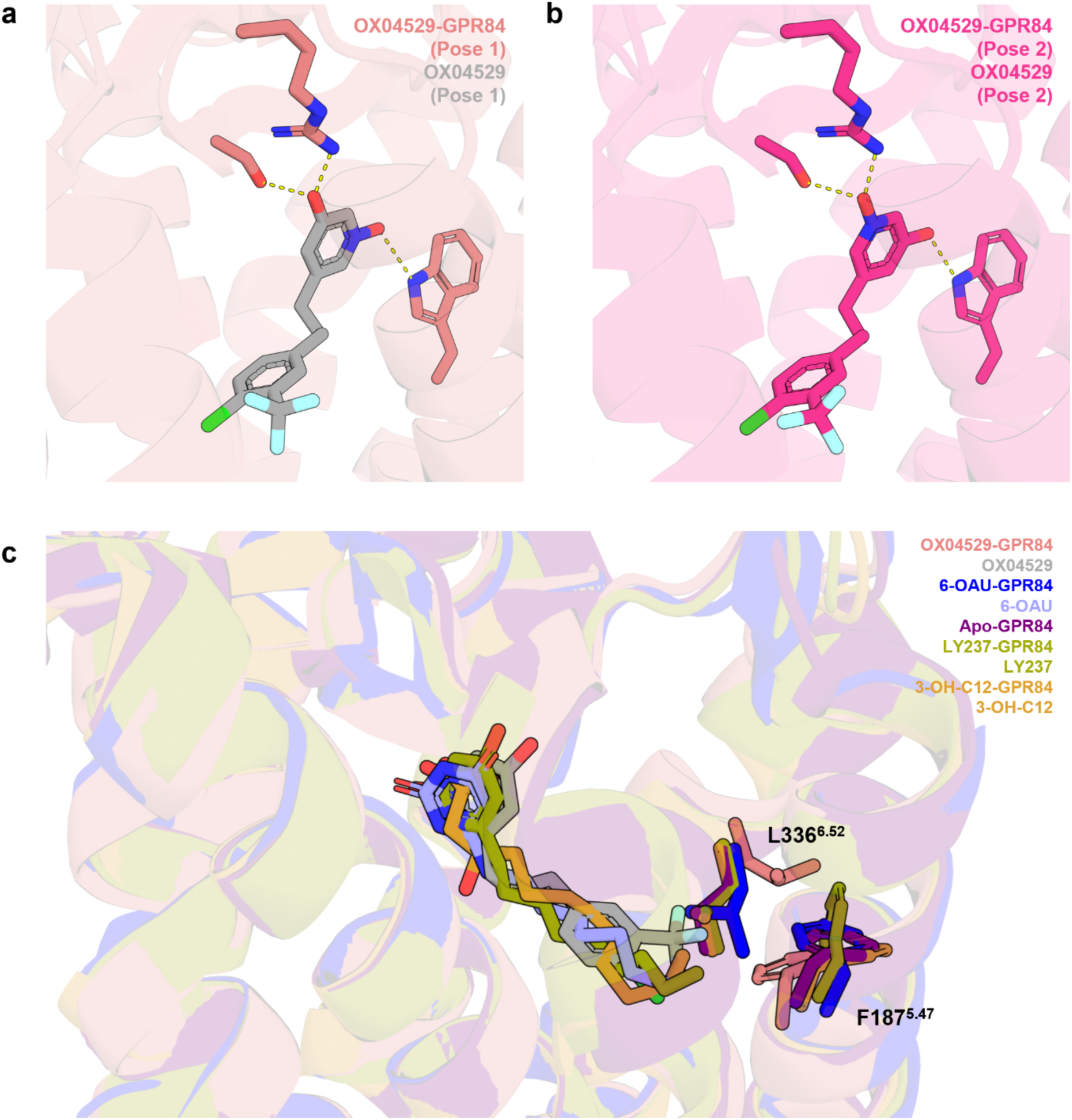
Alternative binding poses of the OX04529 *N*-oxide and overlay of reported cryo-EM structures at L336 and F187. **a, b,** Two alternative orientations of the N-oxide moiety in the binding pocket. In Pose 1 **(a)**, the *N*-oxide group is oriented toward W360, with the hydroxyl group directed toward R172 and S169. In Pose 2 **(b)**, the orientation is reversed. **c,** Overlay of OX04529-GPR84, 6-OAU-GPR84, apo-GPR84, LY237-GPR84 and 3- OH-C12-GPR84 structures zooming at L336 and F187.

**Extended Data Fig. 4:**
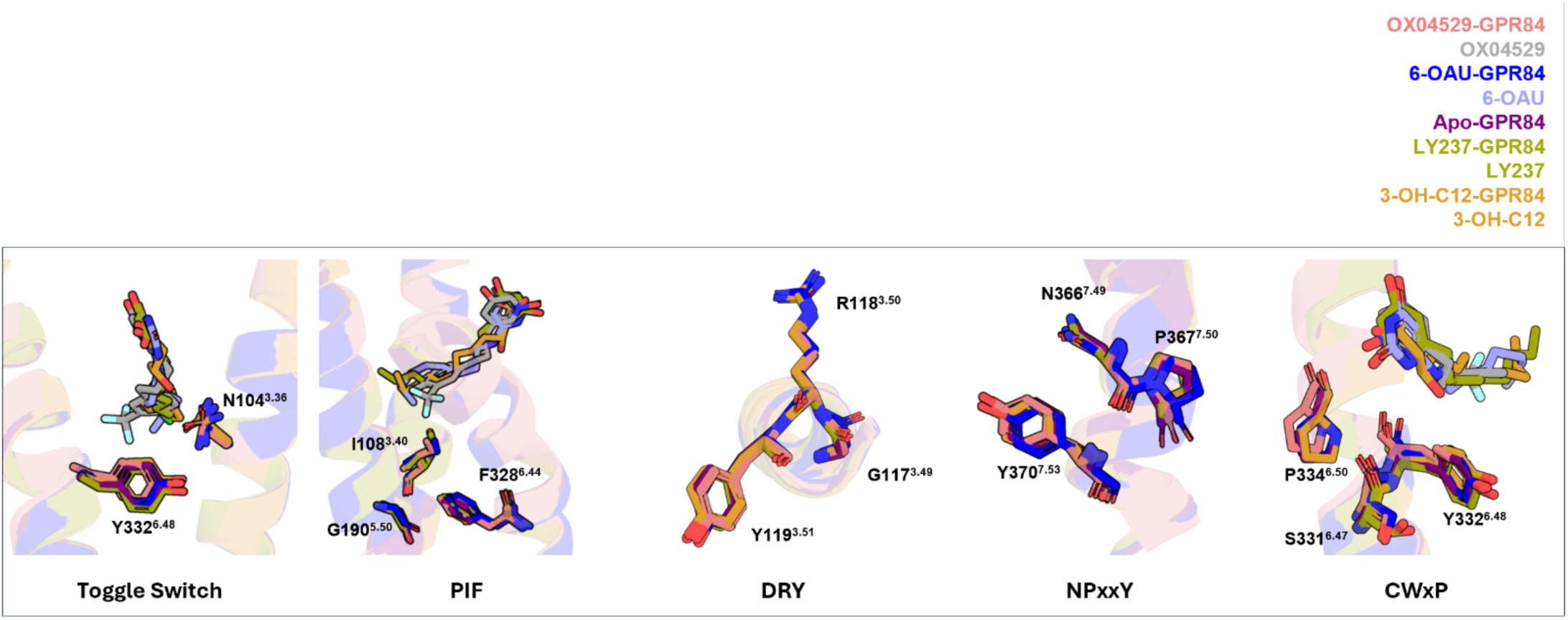
Structural comparison with other GPR84 cryo-EM co-structures. Overlay of OX04529-GPR84, 6-OAU-GPR84, apo-GPR84, LY237-GPR84 and 3-OH-C12-GPR84 structures revealed that key conserved motifs in class A GPCRs, including the toggle switch, PIF motif, DRY motif, NPxxY motif and CWxP motif.

**Extended Data Fig. 5:**
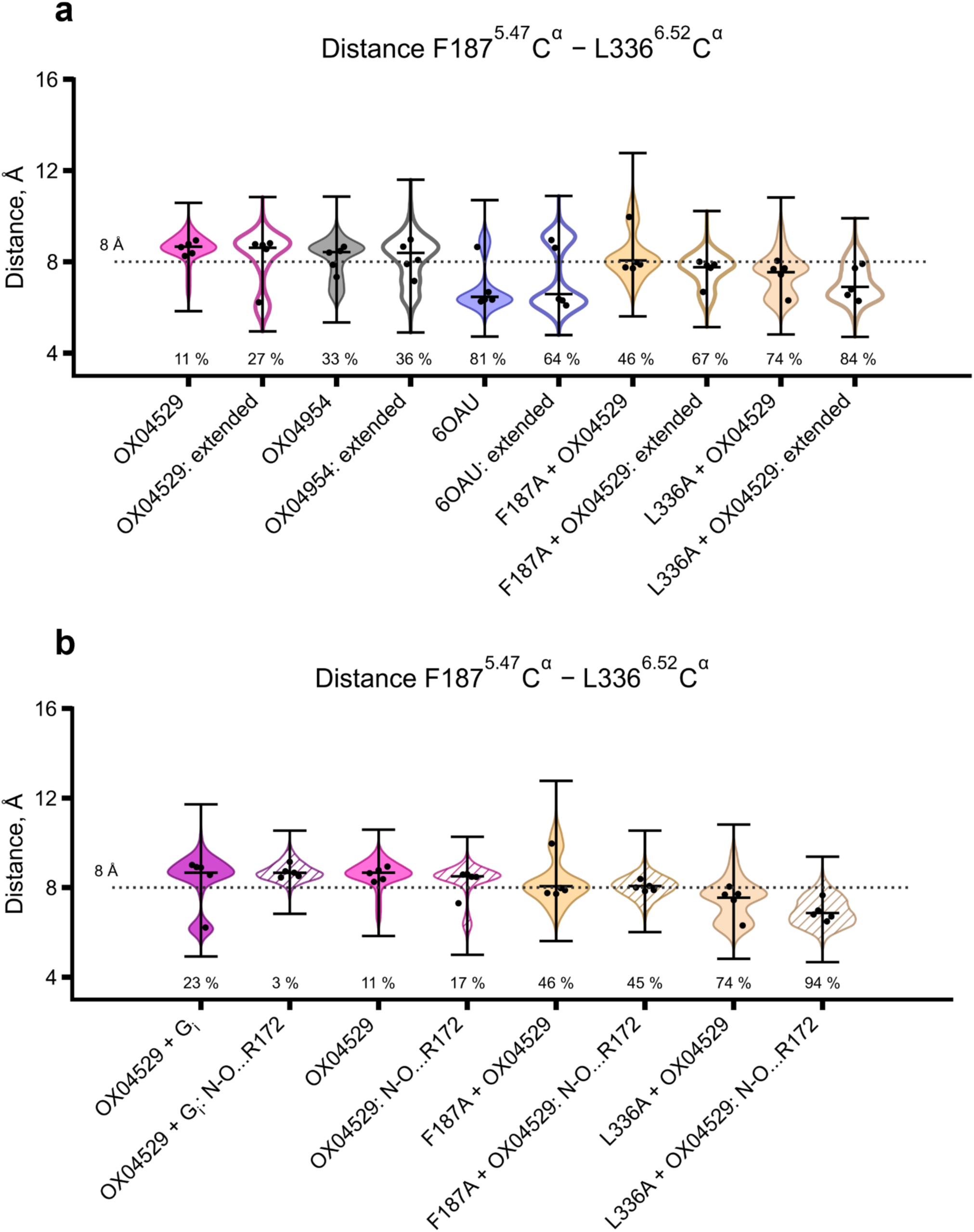
Validation of Phe187^5.47^-Leu336^6.52^separation mechanism across simulation conditions and ligand orientations. **a.** Violin plots comparing Cα-Cα distance distributions from 1-μs (5 replicates) and 2-μs (5 replicates) MD simulations. The dashed line at 8 Å separates compact and extended conformational states. Percentages indicate frames below threshold. Extended sampling confirms that OX04529 maintains the separated state (11-27% compact) while β-arrestin recruiting agonists (6-OAU) favor compaction (64-81% compact). Mutants F187A and L336A consistently restore the compact state (67-84% compact) across both timescales, validating the robustness of the steric mechanism independent of simulation length. This graph shows that both setups (forcefield, protonation, specific receptor model) give same results within the 1st microsecond, and then their results start showing deeper differences. **b.** Violin plots comparing Cα-Cα distance distributions between two possible orientations of the OX04529 3-hydroxypyridine *N*-oxide ring system from 1-μs MD simulations (5 replicates). System names with ‘+G_i_’ indicates simulations with G protein, system names containing ‘N-O…R172’ indicate simulations with the pyridine *N*-oxide oxygen interacting with Arg172^ECL^^2^, and systems labeled with a just ligand name indicates simulations with the hydroxyl group interacting with Arg172^ECL^^2^. The dashed line at 8 Å separates compact and extended states. Percentages indicate frames below threshold. Both orientations of OX04529 maintain the separated conformation (3-23% compact), while mutants Phe187Ala and Leu336Alarestore compaction (74-94% compact) regardless of head group orientation, demonstrating that the steric mechanism is independent of ligand binding pose ambiguity.

**Extended Data Fig. 6:**
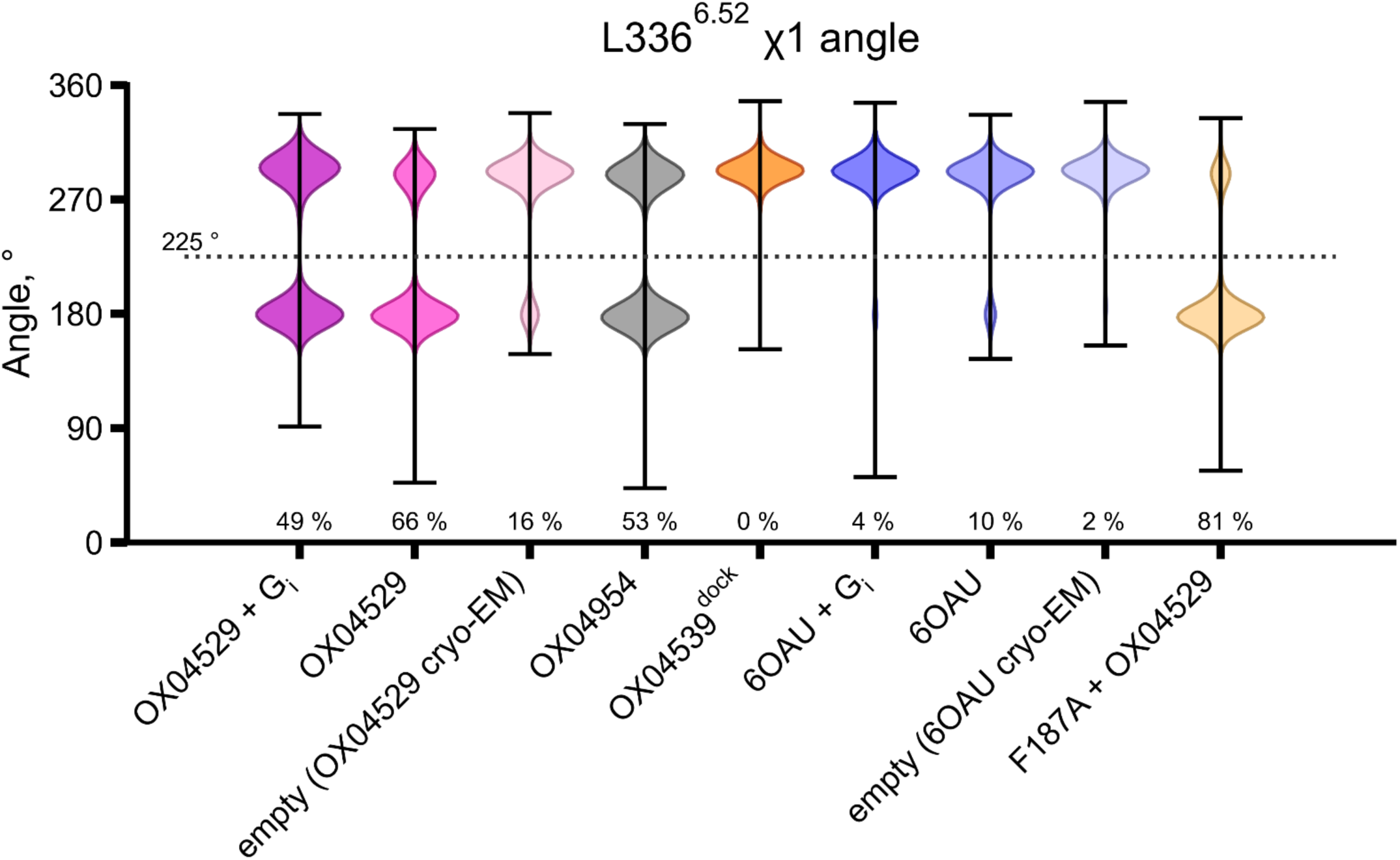
Violin plots illustrate the distribution of χ1 dihedral angles of Leu336^6.52^ obtained from five replicate 1-μs MD simulation trajectories. The analysis compares distinct rotameric states in GPR84 complexes with G protein-biased ligand OX04529, partially biased OX04954 and reference agonists (OX04539 and 6-OAU), two ligand-free receptor conformations derived from cryo-EM structures, and the Phe187^5.47^Ala mutant.

**Extended Data Fig. 7:**
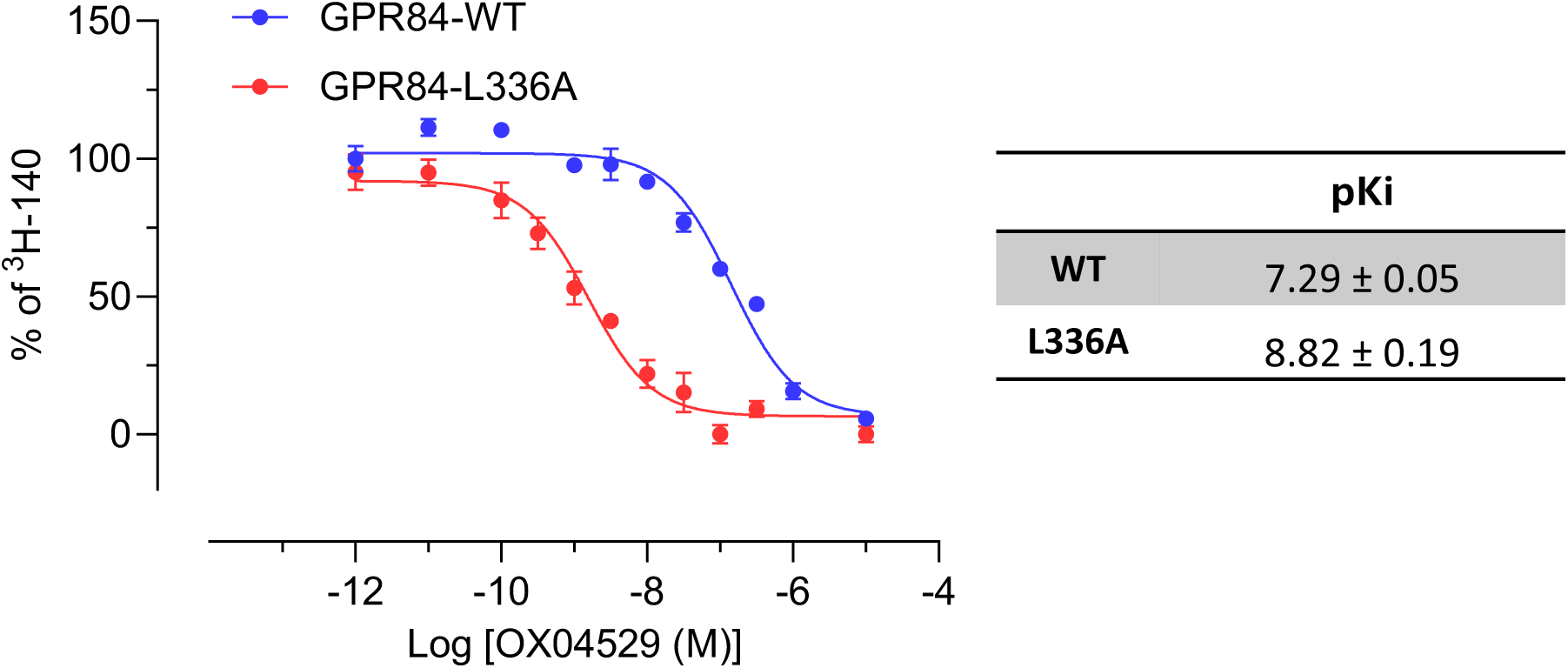
The affinity of OX04529 is markedly increased at Leu^336^Ala GPR84 in competition binding studies using stable cell lines expressing the indicated GPR84-G_i2_ fusion protein Left, data from a representative experiment, right pKi represents means +/- S.E.M. from three independent biological replicates.

**Extended Data Fig. 8:**
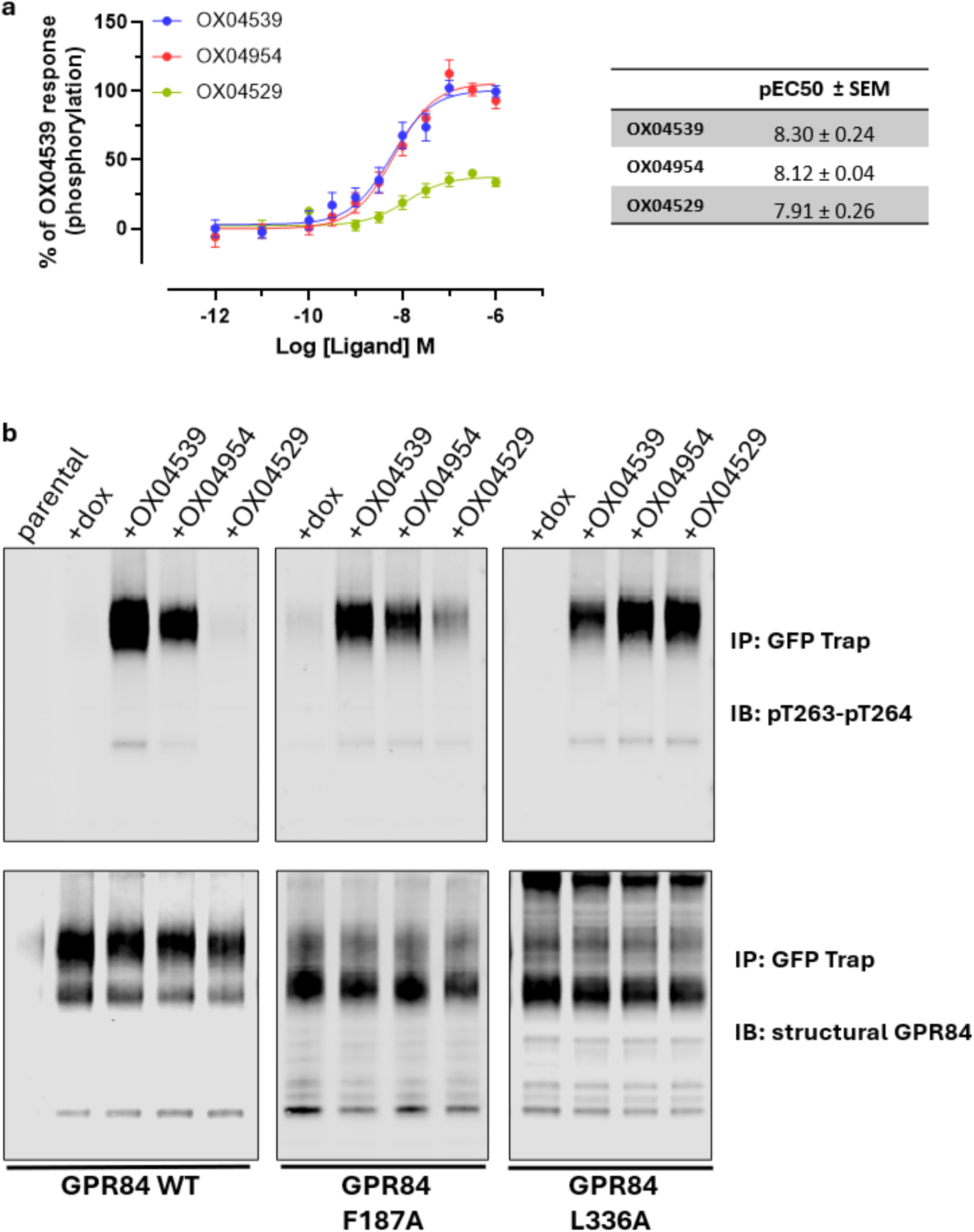
Biased agonist-induced phosphorylation of Thr263^ICL^^3^and Thr264^ICL^^3^in GPR84 is modulated by mutation of Phe187^5.47^and Leu336^6.52^. **a,** The ability of varying concentrations of OX04529, OX04539 and OX04954 to promote phosphorylation of residues Thr263^ICL^^3^ and Thr264^ICL^^3^ of wild type GPR84 was assessed in a magnetic bead capture assay. **b,** Immunoblots show the lack of basal phosphorylation of residues Thr263^ICL^^3^ and Thr264^ICL^^3^ in each of wild type, Phe187^5.47^Ala and Leu336^6.52^Ala GPR84. The ability of OX04529, OX04539 and OX04954 (1 μM) to promote phosphorylation at these sites is illustrated. A representative example of 3 independent experiments is displayed.

**Extended Data Fig. 9.**
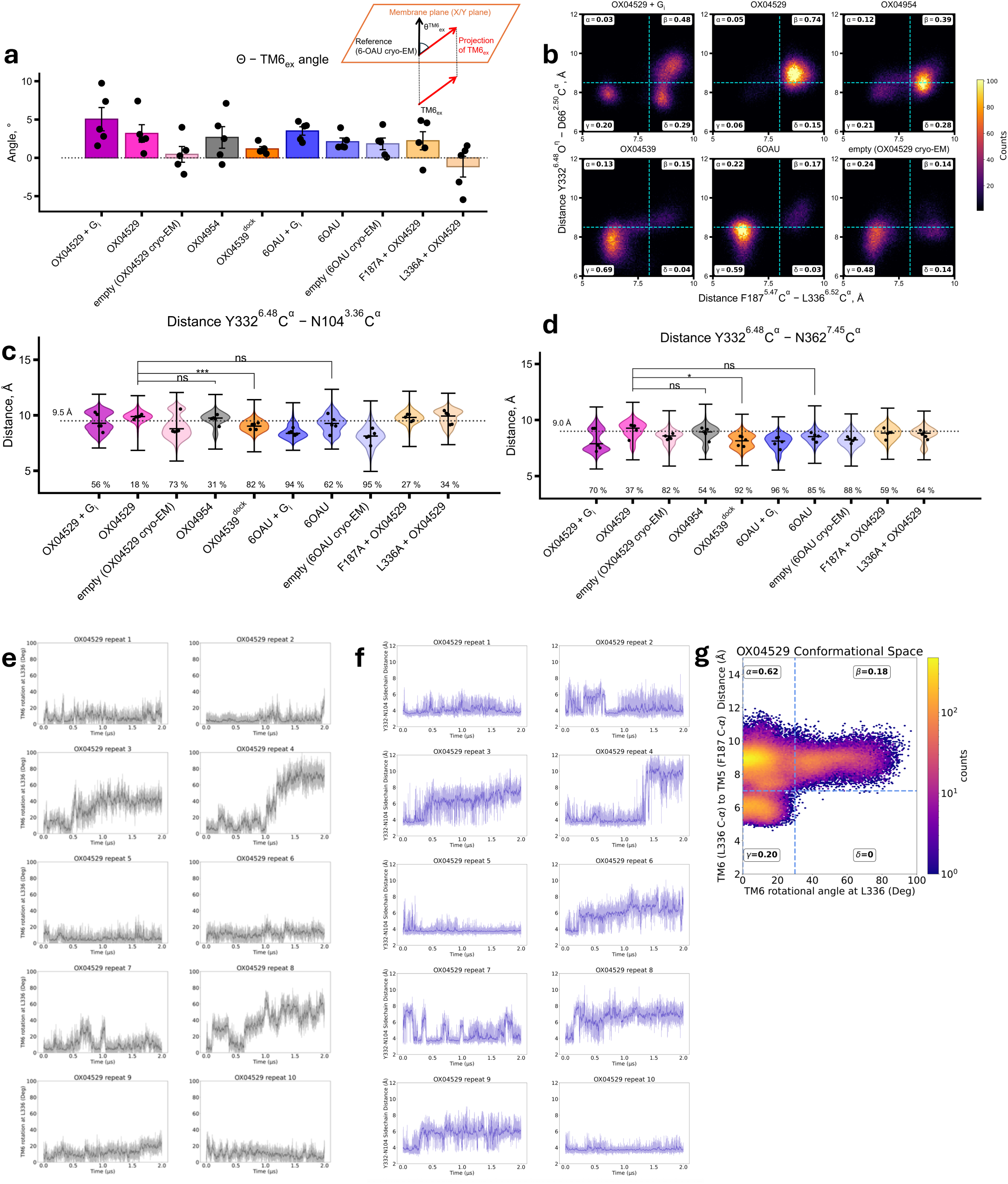
Comprehensive analysis of TM6 conformational dynamics and polar network disruption in OX04529-bound GPR84. **a.** Bar graph showing the angular deviation of the TM6 extracellular segment across different GPR84 systems from five replicate 1-μs MD simulation trajectories, measured relative to the 6-OAU cryo-EM reference structure. The inset diagram illustrates the measurement method: angles represent the projection of vectors (connecting first-to-last helical turn Cα centers of mass in TM6) onto the membrane plane. Note the significant positive angle in OX04529-bound systems, indicating outward TM6 movement away from TM5 toward the lipid interface, contrasting with the more neutral or negative angles observed with β-arrestin recruiting agonists. This conformational difference is enhanced when OX04529 is co-bound with G protein and is disrupted in the Phe187Ala and Leu336Ala mutants. **b.** Two-dimensional density plots showing the correlation between TM6 extracellular rotation angle and Phe187^5.47^-Leu336^6.52^ Cα distance across five replicate 1-μs MD simulation trajectories. Each panel represents a different system as labelled, with the proportion of simulation frames in each quadrant indicated by percentages. The dashed lines at 7 Å (vertical) and 30° (horizontal) demarcate threshold values separating distinct conformational states. OX04529-bound systems show preferential occupancy of the high-distance, high-rotation quadrant, while reference agonists (OX04539, 6-OAU) predominantly occupy the low-distance, low-rotation region. **c.** Distribution of Tyr332^6.48^-Asn104^3.36^ hydrogen bond distances across five replicate 1-μs MD simulations. The dashed line at ∼9.5 Å indicates the hydrogen bonding threshold. Percentages represent the proportion of frames maintaining hydrogen bonding. OX04529 shows the lowest maintenance (56%), demonstrating disruption of this critical toggle-switch interaction. **d.** Distribution of Tyr332^6.48^-Asn362^7.45^ hydrogen bond distances across the same simulation conditions. The dashed line at ∼9.0 Å indicates the hydrogen bonding threshold. OX04529 shows reduced stability (70%) compared to most other systems (54-96%), contributing to polar network disruption. **e.** Individual time-series traces of TM6 rotation angles from ten replicate 2-μs simulations of OX04529, demonstrating the temporal evolution and variability of TM6 conformational changes. **f.** Individual time-series traces of Tyr332^6.48^-Asn104^3.36^ distances from ten replicate 2-μs simulations of OX04529, showing the dynamic disruption of this key hydrogen bond over extended timescales. The measured distance is between the oxygen of Tyr332^6.48^ and the C-gamma of Asn104^3.36^. The quadrant labels in panels **e.** and **f.** represent the fraction of the total simulation frames in a given state of the receptor. **g.** Two-dimensional correlation plot of TM6 rotation angle versus TM5-TM6 separation distance, demonstrating that significant TM6 rotation (>30°) occurs only when the TM5-TM6 separation exceeds 7 Å, establishing the coupling between these conformational changes.

**Extended Data Fig. 10:**
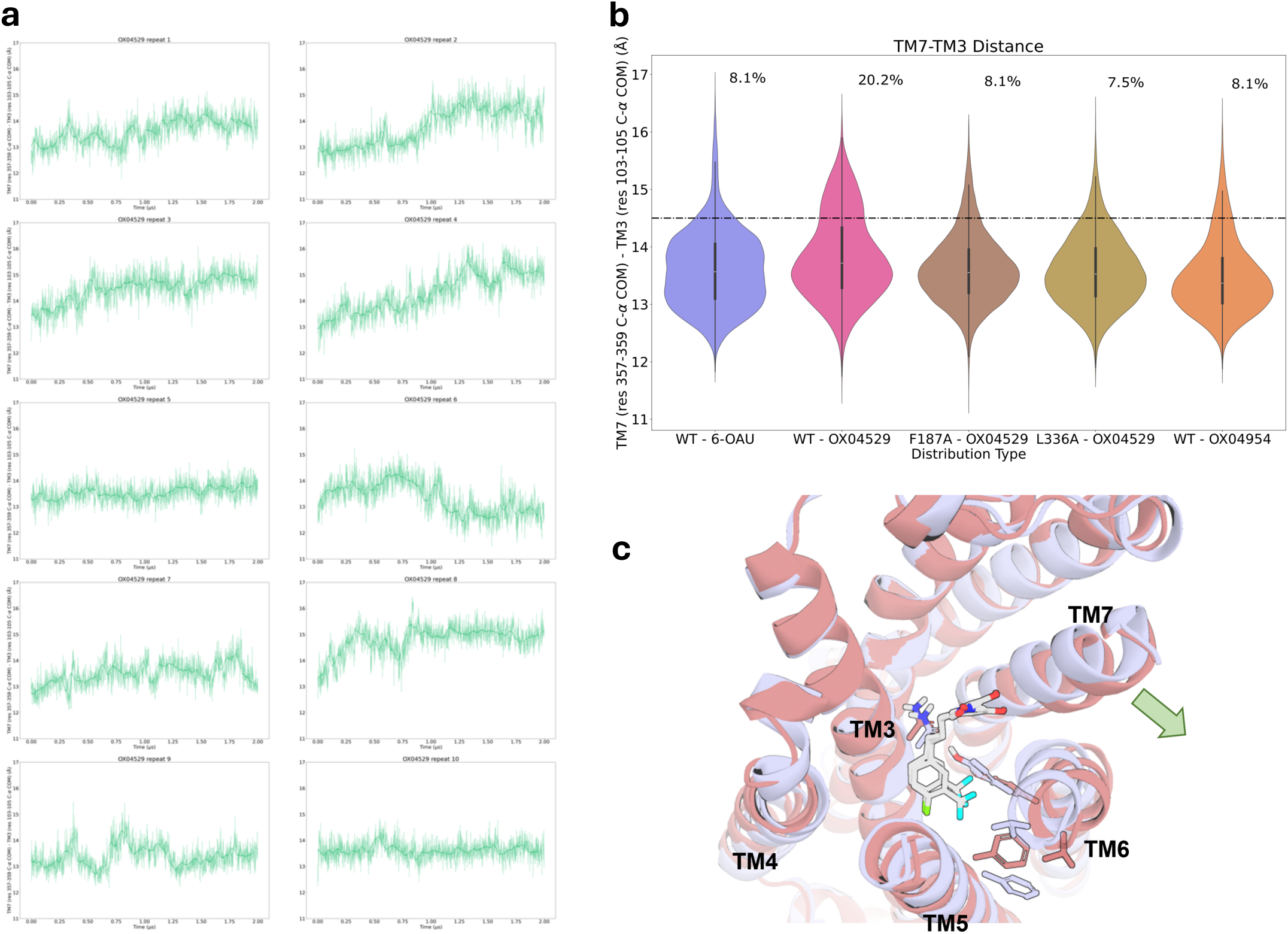
Distribution of TM7-TM3 distances obtained from 10 x 2 μs simulations. **a,** Individual time-series traces of TM7-TM3 distances from ten replicate 2-μs simulations of OX04529. **b,** Distribution of TM7-TM3 distances across ten replicate 2-μs MD simulations. Note OX04529 has a larger occupancy of higher distances. **c,** Two overlaid representative snapshots from molecular dynamics simulations showing the binding poses of G protein-biased OX04529 (gray) in the GPR84 orthosteric pocket, and the outward movement of TM7.

**Extended Data Fig. 11:**
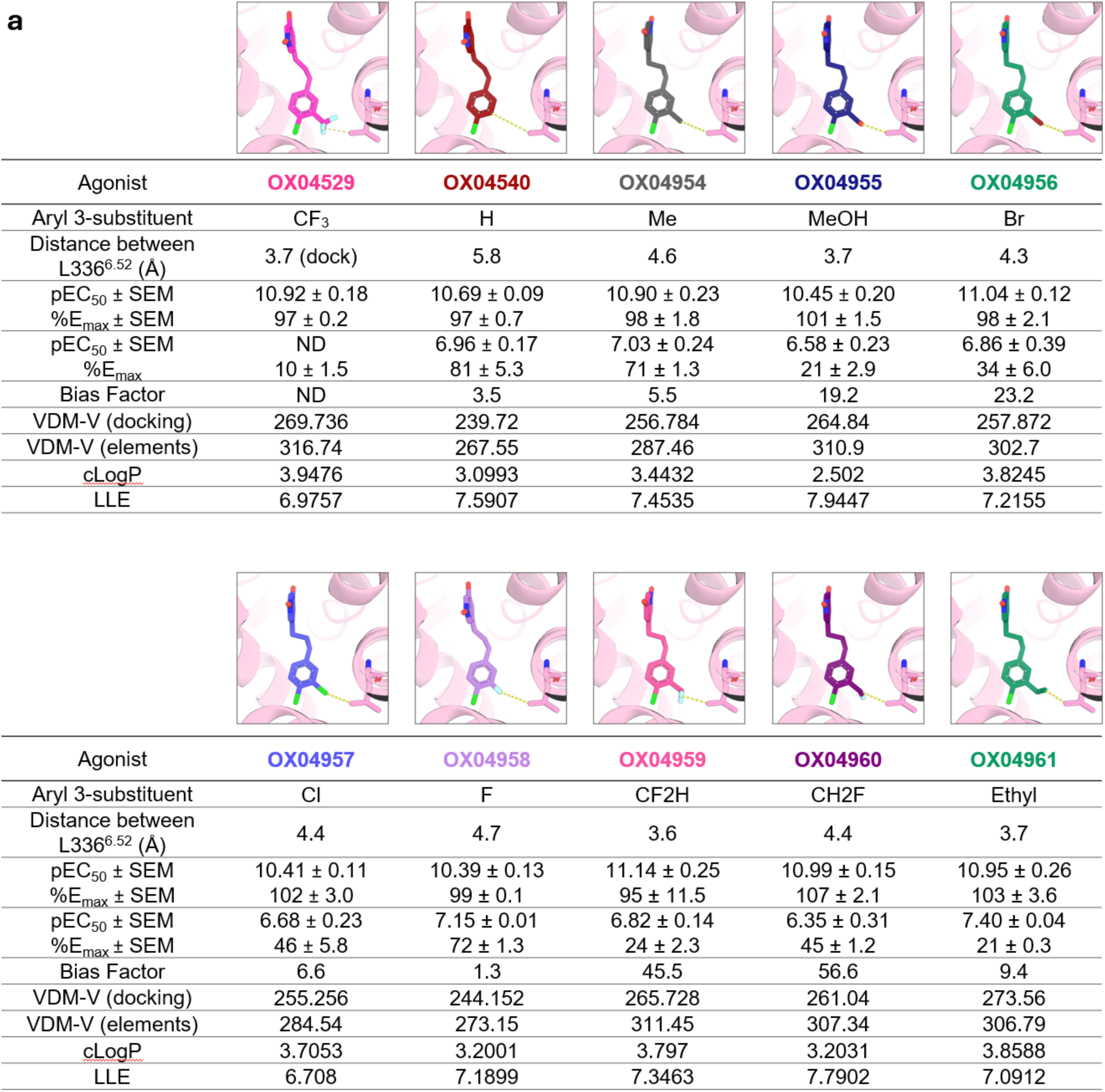
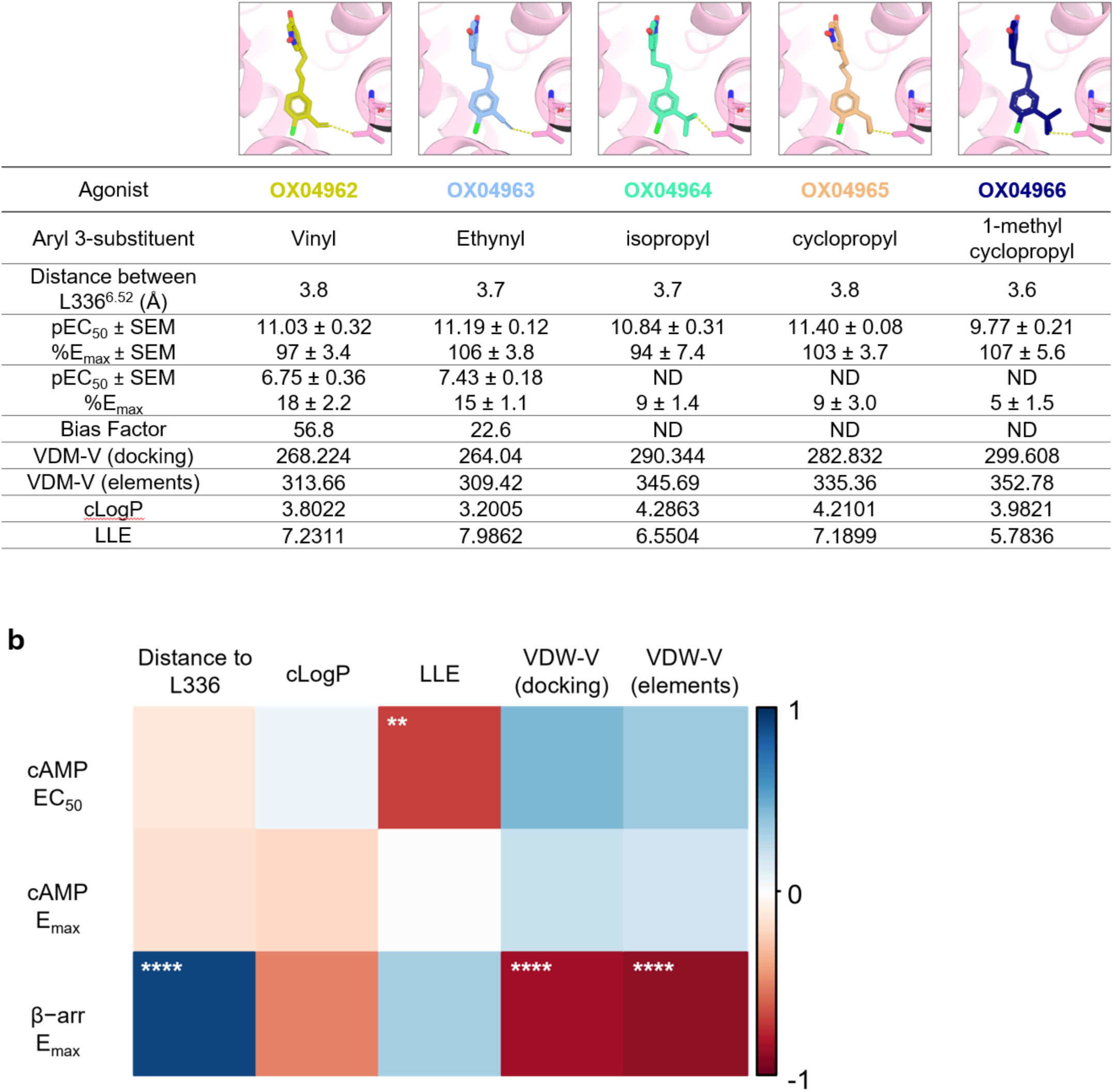
Correlation of cAMP inhibition and β-arrestin-2 recruitment of OX04529 analogs where the aryl 3-substitutent is varied in size and the calculated interatomic distance between the most distal heavy atoms of the aryl 3-substituent and Leu336^6.52^. **a**, Details of measurements of interatomic distance between the most distal heavy atoms of the ligand aryl 3-substituent and the side chain of Leu336^6.52^ taken from docking models using the cryo-EM structure of the OX04529-GPR84 complex. For details of docking studies see Methods. Comparison of interatomic distance, cAMP inhibition and β-arrestin- 2 recruitment potency and efficacy for each ligand. Data are means ± S.E.M. from at least three independent biological experiments, each performed in technical triplicate. **b**, Correlation map of β-arrestin-2 maximum efficacy, van der Waals volume (VDW-V), distance from the most distal heavy (non-H) atom of the ligand arene 3-substituent to Leu336^6.52^, cLogP and cAMP activity. VDW-volume was calculated either from the ligand elements alone (elements) or the 3D structure of the docked pose (docking). Asterisks denote statistical significance (** p < 0.01, **** p< 0.0001) of the correlation. For details of the methodology used for correlation analysis see Methods.

## References

1. Rosenbaum, D. M., Rasmussen, S. G. F. & Kobilka, B. K. The structure and function of G-protein-coupled receptors. Nature 459, 356–363 (2009).

2. Oldham, W. M. & Hamm, H. E. Heterotrimeric G protein activation by G-protein-coupled receptors. Nat. Rev. Mol. Cell Biol. 9, 60–71 (2008).

3. Gurevich, E. V., Tesmer, J. J. G., Mushegian, A. & Gurevich, V. V. G protein-coupled receptor kinases: More than just kinases and not only for GPCRs. Pharmacol. Ther. 133, 40–69 (2012).

4. Tobin, A. B. G-protein-coupled receptor phosphorylation: where, when and by whom. Br. J. Pharmacol. 153, (2008).

5. Pierce, K. L. & Lefkowitz, R. J. Classical and new roles of β-arrestins in the regulation of G-PROTEIN-COUPLED receptors. Nat. Rev. Neurosci. 2, 727–733 (2001).

6. Wang, H., et al. Structure-Based Evolution of G Protein-Biased μ-Opioid Receptor Agonists. Angew. Chem. Int. Ed. 61, (2022).

7. Zhuang, Y. et al. Molecular recognition of morphine and fentanyl by the human μ-opioid receptor. Cell 185, 4361–4375.e19 (2022).

8. Suomivuori, C.-M. et al. Molecular mechanism of biased signaling in a prototypical G protein– coupled receptor. Science 367, 881–887 (2020).

9. Tóth, A. D., Turu, G. & Hunyady, L. Functional consequences of spatial, temporal and ligand bias of G protein-coupled receptors. Nat. Rev. Nephrol. (2024) doi:10.1038/s41581-024-00869-3.

10. Morales, P. et al. Progress on the development of Class A GPCR-biased ligands. Br. J. Pharmacol. bph.17301 (2024) doi:10.1111/bph.17301.

11. Marsango, S. et al. Selective phosphorylation of threonine residues defines GPR84–arrestin interactions of biased ligands. J. Biol. Chem. 298, 101932 (2022).

12. Wang, J., Wu, X., Simonavicius, N., Tian, H. & Ling, L. Medium-chain Fatty Acids as Ligands for Orphan G Protein-coupled Receptor GPR84. J. Biol. Chem. 281, 34457–34464 (2006).

13. Recio, C. et al. Activation of the Immune-Metabolic Receptor GPR84 Enhances Inflammation and Phagocytosis in Macrophages. Front. Immunol. 9, 1419 (2018).

14. Suzuki, M. et al. Medium-chain Fatty Acid-sensing Receptor, GPR84, Is a Proinflammatory Receptor. J. Biol. Chem. 288, 10684–10691 (2013).

15. Recio, C. et al. Activation of the Immune-Metabolic Receptor GPR84 Enhances Inflammation and Phagocytosis in Macrophages. Front. Immunol. 9, 1419 (2018).

16. Lucy, D. et al. A Biased Agonist at Immunometabolic Receptor GPR84 Causes Distinct Functional Effects in Macrophages. ACS Chem. Biol. 14, 2055–2064 (2019).

17. Kamber, R. A. et al. Inter-cellular CRISPR screens reveal regulators of cancer cell phagocytosis. Nature 597, 549–554 (2021).

18. Zhang, X. et al. Pro-phagocytic function and structural basis of GPR84 signaling. Nat. Commun. 14, 5706 (2023).

19. Liu, H. et al. Structural insights into ligand recognition and activation of the medium-chain fatty acid-sensing receptor GPR84. Nat. Commun. 14, 3271 (2023).

20. Wang, P. et al. Development of Highly Potent, G-Protein Pathway Biased, Selective, and Orally Bioavailable GPR84 Agonists. J. Med. Chem. 67, 110–137 (2024).

21. Jenkins, L. et al. Discovery and Characterization of Novel Antagonists of the Proinflammatory Orphan Receptor GPR84. ACS Pharmacol. Transl. Sci. 4, 1598–1613 (2021).

22. Klein Herenbrink, C., et al. The role of kinetic context in apparent biased agonism at GPCRs. Nat. Commun. 7, 10842 (2016).

23. McCorvy, J. D. et al. Structural determinants of 5-HT2B receptor activation and biased agonism. Nat. Struct. Mol. Biol. 25, 787–796 (2018).

24. 24. Ballesteros, J. A. & Weinstein, H. Integrated methods for the construction of three-dimensional models and computational probing of structure-function relations in G protein-coupled receptors. in Methods in Neurosciences vol. 25 366–428 (Elsevier, 1995).

25. Zhou, Q. et al. Common activation mechanism of class A GPCRs. eLife 8, e50279 (2019).

26. Tehan, B. G., Bortolato, A., Blaney, F. E., Weir, M. P. & Mason, J. S. Unifying Family A GPCR Theories of Activation. Pharmacol. Ther. 143, 51–60 (2014).

27. Schönegge, A.-M. et al. Evolutionary action and structural basis of the allosteric switch controlling β2AR functional selectivity. Nat. Commun. 8, 2169 (2017).

28. Allen, J. A. & Roth, B. L. Strategies to Discover Unexpected Targets for Drugs Active at G Protein– Coupled Receptors. Annu. Rev. Pharmacol. Toxicol. 51, 117–144 (2011).

29. Fritze, O. et al. Role of the conserved NPxxY(x) 5,6 F motif in the rhodopsin ground state and during activation. Proc. Natl. Acad. Sci. 100, 2290–2295 (2003).

30. Olivella, M., Caltabiano, G. & Cordomí, A. The role of Cysteine 6.47 in class A GPCRs. BMC Struct. Biol. 13, (2013).

31. Mahmud, Z. A. et al. Three classes of ligands each bind to distinct sites on the orphan G protein-coupled receptor GPR84. Sci. Rep. 7, 17953 (2017).

32. Pillaiyar, T. et al. Diindolylmethane Derivatives: Potent Agonists of the Immunostimulatory Orphan G Protein-Coupled Receptor GPR84. J. Med. Chem. 60, 3636–3655 (2017).

33. Wingler, L. M., McMahon, C., Staus, D. P., Lefkowitz, R. J. & Kruse, A. C. Distinctive Activation Mechanism for Angiotensin Receptor Revealed by a Synthetic Nanobody. Cell 176, 479–490.e12 (2019).

34. Stanek, M. et al. Hybridization of β-Adrenergic Agonists and Antagonists Confers G Protein Bias. J. Med. Chem. 62, 5111–5131 (2019).

35. Teng, X. et al. Ligand recognition and biased agonism of the D1 dopamine receptor. Nat. Commun. 13, 3186 (2022).

36. Manglik, A. et al. Structure-based discovery of opioid analgesics with reduced side effects. Nature 537, 185–190 (2016).

37. Chen, X.-T. et al. Structure–Activity Relationships and Discovery of a G Protein Biased μ Opioid Receptor Ligand, [(3-Methoxythiophen-2-yl)methyl]({2-[(9 R)-9-(pyridin-2-yl)-6-oxaspiro-[4.5]decan- 9-yl]ethyl})amine (TRV130), for the Treatment of Acute Severe Pain. J. Med. Chem. 56, 8019–8031 (2013).

38. Cheng, L. et al. Cryo-EM structure of small-molecule agonist bound delta opioid receptor-Gi complex enables discovery of biased compound. Nat. Commun. 15, 8284 (2024).

39. Faouzi, A. et al. Structure-based design of bitopic ligands for the µ-opioid receptor. Nature 613, 767–774 (2023).

40. Rangari, V. A. et al. A cryptic pocket in CB1 drives peripheral and functional selectivity. Nature (2025) doi:10.1038/s41586-025-08618-7.

41. Wootten, D., Christopoulos, A., Marti-Solano, M., Babu, M. M. & Sexton, P. M. Mechanisms of signalling and biased agonism in G protein-coupled receptors. Nat. Rev. Mol. Cell Biol. 19, 638–653 (2018).

42. Smith, J. S., Lefkowitz, R. J. & Rajagopal, S. Biased signalling: from simple switches to allosteric microprocessors. Nat. Rev. Drug Discov. 17, 243–260 (2018).

43. Luscombe, V. B., Wang, P., Russell, A. J. & Greaves, D. R. Biased agonists of GPR84 and insights into biological control. Br. J. Pharmacol. 181, 1509–1523 (2024).

44. Zhang, X. et al. Allosteric modulation and biased signalling at free fatty acid receptor 2. Nature (2025) doi:10.1038/s41586-025-09186-6.

45. Zhou, Q. et al. Common activation mechanism of class A GPCRs. eLife 8, e50279 (2019).

46. Matthees, E. S. F. et al. GRK specificity and Gβγ dependency determines the potential of a GPCR for arrestin-biased agonism. *Commun*. Biol. 7, (2024).

47. Duan, J. et al. Cryo-EM structure of an activated VIP1 receptor-G protein complex revealed by a NanoBiT tethering strategy. Nat. Commun. 11, 4121 (2020).

48. Liang, Y.-L. et al. Dominant Negative G Proteins Enhance Formation and Purification of Agonist-GPCR-G Protein Complexes for Structure Determination. ACS Pharmacol. Transl. Sci. 1, 12–20 (2018).

49. Maeda, S. et al. Development of an antibody fragment that stabilizes GPCR/G-protein complexes. Nat. Commun. 9, 3712 (2018).

50. Pettersen, E. F. et al. UCSF Chimera—A visualization system for exploratory research and analysis. J. Comput. Chem. 25, 1605–1612 (2004).

51. Emsley, P. & Cowtan, K. *Coot* : model-building tools for molecular graphics. Acta Crystallogr. D Biol. Crystallogr. 60, 2126–2132 (2004).

52. Adams, P. D., et al. *PHENIX* : a comprehensive Python-based system for macromolecular structure solution. Acta Crystallogr. D Biol. Crystallogr. 66, 213–221 (2010).

53. Mahmud, Z. A. et al. Three classes of ligands each bind to distinct sites on the orphan G protein-coupled receptor GPR84. Sci. Rep. 7, 17953 (2017).

54. Schrödinger Release 2021-3: Maestro, Schrödinger, LLC, New York, NY, 2021.

55. He, X., Man, V. H., Ji, B., Xie, X.-Q. & Wang, J. Calculate protein–ligand binding affinities with the extended linear interaction energy method: application on the Cathepsin S set in the D3R Grand Challenge 3. J. Comput. Aided Mol. Des. 33, 105–117 (2019).

56. Friesner, R. A. et al. Extra Precision Glide: Docking and Scoring Incorporating a Model of Hydrophobic Enclosure for Protein−Ligand Complexes. J. Med. Chem. 49, 6177–6196 (2006).

57. Friesner, R. A. et al. Glide: A New Approach for Rapid, Accurate Docking and Scoring. 1. Method and Assessment of Docking Accuracy. J. Med. Chem. 47, 1739–1749 (2004).

58. Halgren, T. A. et al. Glide: A New Approach for Rapid, Accurate Docking and Scoring. 2. Enrichment Factors in Database Screening. J. Med. Chem. 47, 1750–1759 (2004).

59. Case, D. A. et al. Amber 2020.

60. Salomon-Ferrer, R., Götz, A. W., Poole, D., Le Grand, S. & Walker, R. C. Routine Microsecond Molecular Dynamics Simulations with AMBER on GPUs. 2. Explicit Solvent Particle Mesh Ewald. J. Chem. Theory Comput. 9, 3878–3888 (2013).

61. Götz, A. W. et al. Routine Microsecond Molecular Dynamics Simulations with AMBER on GPUs. 1. Generalized Born. J. Chem. Theory Comput. 8, 1542–1555 (2012).

62. Le Grand, S., Götz, A. W. & Walker, R. C. SPFP: Speed without compromise—A mixed precision model for GPU accelerated molecular dynamics simulations. Comput. Phys. Commun. 184, 374–380 (2013).

63. Tian, C. et al. ff19SB: Amino-Acid-Specific Protein Backbone Parameters Trained against Quantum Mechanics Energy Surfaces in Solution. J. Chem. Theory Comput. 16, 528–552 (2020).

64. Dickson, C. J., Walker, R. C. & Gould, I. R. Lipid21: Complex Lipid Membrane Simulations with AMBER. J. Chem. Theory Comput. 18, 1726–1736 (2022).

65. He, X., Man, V. H., Yang, W., Lee, T.-S. & Wang, J. A fast and high-quality charge model for the next generation general AMBER force field. J. Chem. Phys. 153, (2020).

66. Jorgensen, W. L., Chandrasekhar, J., Madura, J. D., Impey, R. W. & Klein, M. L. Comparison of simple potential functions for simulating liquid water. J. Chem. Phys. 79, 926–935 (1983).

67. Søndergaard, C. R., Olsson, M. H. M., Rostkowski, M. & Jensen, J. H. Improved Treatment of Ligands and Coupling Effects in Empirical Calculation and Rationalization of p*K*a Values. J. Chem. Theory Comput. 7, 2284–2295 (2011).

68. Olsson, M. H. M., Søndergaard, C. R., Rostkowski, M. & Jensen, J. H. PROPKA3: Consistent Treatment of Internal and Surface Residues in Empirical p*K*aPredictions. J. Chem. Theory Comput. 7, 525–537 (2011).

69. Johnston, R. C. et al. Epik: p*K*a and Protonation State Prediction through Machine Learning. J. Chem. Theory Comput. 19, 2380–2388 (2023).

70. Schrödinger Release 2021-3: Epik, Schrödinger, LLC, New York, NY, 2021.

71. Jo, S., Kim, T. & Im, W. Automated Builder and Database of Protein/Membrane Complexes for Molecular Dynamics Simulations. PLoS ONE 2, e880 (2007).

72. Lee, J. et al. CHARMM-GUI Input Generator for NAMD, GROMACS, AMBER, OpenMM, and CHARMM/OpenMM Simulations Using the CHARMM36 Additive Force Field. J. Chem. Theory Comput. 12, 405–413 (2016).

73. Kim, S. et al. CHARMM-GUI ligand reader and modeler for CHARMM force field generation of small molecules. J. Comput. Chem. 38, 1879–1886 (2017).

74. Jo, S., Lim, J. B., Klauda, J. B. & Im, W. CHARMM-GUI Membrane Builder for Mixed Bilayers and Its Application to Yeast Membranes. Biophys. J. 97, 50–58 (2009).

75. Wu, E. L. et al. CHARMM-GUI*Membrane Builder*toward realistic biological membrane simulations. J. Comput. Chem. 35, 1997–2004 (2014).

76. Jo, S., Kim, T., Iyer, V. G. & Im, W. CHARMM-GUI: A web-based graphical user interface for CHARMM. J. Comput. Chem. 29, 1859–1865 (2008).

77. Lee, J. et al. CHARMM-GUI supports the Amber force fields. J. Chem. Phys. 153, (2020).

78. Lee, J. et al. CHARMM-GUI *Membrane Builder* for Complex Biological Membrane Simulations with Glycolipids and Lipoglycans. J. Chem. Theory Comput. 15, 775–786 (2019).

79. Lomize, M. A., Pogozheva, I. D., Joo, H., Mosberg, H. I. & Lomize, A. L. OPM database and PPM web server: resources for positioning of proteins in membranes. Nucleic Acids Res. 40, D370–D376 (2012).

80. Pastor, R. W., Brooks, B. R. & Szabo, A. An analysis of the accuracy of Langevin and molecular dynamics algorithms. Mol. Phys. 65, 1409–1419 (1988).

81. Berendsen, H. J. C., Postma, J. P. M., Van Gunsteren, W. F., DiNola, A. & Haak, J. R. Molecular dynamics with coupling to an external bath. J. Chem. Phys. 81, 3684–3690 (1984).

82. El Daibani, A., et al. Molecular mechanism of biased signaling at the kappa opioid receptor. Nat. Commun. 14, 1338 (2023).

83. Jumper, J. et al. Highly accurate protein structure prediction with AlphaFold. Nature 596, 583– 589 (2021).

84. Van Der Spoel, D. et al. GROMACS: Fast, flexible, and free. J. Comput. Chem. 26, 1701–1718 (2005).

85. Bussi, G., Donadio, D. & Parrinello, M. Canonical sampling through velocity rescaling. J. Chem. Phys. 126, (2007).

86. Parrinello, M. & Rahman, A. Crystal Structure and Pair Potentials: A Molecular-Dynamics Study. Phys. Rev. Lett. 45, 1196–1199 (1980).

87. Parrinello, M. & Rahman, A. Polymorphic transitions in single crystals: A new molecular dynamics method. J. Appl. Phys. 52, 7182–7190 (1981).

88. Darden, T., York, D. & Pedersen, L. Particle mesh Ewald: An *N*⋅log(*N*) method for Ewald sums in large systems. J. Chem. Phys. 98, 10089–10092 (1993).

89. Gowers, R. et al. MDAnalysis: A Python Package for the Rapid Analysis of Molecular Dynamics Simulations. in Proceedings of the Python in Science Conference 98–105 (SciPy, Austin, Texas, 2016). doi:10.25080/majora-629e541a-00e.

90. Michaud-Agrawal, N., Denning, E. J., Woolf, T. B. & Beckstein, O. MDAnalysis: A toolkit for the analysis of molecular dynamics simulations. J. Comput. Chem. 32, 2319–2327 (2011).

91. Lomize, A. L., Todd, S. C. & Pogozheva, I. D. Spatial arrangement of proteins in planar and curved membranes by PPM 3.0. Protein Sci. 31, 209–220 (2022).

92. Harris, C. R. et al. Array programming with NumPy. Nature 585, 357–362 (2020).

93. Virtanen, P. et al. SciPy 1.0: fundamental algorithms for scientific computing in Python. Nat. Methods 17, 261–272 (2020).

94. Hunter, J. D. Matplotlib: A 2D Graphics Environment. Comput. Sci. Eng. 9, 90–95 (2007).

95. Waskom, M. seaborn: statistical data visualization. J. Open Source Softw. 6, 3021 (2021).

