## Supporting Information for "Steric control of signaling bias in the immunometabolic receptor GPR84"

Authors and affiliations:

Pinqi Wang[1,2,8], Xuan Zhang[3,8], Abdul-Akim Guseinov[4,5,8], Laura Jenkins[5,8], Carl von Hallerstein[6,8], Jonathan D. Colburn[6], Rowan Ives[1,2,7], Vincent B. Luscombe[7], Sara Marsango[5], Listiana Oktavia[1,2], Arun Raja[1,2], David R. Greaves[7], Philip C. Biggin[6], Graeme Milligan[5*], Cheng Zhang[3*], Irina G. Tikhonova[4*] & Angela J. Russell[1,2*]

[1] Department of Chemistry, Chemistry Research Laboratory, University of Oxford, Mansfield Road, Oxford OX1 3TA, U.K.

[2] Department of Pharmacology, University of Oxford, Mansfield Road, Oxford OX1 3QT, U.K.

[3] Department of Pharmacology and Chemical Biology, School of Medicine, University of Pittsburgh, Pittsburgh, PA 15213, U.S.A.

[4] School of Pharmacy, Queen's University Belfast, Belfast BT9 7BL, Northern Ireland, U.K.

[5] School of Molecular Biosciences, College of Medical, Veterinary and Life Sciences, University of Glasgow, Glasgow, G12 8QQ, U.K.

[6] Department of Biochemistry, University of Oxford, South Parks Road, Oxford, OX1 3QU, U.K.

[7] Sir William Dunn School of Pathology, University of Oxford, South Parks Road, Oxford OX1 3RE, U.K.

[8] These authors contributed equally: Pinqi Wang, Xuan Zhang, Abdul-Akim Guseinov, Laura Jenkins, Carl von Hallerstein

*

#### General information

All reactions involving moisture sensitive reagents were carried out under a nitrogen atmosphere. Anhydrous solvents were dried by passing over an activated alumina column, under an inert atmosphere, using a solvent purification system. All other solvents and reagents were used as supplied (analytical or HPLC grade) without prior purification. Flash column chromatography was performed on Kieselgel 60 silica gel (230-400 mesh particle size) on a glass column or on a Biotage SP4 automated flash column chromatography platform. NMR spectra were recorded on Bruker Advance spectrometers at 400, 500 or 600 MHz in the deuterated solvent stated at room temperature. The field was locked by external referencing to the relevant deuteron resonance. Chemical shifts (δ) are reported in parts per million (ppm) and coupling constants (*J*) are quoted in Hz. Data are reported as follows: chemical shift, multiplicity (s = singlet, d = doublet, t = triplet, q = quartet, sext = sextuplet, hept = heptet, and m = multiplet), coupling constant and integration. Low-resolution mass spectra (*m/z*) were recorded on an Agilent 1260 Infinity II with Diode Array and Single Quadrupole Detectors in solutions of MeOH. A selected peak is reported in Daltons and its intensity given as percentage of the base peak. High resolution mass spectra (HRMS) were run on a Bruker microTOF (ESI and APCI) or on a Waters GCT (EI). Experiments conducted in CROs used their standard equipment. The purity for test compounds was determined by HPLC on a Shimadzu SIL-20AC HT instrument. HPLC conditions were as follows: Atlantis dC18, 100 Å, 5 μm, Column 4.6 × 150 mm, 35−100% MECN in water, 15 min run, flow rate 1.5 mL/min, UV detection (λ = 220, 254, 280 nm). The mass spectra were obtained using Agilent 6120 bench-top single quadrupole instrument using electrospray ionization (ESI).

#### Reaction Schemes

**Scheme 1.** Synthesis of **OX04539**[a]

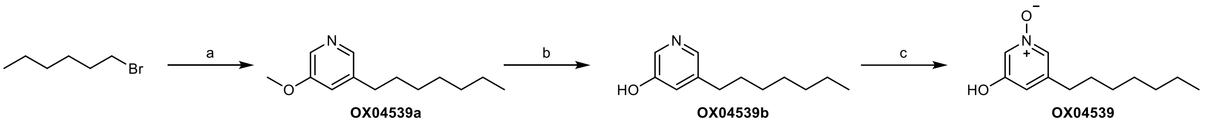


[a]Reagents and conditions: (a) (1) $PPh_3$, Toluene, reflux, 16h, 63%; (2) 5-methoxynicotinaldehyde, NaOtBu, dichloromethane, 0ºC–rt, 91%; (3) Pd/C, $H_2$, MeOH, rt, 1 h, 99%; (b) L-selectride, THF, 70ºC, 72 h, 39%; (c) *m*-CPBA, $CH_2Cl_2$, rt, 2 h, 55%.

**Scheme 2.** Synthesis of **OX04540**[a]

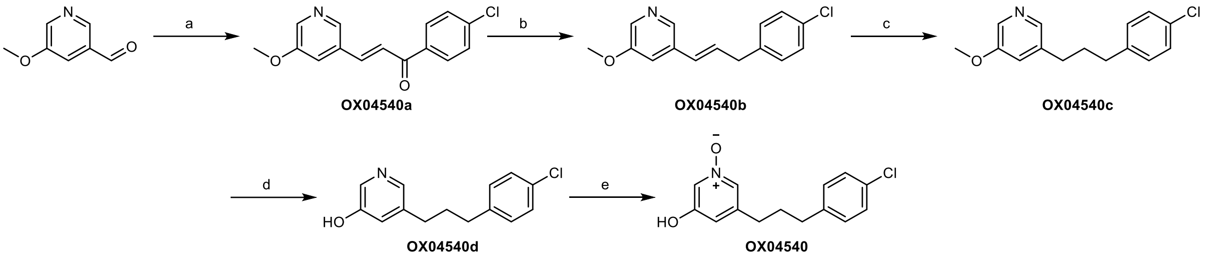


[a]Reagents and conditions: (a) 1-(4-chlorophenyl)ethan-1-one, $BF_3 \cdot Et_2O$, 1,4-dioxane, 105ºC, 16 h, 70%; (b) $Et_3SiH$, TFA, rt, 48 h, 59%; (c) Pd/C, $H_2$, MeOH, rt, 1 h, 79%; (d) 48% HBr, 120ºC, 48 h, 23%; (e) *m*-CPBA, $CH_2Cl_2$, rt, 2 h, 28%.

**Scheme 3.** Synthesis of **OX04954**[a]

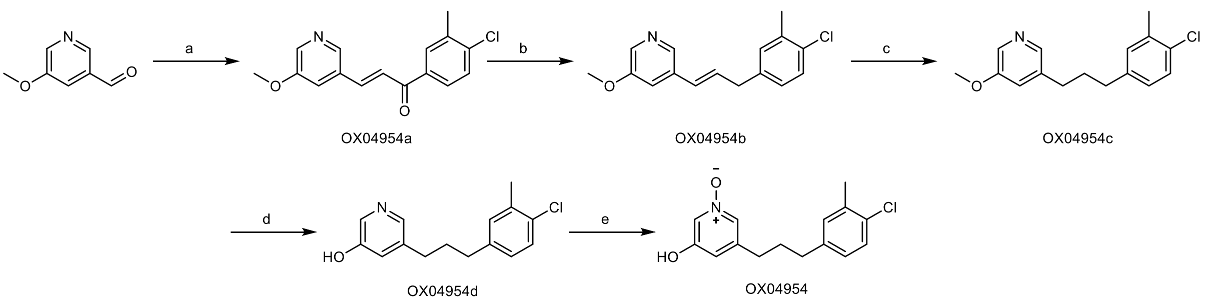


[a]Reagents and conditions: (a) 1-(4-chloro-3-methylphenyl)ethan-1-one, $BF_3 \cdot Et_2O$, 1,4-dioxane, 105ºC, 16 h, 85%; (b) $Et_3SiH$, TFA, rt, 48 h, 82%; (c) Pd/C, $H_2$, MeOH, rt, 30 min, 85%; (d) 48% HBr, 120ºC, 48 h, 79%; (e) *m*-CPBA, $CH_2Cl_2$, rt, 2 h, 87%.

**Scheme 4.** Synthesis of **OX04956**[a]

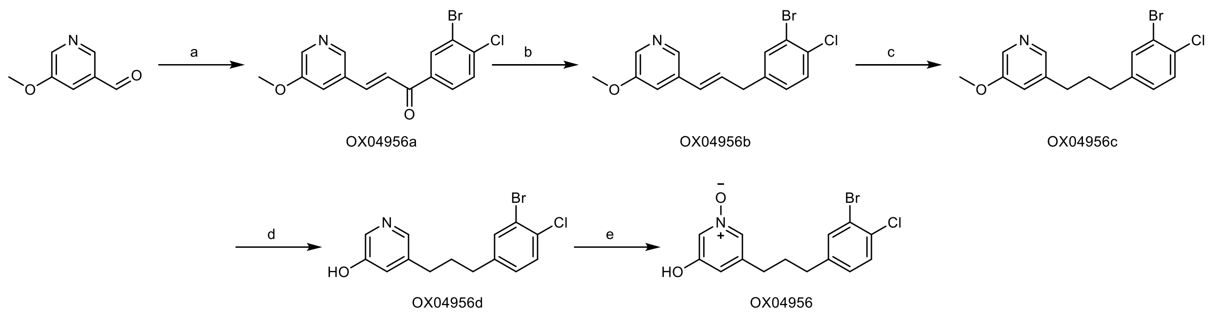


[a]Reagents and conditions: (a) 1-(3-bromo-4-chlorophenyl)ethan-1-one, $BF_3 \cdot Et_2O$, 1,4-dioxane, 105ºC, 16 h, 29%; (b) $Et_3SiH$, TFA, rt, 48 h, 59%; (c) $PtO_2$, $H_2$, MeOH, rt, 1 h, 85%; (d) 48% HBr, 120ºC, 48 h, 28%; (e) *m*-CPBA, $CH_2Cl_2$, rt, 2 h, 52%.

**Scheme 5.** Synthesis of **OX04955**[a]

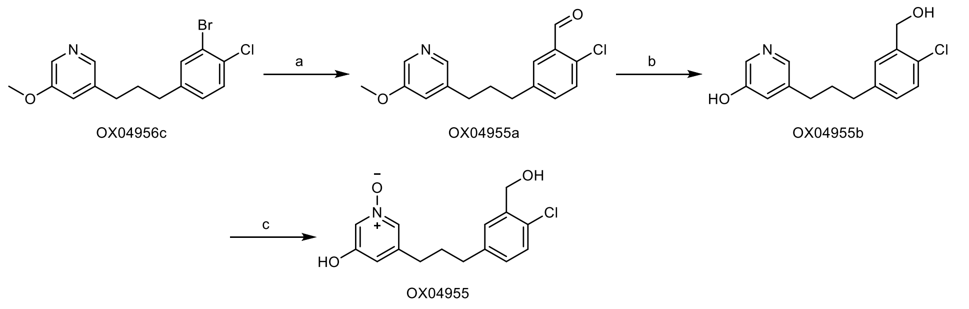


[a]Reagents and conditions: (a) (1) n-BuLi, THF, -78ºC, 5 min; (2) DMF, THF, -78ºC, 15 min, 45%; (b) L-Selectride, THF, 70ºC, 48 h, 39%; (c) *m*-CPBA, $CH_2Cl_2$, rt, 2 h, 64%.

**Scheme 6.** Synthesis of **OX04957**[a]

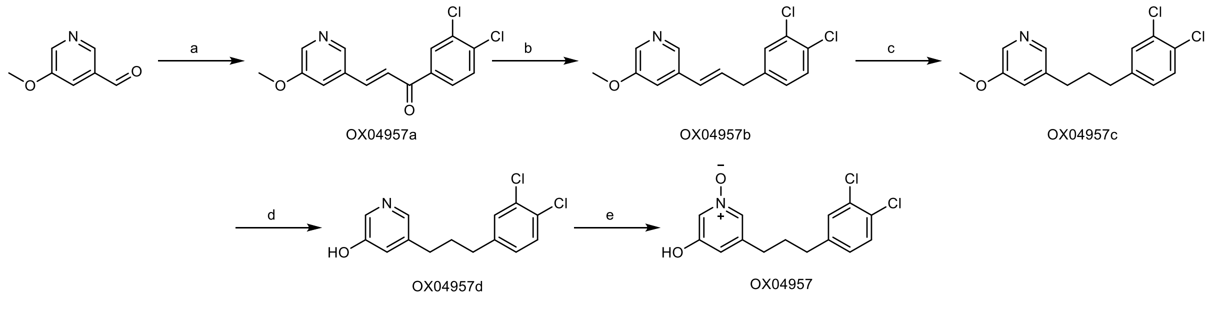


[a]Reagents and conditions: (a) 1-(3,4-dichlorophenyl)ethan-1-one, $BF_3 \cdot Et_2O$, 1,4-dioxane, 105ºC, 16 h, 67%; (b) $Et_3SiH$, TFA, rt, 48 h, 38%; (c) PtO2, $H_2$, MeOH, rt, 1 h, 76%; (d) 48% HBr, 120ºC, 48 h, 46%; (e) *m*-CPBA, $CH_2Cl_2$, rt, 2.5 h, 23%.

**Scheme 7.** Synthesis of **OX04958**[a]

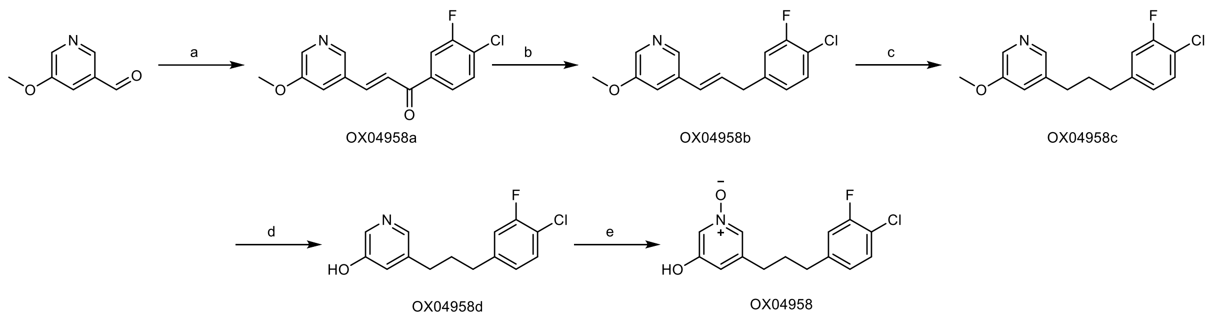


[a]Reagents and conditions: (a) 1-(4-chloro-3-fluorophenyl)ethan-1-one, $BF_3\cdot Et_2O$, 1,4-dioxane, 105ºC, 16 h, 64%; (b) $Et_3SiH$, TFA, rt, 48 h, 38%; (c) PtO2, $H_2$, MeOH, rt, 1 h, 93%; (d) 48% HBr, 120ºC, 48 h, 18%; (e) *m*-CPBA, $CH_2Cl_2$, rt, 2.5 h, 65%.

**Scheme 8.** Synthesis of **OX04959**[a]

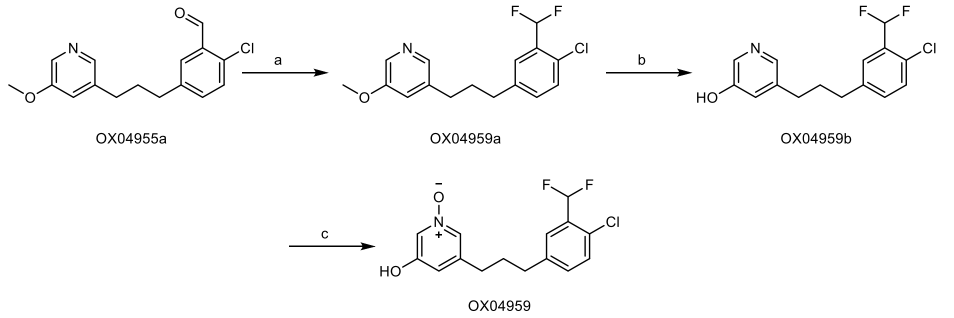


[a]Reagents and conditions: (a) DAST, $CH_2Cl_2$, rt, 16 h, 37%; (b) L-selectride, THF, 70ºC, 48 h, 92%; (c) *m*-CPBA, $CH_2Cl_2$, rt, 2 h, 43%.

**Scheme 9.** Synthesis of **OX04960**[a]

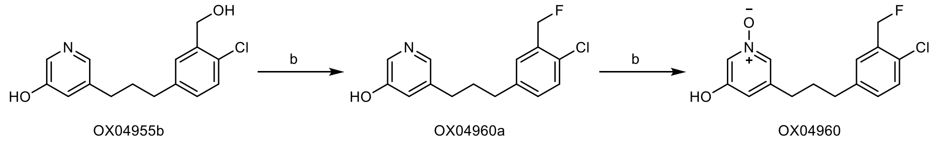


[a]Reagents and conditions: (a) DAST, $CH_2Cl_2$, rt, 24 h, 11%; (b) *m*-CPBA, $CH_2Cl_2$, rt, 2 h, 44%.

**Scheme 10.** Synthesis of **OX04961**[a]

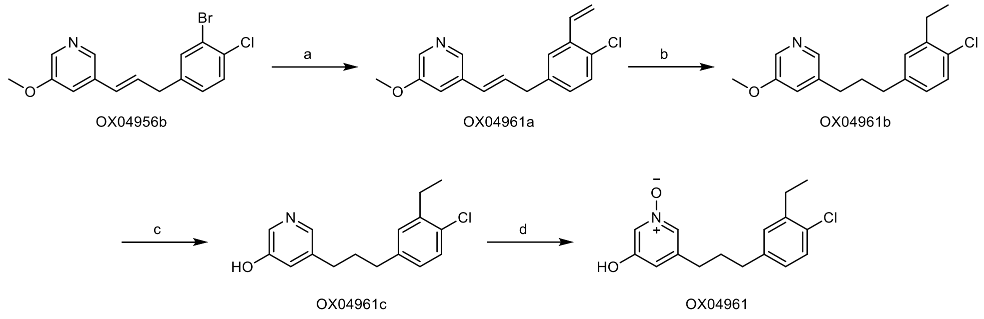


[a]Reagents and conditions: (a) vinyl boronic ester, $Cs_2CO_3$, $PPh_3$, $Pd(OAc)_2$, THF:$H_2O$ 9:1, 70ºC, 25h, 56%; (b) Pd/C, $H_2$, MeOH, rt, 1 h, 39%; (c) 48% HBr, 120ºC, 48 h, 55%; (d) *m*-CPBA, $CH_2Cl_2$, rt, 2 h, 41%.

**Scheme 11.** Synthesis of **OX04962**[a]

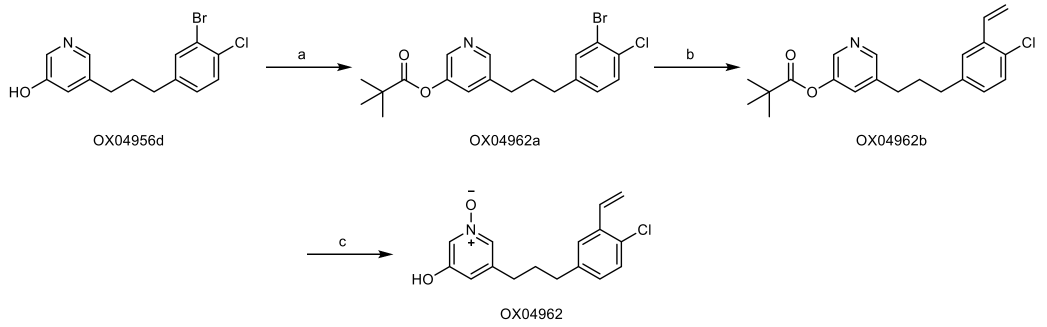


[a]Reagents and conditions: (a) trimethylacetyl chloride, $NEt_3$, $CH_2Cl_2$, rt, 1 h, 98%; (b) tributyl(vinyl)stannane, $Pd(PPh_3)_4$, DMF, 90ºC, 24 h, 34%; (c) (1) *m*-CPBA, $CH_2Cl_2$, rt, 2 h; (2) NaOH, MeOH, rt, 16h, 34%.

**Scheme 12.** Synthesis of **OX04963**[a]

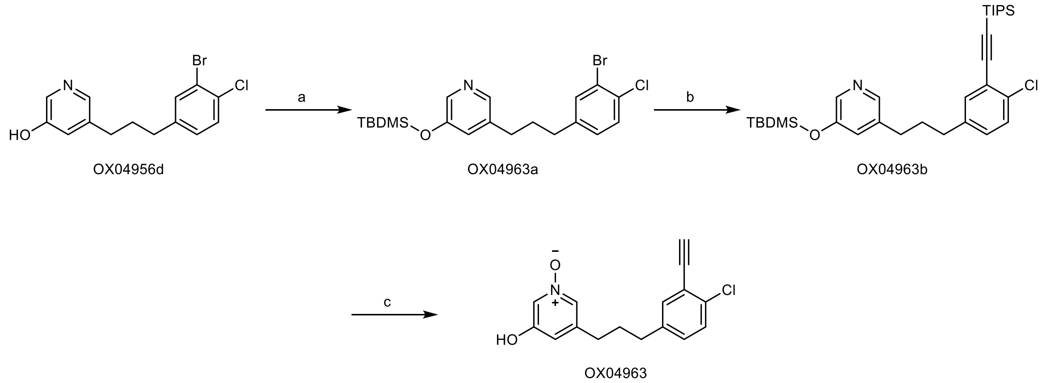


[a]Reagents and conditions: (a) TBDMSCl, imidazole, $CHCl_3$, 60ºC, 16 h, 26%; (b) ethynyltriisopropylsilane, $PPh_3$, $Pd_2(PPh_3)_2Cl_2$, CuI, THF:$NEt_3$ 3:1, 60ºC, 16 h, 69%; (c) (1) *m*-CPBA, $CH_2Cl_2$, rt, 2 h; (2) TBAF, THF, rt, 16h, 22%.

**Scheme 13.** Synthesis of **OX04964**[a]

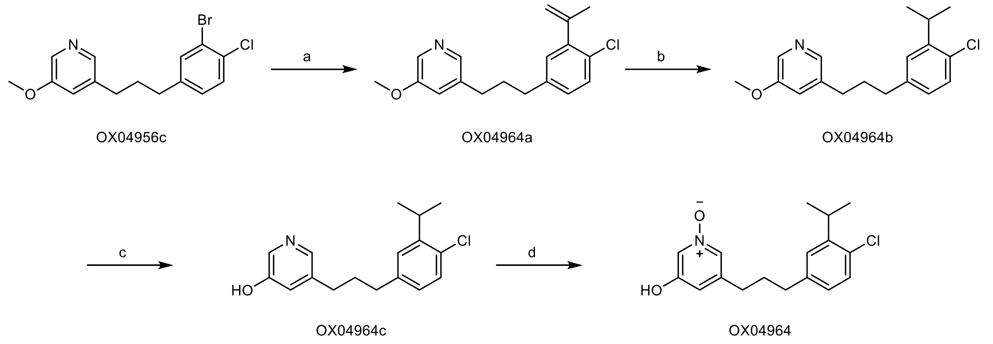


[a]Reagents and conditions: (a) tributyl(prop-1-en-2-yl)stannane, $Pd(PPh_3)_4$, DMF, 90ºC, 24 h, 39%; (b) $PtO_2$, $H_2$, MeOH, rt, 1 h, 64%; (c) 48% HBr, 120ºC, 48 h, 60%; (d) *m*-CPBA, $CH_2Cl_2$, rt, 2 h, 60%.

**Scheme 14.** Synthesis of **OX04965**[a]

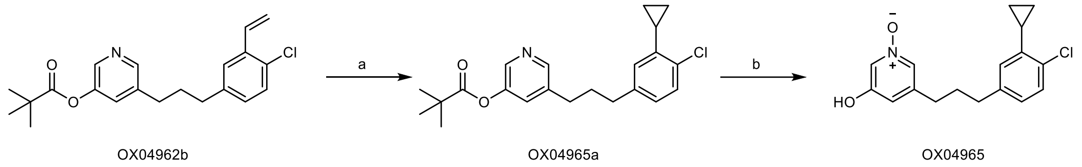


[a]Reagents and conditions: (a) (1) $Et_2Zn$, TFA, CH2Cl2, 0ºC, 30 min; (2) diiodomethane, $CH_2Cl_2$, 0ºC–rt, 16 h, 92%; (b) (1) *m*-CPBA, $CH_2Cl_2$, rt, 2 h; (2) NaOH, MeOH, rt, 16h, 86%.

**Scheme 15.** Synthesis of **OX04966**[a]

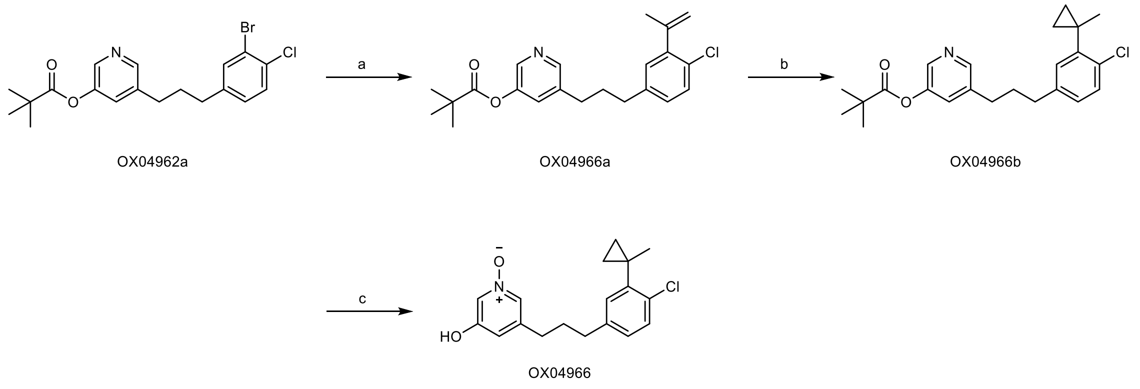


[a]Reagents and conditions: (a) tributyl(vinyl)stannane, $Pd(PPh_3)_4$, DMF, 90ºC, 24 h, 89%; (b) (1) $Et_2Zn$, TFA, $CH_2Cl_2$, 0ºC, 30 min; (2) diiodomethane, $CH_2Cl_2$, 0ºC–rt, 16 h, 31%; (c) (1) *m*-CPBA, $CH_2Cl_2$, rt, 2 h; (2) NaOH, MeOH, rt, 16h, 99%.

**OX04539a**

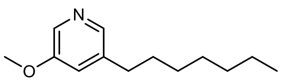

OX04539a

(1) To a solution of 1-bromohexane (825 mg, 5.0 mmol) in 20 mL toluene was added $PPh_3$ (1443 mg, 5.5 mmol), and the mixture was refluxed for 16h. After completion, the mixture was concentrated under vacuum, redissolved in $CH_2Cl_2$ and purified by flash column chromatography (0–10% MeOH in $CH_2Cl_2$) to afford hexyltriphenylphosphonium bromide as a white solid (1351 mg, 63%).

(2) The solid (1042mg, 3.0 mmol) was then dissolved in anhydrous $CH_2Cl_2$ (30 mL), NaOtBu (577 mg, 6.0 mmol) was added at 0ºC in one portion and the mixture was stirred for 15 minutes before adding 5-methoxynicotinaldehyde (274 mg, 2.0 mmol). The mixture was allowed to ward to rt and stirred for 16 h. After completion, the reaction mixture was diluted with $CH_2Cl_2$, washed with aq. sat. $NH_4Cl$ solution, brine, dried ($Na_2SO_4$) and concentrated *in vacuo*. Purification by flash column chromatography (0–30% EtOAc in Pentane) afforded 3-heptyl-5-methoxypyridine as a transparent oil (400 mg, 91%).

(3) The oil (400 mg) was then dissolved in MeOH and the solution was degassed with $N_2$. 40 mg Pd/C (10 wt.%) was added and the atmosphere was replaced with $H_2$. The reaction was stirred for 1h, filtered through Celite®, and dried under vacuum to afford **OX04539a** as a transparent oil (400 mg, 99%). $^{1}$H NMR (400 MHz, $CDCl_3$) δ 8.12 (d, *J* = 2.8 Hz, 1H), 8.05 (d, *J* = 1.8 Hz, 1H), 6.99 (dd, *J* = 2.8, 1.8 Hz, 1H), 3.83 (s, 3H), 2.57 (dd, *J* = 8.7, 6.8 Hz, 2H), 1.66–1.54 (m, 2H), 1.37–1.19 (m, 8H), 0.90–0.83 (m, 3H). $^{13}$C NMR (101 MHz, $CDCl_3$) δ 155.7, 142.6, 138.9, 134.7, 120.8, 55.6, 32.0, 31.9, 31.2, 29.3, 29.2, 22.7, 14.2. HRMS (ESI +ve) $C_{13}H_{21}NO$ $[M+H]^+$ calc. 208.1696, found 208.1693.

**OX04539b**

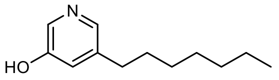

OX04539b

The demethylation using L-Selectride was adapted from a reported route.[1] L-Selectride (6 mL, 1 M in THF) was added to **OX04539a** (400 mg, 1.90 mmol) in THF (10 mL) and then the mixture was stirred at 70ºC for 48 h. After completion, the mixture was cooled to rt, quenched with MeOH, stirred for 30 min and concentrated *in vacuo*. The residue was then purified by flash column chromatography (0–8% MeOH in $CH_2Cl_2$) to afford **OX04539b** as a yellow oil (146 mg, 39%). $^{1}$H NMR (400 MHz, $CDCl_3$) δ 8.73 (d, *J* = 2.5 Hz, 1H), 8.04 (d, *J* = 1.6 Hz, 1H), 7.89 (t, *J* = 2.0 Hz, 1H), 2.72 (t, *J* = 7.8 Hz, 2H), 1.71–1.58 (m, 2H), 1.50–1.02 (m, 8H), 0.90–0.79 (m, 3H). $^{13}$C NMR (101 MHz, $CDCl_3$) δ 157.0, 144.1, 133.4, 130.9, 126.9, 32.9, 31.7, 30.4, 29.0, 29.0, 22.6, 14.1. HRMS (ESI +ve) $C_{12}H_{19}NO$ $[M+H]^+$ calc. 194.1539, found 194.1534.

**OX04539**

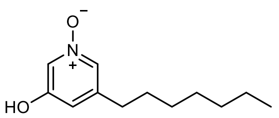

OX04539

To a solution of **OX04539b** (146 mg, 0.76 mmol) in $CH_2Cl_2$ (20 mL) at rt was added *m*-CPBA (1.5 eq.) and the resulting mixture stirred for 2 h. After completion, the reaction was quenched with $NaHCO_3$ (sat, aq.). The organic phase was washed with brine, dried ($Na_2SO_4$), filtered and concentrated *in vacuo*. Purification by flash column chromatography (0–6% MeOH in $CH_2Cl_2$) afforded **OX04539** as a yellow oil (87 mg, 55%). $^1$H NMR (400 MHz, MeOD) δ 7.67 (q, *J* = 1.7 Hz, 2H), 6.87 (t, *J* = 1.6 Hz, 1H), 2.53–2.45 (m, 2H), 1.58–1.46 (m, 2H), 1.25–1.20 (m, 8H), 0.84–0.76 (m, 3H). $^{13}$C NMR (101 MHz, MeOD) δ 157.8, 144.1, 131.8, 126.9, 119.5, 33.5, 32.9, 31.6, 30.1, 30.0, 23.7, 14.4. HRMS (ESI +ve) $C_{12}H_{19}NO_2$ [M+H$^+$] calc. 210.1489, found 210.1487. HPLC 98% (AUC), $t_R$ = 5.1 min.

**OX04540a**

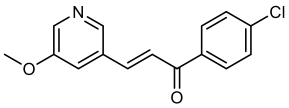

OX04540a

To a solution of 5-methoxynicotinaldehyde (500 mg, 3.65 mmol) and 1-(4-chlorophenyl)ethan-1-one (562 mg, 3.65 mmol) in 1,4-dioxane (10 mL) was added $BF_3{\cdot}Et_2O$ (1.9 mL) under $N_2$ and then the mixture was stirred at 105ºC for 16 h. After completion of the reaction, cooled to rt, the mixture was diluted with sat. aq. $NaHCO_3$ (50 mL) and extracted with EtOAc (3 × 30 mL). The combined organic extracts were washed with brine (100 mL), dried over $Na_2SO_4$, filtered, and concentrated under reduced pressure. The resulting residue was chromatographed over silica gel (0 − 60% EtOAc in pentane) to give **OX04540a** as a white solid (700 mg, 70%). $^1$H NMR (400 MHz, $CDCl_3$) δ 8.46 (d, *J* = 1.7 Hz, 1H), 8.35 (d, *J* = 2.8 Hz, 1H), 8.00–7.93 (m, 2H), 7.77 (d, *J* = 15.8 Hz, 1H), 7.56–7.46 (m, 3H), 7.39 (dd, *J* = 2.8, 1.8 Hz, 1H), 3.92 (s, 3H). $^{13}$C NMR (101 MHz, $CDCl_3$) δ 188.7, 155.9, 142.6, 141.6, 139.8, 139.7, 136.2, 131.2, 130.1, 129.2, 123.6, 118.4, 77.5, 77.2, 76.8, 55.9. **HRMS (ESI+ve)** $C_{15}H_{12}ClNO_2$ [M+H$^+$] calc. 274.0629, found 274.0623.

**OX04540b**

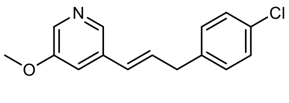

OX04540b

**OX04540a** (273 mg, 1.0 mmol) was dissolved in TFA (6 mL) using sonication. Triethylsilane (12 mL) was added and the reaction was vigorously stirred for 48h. Upon completion, the reaction mixture was poured into sat. aq. $NaHCO_3$ and stirred until bubbling ceased. This solution was taken up into a separatory funnel and extracted with $CH_2Cl_2$ (3 × 40 mL), washed with brine, and dried under $Na_2SO_4$. Organic extracts were dried under vacuum. Purification by flash column chromatography (30% EtOAc in Pentane) afforded

**OX04540b** as a transparent oil (627 mg, 59%). $^1$H NMR (400 MHz, $CDCl_3$) δ 8.10 (dd, *J* = 8.6, 2.3 Hz, 2H), 7.25–7.17 (m, 2H), 7.11–7.06 (m, 3H), 6.34–6.29 (m, 2H), 3.77 (s, 3H), 3.48–3.43 (m, 2H). $^{13}$C NMR (101 MHz, $CDCl_3$) δ 155.8, 140.8, 138.0, 136.5, 133.5, 132.3, 131.4, 130.1, 129.8, 128.8, 128.6, 127.9, 116.8, 55.6, 38.7.

**OX04540c**

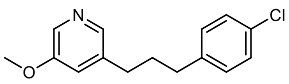


OX04540c

40 mg Pd/C (10% M/w) was added in one portion to a $N_2$ degassed solution of the **OX04540b** (400 mg, 1.5 mmol) in MeOH (50 mL) at rt, the atmosphere was replaced with $H_2$, and the reaction stirred for 1 h. After completion, the reaction was filtered through Celite® using MeOH as an eluent, concentrated *in vacuo* and purified by flash column chromatography (30% EtOAc in Pentane) to afford the **OX04540c** as a yellow oil (320 mg, 79%). $^1$H NMR (400 MHz, $CDCl_3$) δ 8.15 (d, *J* = 2.8 Hz, 1H), 8.06 (d, *J* = 1.8 Hz, 1H), 7.28–7.21 (m, 2H), 7.13–7.05 (m, 2H), 6.98 (dd, *J* = 2.7, 1.8 Hz, 1H), 3.84 (s, 3H), 2.61 (td, *J* = 7.7, 3.0 Hz, 4H), 1.99–1.87 (m, 2H). $^{13}$C NMR (101 MHz, $CDCl_3$) δ 155.7, 142.5, 140.2, 138.0, 135.0, 131.8, 129.9, 128.6, 120.8, 55.6, 34.7, 32.5, 32.3. **HRMS (ESI+ve)** $C_{15}H_{16}ClNO$ [M+H$^+$] calc. **262.0993**, found **262.0990**.

**OX04540d**

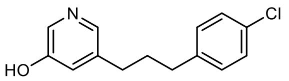


OX04540d

3 mL 48% HBr in $H_2O$ was added to **OX04540c** (100 mg, 0.4 mmol) and then the mixture was stirred at 120ºC for 48 h. After completion of the reaction, cooled to rt, the mixture was neutralized with sat. aq. $NaHCO_3$ and extracted with EtOAc (3 × 30 mL). The combined organic extracts were washed with brine (100 mL), dried over $Na_2SO_4$, filtered, and concentrated under reduced pressure. The resulting residue was purified by flash column chromatography (5% MeOH in $CH_2Cl_2$) afforded the **OX04540d** as a transparent oil (22 mg, 23%). $^1$H NMR (400 MHz, $CDCl_3$) δ 8.27–8.21 (m, 1H), 8.02 (s, 1H), 7.88 (s, 1H), 7.39 (t, *J* = 2.0 Hz, 1H), 7.26–7.22 (m, 2H), 7.11–7.06 (m, 2H), 2.63 (q, *J* = 7.1 Hz, 4H), 1.92 (dq, *J* = 9.3, 7.7 Hz, 2H). $^{13}$C NMR (101 MHz, $CDCl_3$) δ 155.2, 140.1, 139.7, 139.6, 134.1, 131.8, 129.9, 128.7, 125.1, 34.7, 32.3, 32.2. HRMS (ESI+ve) $C_{14}H_{14}ClNO$ [M+H$^+$] calc. 248.0837, found 248.0835.

**OX04540**

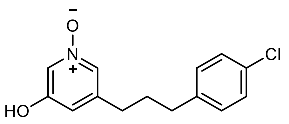


OX04540

To a solution of **OX04540d** (20 mg, 0.08 mmol) in $CH_2Cl_2$ (10 mL) at rt was added *m*-CPBA (1.5 eq.) and the resulting mixture stirred for 2 h. After completion, the reaction was quenched with $NaHCO_3$ (sat, aq.). The organic phase was washed with brine, dried ($Na_2SO_4$), filtered and concentrated *in vacuo*. Purification by flash column chromatography (0–100% EtOAc in Pentane then 0–6% MeOH in $CH_2Cl_2$) afforded **OX04540** as a yellow oil (6 mg, 28%). $^1H$ NMR (600 MHz, $CDCl_3$) δ 8.06 (t, *J* = 1.9 Hz, 1H), 7.63 (t, *J* = 1.5 Hz, 1H), 7.26–7.23 (m, 2H), 7.10–7.05 (m, 2H), 6.98 (t, *J* = 1.7 Hz, 1H), 2.60 (t, *J* = 7.6 Hz, 2H), 2.56–2.50 (m, 2H), 1.89 (tt, *J* = 9.2, 6.8 Hz, 2H). $^{13}C$ NMR (151 MHz, $CDCl_3$) δ 157.6, 141.7, 139.6, 132.1, 129.8, 129.7, 128.8, 126.6, 120.7, 34.5, 32.2, 31.7. **HRMS (ESI +ve)** $C_{14}H_{14}ClNO_2$ [$M+H^+$] calc. **264.0786**, found 2**64.0784**. HPLC 98% (AUC), $t_R$ = 6.3 min.

**OX04954a**

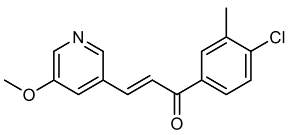


OX04954a

To a solution of 5-methoxynicotinaldehyde (609 mg, 4.45 mmol) and 1-(4-chloro-3-methylphenyl)ethan-1-one (500 mg, 2.96 mmol) in 1,4-dioxane (8 mL) was added $BF_3{\cdot}Et_2O$ (1.5 mL) under $N_2$ and then the mixture was stirred at 105°C for 16 h. After completion of the reaction, cooled to rt, the mixture was diluted with sat. aq. $NaHCO_3$ (50 mL) and extracted with EtOAc (3 × 30 mL). The combined organic extracts were washed with brine (100 mL), dried over $Na_2SO_4$, filtered, and concentrated under reduced pressure. The resulting residue was chromatographed over silica gel (0 − 60% EtOAc in pentane) to give **OX04954a** as a white solid (720 mg, 85%). $^1H$ NMR (400 MHz, $CDCl_3$) δ 8.47 (d, J = 1.7 Hz, 1H), 8.35 (d, J = 2.8 Hz, 1H), 7.89 (dd, J = 2.2, 0.9 Hz, 1H), 7.81–7.73 (m, 2H), 7.52 (d, J = 15.7 Hz, 1H), 7.48 (d, J = 8.3 Hz, 1H), 7.43–7.37 (m, 1H), 3.93 (s, 3H), 2.47 (s, 3H). $^{13}C$ NMR (101 MHz, $CDCl_3$) δ 189.0, 155.9, 142.5, 141.4, 139.9, 139.5, 137.0, 136.3, 131.3, 131.1, 129.6, 127.4, 123.8, 118.6, 55.9, 20.3. **LRMS** (ESI +ve) 288 [M+H]+. **HRMS** (ESI +ve) $C_{16}H_{14}ClNO_2$ [$M+H^+$] calc. 288.0822, found 288.0786.

**OX04954b**

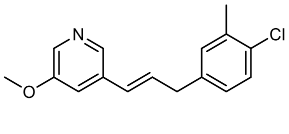


OX04954b

To a solution of **OX04954a** (700 mg, 2.43 mmol) in TFA (3.0 mL) was added Triethylsilane (3.1 mL) under $N_2$ and then the mixture was stirred at room temperature for 48 h. After completion of the reaction, the mixture was diluted with $CH_2Cl_2$ (50 mL), quenched with sat. aq. 1N NaOH (30 mL) and extracted with $CH_2Cl_2$ (3 × 30 mL). The combined organic extracts were washed with brine (100 mL), dried over $Na_2SO_4$, filtered, and concentrated under reduced pressure. The resulting residue was chromatographed over silica gel (0 − 60% EtOAc in pentane) to give **OX04954b** as a colourless oil (540 mg, 82%). $^1H$ NMR (400 MHz,

$CDCl_3$) δ 8.17 (dd, J = 10.9, 2.3 Hz, 2H), 7.27 (d, J = 8.1 Hz, 1H), 7.15 (dd, J = 2.8, 1.8 Hz, 1H), 7.08 (d, J = 2.2 Hz, 1H), 6.99 (dd, J = 8.1, 2.3 Hz, 1H), 6.41–6.36 (m, 2H), 3.84 (s, 3H), 3.55–3.44 (m, 2H), 2.36 (s, 3H). $^{13}$C NMR (101 MHz, $CDCl_3$) δ 155.8, 140.8, 138.1, 136.5, 136.2, 133.6, 132.4, 131.6, 131.4, 129.2, 127.7, 127.5, 116.8, 55.6, 38.8, 20.1. LRMS (ESI +ve) 274 [M+H]+. HRMS (ESI +ve) $C_{16}H_{16}ClNO$ [M+H$^+$] calc. 274.0993, found 274.1040.

**OX04954c**

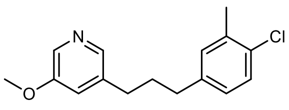

OX04954c

A mixture of **OX04954b** (500 mg, 1.82 mmol) and 50 mg $PtO_2$ (10% w/w) in MeOH (10 mL) was stirred at rt under an atmosphere of $H_2$ gas via balloon (1 atm of pressure) for 30 min. The mixture was filtered through a pad of Celite® with repeat rinsing using anhydrous MeOH. The filtrate was concentrated under reduced pressure, and the resulting crude residue was chromatographed over silica gel (0 − 40% EtOAc in pentane) to give **OX04954c** as a white solid (430 mg, 85%). $^1$H NMR (400 MHz, $CDCl_3$) δ 8.14 (d, J = 2.8 Hz, 1H), 8.05 (d, J = 1.8 Hz, 1H), 7.22 (d, J = 8.1 Hz, 1H), 7.03–6.96 (m, 2H), 6.92 (dd, J = 8.2, 2.2 Hz, 1H), 3.83 (s, 3H), 2.59 (dt, J = 10.4, 7.6 Hz, 4H), 2.33 (s, 3H), 2.01–1.85 (m, 2H). $^{13}$C NMR (101 MHz, $CDCl_3$) δ 155.7, 142.4, 140.3, 138.0, 135.9, 134.9, 131.8, 131.1, 129.0, 127.2, 120.7, 55.5, 34.6, 32.5, 32.3, 20.1. **LRMS** (ESI +ve) 276 [M+H]+. **HRMS** (ESI +ve) $C_{16}H_{18}ClNO$ [M+H$^+$] calc. 276.1150, found 276.1181.

**OX04954d**

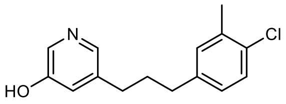

OX04954d

To 3-(3-(4-chloro-3-methylphenyl)propyl)-5-methoxypyridine **OX04954c (**250 mg, 0.906 mmol) in a vial was added 48% HBr in water (5 mL) and then the vial was sealed and the mixture was stirred at 120ºC for 48 h. After completion of the reaction, cooled to rt, the mixture was neutralized with sat. aq. $NaHCO_3$ and extracted with EtOAc (3 × 30 mL). The combined organic extracts were washed with brine (100 mL), dried over $Na_2SO_4$, filtered, and concentrated under reduced pressure. The resulting residue was chromatographed over silica gel (0 − 10% MeOH in $CH_2Cl_2$) to give **OX04954d** as a white solid (187 mg, 79%). $^1$H NMR (400 MHz, Acetone) δ 8.84 (s, 1H), 8.00 (dd, J = 34.7, 2.3 Hz, 2H), 7.31–7.13 (m, 2H), 7.11–7.00 (m, 2H), 2.61 (td, J = 7.8, 3.8 Hz, 4H), 2.32 (s, 3H), 1.97–1.88 (m, 2H). $^{13}$C NMR (101 MHz, Acetone) δ 154.4, 142.0, 142.0, 139.1, 136.6, 136.3, 132.1, 132.0, 129.6, 128.3, 122.7, 35.2, 33.5, 32.7, 20.0.. **LRMS** (ESI +ve) 262 [M+H]+. **HRMS** (ESI +ve) $C_{15}H_{16}ClNO$ [M+H$^+$] calc. 262.0993, found 262.1028

**OX04954**

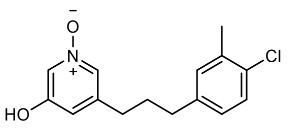

OX04954

To a solution of **OX04954a** (150 mg, 0.573 mmol) in $CH_2Cl_2$ (10 mL) at rt was added *m*-CPBA (1.5 eq.) and the resulting mixture stirred for 2 h. After completion, the reaction was quenched with $NaHCO_3$ (sat, aq.). The organic phase was washed with brine, dried ($Na_2SO_4$), filtered and concentrated *in vacuo*. Purification by flash column chromatography (0–6% MeOH in $CH_2Cl_2$) afforded **OX04954** as a white solid (138 mg, 87%). $^1$H NMR (400 MHz, $CDCl_3$) δ 7.99 (t, J = 1.9 Hz, 1H), 7.62 (t, J = 1.5 Hz, 1H), 7.22 (d, J = 8.1 Hz, 1H), 7.00 (d, J = 2.2 Hz, 1H), 6.94–6.85 (m, 2H), 2.53 (dt, J = 16.2, 7.7 Hz, 4H), 2.37 (s, 3H), 1.95–1.81 (m, 2H). $^{13}$C NMR (101 MHz, $CDCl_3$) δ 157.5, 141.6, 139.7, 136.1, 132.1, 131.1, 129.8, 129.1, 127.2, 126.6, 119.8, 34.4, 32.2, 31.8, 20.1. **LRMS** (ESI +ve) 278 [M+H]+. **HRMS** (ESI +ve) $C_{15}H_{16}ClNO_2$ [M+H$^+$] calc. 278.0942, found 278.0992. HPLC 97% (AUC), $t_R$ = 7.4 min.

**OX04956a**

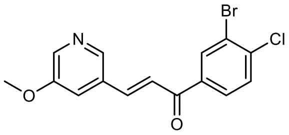

OX04956a

To a solution of 5-methoxynicotinaldehyde (1181 mg, 8.62 mmol) and 1-(3-bromo-4-chlorophenyl)ethan-1-one (1000 mg, 4.31 mmol) in 1,4-dioxane (10 mL) was added $BF_3{\cdot}Et_2O$ (2.2 mL) under $N_2$ and then the mixture was stirred at 105°C for 16 h. After completion of the reaction, cooled to rt, the mixture was diluted with sat. aq. $NaHCO_3$ (50 mL) and extracted with EtOAc (3 × 30 mL). The combined organic extracts were washed with brine (100 mL), dried over $Na_2SO_4$, filtered, and concentrated under reduced pressure. The resulting residue was chromatographed over silica gel (0 − 60% EtOAc in pentane) to give **OX04956a** as a white solid (1168 mg, 29%). $^1$H NMR (400 MHz, DMSO) δ 8.79 (d, *J* = 1.7 Hz, 1H), 8.52 (dd, *J* = 5.8, 2.4 Hz, 2H), 8.27 (dd, *J* = 2.7, 1.7 Hz, 1H), 8.22–8.14 (m, 2H), 7.90–7.80 (m, 2H), 3.97 (s, 3H). $^{13}$C NMR (101 MHz, DMSO) δ 186.7, 156.4, 140.2, 140.2, 138.5, 137.2, 136.4, 133.7, 132.4, 131.1, 129.3, 124.8, 122.4, 122.3, 56.5, 39.9, 39.7, 39.5, 39.3, 39.1.

**OX04956b**

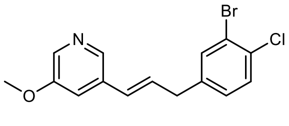

OX04956b

**OX04956a** (1100 mg, 3.1 mmol) was dissolved in TFA (10 mL) using sonication. Triethylsilane (20 mL) was added and the reaction was vigorously stirred for 48h. Upon completion, the reaction mixture was poured into sat. aq. $NaHCO_3$ and stirred until bubbling ceased. This solution was taken up into a separatory funnel and extracted with $CH_2Cl_2$ (3 × 70 mL), washed with brine, and dried under $Na_2SO_4$. Organic extracts were dried under vacuum. Purification by flash column chromatography (30% EtOAc in Pentane) afforded **OX04956b** as a transparent oil (627 mg, 59%). $^1$H NMR (400 MHz, $CDCl_3$) δ 8.17 (dd, *J* = 6.7, 2.3 Hz, 2H), 7.48 (d, *J* = 2.0 Hz, 1H), 7.37 (d, *J* = 8.2 Hz, 1H), 7.15 (t, *J* = 2.3 Hz, 1H), 7.10 (dd, *J* = 8.2, 2.1 Hz, 1H), 6.43–6.29 (m, 2H), 3.84 (s, 3H), 3.50 (d, *J* = 6.1 Hz, 2H). $^{13}$C NMR (101 MHz, $CDCl_3$) δ 155.8, 140.8, 139.9, 136.6, 133.9, 133.3, 132.5, 130.5, 130.4, 128.9, 128.4, 122.6, 116.9, 55.6, 38.4. **HRMS (ESI+ve)** $C_{15}H_{13}BrClNO$ [M+H$^+$] calc. **337.9942**, found **337.9936**.

**OX04956c**

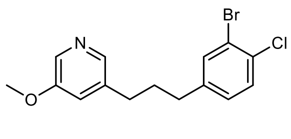

OX04956c

30 mg PtO2 was added in one portion to a $N_2$ degassed solution of the **OX04956b** (600 mg, 1.8 mmol) in MeOH (80 mL) at rt, the atmosphere was replaced with $H_2$, and the reaction stirred for 1 h. After completion, the reaction was filtered through Celite® using MeOH as an eluent, concentrated *in vacuo* and purified by flash column chromatography (30% EtOAc in Pentane) to afford **OX04956c** as a transparent oil (514 mg, 85%). $^1$H NMR (400 MHz, $CDCl_3$) δ 8.12 (d, *J* = 2.8 Hz, 1H), 8.02 (d, *J* = 1.8 Hz, 1H), 7.40 (d, *J* = 2.1 Hz, 1H), 7.31 (d, *J* = 8.2 Hz, 1H), 7.02 (dd, *J* = 8.2, 2.1 Hz, 1H), 6.96 (dd, *J* = 2.7, 1.8 Hz, 1H), 3.82 (s, 3H), 2.58 (q, *J* = 7.3 Hz, 4H), 1.95–1.85 (m, 2H). $^{13}$C NMR (101 MHz, $CDCl_3$) δ 155.7, 142.2, 142.1, 137.7, 134.0, 133.6, 131.8, 130.2, 128.6, 122.3, 120.7, 55.5, 34.3, 32.2. **HRMS (ESI+ve)** $C_{15}H_{15}BrClNO$ [M+H$^+$] calc. **340.0098**, found **340.0094**.

**OX04956d**

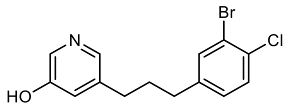

OX04956d

6 mL 48% HBr in $H_2O$ was added to **OX04956c** (440 mg, 1.3 mmol) and then the mixture was stirred at 120ºC for 48 h. After completion of the reaction, cooled to rt, the mixture was neutralised with sat. aq. $NaHCO_3$ and extracted with EtOAc (3 × 40 mL). The combined organic extracts were washed with brine (100 mL), dried over $Na_2SO_4$, filtered, and concentrated under reduced pressure. The resulting residue was purified by flash column chromatography (5% MeOH in $CH_2Cl_2$) afforded **OX04956d** as a transparent oil (120 mg, 28%). $^1$H NMR (400 MHz, MeOD) δ 7.92 (d, *J* = 2.7 Hz, 1H), 7.84 (d, *J* = 1.8 Hz, 1H), 7.48 (d, *J* = 2.0 Hz, 1H), 7.37 (d, *J* = 8.2 Hz, 1H), 7.13 (dd, *J* = 8.2, 2.1 Hz, 1H), 7.07 (dd, *J* = 2.6, 1.8 Hz, 1H), 2.59 (td, *J* = 7.7, 5.1 Hz, 4H), 1.96–1.82 (m, 2H). $^{13}$C NMR (101 MHz, MeOD) δ 155.7, 144.1, 140.9, 140.3, 135.9, 134.7, 132.6, 131.3, 130.1, 124.4, 122.9, 35.2, 33.4, 32.9. **HRMS (ESI+ve)** $C_{14}H_{13}BrClNO$ [M+H$^+$] calc. **325.9942**, found **325.9936**.

**OX04956**

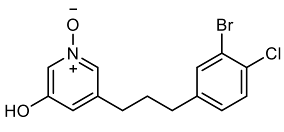

OX04956

To a solution of **OX04956d** (20 mg, 0.035 mmol) in $CH_2Cl_2$ (10 mL) at rt was added *m*-CPBA (1.5 eq.) and the resulting mixture stirred for 2 h. After completion, the reaction was quenched with $NaHCO_3$ (sat., aq.). The organic phase was washed with brine, dried ($Na_2SO_4$), filtered and concentrated *in vacuo*. Purification by flash column chromatography (0–100% EtOAc in Pentane then 0–6% MeOH in $CH_2Cl_2$) afforded **OX04956** as a yellow oil (11 mg, 52%). $^1$H NMR (400 MHz, MeOD) δ 7.77 (d, $J$ = 1.8 Hz, 2H), 7.53 (d, $J$ = 2.1 Hz, 1H), 7.40 (d, $J$ = 8.2 Hz, 1H), 7.17 (dd, $J$ = 8.2, 2.1 Hz, 1H), 6.96 (t, $J$ = 1.8 Hz, 1H), 2.62 (dt, $J$ = 13.4, 7.7 Hz, 4H), 1.97–1.86 (m, 2H). $^{13}$C NMR (101 MHz, MeOD) δ 157.8, 143.9, 143.4, 134.8, 132.8, 131.8, 131.3, 130.1, 127.1, 123.0, 119.4, 49.6, 49.4, 49.2, 49.0, 48.8, 48.6, 48.4, 35.1, 33.0, 32.8. **HRMS (ESI +ve)** $C_{14}H_{13}BrClNO_2$ [M+H$^+$] calc. **339.9745**, found **339.9745**. HPLC 98% (AUC), $t_R$ = 7.5 min.

**OX04955a**

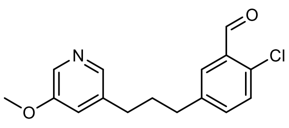

OX04955a

To a solution of **OX4956c** (555 mg, 1.6 mmol) in THF (4 mL) was added *n*-BuLi (1.4 mL, 2.5 M in hexane) dropwise over 10 min at -78ºC. The reaction was stirred for 5 min before adding DMF:THF mixture (1:9, 4 mL) in one portion at -78ºC. The reaction was stirred for 15 min then quenched with brine (5 mL) at -78ºC, and the mixture was allowed to warm to room temperature. The organic layer was washed with brine (8 mL × 2), dried ($Na_2SO_4$), concentrated *in vacuo* and purified with flash column chromatography (10–30% EtOAc in petroleum ether) to afford the **OX04955a** as a transparent oil (212 mg, 45%). $^1$H NMR (400 MHz, $CDCl_3$) δ 10.46 (s, 1H), 8.15 (d, $J$ = 2.8 Hz, 1H), 8.05 (s, 1H), 7.73 (dd, $J$ = 2.1, 0.7 Hz, 1H), 7.38–7.31 (m, 2H), 6.98 (dd, $J$ = 2.7, 1.8 Hz, 1H), 3.84 (s, 3H), 2.65 (dt, $J$ = 20.7, 7.7 Hz, 4H), 2.01–1.90 (m, 2H). $^{13}$C NMR (101 MHz, $CDCl_3$) δ 190.1, 155.8, 142.4, 141.4, 137.7, 135.7, 135.5, 135.2, 132.4, 130.7, 129.0, 120.7, 55.6, 34.6, 32.3, 32.3. LRMS (ESI +ve) 290 [M+H]$^+$. HRMS (ESI +ve) $C_{16}H_{16}ClNO_2$ [M+H]$^+$ calc. 290.0942, found 290.0941.

**OX04955b**

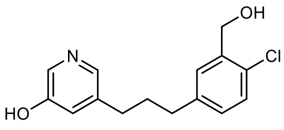

OX04955b

The demethylation using L-Selectride was adapted from a reported route.[1] L-Selectride (1.2 mL, 1 M in THF) was added to **OX04955a** (120 mg, 0.4 mmol) in THF (4 mL) and then the mixture was stirred at 70ºC for 48 h. After completion, the mixture was cooled to rt, quenched with MeOH, stirred for 30 min and concentrated *in vacuo*. The residue was then purified by flash column chromatography (0–8% MeOH in $CH_2Cl_2$) to afford **OX04955b** as a transparent oil (45 mg, 39%). $^1$H NMR (400 MHz, MeOD) δ 7.91 (d, $J$ = 2.6 Hz, 1H), 7.84 (d, $J$ = 1.8 Hz, 1H), 7.37 (d, $J$ = 2.2 Hz, 1H), 7.25 (d, $J$ = 8.1 Hz, 1H), 7.11–7.03 (m, 2H), 4.67 (s, 2H), 2.62 (dt, $J$ = 17.9, 7.8 Hz, 4H), 1.99–1.85 (m, 2H). $^{13}$C NMR (101 MHz, MeOD) δ 155.7, 142.2, 140.9, 140.6, 139.8, 135.8, 130.8, 130.0, 129.6, 129.5, 124.3, 62.3, 35.7, 33.7, 33.0. LRMS (ESI +ve) 278 $[M+H]^+$. HRMS (ESI +ve) $C_{15}H_{16}ClNO_2$ $[M+H]^+$ calc. 278.0942, found 278.0932.

**OX04955**

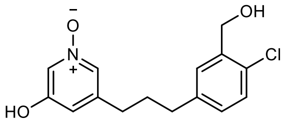

OX04955

To a solution of **OX04955b** (25 mg, 0.1 mmol) in $CH_2Cl_2$ (15 mL) at rt was added *m*-CPBA (1.5 eq.) and the resulting mixture stirred for 2 h. After completion, the reaction was quenched with $NaHCO_3$ (sat., aq.). The organic phase was washed with brine, dried ($Na_2SO_4$), filtered and concentrated *in vacuo*. Purification by flash column chromatography (0–10% MeOH in $CH_2Cl_2$) afforded **OX04955** as a yellow solid (17 mg, 64%). $^1$H NMR (400 MHz, MeOD) δ 7.76 (p, $J$ = 1.7 Hz, 2H), 7.38 (d, $J$ = 2.2 Hz, 1H), 7.26 (d, $J$ = 8.1 Hz, 1H), 7.10 (dd, $J$ = 8.1, 2.3 Hz, 1H), 6.95 (dd, $J$ = 2.1, 1.4 Hz, 1H), 4.67 (s, 2H), 2.68 (t, $J$ = 7.6 Hz, 2H), 2.65–2.59 (m, 2H), 2.01–1.91 (m, 2H). $^{13}$C NMR (151 MHz, MeOD) δ 157.8, 143.6, 141.9, 139.9, 131.9, 130.9, 130.1, 129.6, 129.5, 127.0, 119.4, 62.3, 35.5, 33.0, 33.0. LRMS (ESI +ve) 294 $[M+H]^+$. HRMS (ESI +ve) $C_{15}H_{16}ClNO_3$ $[M+H]^+$ calc. 294.0892, found 294.0890. HPLC 96% (AUC), $t_R$ = 3.2 min.

**OX04957a**

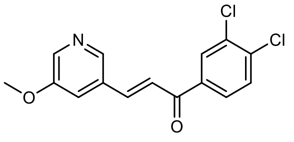

OX04957a

To a solution of 5-methoxynicotinaldehyde (1.00 g, 7.3 mmol) in 10 mL 1,4-dioxane was added 1-(3,4-dichlorophenyl)ethan-1-one (1.15 g, 6.1 mmol). $BF_3·OEt_2$ (6.0 mL) was added and the mixture was heated at 105ºC for 16h. After completion, the reaction was cooled to room temperature and slowly poured into aq. sat. $NaHCO_3$ and stirred until bubbling ceased. This solution was taken up into a separatory funnel and extracted with $CH_2Cl_2$. The organic layer was dried over $Na_2SO_4$ then under vacuum. To the crude residue, 5 mL MeOH was added. This mixture was diluted with pentane and allowed to stir for 20 minutes. The resulting suspension was filtered and washed with pentane to afford **OX04957a** as a yellow-brown solid (1.52 g, 67%). $^1$H NMR (400 MHz, $CDCl_3$) δ 8.47 (d, *J* = 1.8 Hz, 1H), 8.36 (d, *J* = 2.8 Hz, 1H), 8.10 (d, *J* = 2.0 Hz, 1H), 7.85 (dd, *J* = 8.3, 2.1 Hz, 1H), 7.80 (d, *J* = 15.7 Hz, 1H), 7.60 (d, *J* = 8.4 Hz, 1H), 7.48 (d, *J* = 15.7 Hz, 1H), 7.41 (dd, *J* = 2.7, 2.0 Hz, 1H), 3.93 (s, 3H). $^{13}$C NMR (101 MHz, $CDCl_3$) δ 187.6, 156.0, 142.7, 142.3, 139.9, 137.9, 137.4, 133.6, 131.0, 131.0, 130.7, 127.7, 123.1, 118.5, 55.9. HRMS (ESI +ve) $C_{15}H_{11}Cl_2NO_2$ $[M+H]^+$ calc. 308.0240, found 308.0255.

**OX04957b**

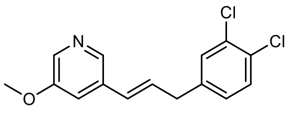


OX04957b

**OX04957a** (1.52 g, 4.9 mmol) was dissolved in TFA (10 mL) using sonication. Triethylsilane (20 mL) was added and the reaction was vigorously stirred for 48h. Upon completion, the reaction mixture was poured into sat. aq. $NaHCO_3$ and stirred until bubbling ceased. This solution was taken up into a separatory funnel and extracted with $CH_2Cl_2$, washed with brine, and dried under $Na_2SO_4$. Organic extracts were dried under vacuum. Purification by flash column chromatography (0–50% EtOAc in pentane) afforded **OX04957b** as a pale yellow solid (547 mg, 38%). $^1$H NMR (400 MHz, $CDCl_3$) δ 8.19–8.16 (m, 2H), 7.37 (d, *J* = 8.2 Hz, 1H), 7.31 (d, *J* = 2.1 Hz, 1H), 7.15 (dd, *J* = 2.8, 1.8 Hz, 1H), 7.08–7.03 (m, 1H), 6.41 (d, *J* = 15.9 Hz, 1H), 6.34 (dt, *J* = 15.8, 6.1 Hz, 1H), 3.85 (s, 3H), 3.51 (d, *J* = 6.0 Hz, 2H). $^{13}$C NMR (101 MHz, $CDCl_3$) δ 155.5, 140.4, 139.5, 136.3, 133.0, 132.3, 130.4, 130.2, 130.2, 130.2, 128.1, 127.9, 116.6, 55.2, 38.1. HRMS (ESI +ve) $C_{15}H_{13}Cl_2NO$ $[M+H]^+$ calc. 294.0447, found 294.0449.

**OX04957c**

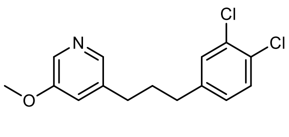


OX04957c

**OX04957b** (547 mg, 1.83 mmol) was dissolved in MeOH and the solution was degassed with $N_2$. $PtO_2$ (27 mg, 5 mol%) was added and the atmosphere was replaced with $H_2$. The reaction was stirred for 1h, filtered through Celite®, and dried under vacuum. Purification by flash column chromatography (0–40% EtOAc in pentane) afforded **OX04957c** as a pale yellow oil (411 mg, 76%). $^1$H NMR (400 MHz, $CDCl_3$) δ 8.16 (d, *J* = 2.8 Hz, 1H), 8.06 (d, *J* = 1.8 Hz, 1H), 7.34 (d, *J* = 8.2 Hz, 1H), 7.26 (s, 1H), 7.00 (dd, *J* = 8.1, 2.1 Hz, 2H), 3.85 (s, 3H), 2.62 (d, *J* = 4.9 Hz, 4H), 1.99–1.89 (m, 2H). $^{13}$C NMR (101 MHz, $CDCl_3$) δ 155.8, 142.3, 142.1, 137.8, 135.0, 132.4, 130.5, 130.5, 130.1, 128.0, 120.9, 55.7, 34.5, 32.3, 32.3. HRMS (ESI +ve) $C_{15}H_{15}Cl_2NO$ $[M+H^+]$ calc. 296.0604, found 296.0609.

**OX04957d**

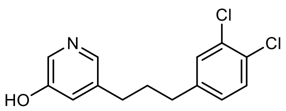

OX04957d

6 mL 48% HBr in $H_2O$ was added to **OX04957c** (411 mg, 1.39 mmol), and the reaction was heated to 120℃ and stirred for 48 h. Upon completion, the reaction mixture was poured into aq. sat. $NaHCO_3$ and stirred until bubbling ceased. This solution was taken up into a separatory funnel and extracted with EtOAc, washed with brine, and dried under $Na_2SO_4$. Organic extracts were dried under vacuum. Purification by flash column chromatography (0–100% EtOAc in pentane) afforded **OX04957d** as a pale orange solid (180 mg, 46%). $^1$H NMR (400 MHz, $CDCl_3$) δ 8.12 (d, *J* = 2.6 Hz, 1H), 7.91 (d, *J* = 1.8 Hz, 1H), 7.33 (d, *J* = 8.2 Hz, 1H), 7.25 (d, *J* = 2.1 Hz, 1H), 7.14 (t, *J* = 2.1 Hz, 1H), 6.99 (dd, *J* = 8.2, 2.1 Hz, 1H), 2.61 (td, *J* = 8.3, 3.4 Hz, 4H), 1.98–1.86 (m, 2H). $^{13}$C NMR (101 MHz, $CDCl_3$) δ 155.4, 141.9, 139.6, 139.2, 133.9, 132.4, 130.5, 130.4, 130.1, 128.0, 125.3, 34.5, 32.2, 32.1. HRMS (ESI +ve) $C_{14}H_{13}Cl_2NO$ [M+H$^+$] calc. 282.0447, found 282.0459.

**OX04957**

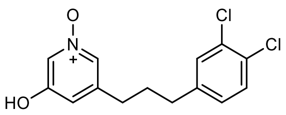

OX04957

To a solution of **OX04957d** (50 mg, 0.18 mmol) in $CH_2Cl_2$ (10 mL) was added 77% *meta*-chloroperoxybenzoic acid (60 mg, 0.27 mmol) and the reaction was stirred at room temperature for 2 h. Another portion of 77% *meta*-chloroperoxybenzoic acid (40 mg, 0.18 mmol) was added and the reaction was further stirred for 30 mins. Upon completion, aq. 2M NaOH (5 mL) was added to the reaction mixture. The pH was adjusted to 3 – 4 using aq. 1M HCl, and a precipitate formed in the organic phase. The precipitate was filtered and washed with $CH_2Cl_2$ to afford **OX04957** as a white solid (12 mg, 23%). $^1$H NMR (400 MHz, DMSO) δ 10.38 (s, 1H), 7.68 (t, *J* = 1.5 Hz, 1H), 7.61 (t, *J* = 1.9 Hz, 1H), 7.53 (d, *J* = 8.3 Hz, 1H), 7.51 (d, *J* = 2.1 Hz, 1H), 7.22 (dd, *J* = 8.2, 2.1 Hz, 1H), 6.67 (t, *J* = 1.7 Hz, 1H), 2.63–2.56 (m, 2H), 2.47 (t, *J* = 6.9 Hz, 2H), 1.84 (tt, *J* = 9.5, 6.6 Hz, 2H). $^{13}$C NMR (101 MHz, $CDCl_3$) δ 157.6, 141.4, 141.3, 132.5, 130.6, 130.4, 130.3, 129.8, 128.0, 126.7, 119.8, 34.3, 32.2, 31.5. HRMS (ESI +ve) $C_{14}H_{13}Cl_2NO_2$ [M+H$^+$] calc. 298.0396, found 298.0408. HPLC 98% (AUC), $t_R$ = 7.3 min.

**OX04958a**

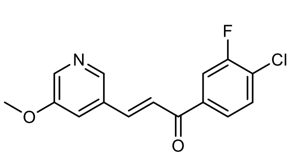

OX04958a

To a solution of 5-methoxynicotinaldehyde (1.00 g, 7.3 mmol) in 10 mL 1,4-dioxane was added 1-(4-chloro-3-fluorophenyl)ethan-1-one (1.05 g, 6.1 mmol). $BF_3{\cdot}OEt_2$ (6.0 mL) was added and the mixture was heated to 105°C for 16h. After completion, the reaction was cooled to room temperature and slowly poured into aq. sat. $NaHCO_3$ and stirred until bubbling ceased. This solution was taken up into a separatory funnel and extracted with $CH_2Cl_2$. Organic extracts were dried under vacuum. To the crude residue, a small amount of MeOH was added. This mixture was diluted with pentane, and allowed to stir for 20 minutes. The resulting suspension was filtered and washed with pentane to afford **OX04958a** as a yellow-brown solid (1.13 g, 64%). $^1H$ NMR (400 MHz, $CDCl_3$) δ 8.47 (d, *J* = 1.8 Hz, 1H), 8.36 (d, *J* = 2.8 Hz, 1H), 7.82–7.73 (m, 3H), 7.55 (dd, *J* = 8.2, 7.1 Hz, 1H), 7.48 (d, *J* = 15.7 Hz, 1H), 7.42–7.38 (m, 1H), 3.93 (s, 3H). $^{13}C$ NMR (101 MHz, $CDCl_3$) δ 187.5, 158.4 (d, *J* = 251.3 Hz), 156.0, 142.6, 142.3, 139.9, 138.1 (d, *J* = 5.4 Hz), 131.2, 131.0, 126.7 (d, *J* = 17.9 Hz), 125.0 (d, *J* = 3.7 Hz), 118.5, 116.7 (d, *J* = 22.1 Hz), 116.6, 55.9. $^{19}F$ NMR (376 MHz, $CDCl_3$) δ -113.42 (dd, *J* = 9.4, 7.1 Hz). HRMS (ESI +ve) $C_{15}H_{11}ClFNO_2$ $[M+H]^+$ calc. 292.0535, found 292.0532.

**OX04958b**

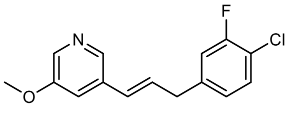


OX04958b

**OX04958a** (1.13 g, 3.9 mmol) was dissolved in TFA (8 mL) using sonication. Triethylsilane (16 mL) was added and the reaction was vigorously stirred for 48h. Upon completion, the reaction mixture was poured into aq. sat. $NaHCO_3$ and stirred until bubbling ceased. This solution was taken up into a separatory funnel and extracted with $CH_2Cl_2$, washed with brine, and dried under $Na_2SO_4$. Organic extracts were dried under vacuum. Purification by flash column chromatography (0–40% EtOAc in pentane) afforded **OX04958b** as a pale yellow solid (406 mg, 38%). $^1H$ NMR (400 MHz, $CDCl_3$) δ 8.19 (d, *J* = 1.8 Hz, 1H), 8.17 (d, *J* = 2.8 Hz, 1H), 7.33 (t, *J* = 7.9 Hz, 1H), 7.16 (dd, *J* = 2.8, 1.8 Hz, 1H), 7.02 (dd, *J* = 9.9, 2.0 Hz, 1H), 6.96 (ddd, *J* = 8.2, 2.0, 0.8 Hz, 1H), 6.42 (d, *J* = 16.0 Hz, 1H), 6.35 (dt, *J* = 15.8, 6.0 Hz, 1H), 3.86 (s, 3H), 3.54 (d, *J* = 5.9 Hz, 2H). $^{13}C$ NMR (101 MHz, $CDCl_3$) δ 158.1 (d, *J* = 248.8 Hz), 155.8, 140.6, 140.4 (d, *J* = 6.3 Hz), 136.5, 133.3, 130.6, 130.5, 128.3, 125.1 (d, *J* = 3.5 Hz), 118.9 (d, *J* = 8.1 Hz), 116.9, 116.9 (d, *J* = 21.2 Hz), 55.6, 38.6 (d, *J* = 1.6 Hz). $^{19}F$ NMR (376 MHz, $CDCl_3$) δ -115.44 (dd, *J* = 10.0, 7.6 Hz). HRMS (ESI +ve) $C_{15}H_{13}ClFNO$ $[M+H]^+$ calc. 278.0743, found 278.0736

**OX04958c**

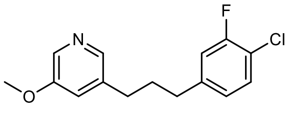


OX04958c

**OX04958b** (406 mg, 1.46 mmol) was dissolved in MeOH and the solution was degassed with $N_2$. $PtO_2$ (20 mg, 5 mol%) was added and the atmosphere was replaced with $H_2$. The reaction was stirred for 1h, filtered through Celite®, and dried under vacuum to afford **OX04958c** as a pale yellow oil (380 mg, 93%). $^1$H NMR (400 MHz, $CDCl_3$) δ 8.16 (d, $J$ = 2.8 Hz, 1H), 8.06 (d, $J$ = 1.7 Hz, 1H), 7.29 (t, $J$ = 7.9 Hz, 1H), 7.00–6.98 (m, 1H), 6.96 (dd, $J$ = 10.1, 2.0 Hz, 1H), 6.89 (ddd, $J$ = 8.2, 2.1, 0.8 Hz, 1H), 3.85 (s, 3H), 2.63 (dd, $J$ = 7.4, 1.7 Hz, 4H), 1.99–1.88 (m, 2H). $^{13}$C NMR (151 MHz, $CDCl_3$) δ 158.2 (d, $J$ = 248.5 Hz), 155.8, 142.8 (d, $J$ = 6.5 Hz), 142.4, 137.8, 135.1, 130.5, 125.0 (d, $J$ = 3.4 Hz), 120.9, 118.4 (d, $J$ = 17.7 Hz), 116.6 (d, $J$ = 20.4 Hz), 55.7, 34.7, 32.3, 32.3. $^{19}$F NMR (376 MHz, $CDCl_3$) δ -115.94 (dd, $J$ = 10.1, 7.6 Hz). HRMS (ESI +ve) $C_{15}H_{15}ClFNO$ [M+H$^+$] calc. 280.0899, found 280.0913

**OX04958d**

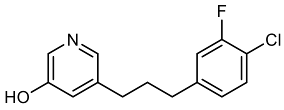


OX04958d

5 mL 48% HBr in $H_2O$ was added to **OX04958c** (380 mg, 1.36 mmol), and the reaction was heated to 120℃ and stirred for 48h. Upon completion, the reaction mixture was poured into aq. sat. $NaHCO_3$ and stirred until bubbling ceased. This solution was taken up into a separatory funnel and extracted with EtOAc, washed with brine, and dried under $Na_2SO_4$. Organic extracts were dried under vacuum. Purification by flash column chromatography (0–100% EtOAc in pentane) afforded **OX04958d** as a pale orange oil (63.8 mg, 18%). $^1$H NMR (400 MHz, $CDCl_3$) δ 8.15–8.11 (m, 1H), 7.92 (s, 1H), 7.28 (t, $J$ = 8.0 Hz, 1H), 7.15 (t, $J$ = 2.1 Hz, 1H), 6.95 (dd, $J$ = 10.1, 2.0 Hz, 1H), 6.88 (ddd, $J$ = 8.1, 2.0, 0.8 Hz, 1H), 2.64 (d, $J$ = 2.8 Hz, 2H), 2.60 (d, $J$ = 2.9 Hz, 2H), 1.99–1.87 (m, 2H). $^{13}$C NMR (101 MHz, $CDCl_3$) δ 158.1 (d, $J$ = 248.6 Hz), 155.4, 142.6 (d, $J$ = 6.5 Hz), 139.6, 139.1, 133.9, 130.6, 125.4, 125.0 (d, $J$ = 3.4 Hz), 118.5 (d, $J$ = 17.5 Hz), 116.6 (d, $J$ = 20.5 Hz), 34.6, 32.2, 32.1. $^{19}$F NMR (376 MHz, $CDCl_3$) δ -115.85 (dd, $J$ = 10.1, 7.6 Hz). HRMS (ESI +ve) $C_{14}H_{13}ClFNO$ [M+H$^+$] calc. 266.0743, found 266.0751

**OX04958**

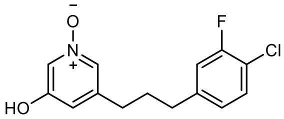


OX04958

To a solution of **OX04958c** (60 mg, 0.23 mmol) in $CH_2Cl_2$ (10 mL). 77% *meta*-chloroperoxybenzoic acid (83 mg, 0.36 mmol) was added and the reaction was stirred at room temperature for 2 h. Upon completion, aq. 2M NaOH (5 mL) was added to the reaction mixture. The pH was adjusted to 3-4 using aq. 1M HCl, and the aqueous phase was extracted with 2:1 $CHCl_3$:EtOH. The organic extract was dried and purified by flash column chromatography (0–5% MeOH in $CH_2Cl_2$) to afford **OX04958** as a pale yellow solid (40.3 mg, 65%). $^1$H NMR (400 MHz, $CDCl_3$) δ 8.00 (d, $J$ = 2.1 Hz, 1H), 7.64 (s, 1H), 7.28 (t, $J$ = 7.2 Hz, 1H), 6.94–

6.90 (m, 2H), 6.86 (dd, $J$ = 8.2, 2.0 Hz, 1H), 2.60 (t, $J$ = 7.6 Hz, 2H), 2.53 (t, $J$ = 7.7 Hz, 2H), 1.93–1.84 (m, 2H). $^{13}$C NMR (101 MHz, $CDCl_3$) δ 158.1 (d, $J$ = 249.5 Hz), 157.5, 142.1 (d, $J$ = 6.4 Hz), 141.3, 130.6, 129.8, 126.7, 124.9 (d, $J$ = 3.5 Hz), 119.8, 118.6 (d, $J$ = 17.7 Hz), 116.6 (d, $J$ = 20.6 Hz), 34.4, 32.1, 31.4. $^{19}$F NMR (377 MHz, $CDCl_3$) δ -115.60 (dd, $J$ = 10.1, 7.8 Hz). HRMS (ESI +ve) $C_{14}H_{13}ClFNO_2$ [M+H$^+$] calc. 282.0692, found 282.0701. HPLC 96% (AUC), $t_R$ = 6.5 min.

**OX04959a**

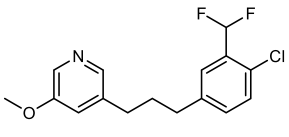

OX04959a

To a solution of **OX04955a** (100 mg, 0.35 mmol) in $CH_2Cl_2$ (8 mL) was added DAST (0.11 mL, 0.84 mmol) dropwise. The reaction was stirred for 16 h at rt before quenched with $NaHCO_3$ (10 mL, sat., aq.) and the two phases were separated. The organic layer was washed with $NaHCO_3$ (10 mL, sat., aq.), brine (15 mL), dried ($Na_2SO_4$), filtered and concentrated *in vacuo*. The residue was purified by flash column chromatography (10–30% EtOAc in petroleum ether) to afford **OX04959a** as a yellow oil (40 mg, 37%). $^1$H NMR (400 MHz, $CDCl_3$) δ 8.16 (d, $J$ = 2.8 Hz, 1H), 8.06 (d, $J$ = 1.8 Hz, 1H), 7.47 (d, $J$ = 2.2 Hz, 1H), 7.33 (dt, $J$ = 8.3, 1.2 Hz, 1H), 7.21 (ddt, $J$ = 8.2, 1.9, 0.9 Hz, 1H), 7.09–6.77 (m, 2H), 3.85 (s, 3H), 2.65 (dt, $J$ = 17.4, 7.7 Hz, 4H), 2.03–1.91 (m, 2H). $^{13}$C NMR (101 MHz, $CDCl_3$) δ 155.8, 142.4, 141.3, 137.7, 135.2, 132.1 (t, $J$ = 2.0 Hz), 131.7, 130.3, 130.0, 126.8 (t, $J$ = 5.8 Hz), 120.8, 112.2 (t, $J$ = 238.0 Hz), 55.6, 34.7, 32.4, 32.3. $^{19}$F NMR (376 MHz, $CDCl_3$) δ -115.0. LRMS (ESI +ve) 312 $[M+H]^+$. HRMS (ESI +ve) $C_{16}H_{16}ClF_2NO$ $[M+H]^+$ calc. 312.0961, found 312.0967.

**OX04959b**

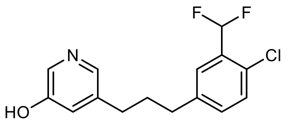

OX04959b

The demethylation using L-Selectride was adapted from a reported route.[1] L-Selectride (0.4 mL, 1 M in THF) was added to **OX04959a** (40 mg, 0.13 mmol) in THF (3 mL) and then the mixture was stirred at 70ºC for 48 h. After completion, the mixture was cooled to rt, quenched with MeOH, stirred for 30 min and concentrated *in vacuo*. The residue was then purified by flash column chromatography (0–8% MeOH in $CH_2Cl_2$) to afford **OX04959b** as a white solid (40 mg, 92%). $^1$H NMR (400 MHz, $CDCl_3$) δ 8.12 (d, $J$ = 2.7 Hz, 1H), 7.91 (d, $J$ = 1.8 Hz, 1H), 7.45 (d, $J$ = 2.1 Hz, 1H), 7.32 (dt, $J$ = 8.2, 1.2 Hz, 1H), 7.22–7.18 (m, 1H), 7.14 (t, $J$ = 2.2 Hz, 1H), 6.92 (t, $J$ = 55.0 Hz, 1H), 2.64 (dt, $J$ = 19.9, 7.7 Hz, 4H), 2.00–1.90 (m, 2H). $^{13}$C NMR (101 MHz, $CDCl_3$) δ 155.4, 141.2, 139.5, 139.3, 134.1, 132.1 (t, $J$ = 1.8 Hz), 130.0, 129.1, 127.2, 126.8 (t, $J$ = 5.9 Hz), 125.2, 112.2 (t, $J$ = 237.9 Hz), 34.8, 32.3, 32.2. $^{19}$F NMR (376 MHz, $CDCl_3$) δ -115.0. HRMS (ESI +ve) $C_{15}H_{14}ClF_2NO$ $[M+H]^+$ calc. 298.0805, found 298.0814.

**OX04959**

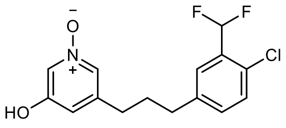


OX04959

To a solution of **OX04959b** (20 mg, 0.07 mmol) in $CH_2Cl_2$ (10 mL) at rt was added *m*-CPBA (1.5 eq.) and the resulting mixture stirred for 2 h. After completion, the reaction was quenched with $NaHCO_3$ (sat, aq.). The organic phase was washed with brine, dried ($Na_2SO_4$), filtered and concentrated *in vacuo*. Purification by flash column chromatography (0–6% MeOH in $CH_2Cl_2$) afforded **OX04959** as a white solid (9 mg, 43%). $^1$H NMR (400 MHz, $CDCl_3$) δ 8.01 (s, 1H), 7.63 (s, 1H), 7.44 (d, *J* = 2.2 Hz, 1H), 7.33 (d, *J* = 8.2 Hz, 1H), 7.19 (dd, *J* = 8.2, 2.1 Hz, 1H), 7.08–6.77 (m, 2H), 2.65 (t, *J* = 7.7 Hz, 2H), 2.55 (t, *J* = 7.7 Hz, 2H), 1.97–1.85 (m, 2H). $^{13}$C NMR (151 MHz, CDCl3) δ 157.5, 141.3, 140.7, 132.0, 131.9, 130.5, 130.1, 130.1, 126.8 (t, *J* = 5.9 Hz), 126.7, 119.8, 112.2 (t, *J* = 237.1 Hz), 34.6, 32.3, 31.6. $^{19}$F NMR (376 MHz, $CDCl_3$) δ -115.0. LRMS (ESI +ve) 314 $[M+H]^+$. HRMS (ESI +ve) $C_{15}H_{14}ClF_2NO_2$ $[M+H]^+$ calc. 314.0754, found 314.0768. HPLC 98% (AUC), $t_R$ = 6.8 min.

**OX04960a**

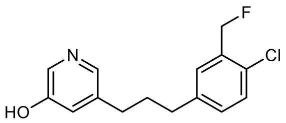


OX04960a

**OX04955b** (110 mg, 0.4 mmol) was dissolved in anhydrous $CH_2Cl_2$ ($CH_2Cl_2$) (4 mL) at room temperature under inert conditions. The solution was then cooled to 0°C and stirred for 5 minutes. Diethylaminosulfur trifluoride (DAST) (0.2 mL, 1.2 mmol) was added dropwise, and the mixture was allowed to warm gradually to room temperature. The reaction was stirred for 24 h. Upon completion, $NaHCO_3$ solution (4 mL) was added, followed by EtOAc. The organic layer was separated, dried over $Na_2SO_4$, and concentrated. Purification by flash chromatography (0–10% MeOH in $CH_2Cl_2$) afforded **OX04960a** as a brown solid (12 mg, 11% yield). $^1$H NMR (400 MHz, $CDCl_3$) δ 8.14 (d, *J* = 2.7 Hz, 1H), 7.93 (d, *J* = 1.8 Hz, 1H), 7.28 (dd, *J* = 8.2, 1.3 Hz, 2H), 7.14–7.11 (m, 1H), 7.09 (dt, *J* = 7.9, 1.6 Hz, 1H), 5.54 (s, 1H), 5.42 (s, 1H), 2.63 (dt, *J* = 13.3, 7.7 Hz, 4H), 2.01–1.89 (m, 2H). $^{13}$C NMR (101 MHz, $CDCl_3$) δ 154.8, 140.7, 139.7, 139.4, 134.1, 129.7, 129.7, 129.4, 128.6, 128.5, 124.8, 81.7 (d, *J* = 168.5 Hz), 34.6, 32.2, 32.1. $^{19}$F NMR (377 MHz, $CDCl_3$) δ -216.85. HRMS (ESI +ve) $C_{15}H_{15}ClFNO$ $[M+H^+]$ calc. 280.0899, found 280.0964

**OX04960**

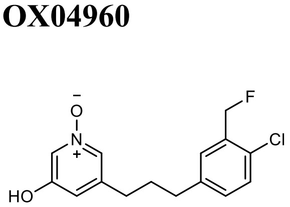


OX04960

To a solution of **OX04960a** (11.7 mg, 0.04 mmol, 1.0 eq.) in $CH_2Cl_2$ (7 mL), *meta*-chloroperoxybenzoic acid (1.5 eq.) was added at room temperature. The reaction progress was monitored by TLC. After 2 hours, $NaHCO_3$ solution (10 mL) was added. The mixture was diluted with $CH_2Cl_2$, and the organic layer was separated, washed with brine, and dried over $Na_2SO_4$. Purification by flash chromatography (0–10% MeOH in $CH_2Cl_2$) yielded **OX04960** (5 mg, 44% yield) as a light brown solid. $^1$H NMR (600 MHz, $CDCl_3$) δ 7.99 (s, 1H), 7.63 (s, 1H), 7.29 (d, *J* = 8.1 Hz, 1H), 7.27 (s, 1H), 7.07 (d, *J* = 7.5 Hz, 1H), 6.92 (s, 1H), 5.48 (d, *J* = 47.2 Hz, 2H), 2.58 (dt, *J* = 54.0, 7.7 Hz, 4H), 1.91 (p, *J* = 7.7 Hz, 2H). $^{13}$C NMR (151 MHz, $CDCl_3$) δ 157.3, 141.3, 140.1, 134.1 (d, *J* = 17.7 Hz), 129.9 (d, *J* = 5.3 Hz), 129.7, 129.6 (d, *J* = 3.0 Hz), 129.4, 128.3 (d, *J* = 8.7 Hz), 126.4, 119.8, 81.6 (d, *J* = 168.3 Hz), 34.3, 32.0, 31.4. $^{19}$F NMR (377 MHz, CDCl3) δ -217.37. **HRMS (ESI +ve)** $C_{15}H_{16}ClFNO_2$ [M+H$^+$] calc. **296.0848**, found **296.0829**. HPLC 95% (AUC), $t_R$ = 7.3 min.

**OX04961a**

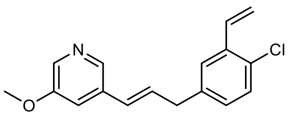

OX04961a

To a nitrogen-degassed microwave vial with **OX04956b** (1150 mg, 3.4 mmol), vinyl boronic ester (787 mg, 5.1 mmol) and $Cs_2CO_3$ (1111 mg, 10.2 mmol) in THF:$H_2O$ mixture (9:1, 10 mL) was added $PPh_3$ (89 mg, 0.3 mmol) and Pd(OAc)$_2$ (38 mg, 0.2 mmol). After degassing 5 min with nitrogen, the vial was sealed and the mixture was placed on a pre-heated heating plate and stirred at 70ºC for 25 h and then the mixture was diluted with EtOAc and filtered through Celite®. The organic layer was washed with brine (30 mL × 2), dried over $Na_2SO_4$ and concentrated *in vacuo*. **OX04961a** was purified via column chromatography (0–30% EtOAc in petroleum) as a yellow oil (543 mg, 56%). $^1$H NMR (400 MHz, $CDCl_3$) δ 8.19 (d, *J* = 1.8 Hz, 1H), 8.16 (d, *J* = 2.8 Hz, 1H), 7.42 (d, *J* = 2.2 Hz, 1H), 7.30 (d, *J* = 8.2 Hz, 1H), 7.15 (dd, *J* = 2.8, 1.8 Hz, 1H), 7.13–7.05 (m, 2H), 6.42–6.38 (m, 2H), 5.74 (dd, *J* = 17.6, 1.1 Hz, 1H), 5.38 (dd, *J* = 11.0, 1.1 Hz, 1H), 3.85 (s, 3H), 3.56–3.52 (m, 2H). $^{13}$C NMR (101 MHz, $CDCl_3$) δ 155.8, 140.8, 138.3, 136.5, 135.9, 133.6, 133.3, 131.4, 131.3, 129.9, 129.4, 127.9, 126.9, 116.9, 116.8, 55.7, 38.9. LRMS (ESI +ve) 286 [M+H]$^+$. HRMS (ESI +ve) $C_{17}H_{16}ClNO$ [M+H]$^+$ calc. 286.0993, found 286.0971.

**OX04961b**

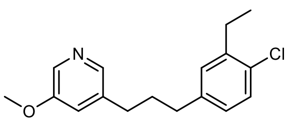

OX04961b

54 mg Pd/C (10% M/w) was added in one portion to a $N_2$ degassed solution of the **OX04961a** (540 mg, 1.9 mmol) in MeOH (60 mL) at rt, the atmosphere was replaced with $H_2$, and the reaction stirred for 1 h. After

completion, the reaction was filtered through Celite® using MeOH as an eluent, concentrated *in vacuo* and purified by flash column chromatography (30% EtOAc in Pentane) to afford the **OX04961b** as a yellow oil (215 mg, 39%). $^{1}$H NMR (400 MHz, $CDCl_3$) δ 8.17 (d, *J* = 2.8 Hz, 1H), 8.09 (d, *J* = 1.8 Hz, 1H), 7.30–7.23 (m, 1H), 7.06–7.01 (m, 2H), 6.96 (dd, *J* = 8.1, 2.3 Hz, 1H), 3.87 (s, 3H), 2.74 (q, *J* = 7.5 Hz, 2H), 2.64 (q, *J* = 7.7 Hz, 4H), 2.01–1.90 (m, 2H), 1.24 (t, *J* = 7.5 Hz, 3H). $^{13}$C NMR (101 MHz, $CDCl_3$) δ 155.8, 142.2, 141.6, 140.5, 138.3, 134.6, 131.4, 129.8, 129.4, 127.2, 121.2, 55.7, 34.8, 32.6, 32.4, 26.9, 14.3. LRMS (ESI +ve) 290 $[M+H]^+$. HRMS (ESI +ve) $C_{17}H_{20}ClNO$ $[M+H]^+$ calc. 290.1306, found 290.1323.

**OX04961c**

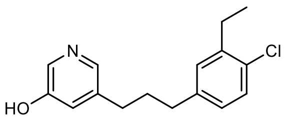


OX04961c

5 mL 48% HBr in $H_2O$ was added to **OX04961b** (215 mg, 0.7 mmol) and then the mixture was stirred at 120ºC for 48 h. After completion of the reaction, cooled to rt, the mixture was neutralized with sat. aq. $NaHCO_3$ and extracted with EtOAc (3 × 40 mL). The combined organic extracts were washed with brine (100 mL), dried over $Na_2SO_4$, filtered, and concentrated under reduced pressure. The resulting residue was purified by flash column chromatography (0 – 6% MeOH in $CH_2Cl_2$) afforded **OX04961c** as a yellow oil (113 mg, 55%). $^{1}$H NMR (400 MHz, $CDCl_3$) δ 11.27 (s, 1H), 8.10 (d, *J* = 2.6 Hz, 1H), 7.90 (d, *J* = 1.8 Hz, 1H), 7.22 (d, *J* = 8.1 Hz, 1H), 7.14 (t, *J* = 2.2 Hz, 1H), 7.01 (d, *J* = 2.2 Hz, 1H), 6.91 (dd, *J* = 8.1, 2.2 Hz, 1H), 2.71 (q, *J* = 7.5 Hz, 2H), 2.59 (td, *J* = 7.8, 2.4 Hz, 4H), 1.97–1.86 (m, 2H), 1.21 (t, *J* = 7.5 Hz, 3H). $^{13}$C NMR (101 MHz, $CDCl_3$) δ 155.5, 141.6, 140.4, 139.9, 139.1, 133.8, 131.4, 129.7, 129.4, 127.2, 125.3, 34.8, 32.4, 32.3, 26.9, 14.3. **LRMS (ESI +ve)** 276 $[M+H]^+$. **HRMS (ESI +ve)** $C_{16}H_{18}ClNO$ $[M+H^+]$ calc. 276.1150, found 276.1124.

**OX04961**

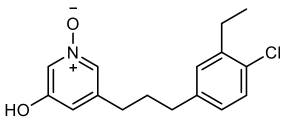


OX04961

**3-(3-(4-chloro-3-ethylphenyl)propyl)-5-hydroxypyridine 1-oxide**

To a solution of **OX04961c** (64 mg, 0.2 mmol) in $CH_2Cl_2$ (20 mL) at rt was added *m*-CPBA (1.5 eq.) and the resulting mixture stirred for 2 h. After completion, the reaction was quenched with $NaHCO_3$ (sat., aq.). The organic phase was washed with brine, dried ($Na_2SO_4$), filtered and concentrated *in vacuo*. Purification by flash column chromatography (0–6% MeOH in $CH_2Cl_2$) afforded **OX04961** as a yellow oil (28 mg, 41%). $^{1}$H NMR (600 MHz, $CDCl_3$) δ 7.99 (s, 1H), 7.63 (s, 1H), 7.23 (d, *J* = 8.1 Hz, 1H), 7.00 (d, *J* = 2.2 Hz, 1H), 6.93–6.88 (m, 2H), 2.72 (q, *J* = 7.5 Hz, 2H), 2.58 (t, *J* = 7.6 Hz, 2H), 2.53 (t, *J* = 7.7 Hz, 2H), 1.89 (h, *J*

= 6.9 Hz, 2H), 1.22 (t, *J* = 7.5 Hz, 3H). $^{13}C$ NMR (151 MHz, $CDCl_3$) δ 157.5, 141.8, 141.6, 139.9, 131.6, 130.0, 129.7, 129.5, 127.2, 126.6, 119.9, 34.6, 32.3, 31.8, 26.9, 14.3. LRMS (ESI +ve) 292 $[M+H]^+$. HRMS (ESI +ve) $C_{16}H_{18}ClNO_2$ $[M+H]^+$ calc. 292.1099, found 292.1100. HPLC 96% (AUC), $t_R$ = 3.0 min.

**OX04962a**

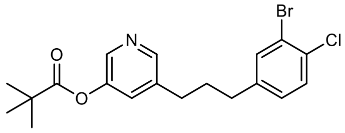


OX04962a

To a slurry of **OX04956d** (2.2 g, 6.8 mmol) and $NEt_3$ (1.43 mL, 10.2 mmol) in $CH_2Cl_2$ (80 mL) was added trimethylacetyl chloride (1.09 mL, 8.9 mmol) dropwise at 0ºC over 10 min. The mixture was then stirred at rt for 3 h before poured into $H_2O$ (100 mL). The organic layer was separated, dried ($Na_2SO_4$) and concentrated *in vacuo* to afford **OX04962a** as an orange oil (2.7 g, 98%). $^1H$ NMR (400 MHz, $CDCl_3$) δ 8.32–8.28 (m, 1H), 8.23 (d, *J* = 2.4 Hz, 1H), 7.43 (d, *J* = 2.1 Hz, 1H), 7.35 (d, *J* = 8.2 Hz, 1H), 7.26–7.23 (m, 1H), 7.04 (dd, *J* = 8.2, 2.1 Hz, 1H), 2.63 (dt, *J* = 17.2, 7.7 Hz, 4H), 2.01–1.89 (m, 2H), 1.37 (s, 9H). $^{13}C$ NMR (101 MHz, $CDCl_3$) δ 176.8, 147.8, 147.0, 142.0, 141.1, 137.9, 133.7, 132.0, 130.3, 129.1, 128.7, 122.4, 39.3, 34.3, 32.2, 32.0, 27.2. LRMS (ESI +ve) 410 $[M+H]^+$. HRMS (ESI +ve) $C_{19}H_{21}BrClNO_2$ $[M+H]^+$ calc. 410.0517, found 410.0513.

**OX04962b**

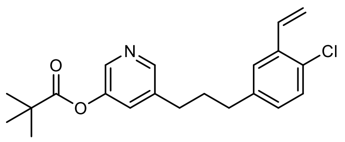


OX04962b

To a degassed microwave vial with **OX04962a** (701 mg, 1.7 mmol) and tributyl(vinyl)stannane (599 mg, 1.9 mmol) in DMF (4 mL) was added $Pd(PPh_3)4$ (109 mg, 0.1 mmol). The mixture was degassed for 5 min before the vial was sealed and placed on a pre-heated heating plate. The mixture was stirred for 24 h at 90ºC. After completion, the mixture was diluted with EtOAc (5 mL), poured into KF (5 mL, sat., aq.) then the mixture was stirred for 30 min before filtered through Celite®. The organic layer was washed with $NH_4Cl$ (20 mL × 2, sat., aq.), dried ($Na_2SO_4$) and concentrated *in vacuo*. The crude product was purified with flash column chromatography (0–17% EtOAc in petroleum ether) to afford **OX04962b** as a colourless oil (207 mg, 34%). $^1H$ NMR (400 MHz, $CDCl_3$) δ 8.31 (d, *J* = 1.9 Hz, 1H), 8.24 (d, *J* = 2.5 Hz, 1H), 7.36 (d, *J* = 2.2 Hz, 1H), 7.29–7.24 (m, 2H), 7.09 (dd, *J* = 17.5, 11.0 Hz, 1H), 7.02 (dd, *J* = 8.2, 2.2 Hz, 1H), 5.74 (dd, *J* = 17.5, 1.1 Hz, 1H), 5.37 (dd, *J* = 11.0, 1.1 Hz, 1H), 2.66 (dt, *J* = 10.9, 7.6 Hz, 4H), 2.03–1.92 (m, 2H), 1.37 (s, 9H). $^{13}C$ NMR (101 MHz, $CDCl_3$) δ 176.8, 147.7, 147.0, 141.0, 140.4, 138.1, 135.6, 133.4, 130.9, 129.7,

129.1, 129.1, 126.6, 116.5, 39.3, 34.8, 32.4, 32.1, 27.2. LRMS (ESI +ve) 358 $[M+H]^+$. HRMS (ESI +ve) $C_{21}H_{24}ClNO_2$ $[M+H]^+$ calc. 358.1568, found 358.1576.

**OX04962**

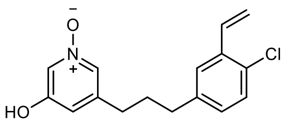


OX04962

To a mixture of **OX04962b** (30 mg, 0.1 mmol) in $CH_2Cl_2$ (10 mL) was added *m*-CPBA (1.5 eq.) in one portion at rt. The mixture was stirred at rt for 2 h then concentrated *in vacuo*, followed by purification with flash column chromatography (20–100% EtOAc in petroleum ether then 0–20% MeOH in EtOAc). The MeOH in EtOAc fractions were checked by thin layer chromatography, collected, concentrated *in vacuo* and dissolved in MeOH (10 mL). 4M aq. NaOH (2 mL) was added subsequently and the mixture was stirred for 16 h at rt. After completion, the mixture was concentrated *in vacuo*, taken up with sat. aq. $NH_4Cl$ to adjust pH < 8 and extracted with $CH_2Cl_2$ (3 × 20 mL). The organic layer was washed with brine, dried with anhydrous $Na_2SO_4$, concentrated *in vacuo* and purified with flash column chromatography (0–10% MeOH in $CH_2Cl_2$) to afford **OX04962** as a brown solid (8.3 mg, 34%). $^1$H NMR (400 MHz, $CDCl_3$) δ 8.00 (s, 1H), 7.63 (s, 1H), 7.33 (d, *J* = 2.1 Hz, 1H), 7.27–7.25 (m, 2H), 7.07 (dd, *J* = 17.5, 11.0 Hz, 1H), 6.97 (dd, *J* = 8.2, 2.2 Hz, 1H), 6.91 (s, 1H), 5.73 (dd, *J* = 17.6, 1.1 Hz, 1H), 5.37 (dd, *J* = 11.0, 1.1 Hz, 1H), 2.60 (t, *J* = 7.6 Hz, 2H), 2.54 (t, *J* = 7.8 Hz, 2H), 1.95–1.85 (m, 2H). $^{13}$C NMR (151 MHz, $CDCl_3$) δ 157.6, 141.5, 139.9, 135.8, 133.3, 131.0, 129.9, 129.8, 129.0, 126.7, 126.6, 119.9, 116.7, 34.6, 32.3, 31.7. LRMS (ESI +ve) 290 $[M+H]^+$. HRMS (ESI +ve) $C_{16}H_{16}ClNO_2$ $[M+H]^+$ calc. 290.0942, found 290.0947. HPLC 95% (AUC), $t_R$ = 8.0 min.

**OX04963a**

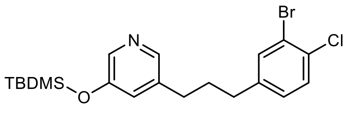


OX04963a

To a solution of **OX04956d** (4.8 g, 14.8 mmol) in chloroform (60 mL) was added TBDMSCl (3.3 g, 22.2 mmol) and imidazole (4.1 g, 59.1 mmol). The organic mixture was stirred at 60ºC for 16 h before washed with $H_2O$ (50 mL × 2), brine (50 mL × 1), dried ($Na_2SO_4$), concentrated *in vacuo*. Purification with flash column chromatography (0–30% EtOAc in petroleum ether) afforded **OX04963a** as a yellow oil (1.7 g, 26%). $^1$H NMR (400 MHz, $CDCl_3$) δ 8.07 (d, *J* = 2.7 Hz, 1H), 8.05 (d, *J* = 1.8 Hz, 1H), 7.43 (d, *J* = 2.0 Hz, 1H), 7.35 (d, *J* = 8.2 Hz, 1H), 7.04 (dd, *J* = 8.2, 2.1 Hz, 1H), 6.94 (dd, *J* = 2.6, 1.9 Hz, 1H), 2.59 (t, *J* = 7.7 Hz, 4H), 1.96–1.86 (m, 2H), 0.99 (s, 9H), 0.21 (s, 6H). $^{13}$C NMR (101 MHz, $CDCl_3$) δ 157.6, 144.1, 141.6, 139.8, 133.5, 131.4, 129.9, 129.7, 127.5, 126.6, 119.9, 34.6, 32.4, 31.8, 25.2, 20.9, 14.3. LRMS (ESI +ve) 440 $[M+H]^+$. HRMS (ESI +ve) $C_{20}H_{27}BrClNOSi$ $[M+H]^+$ calc. 440.0807, found 440.0815.

**OX04963b**

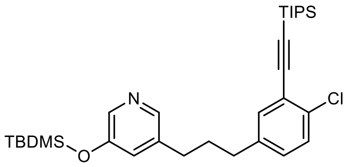

OX04963b

To a microwave vial was added **OX04963a** (773 mg, 1.8 mmol), $PPh_3$ (92 mg, 0.4 mmol) and ethynyltriisopropylsilane (352 mg, 1.9 mmol) sequentially before the addition of degassed THF/$NEt_3$- mixture (3:1, 8 mL). The mixture was degassed for 5 min before adding $Pd_2(PPh_3)_2Cl_2$ (123 mg, 0.2 mmol) and CuI (134 mg, 0.7 mmol). The mixture was degassed for another 5 min then the vial was sealed, placed on a 60°C pre-heated heating plate and stirred for 16 h. After completion, the mixture was diluted with EtOAc (40 mL), filtered through Celite® and concentrated *in vacuo*. The dried mixture was taken up with $CH_2Cl_2$ (50 mL), washed with brine (50 mL × 2), dried ($Na_2SO_4$) and concentrated *in vacuo*. The residue was then purified by flash column chromatography (0–100% $CH_2Cl_2$ in petroleum ether then 0–10% EtOAc in $CH_2Cl_2$) to afford **OX04963b** as a transparent oil (660 mg, 69%). $^1$H NMR (400 MHz, $CDCl_3$) δ 8.07 (d, *J* = 2.7 Hz, 2H), 7.31 (d, *J* = 2.2 Hz, 1H), 7.29 (d, *J* = 8.3 Hz, 1H), 7.03 (dd, *J* = 8.2, 2.2 Hz, 1H), 6.95 (dd, *J* = 2.6, 1.9 Hz, 1H), 2.63–2.55 (m, 4H), 1.96–1.86 (m, 2H), 1.15 (s, 21H), 0.99 (s, 9H), 0.21 (s, 6H). $^{13}$C NMR (101 MHz, $CDCl_3$) δ 152.2, 143.0, 140.3, 140.3, 138.0, 134.0, 133.6, 129.6, 129.3, 127.0, 123.4, 103.4, 96.6, 34.6, 32.5, 32.2, 25.8, 18.8, 18.4, 11.4, -4.3. LRMS (ESI +ve) 542 $[M+H]^+$. HRMS (ESI +ve) $C_{31}H_{48}ClNOSi_2$ $[M+H]^+$ calc. 542.3036, found 542.3040.

**OX04963**

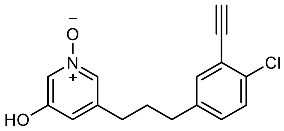

OX04963

To a solution of **OX04963b** (60 mg, 0.1 mmol) in $CH_2Cl_2$ (10 mL) was added *m*-CPBA (60 mg, 0.27 mmol) in one portion at rt. The mixture was stirred for 2 h before adding $NaHCO_3$ (20 mL, sat., aq.) to quench the reaction. The organic layer was separated and washed with $NaHCO_3$ (25 mL × 2), brine (25 mL), dried ($Na_2SO_4$) and concentrated *in vacuo*. The resulting while solid was dissolved in THF (10 mL) and TBAF (0.3 mL, 1M in THF) was added at 0ºC. The mixture was stirred for 1 h and MeOH (10 mL) was added, then the mixture was concentrated *in vacuo*. Flash column chromatography (20–100% EtOAc in petroleum ether, then 0–15% MeOH in EtOAc) afforded **OX04963** as a yellow oil (7.0 mg, 22%). $^1$H NMR (400 MHz, $CDCl_3$) δ 7.93 (t, *J* = 1.9 Hz, 1H), 7.56 (d, *J* = 1.5 Hz, 1H), 7.26 (s, 1H), 7.22 (d, *J* = 14.1 Hz, 1H), 7.00 (dd, *J* = 8.3, 2.2 Hz, 1H), 6.85 (d, *J* = 1.6 Hz, 1H), 3.29 (s, 1H), 2.49 (dt, *J* = 20.6, 7.6 Hz, 4H), 1.82 (tt, *J* = 9.2, 6.7 Hz, 2H). $^{13}$C NMR (101 MHz, $CDCl_3$) δ 157.6, 141.3, 139.8, 134.1, 133.9, 130.2, 129.8, 129.5, 126.7,

122.1, 119.8, 82.5, 80.4, 34.3, 32.2, 31.5. LRMS (ESI +ve) 288 $[M+H]^+$. HRMS (ESI +ve) $C_{16}H_{14}ClNO_2$ $[M+H]^+$ calc. 288.0786, found 288.0800. HPLC 99% (AUC), $t_R$ = 6.6 min.

**OX04964a**

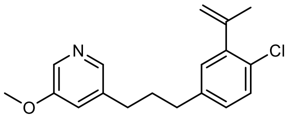

OX04964a

To a degassed microwave vial with **OX04956c** (339 mg, 1.45 mmol) and tributyl(prop-1-en-2-yl)stannane (660 mg, 2.0 mmol) in DMF (4 mL) was added $Pd(PPh_3)_4$ (173 mg, 0.15 mmol). The mixture was degassed for 5 min before the vial was sealed and placed on a pre-heated heating plate. The mixture was stirred for 24 h at 90ºC. After completion, the mixture was diluted with EtOAc (5 mL), poured into KF (5 mL, sat., aq.) then the mixture was stirred for 30 min before filtered through Celite®. The organic layer was washed with $NH_4Cl$ (20 mL × 2, sat., aq.), dried ($Na_2SO_4$) and concentrated *in vacuo*. The crude product was purified with flash column chromatography (0–30% EtOAc in petroleum ether) to afford **OX04964a** as a colourless oil (170 mg, 39%). $^1H$ NMR (400 MHz, $CDCl_3$) δ 8.15 (d, *J* = 2.8 Hz, 1H), 8.07 (d, *J* = 1.8 Hz, 1H), 7.30–7.22 (m, 1H), 7.00 (td, *J* = 3.0, 1.2 Hz, 3H), 5.22 (p, *J* = 1.6 Hz, 1H), 4.96 (dq, *J* = 1.9, 1.0 Hz, 1H), 3.85 (s, 3H), 2.62 (q, *J* = 7.2 Hz, 4H), 2.10 (dd, *J* = 1.5, 0.9 Hz, 3H), 1.98–1.91 (m, 2H). $^{13}C$ NMR (101 MHz, $CDCl_3$) δ 155.7, 144.6, 142.7, 142.5, 140.5, 138.0, 135.0, 129.9, 129.6, 129.4, 128.3, 120.8, 116.2, 55.6, 34.7, 32.5, 32.4, 23.5. LRMS (ESI +ve) 302 $[M+H]^+$. HRMS (ESI +ve) $C_{18}H_{20}ClNO$ $[M+H]^+$ calc. 302.1306, found 302.1311.

**OX04964b**

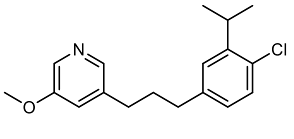

OX04964b

7 mg $PtO_2$ was added in one portion to a $N_2$ degassed solution of the **OX04964a** (130 mg, 0.43 mmol) in MeOH (180 mL) at rt, the atmosphere was replaced with $H_2$, and the reaction stirred for 1 h. After completion, the reaction was filtered through Celite® using MeOH as an eluent, concentrated *in vacuo* and purified by flash column chromatography (30% EtOAc in Pentane) to afford **OX04964b** as a colourless oil (84 mg, 64%). $^1H$ NMR (400 MHz, $CDCl_3$) δ 8.15 (d, *J* = 2.8 Hz, 1H), 8.07 (d, *J* = 1.8 Hz, 1H), 7.24 (d, *J* = 8.1 Hz, 1H), 7.07 (d, *J* = 2.2 Hz, 1H), 7.03–6.99 (m, 1H), 6.92 (dd, *J* = 8.1, 2.2 Hz, 1H), 3.85 (s, 3H), 3.38 (p, *J* = 6.9 Hz, 1H), 2.62 (td, *J* = 7.7, 3.8 Hz, 4H), 2.00–1.88 (m, 2H), 1.23 (d, *J* = 6.9 Hz, 6H). $^{13}C$ NMR (101 MHz, $CDCl_3$) δ 155.8, 145.6, 142.3, 140.6, 138.2, 134.7, 131.0, 129.4, 127.0, 126.7, 121.0, 55.6, 35.0,

32.6, 32.4, 30.2, 22.8. LRMS (ESI +ve) 304 $[M+H]^+$. HRMS (ESI +ve) $C_{18}H_{22}ClNO$ $[M+H]^+$ calc. 304.1463, found 304.1467.

**OX04964c**

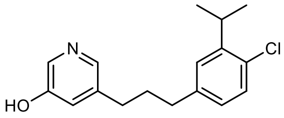

OX04964c

3 mL 48% HBr in $H_2O$ was added to **OX04964b** (80 mg, 0.26 mmol) and then the mixture was stirred at 120ºC for 48 h. After completion of the reaction, cooled to rt, the mixture was neutralized with sat. aq. $NaHCO_3$ and extracted with EtOAc (3 × 30 mL). The combined organic extracts were washed with brine (100 mL), dried over $Na_2SO_4$, filtered, and concentrated under reduced pressure. The resulting residue was purified by flash column chromatography (5% MeOH in $CH_2Cl_2$) afforded **OX04964c** as a brown solid (46 mg, 60%). $^1H$ NMR (400 MHz, $CDCl_3$) δ 8.14 (d, *J* = 2.7 Hz, 1H), 7.92 (d, *J* = 1.8 Hz, 1H), 7.23 (d, *J* = 8.1 Hz, 1H), 7.16–7.13 (m, 1H), 7.06 (d, *J* = 2.2 Hz, 1H), 6.91 (dd, *J* = 8.1, 2.2 Hz, 1H), 3.37 (hept, *J* = 6.7 Hz, 1H), 2.66–2.57 (m, 4H), 1.99–1.87 (m, 2H), 1.23 (d, *J* = 6.9 Hz, 6H). $^{13}C$ NMR (101 MHz, $CDCl_3$) δ 155.2, 145.7, 140.5, 139.8, 139.6, 134.0, 131.0, 129.5, 127.0, 126.8, 125.2, 35.0, 32.4, 32.3, 30.2, 22.8. LRMS (ESI +ve) 290 $[M+H]^+$. HRMS (ESI +ve) $C_{17}H_{20}ClNO$ $[M+H]^+$ calc. 290.1306

, found 290.1313.

**OX04964**

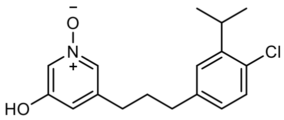

OX04964

To a solution of **OX04964c** (30 mg, 0.1 mmol) in $CH_2Cl_2$ (10 mL) at rt was added *m*-CPBA (1.5 eq.) and the resulting mixture stirred for 2 h. After completion, the reaction was quenched with $NaHCO_3$ (sat., aq.). The organic phase was washed with brine, dried ($Na_2SO_4$), filtered and concentrated *in vacuo*. Purification by flash column chromatography (0–8% MeOH in $CH_2Cl_2$) afforded **OX04964** as a yellow oil (19 mg, 60%). $^1H$ NMR (400 MHz, $CDCl_3$) δ 8.00 (s, 1H), 7.63 (s, 1H), 7.24 (d, *J* = 8.1 Hz, 1H), 7.05 (d, *J* = 2.2 Hz, 1H), 6.93–6.87 (m, 2H), 3.37 (p, *J* = 6.8 Hz, 1H), 2.56 (dt, *J* = 23.1, 7.7 Hz, 4H), 1.90 (h, *J* = 6.8 Hz, 2H), 1.23 (d, *J* = 6.9 Hz, 6H). $^{13}C$ NMR (151 MHz, $CDCl_3$) δ 157.5, 145.8, 141.6, 140.0, 131.3, 130.0, 129.6, 126.9, 126.7, 126.6, 119.9, 34.8, 32.3, 31.8, 30.2, 22.8. **LRMS (ESI +ve)** 306 $[M+H]^+$. **HRMS (ESI +ve)** $C_{17}H_{20}ClNO_2$ $[M+H^+]$ calc. 306.1255, found 306.1267. HPLC 99% (AUC), $t_R$ = 6.5 min.

**OX04965a**

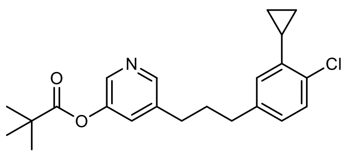

OX04965a

To $CH_2Cl_2$ (4 mL) was added $Et_2Zn$ (1.3 mL, 1.0 M in hexane). The solution was cooled in an ice bath and TFA (140 mg, 1.2 mmol) was then added dropwise into the reaction mixture. Upon stirring for 30 min, a solution of $CH_2I_2$ (330 mg, 1.2 mmol) was added at 0ºC. After an additional 20 min of stirring, a solution of **OX04962b** (44 mg, 0.1 mmol) in $CH_2Cl_2$ (4 mL) was added, and the ice bath was removed. The reaction mixture was stirred for 16 h before it was quenched with $NaHCO_3$ (10 mL, sat., aq.) and the phases were separated. The organic layer was washed with $NH_4Cl$ (20 mL × 2, sat., aq.), brine (15 mL) and then dried ($Na_2SO_4$), filtered and concentrated *in vacuo*. The residue was purified by flash column chromatography (10–18% EtOAc in petroleum ether) to afford **OX04965a** as a beige oil (32 mg, 92%). $^1$H NMR (400 MHz, $CDCl_3$) δ 8.30 (d, $J$ = 1.8 Hz, 1H), 8.23 (d, $J$ = 2.5 Hz, 1H), 7.27–7.20 (m, 2H), 6.91 (dd, $J$ = 8.1, 2.2 Hz, 1H), 6.72 (d, $J$ = 2.2 Hz, 1H), 2.64 (t, $J$ = 7.7 Hz, 2H), 2.58 (t, $J$ = 7.7 Hz, 2H), 2.22–2.12 (m, 1H), 1.98–1.87 (m, 2H), 1.37 (s, 9H), 1.03–0.96 (m, 2H), 0.70–0.63 (m, 2H). $^{13}$C NMR (101 MHz, $CDCl_3$) δ 176.8, 147.7, 147.1, 141.0, 140.8, 140.2, 138.2, 132.9, 129.1, 129.1, 126.8, 126.3, 39.3, 34.9, 32.4, 32.1, 27.2, 13.4, 8.1. LRMS (ESI +ve) 372 $[M+H]^+$. HRMS (ESI +ve) $C_{22}H_{26}ClNO_2$ $[M+H]^+$ calc. 372.1725, found 372.1713.

**OX04965**

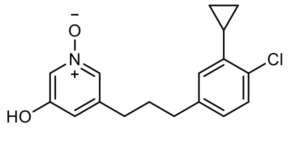

OX04965

To a mixture of **OX04965a** (20 mg, 0.05 mmol) in $CH_2Cl_2$ (10 mL) was added *m*-CPBA (1.5 eq.) in one portion at rt. The mixture was stirred at rt for 2 h then concentrated *in vacuo*, followed by purification with flash column chromatography (20–100% EtOAc in petroleum ether then 0–20% MeOH in EtOAc). The MeOH in EtOAc fractions were checked by thin layer chromatography, collected, concentrated *in vacuo* and dissolved in MeOH (10 mL). 4M aq. NaOH (2 mL) was added subsequently and the mixture was stirred for 16 h at rt. After completion, the mixture was concentrated *in vacuo*, taken up with sat. aq. $NH_4Cl$ to adjust pH < 8 and extracted with $CH_2Cl_2$ (3 × 20 mL). The organic layer was washed with brine, dried with anhydrous $Na_2SO_4$, concentrated *in vacuo* and purified with flash column chromatography (0–10% MeOH in $CH_2Cl_2$) to afford **OX04965** as a white solid (14 mg, 86%). $^1$H NMR (400 MHz, $CDCl_3$) δ 8.01 (t, $J$ = 1.9 Hz, 1H), 7.63 (d, $J$ = 1.5 Hz, 1H), 7.24 (d, $J$ = 8.1 Hz, 1H), 6.93–6.83 (m, 2H), 6.69 (d, $J$ = 2.1 Hz, 1H), 2.52 (dt, $J$ = 14.7, 7.7 Hz, 4H), 2.16 (tt, $J$ = 8.5, 5.3 Hz, 1H), 1.85 (tt, $J$ = 9.3, 6.7 Hz, 2H), 1.05–0.93 (m, 2H), 0.72–0.62 (m, 2H). $^{13}$C NMR (101 MHz, $CDCl_3$) δ 157.5, 141.6, 141.0, 139.8, 133.1, 129.9, 129.2, 126.7,

126.6, 126.3, 119.9, 34.7, 32.3, 31.8, 13.4, 8.1. LRMS (ESI +ve) 304 $[M+H]^+$. HRMS (ESI +ve) $C_{17}H_{18}ClNO_2$ $[M+H]^+$ calc. 304.1099, found 304.1097. HPLC 95% (AUC), $t_R$ = 8.0 min.

**OX04966a**

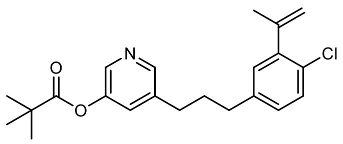

OX04966a

To an N2 degassed mixture of **OX04962a** (839 mg, 2.05 mmol) and tributyl(1-methylethenyl)Stannane (1362 mg, 4.10 mmol) in DMF (4.5 mL) was added $Pd(PPh_3)_4$ (118 mg, 0.10 mmol). The mixture was subsequently degassed for another 5 min before the mixture was heated to 90ºC and stirred for 72 h. After completion, the mixture was poured into sat. aq. KF (10 mL), stirred for 30 min and EtOAc (2 × 30 mL) was added to extract the organic layer. The organic layer was then washed with sat. aq. KF (1 × 10 mL), water (3 × 30 mL), brine (1 × 50 mL), dried over $Na_2SO_4$ and saturated *in vacuo*. The resulting brown oil was purified with flash column chromatography (0–30% EtOAc in petroleum ether) to afford **OX04966a** as a yellow oil (677 mg, 89%). $^1$H NMR (400 MHz, $CDCl_3$) δ 8.30 (s, 1H), 8.22 (s, 1H), 7.26–7.23 (m, 2H), 7.02–6.98 (m, 2H), 5.21 (p, *J* = 1.6 Hz, 1H), 4.95 (dq, *J* = 1.9, 0.9 Hz, 1H), 2.64 (dt, *J* = 17.2, 7.7 Hz, 4H), 2.09 (dd, *J* = 1.5, 0.9 Hz, 3H), 2.01–1.90 (m, 2H), 1.36 (s, 9H). $^{13}$C NMR (101 MHz, $CDCl_3$) δ 176.8, 147.1, 144.6, 142.8, 141.0, 140.3, 129.9, 129.6, 129.4, 129.1, 128.3, 116.2, 39.3, 34.7, 32.4, 32.2, 27.2, 23.5. **HRMS (ESI +ve)** $C_{22}H_{26}ClNO_2$ $[M+H^+]$ calc. 372.1725, found 372.1742.

**OX04966b**

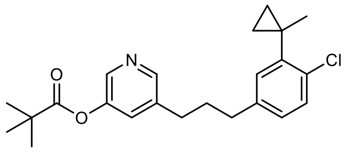

OX04966b

To an mixture of $ZnEt_2$ (8.1 mL, 1M in hexane) in $CH_2Cl_2$ (18 mL) was added TFA (0.62 mL) dropwise at 0ºC and the resulting white slurry was stirred vigorously for 30 min at 0ºC. Diiodomethane (0.65 mL) was added dropwise to the mixture at 0ºC to from a Clear solution, followed by stirring for 30 min. Then **OX04966a** (300 mg, 0.8 mmol) in $CH_2Cl_2$ (5 mL) was added to the mixture at 0ºC, the ice bath was removed and the reaction mixture was stirred for 16 h. After completion, the mixture was poured into sat. $NaHCO_3$ (50 mL) and separated. The organic layer was washed with sat. aq. $NH_4Cl$ (2 × 50 mL), brine (1 × 30 mL) and concentrated *in vacuo*. Purification with flash column chromatography (0–20% EtOAc in petroleum ether) afforded **OX04966b** as a colorless oil (98 mg, 31%). $^1$H NMR (400 MHz, $CDCl_3$) δ 8.23 (d, *J* = 1.9 Hz, 1H), 8.15 (d, *J* = 2.5 Hz, 1H), 7.18 (d, *J* = 2.4 Hz, 1H), 7.14 (d, *J* = 8.1 Hz, 1H), 7.07 (d, *J* = 2.2 Hz, 1H), 6.86 (dd, *J* = 8.1, 2.3 Hz, 1H), 2.55 (dt, *J* = 20.9, 7.7 Hz, 4H), 1.92–1.83 (m, 2H), 1.29 (s, 9H), 1.26 (s, 3H), 0.76–0.70 (m, 2H), 0.70–0.63 (m, 2H). $^{13}$C NMR (101 MHz, $CDCl_3$) δ 176.8, 147.7, 147.0, 143.9,

140.9, 140.2, 138.2, 133.2, 131.5, 129.6, 129.0, 127.6, 39.3, 34.7, 32.4, 32.2, 27.2, 25.2, 20.9, 14.2. HRMS (ESI +ve) $C_{23}H_{28}ClNO_2$ $[M+H]^+$ calc. 386.1881, found 386.1906.

**OX04966**

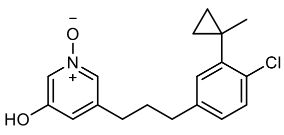

OX04966

To a mixture of **OX04966b** (98 mg, 0.3 mmol) in $CH_2Cl_2$ (30 mL) was added *m*-CPBA (1.5 eq.) in one portion at rt. The mixture was stirred at rt for 2 h then concentrated *in vacuo*, followed by purification with flash column chromatography (20–100% EtOAc in petroleum ether then 0–20% MeOH in EtOAc). The MeOH in EtOAc fractions were checked by thin layer chromatography, collected, concentrated *in vacuo* and dissolved in MeOH (10 mL). 4M aq. NaOH (5 mL) was added subsequently and the mixture was stirred for 16 h at rt. After completion, the mixture was concentrated *in vacuo*, taken up with sat. aq. $NH_4Cl$ to adjust pH < 8 and extracted with $CH_2Cl_2$ (3 × 20 mL). The organic layer was washed with brine, dried with anhydrous $Na_2SO_4$, concentrated *in vacuo* and purified with flash column chromatography (0–10% MeOH in $CH_2Cl_2$) to afford **OX04966** as a white solid (80 mg, 99%). $^1$H NMR (400 MHz, $CDCl_3$) δ 8.01 (s, 1H), 7.63 (s, 1H), 7.22 (d, $J$ = 8.1 Hz, 1H), 7.13 (d, $J$ = 2.3 Hz, 1H), 6.94–6.89 (m, 2H), 2.55 (dt, $J$ = 15.1, 7.7 Hz, 4H), 1.95–1.84 (m, 2H), 1.33 (s, 3H), 0.83–0.77 (m, 2H), 0.77–0.71 (m, 2H). $^{13}$C NMR (101 MHz, $CDCl_3$) δ 157.6, 144.1, 141.6, 139.8, 133.5, 131.4, 129.9, 129.7, 127.5, 126.6, 119.9, 34.6, 32.4, 31.8, 25.2, 20.9, 14.3. LRMS (ESI +ve) 318 $[M+H]^+$. HRMS (ESI +ve) $C_{18}H_{20}ClNO_2$ $[M+H]^+$ calc. 318.1255, found 318.1253. HPLC 99% (AUC), $t_R$ = 8.6 min.

**References**


1. Makino, K. *et al.* Chemoselective Demethylation of Methoxypyridine. *Synlett* **30**, 951–954 (2019).

### NMR

# OX04539a

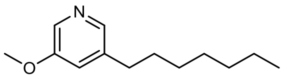

**OX04539a**

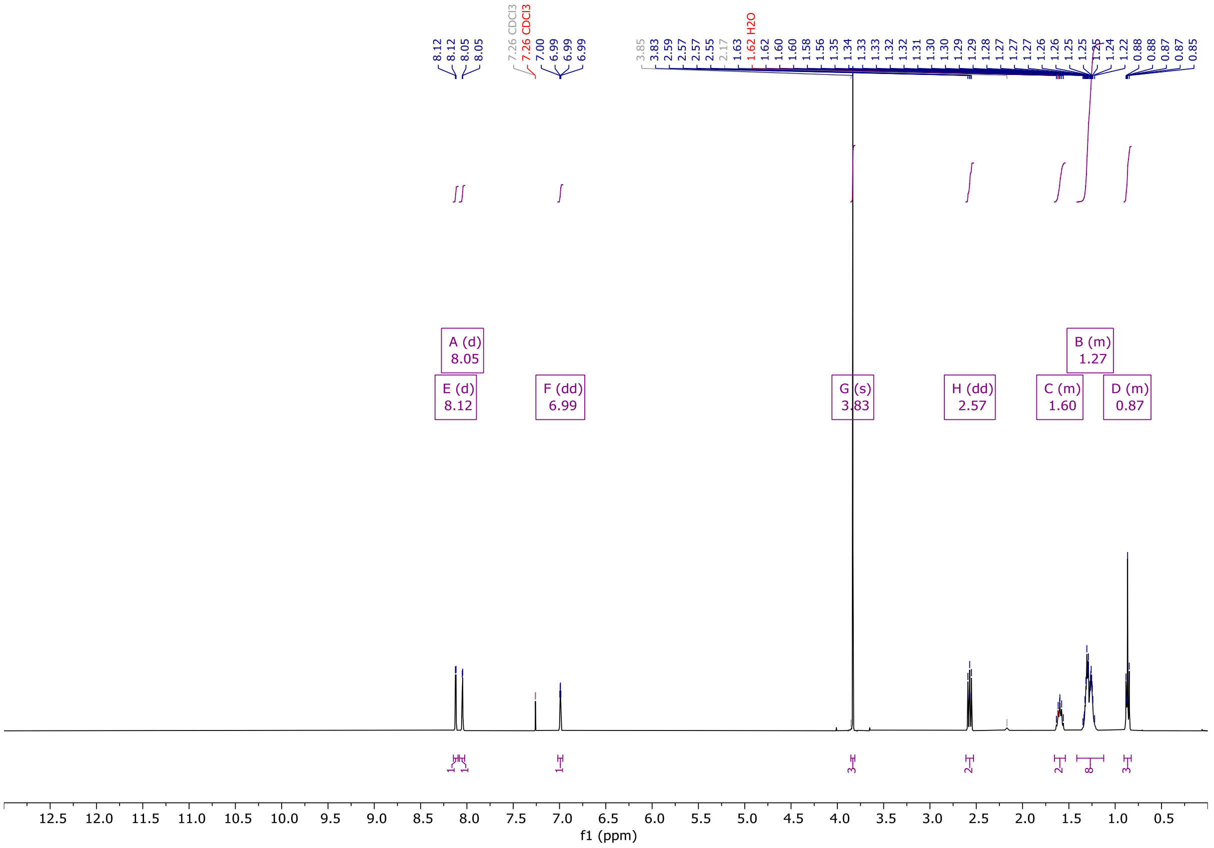

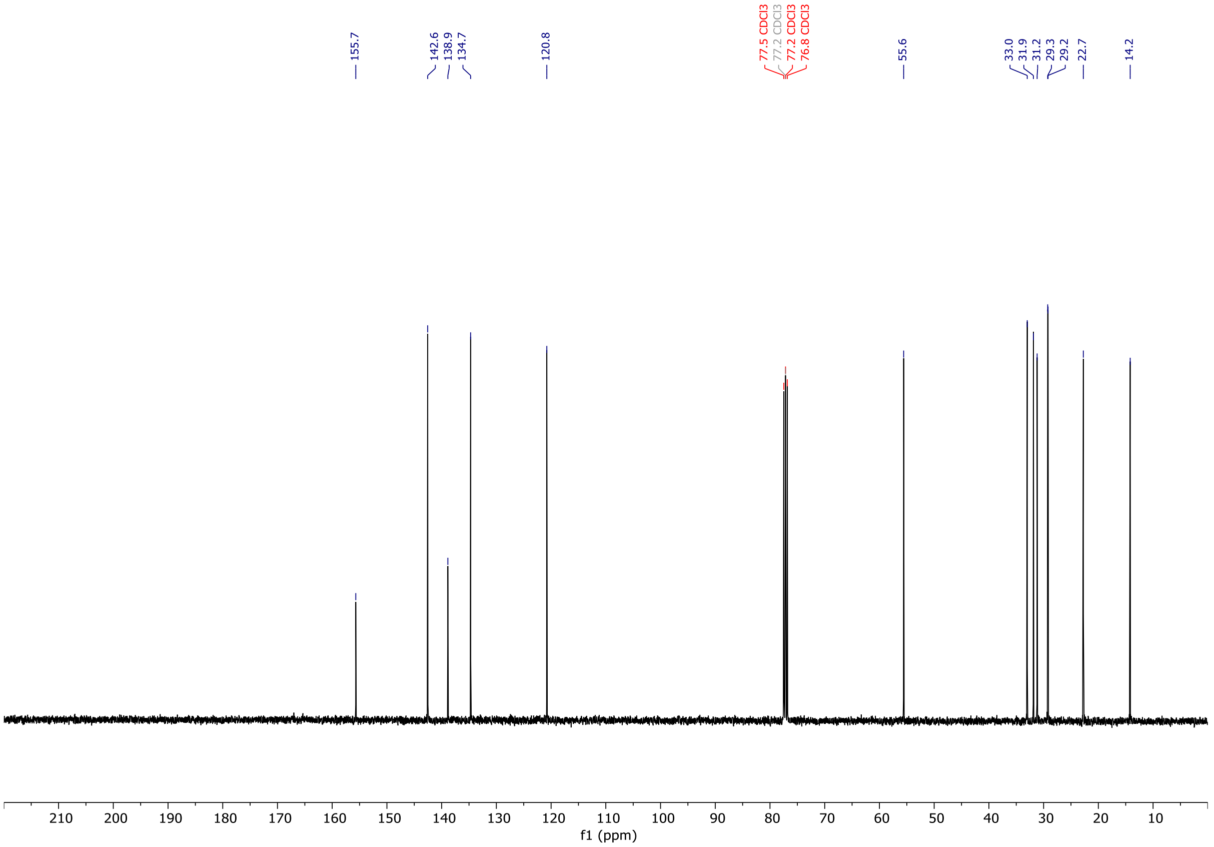

# OX04539b

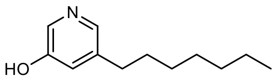


**OX04539b**

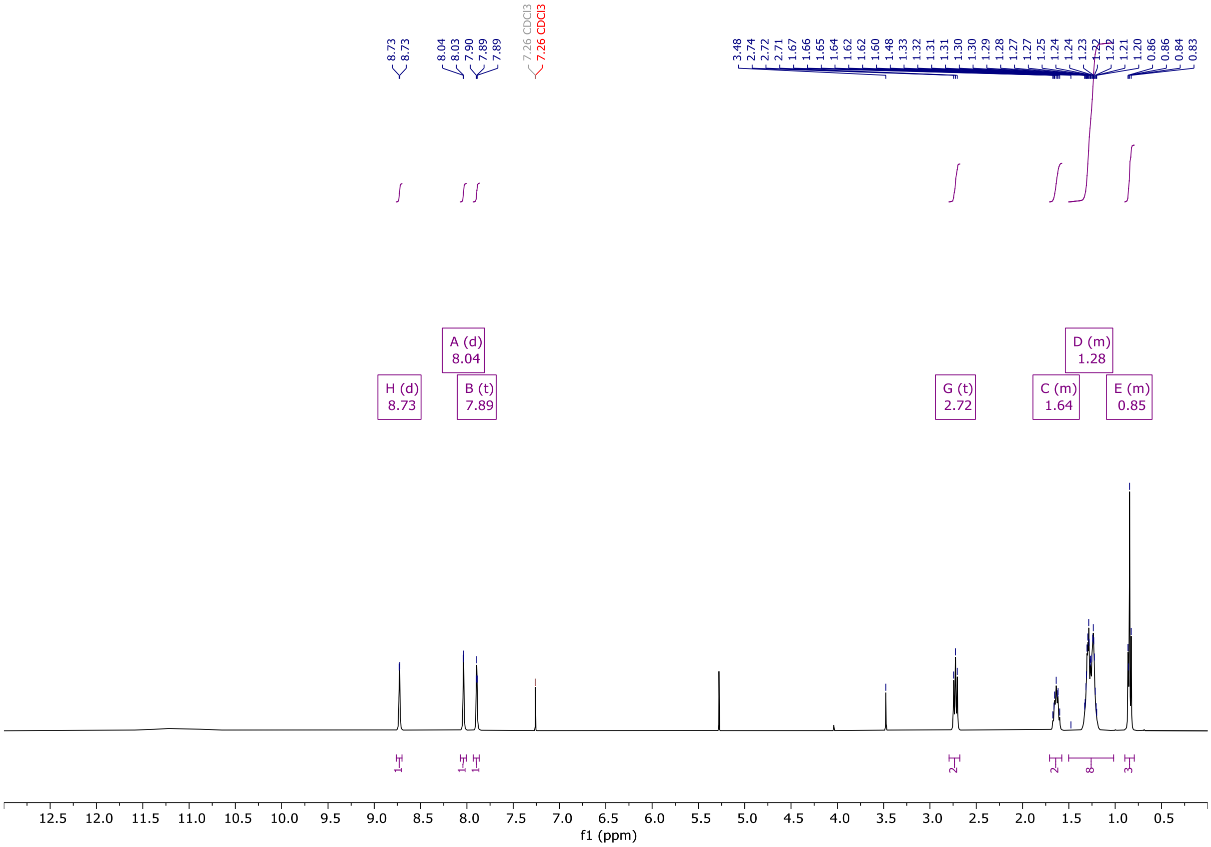


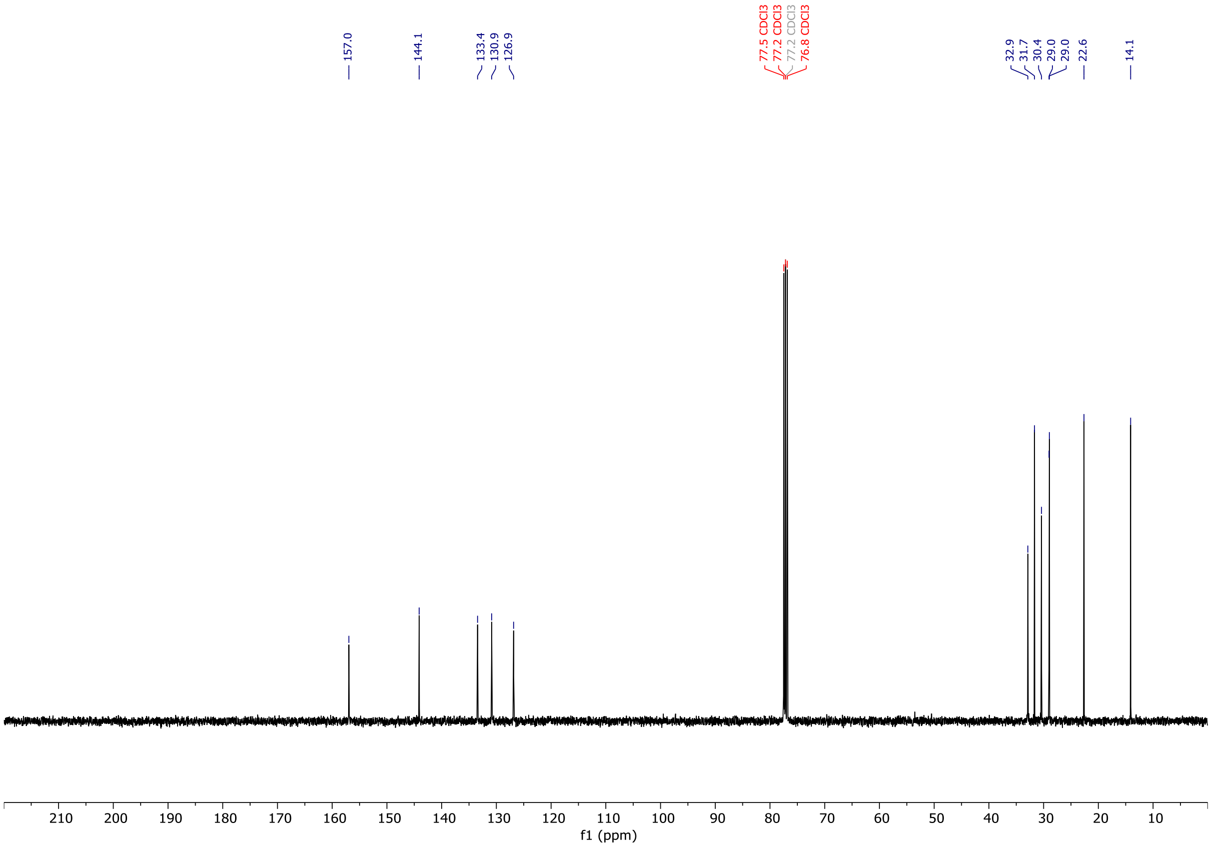

# OX04539

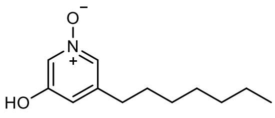


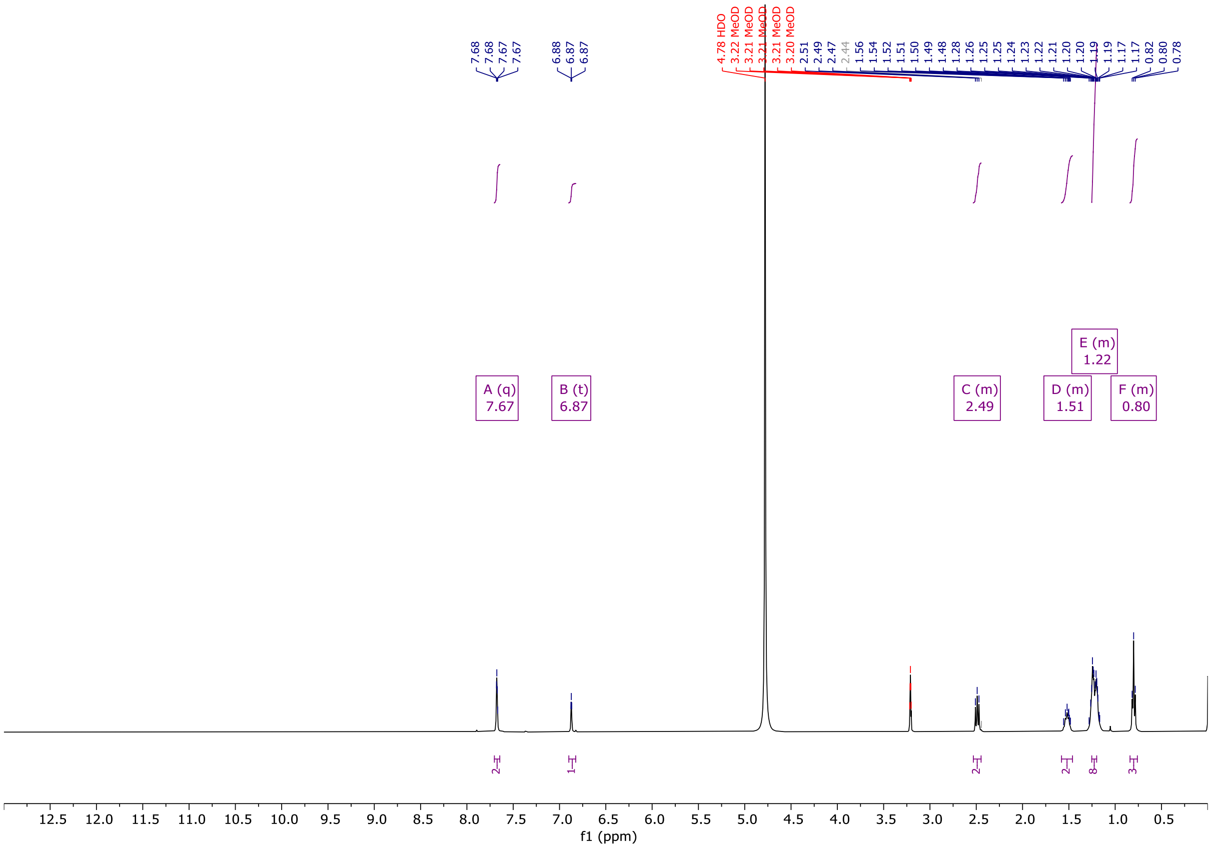

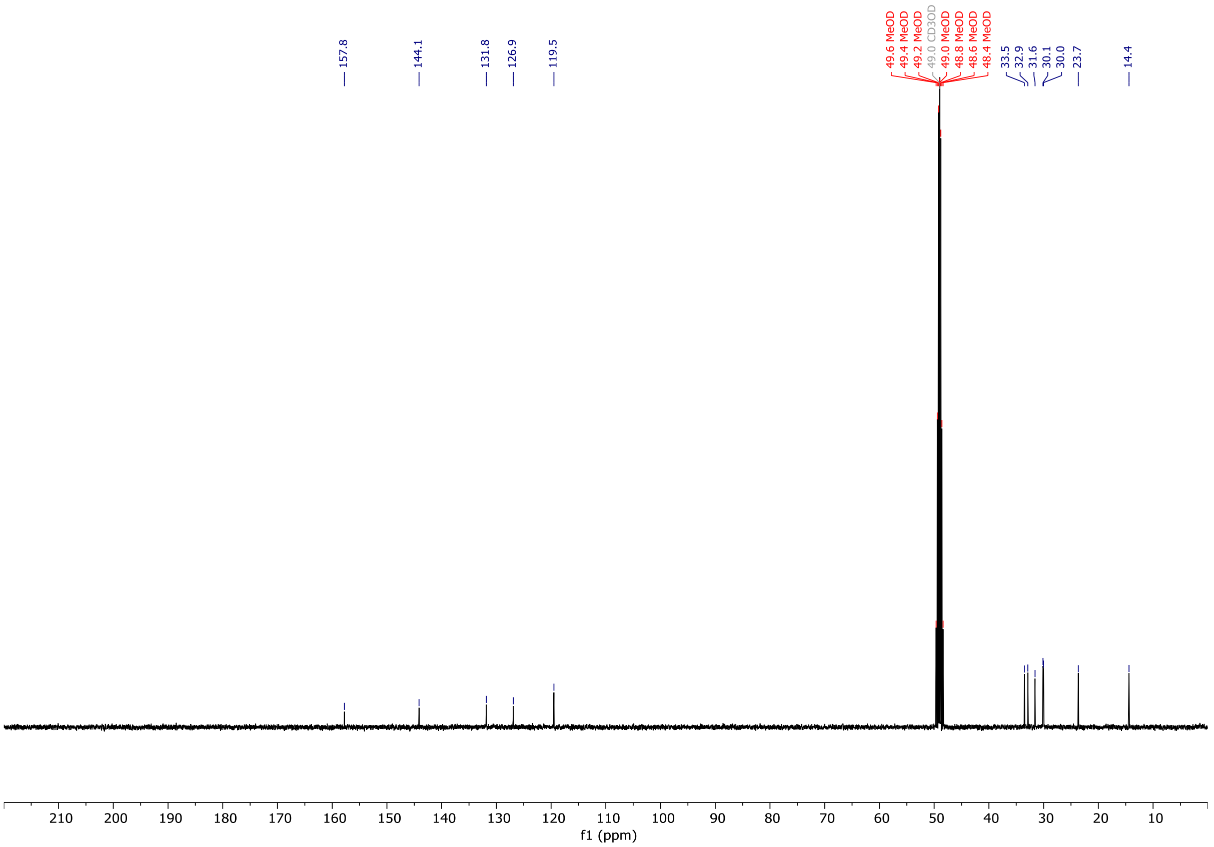

# OX04540a

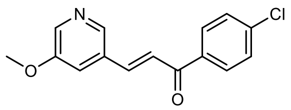


**OX04540a**

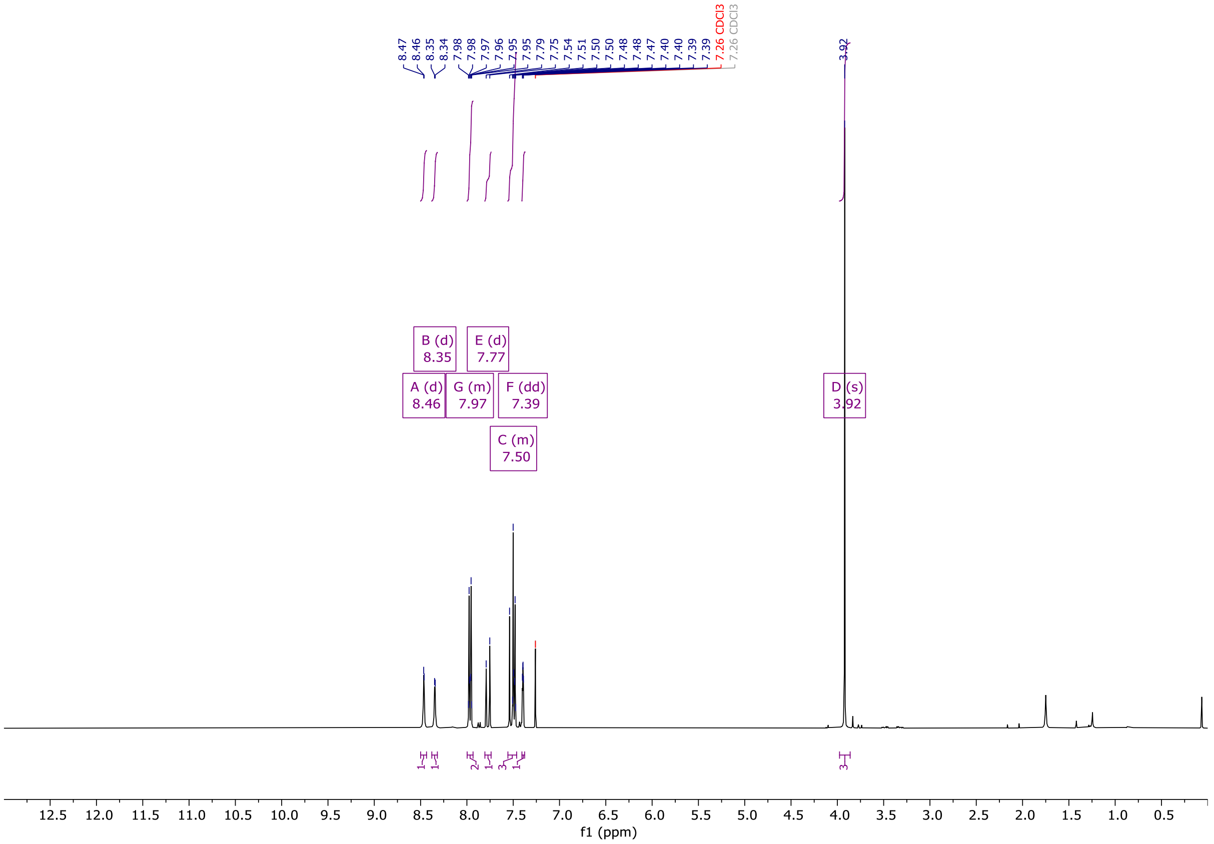


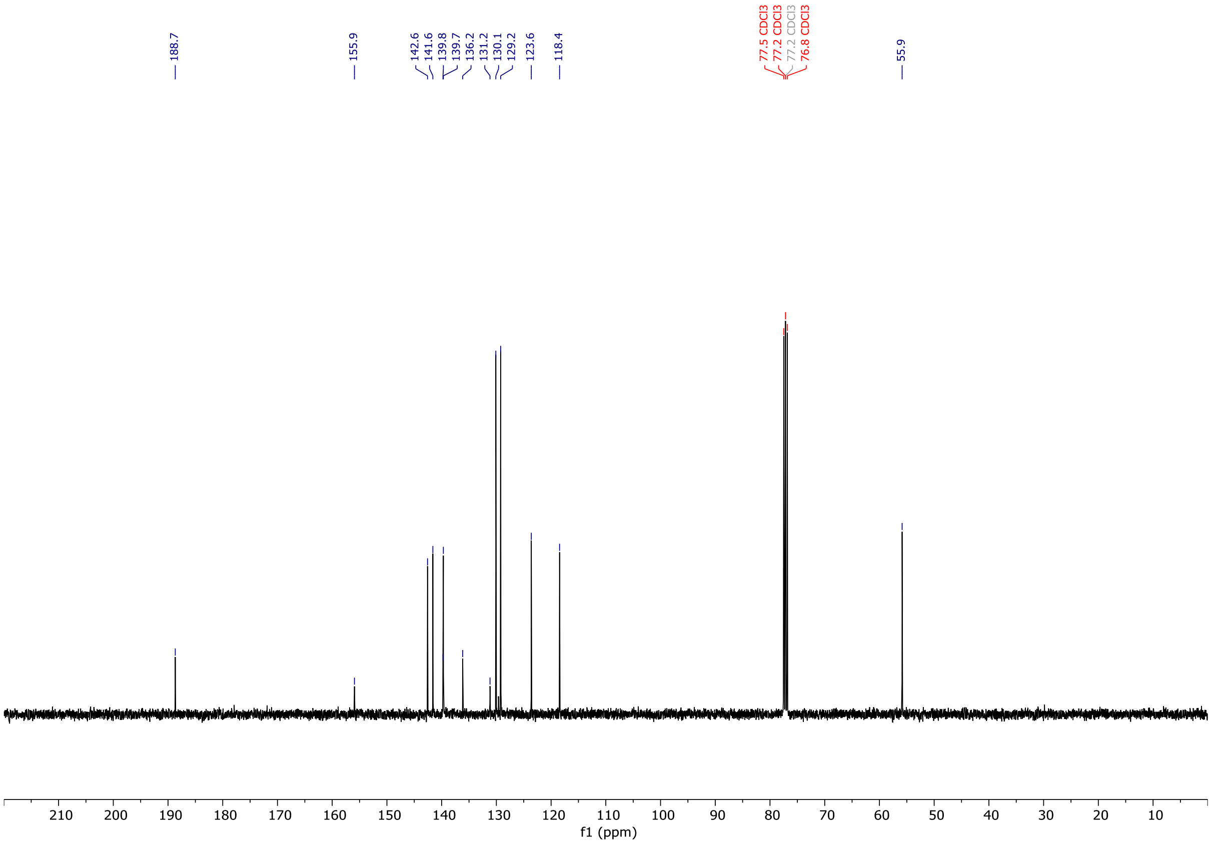

# OX04540b

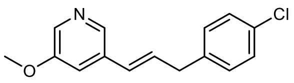

**OX04540b**

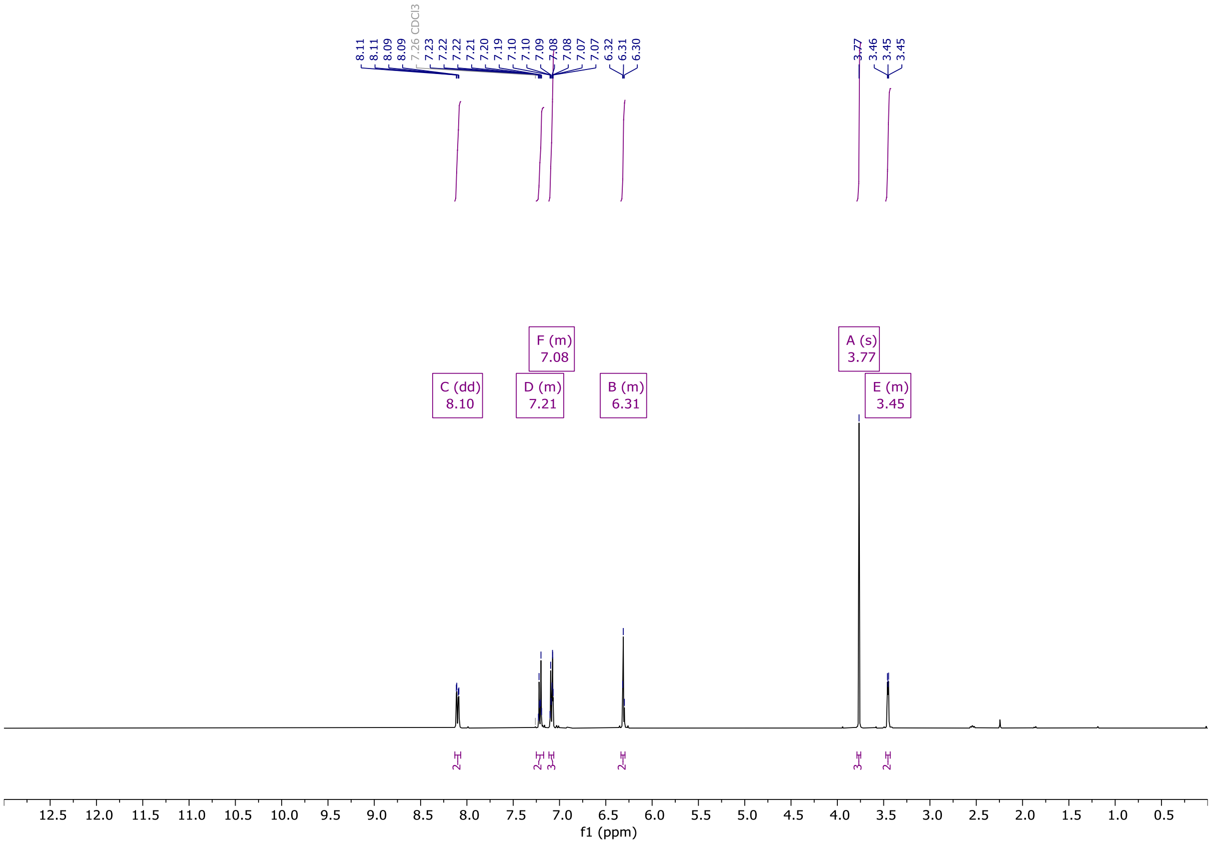


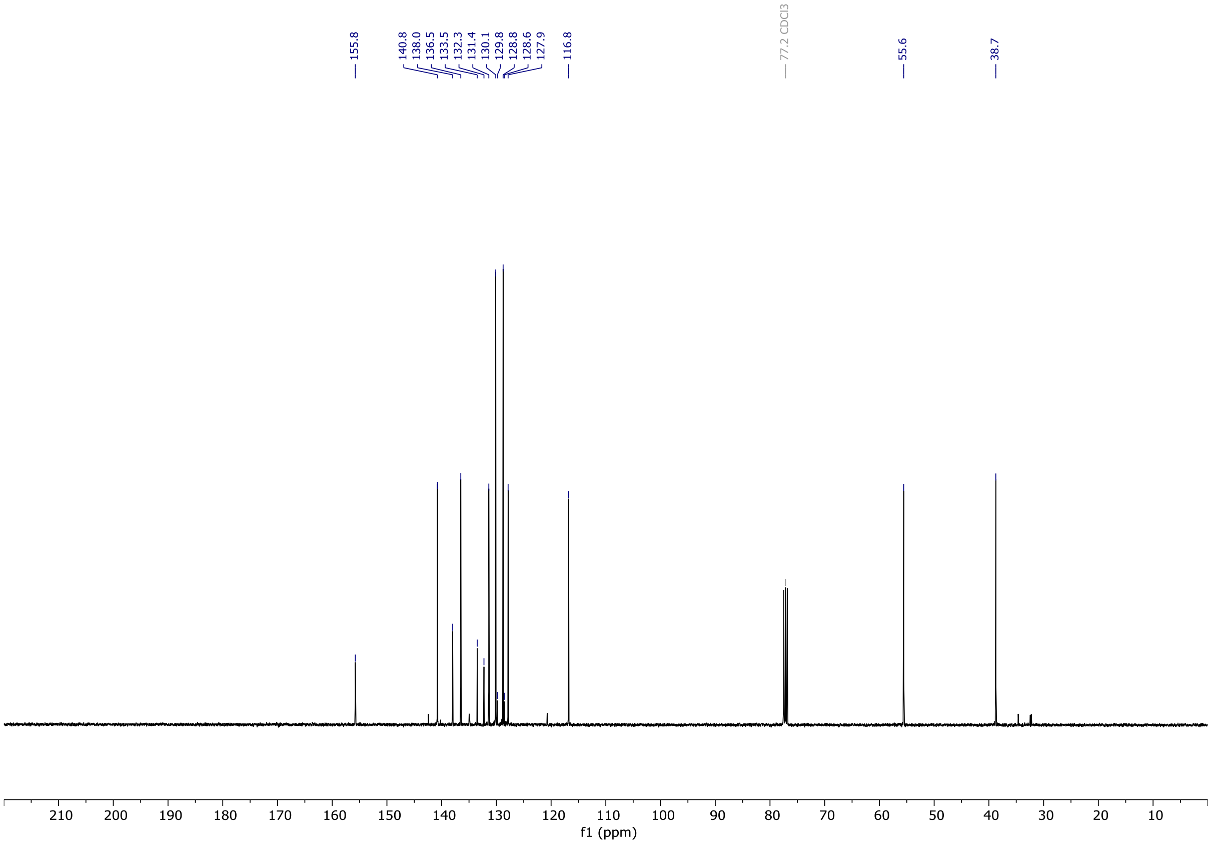

# OX04540c

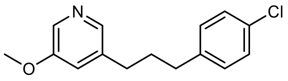


**OX04540c**

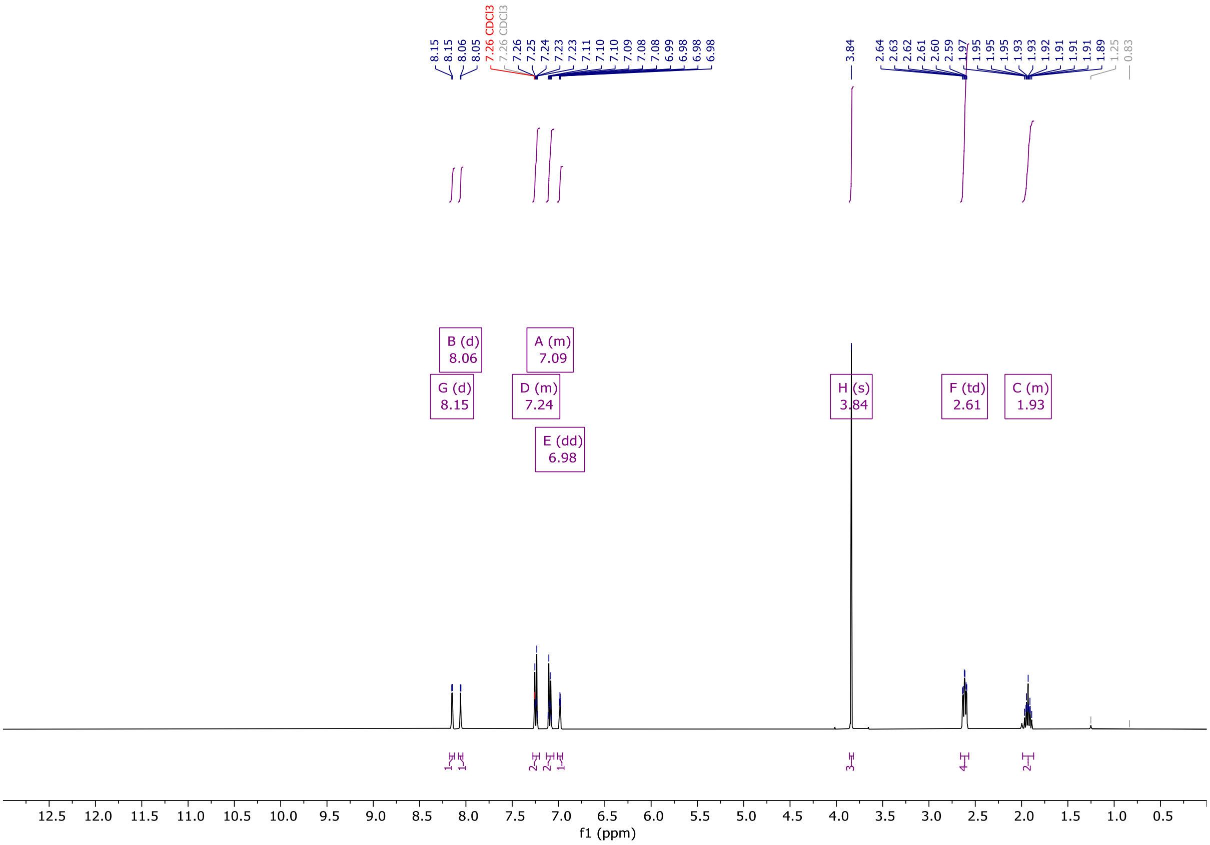


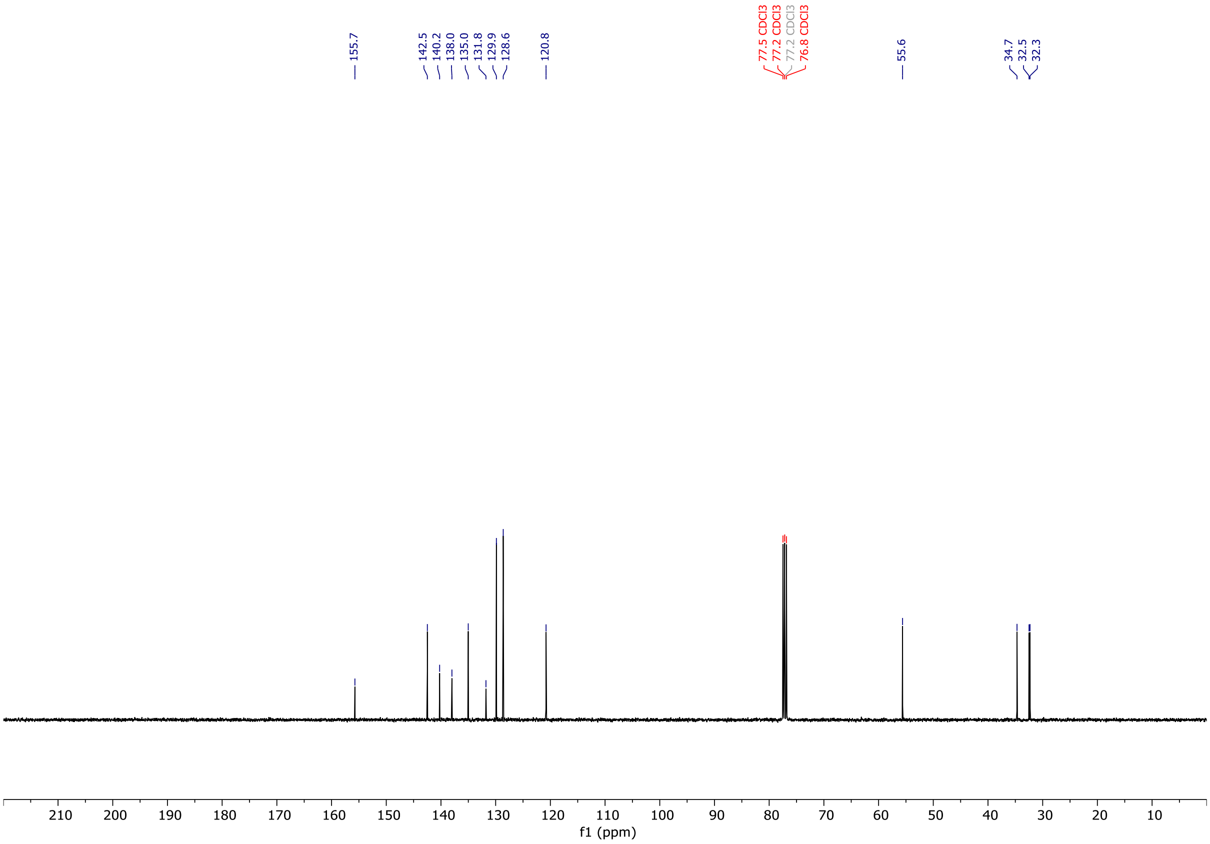

# OX04540d

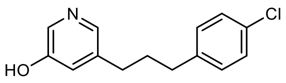

OX04540d

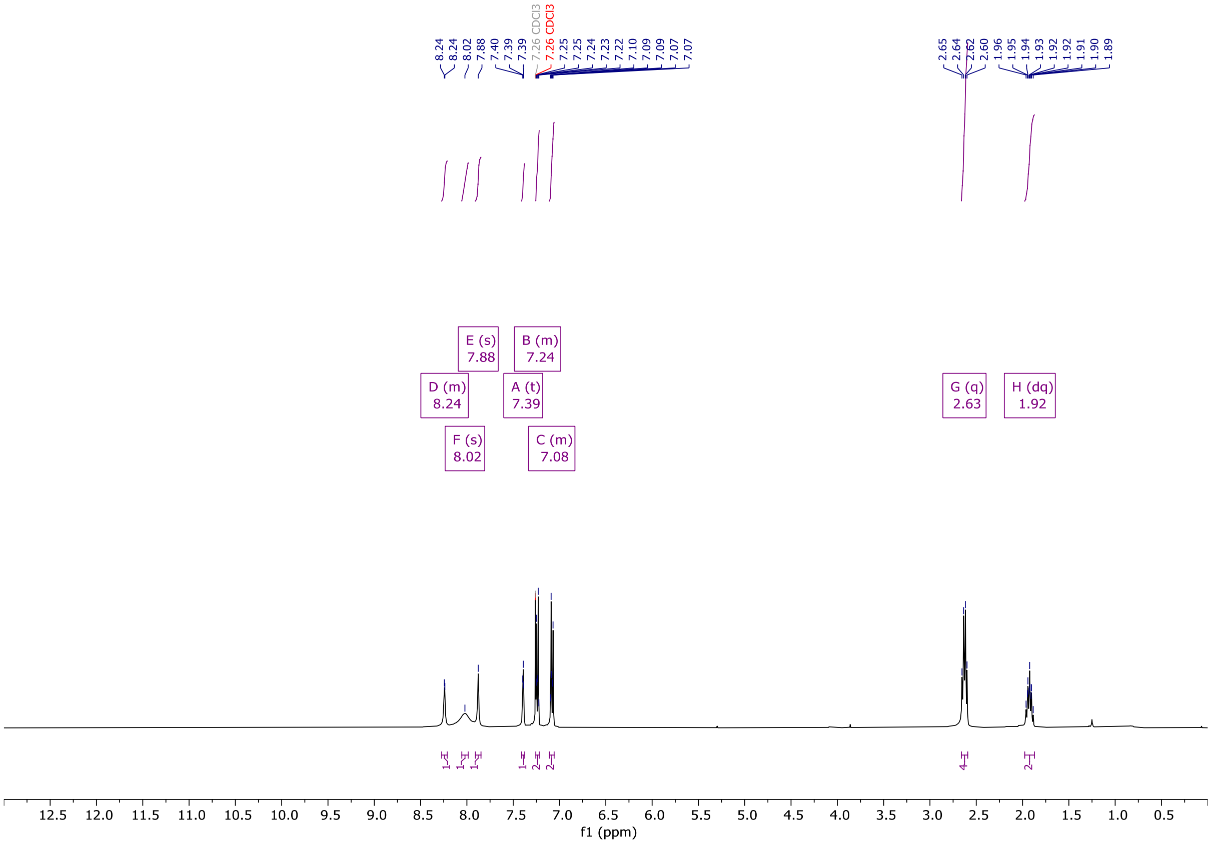

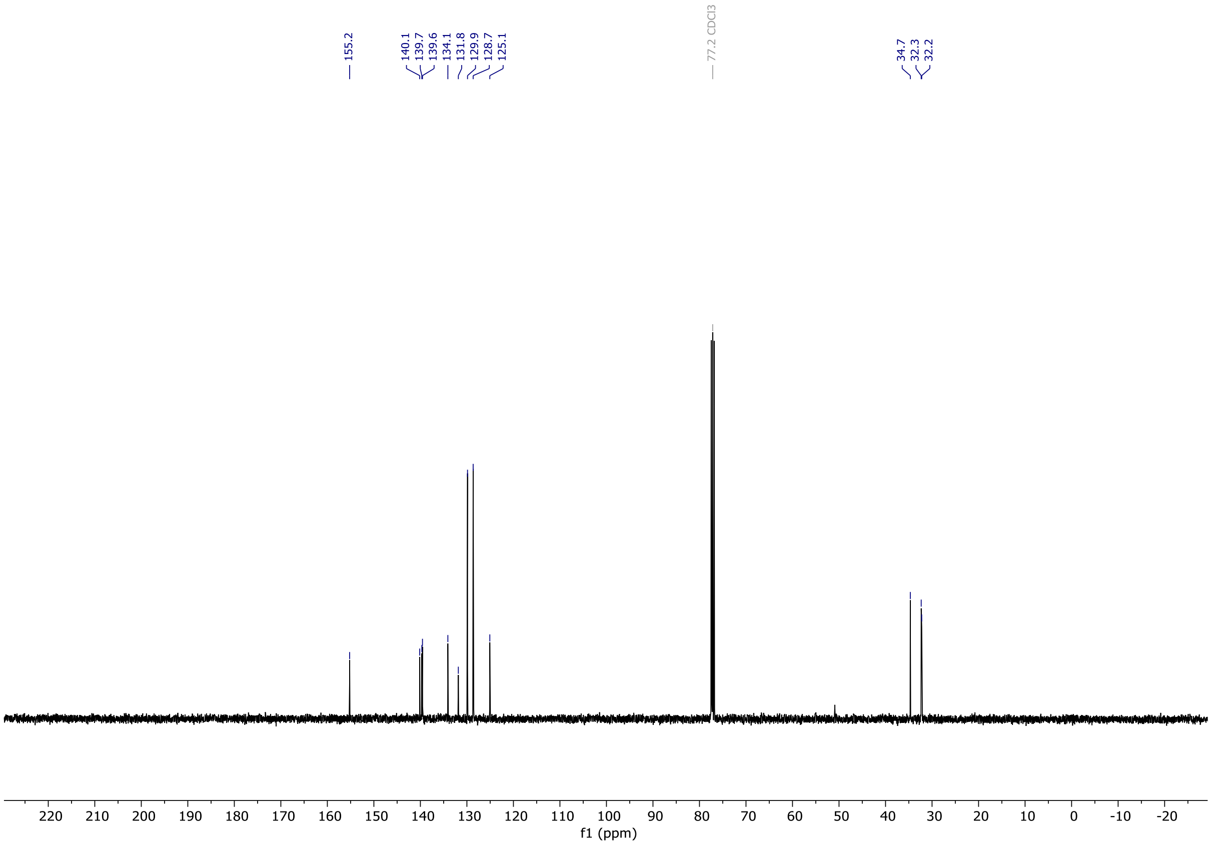

# OX04540

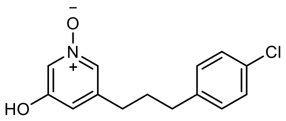


**OX04540**

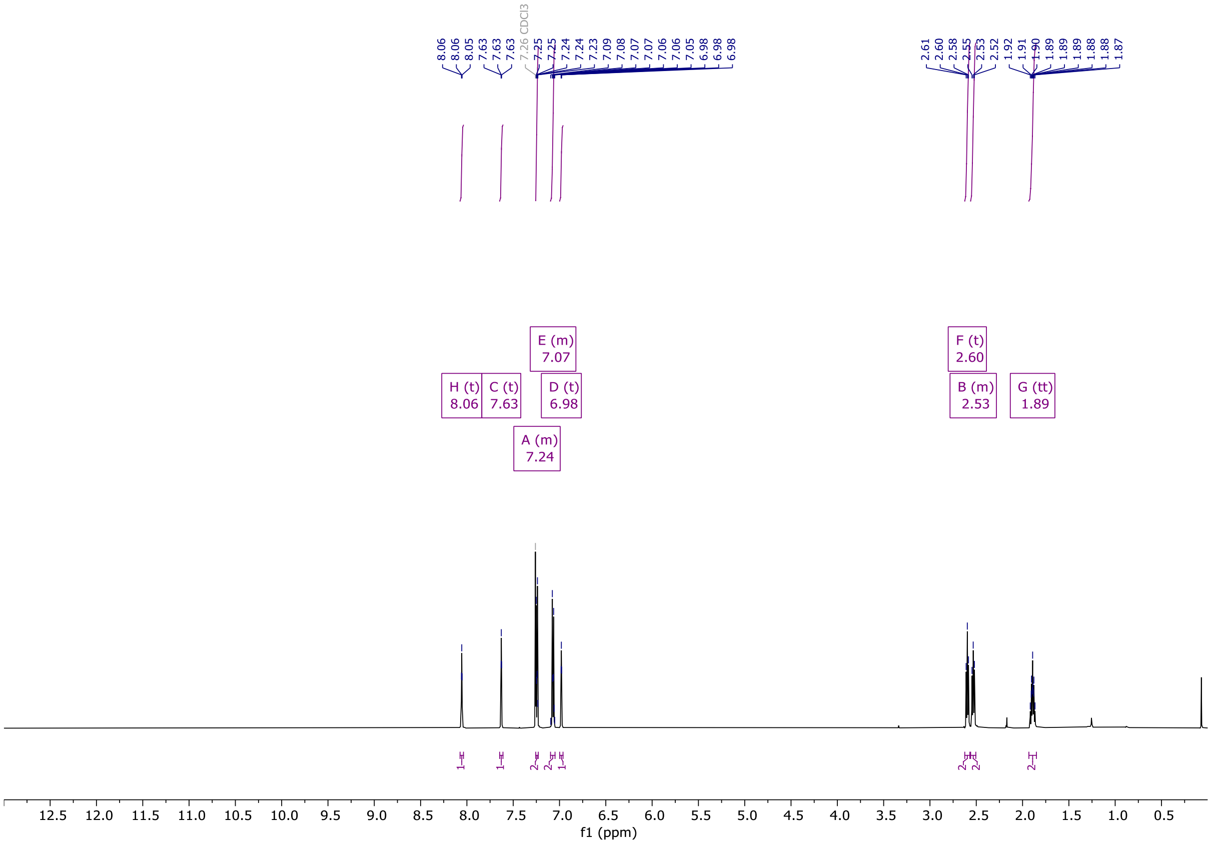


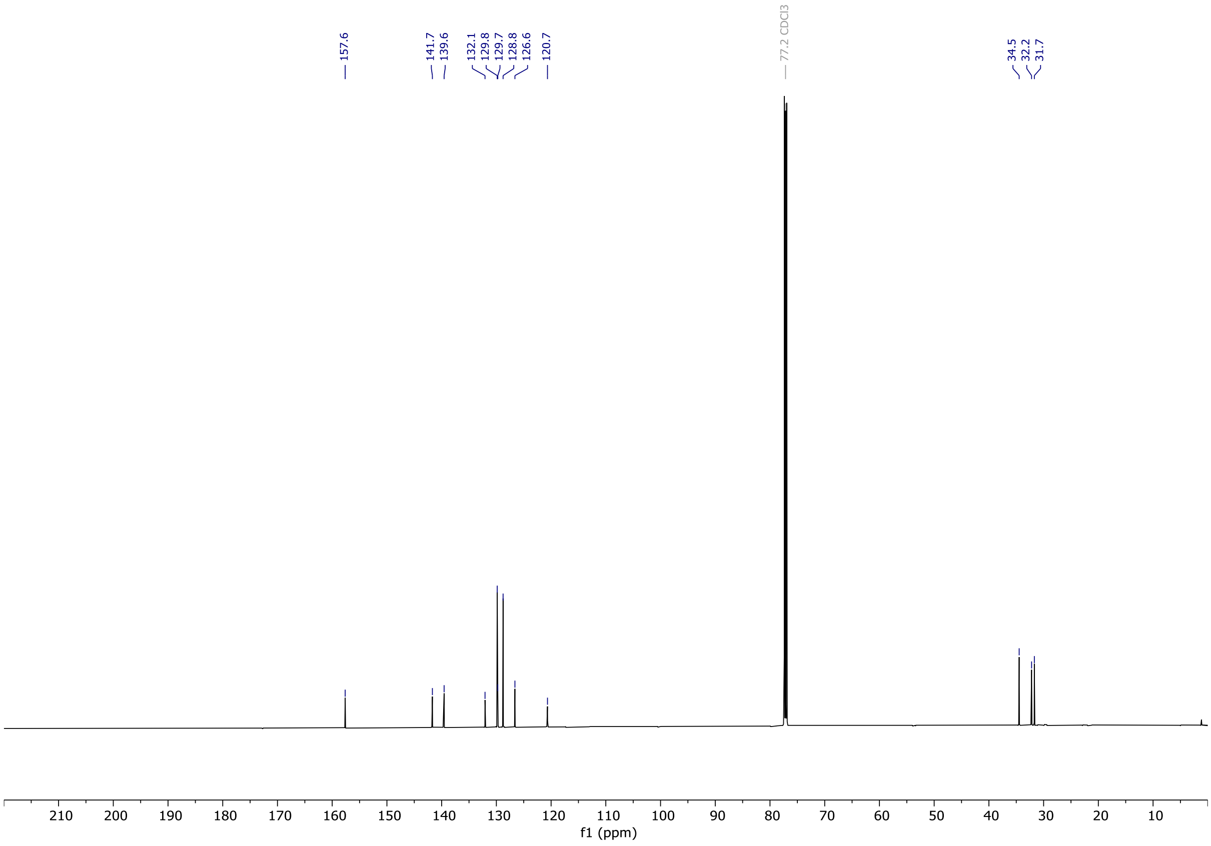

**OX04954a**

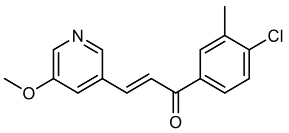


OX04954a

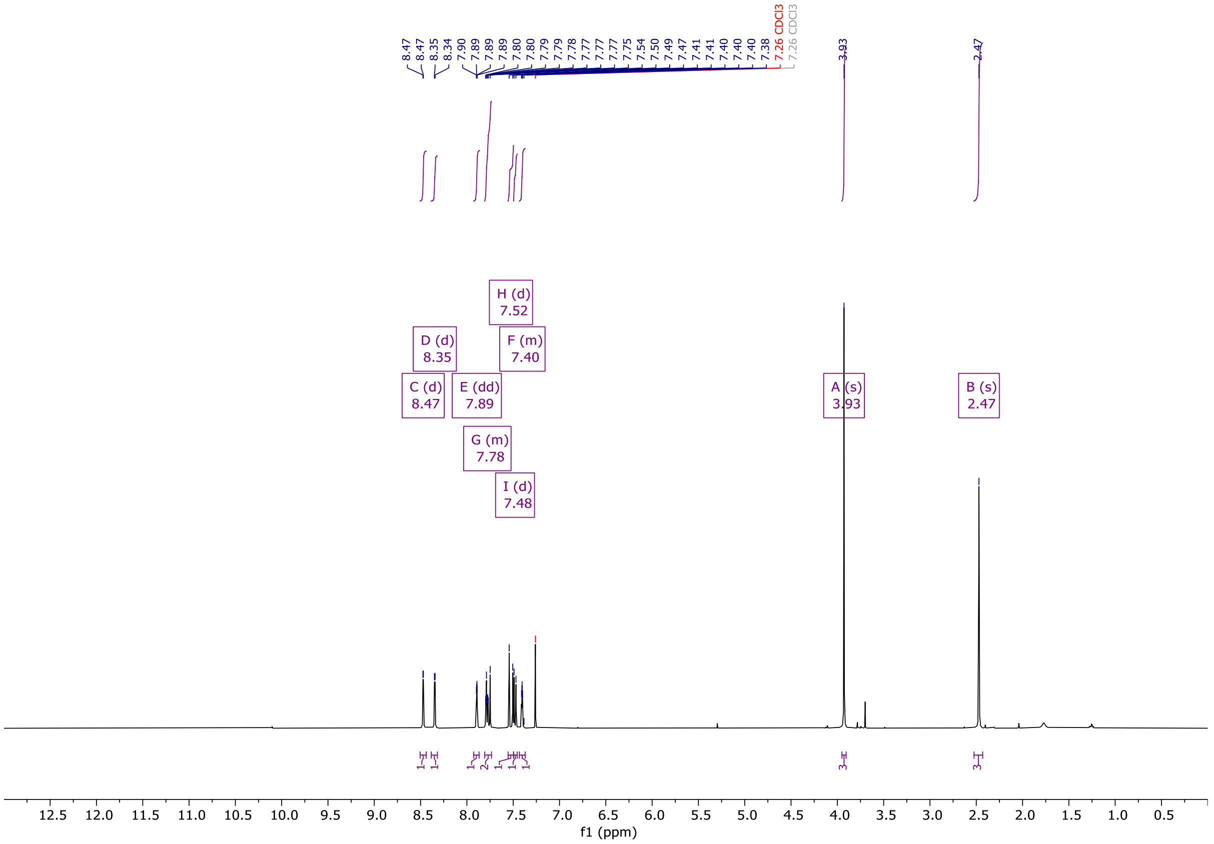


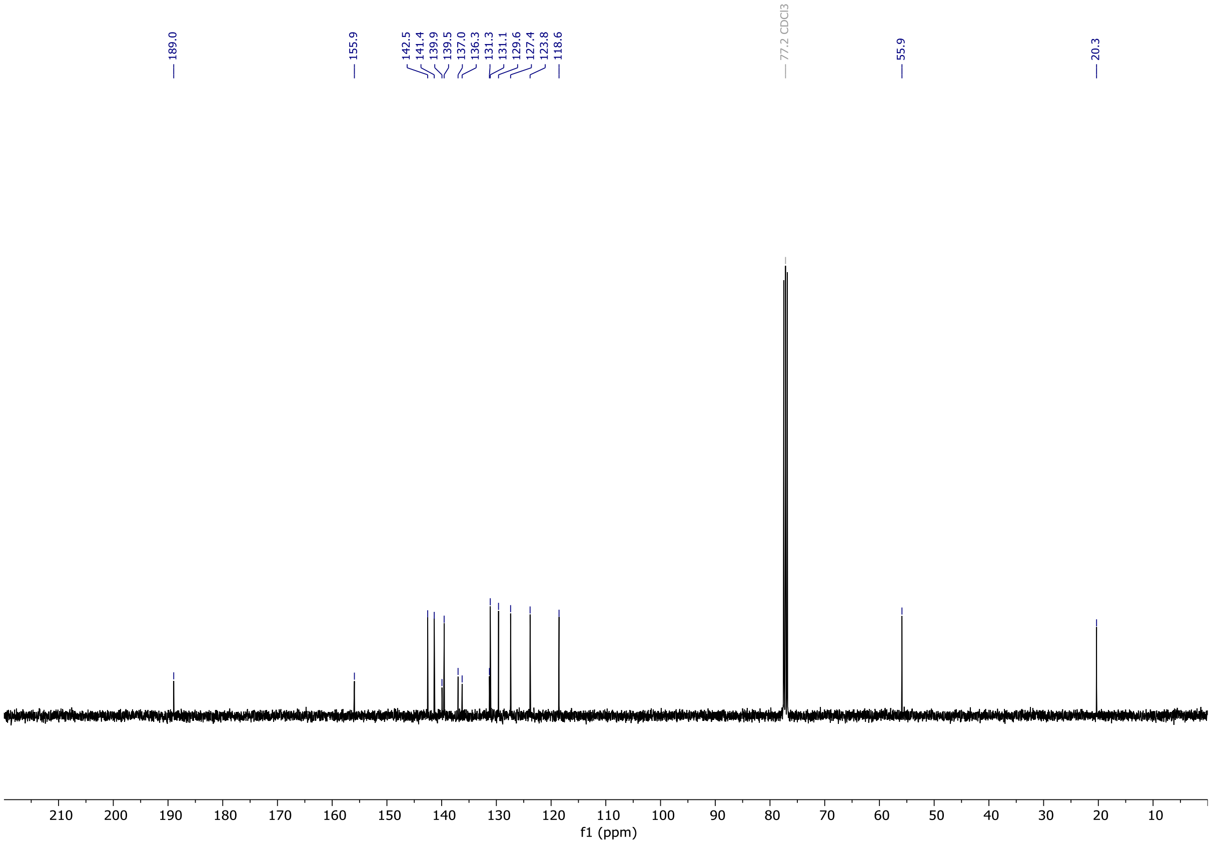

**OX04954b**

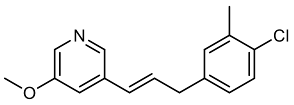

OX04954b

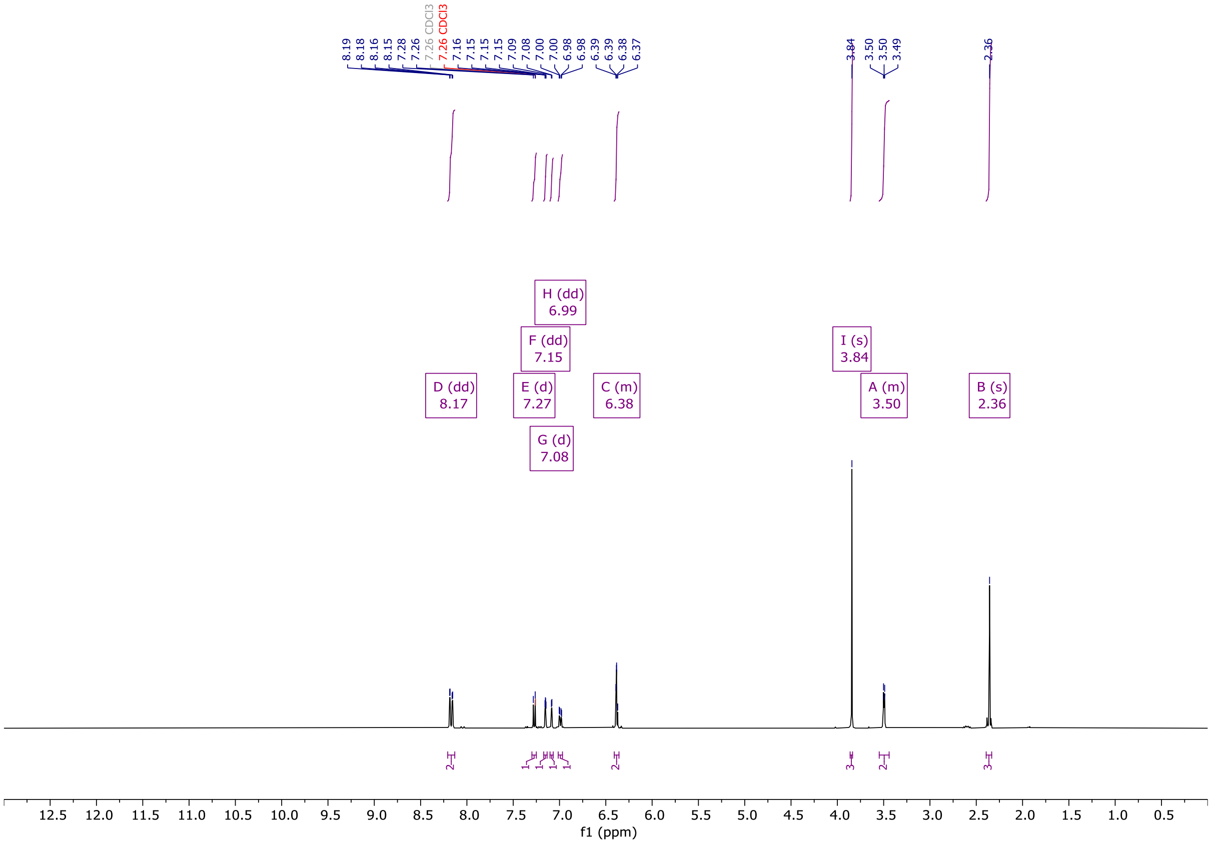


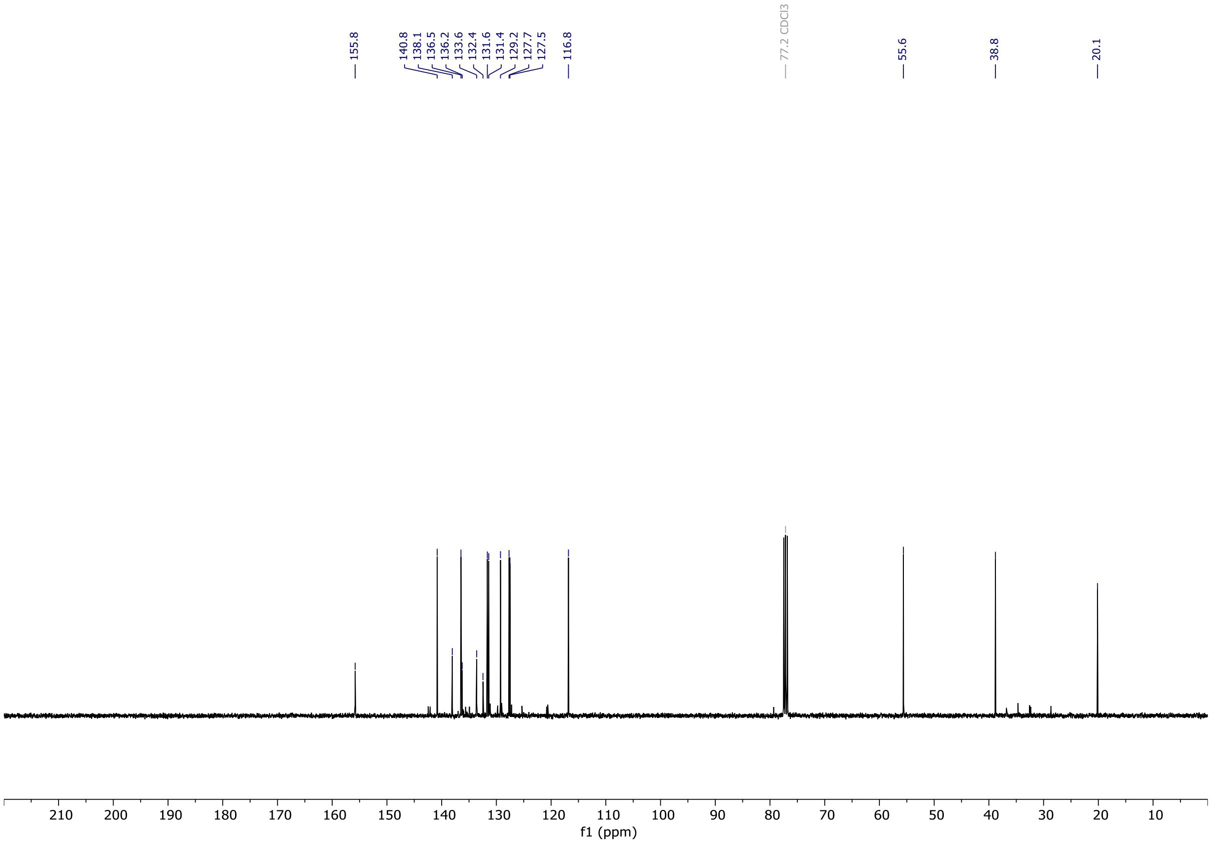

# OX04954c

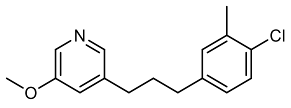


OX04954c

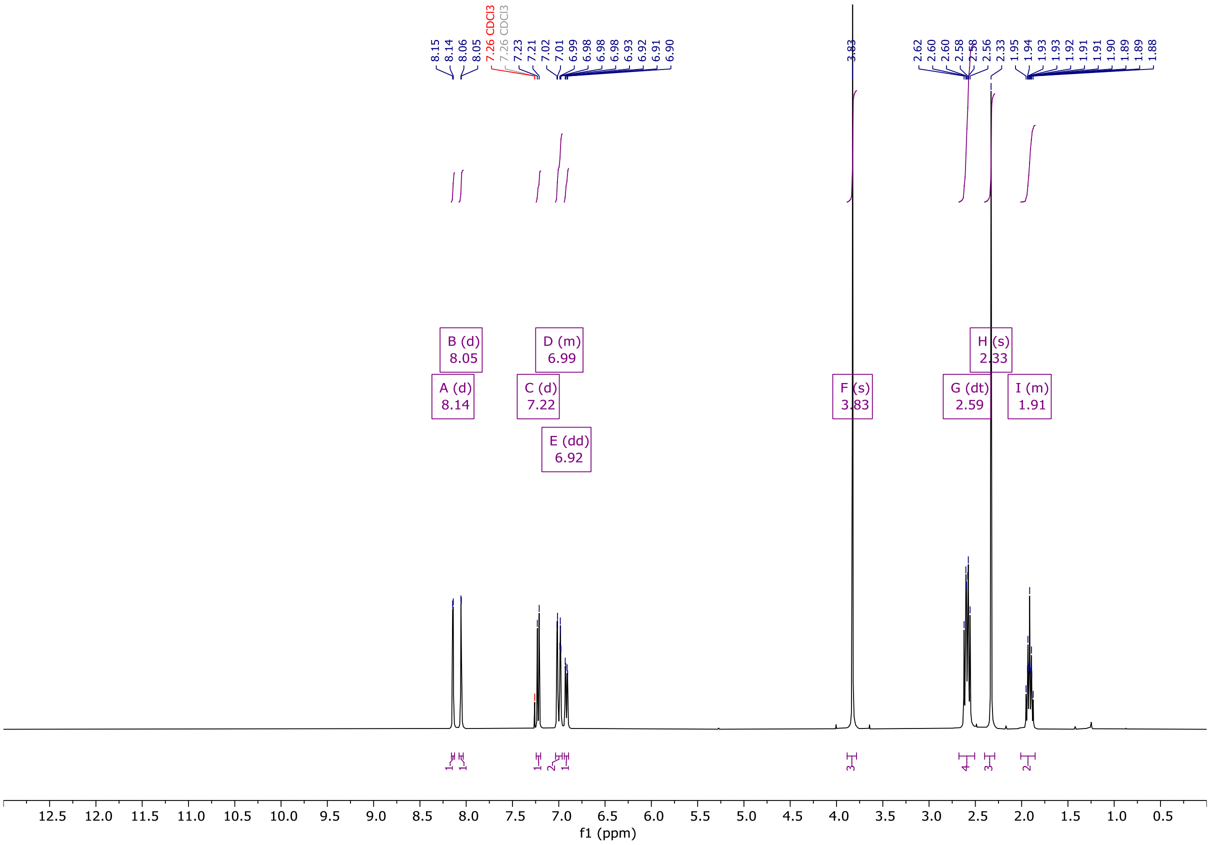


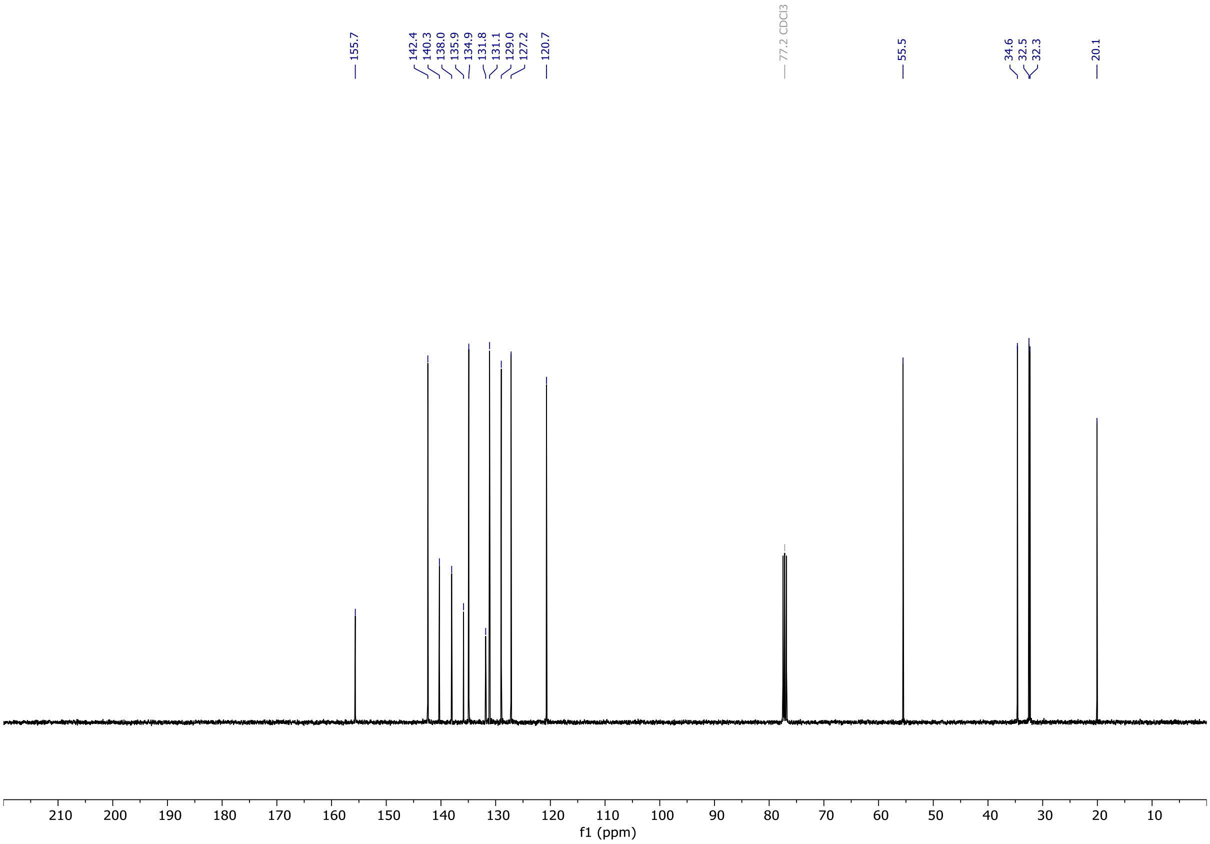

# OX04954d

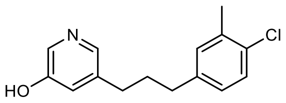

OX04954d

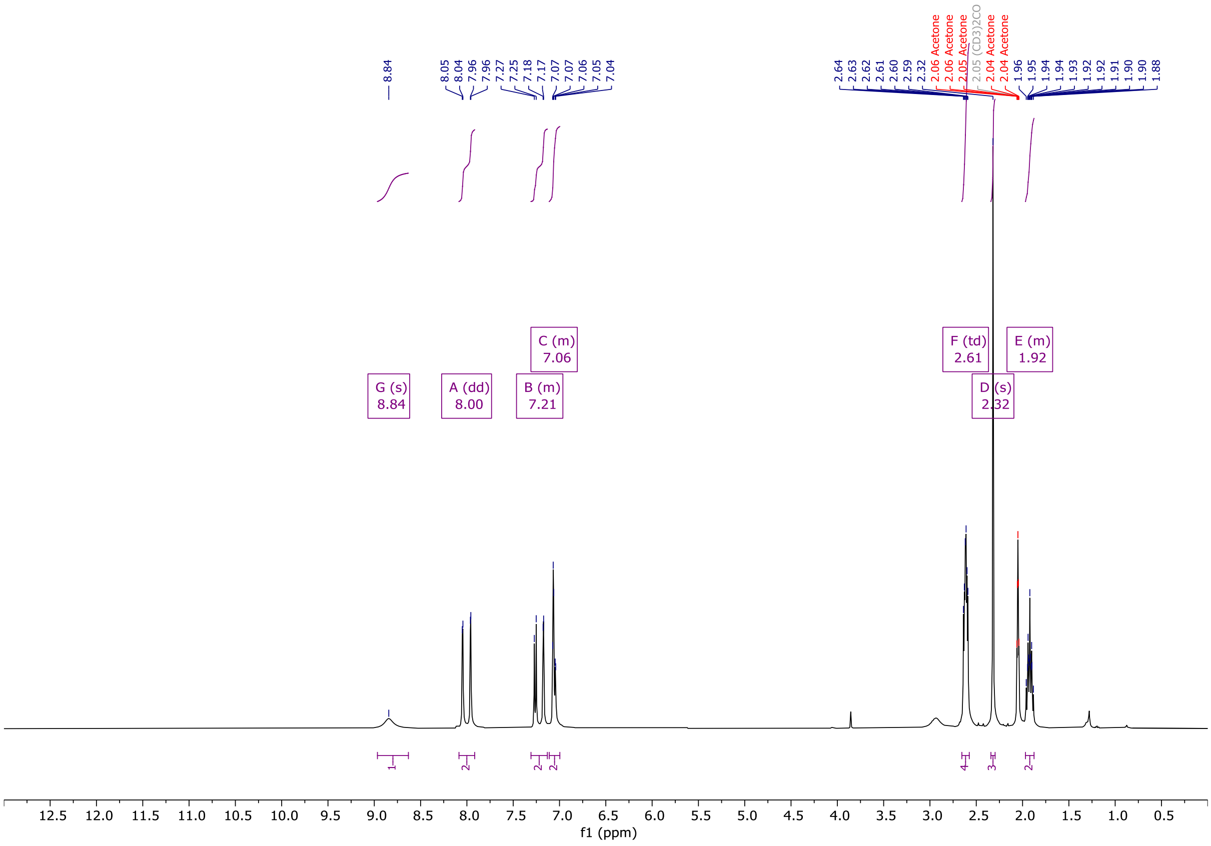


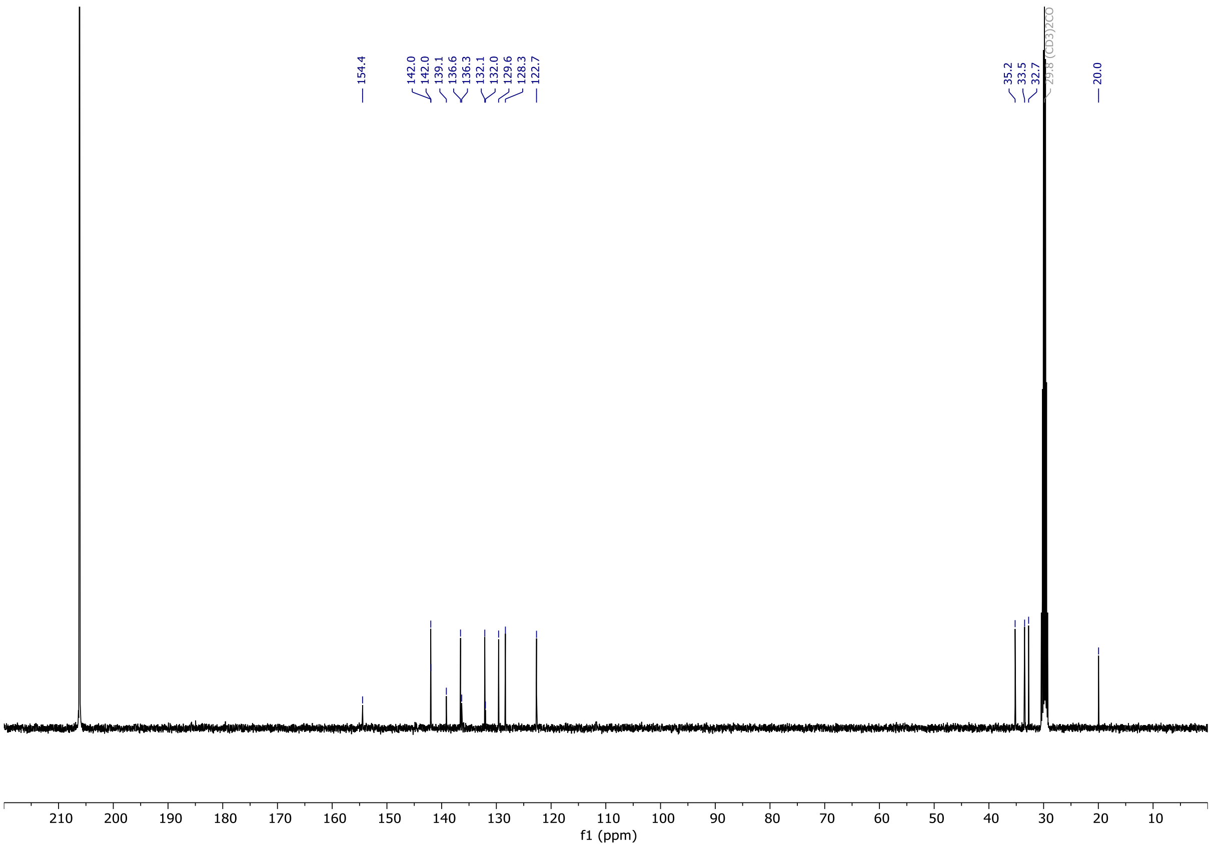

# OX04954

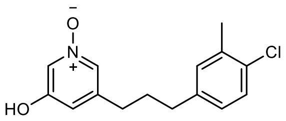


OX04954

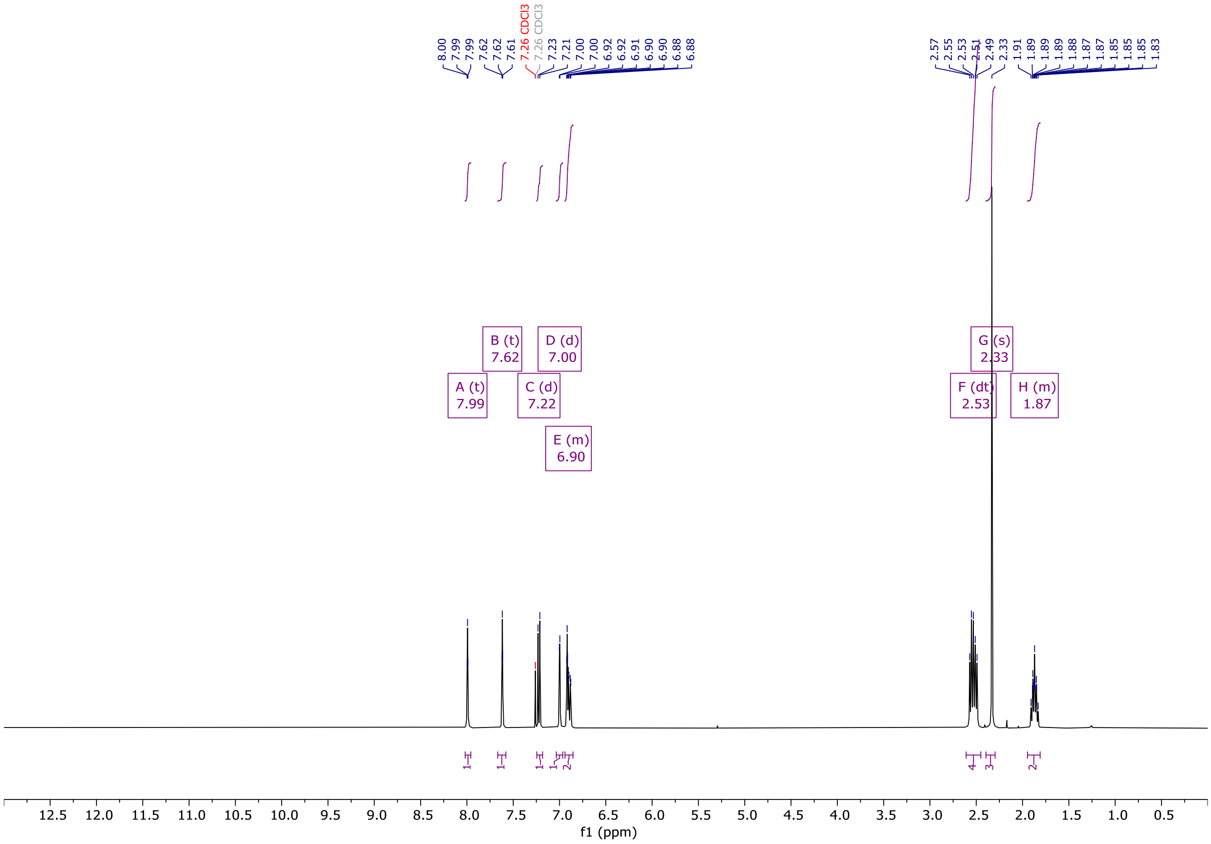


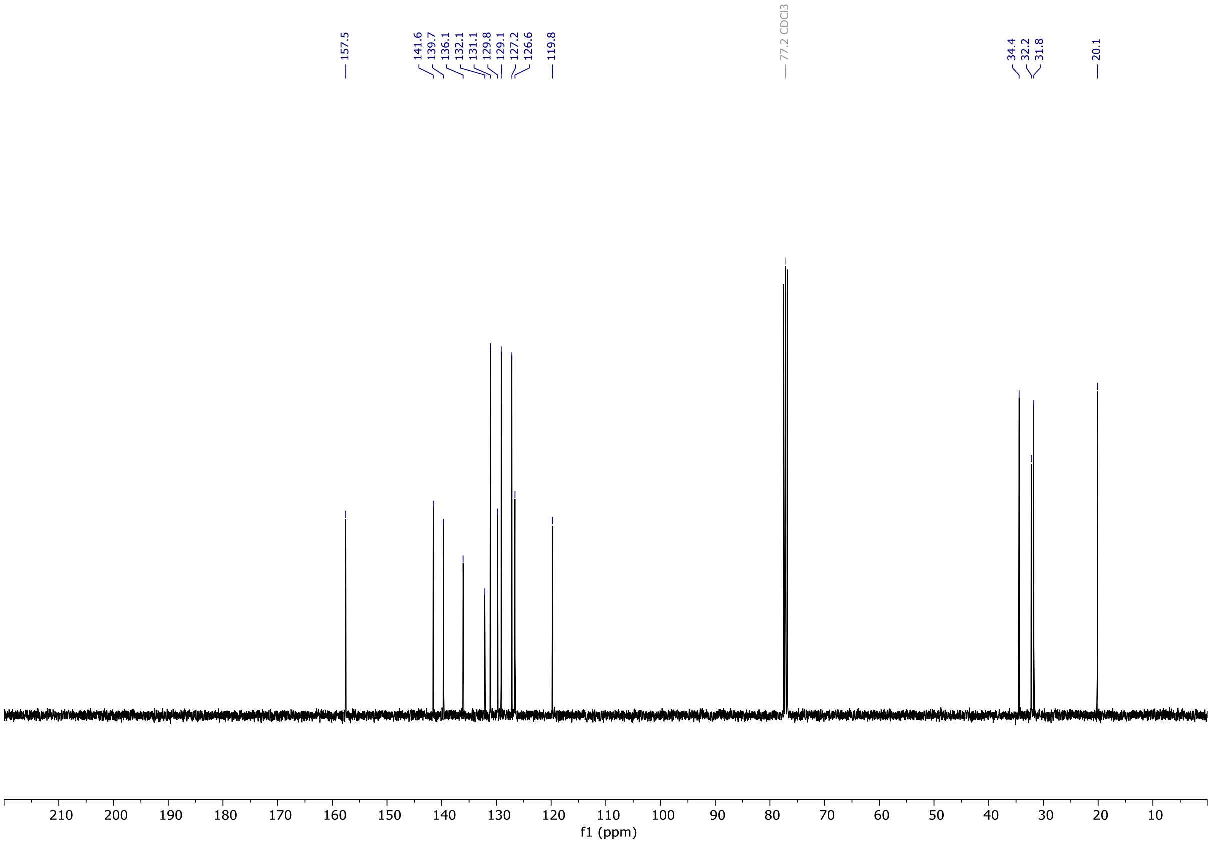

# OX04956a

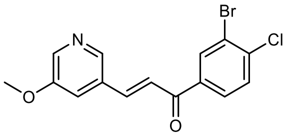

OX04956a

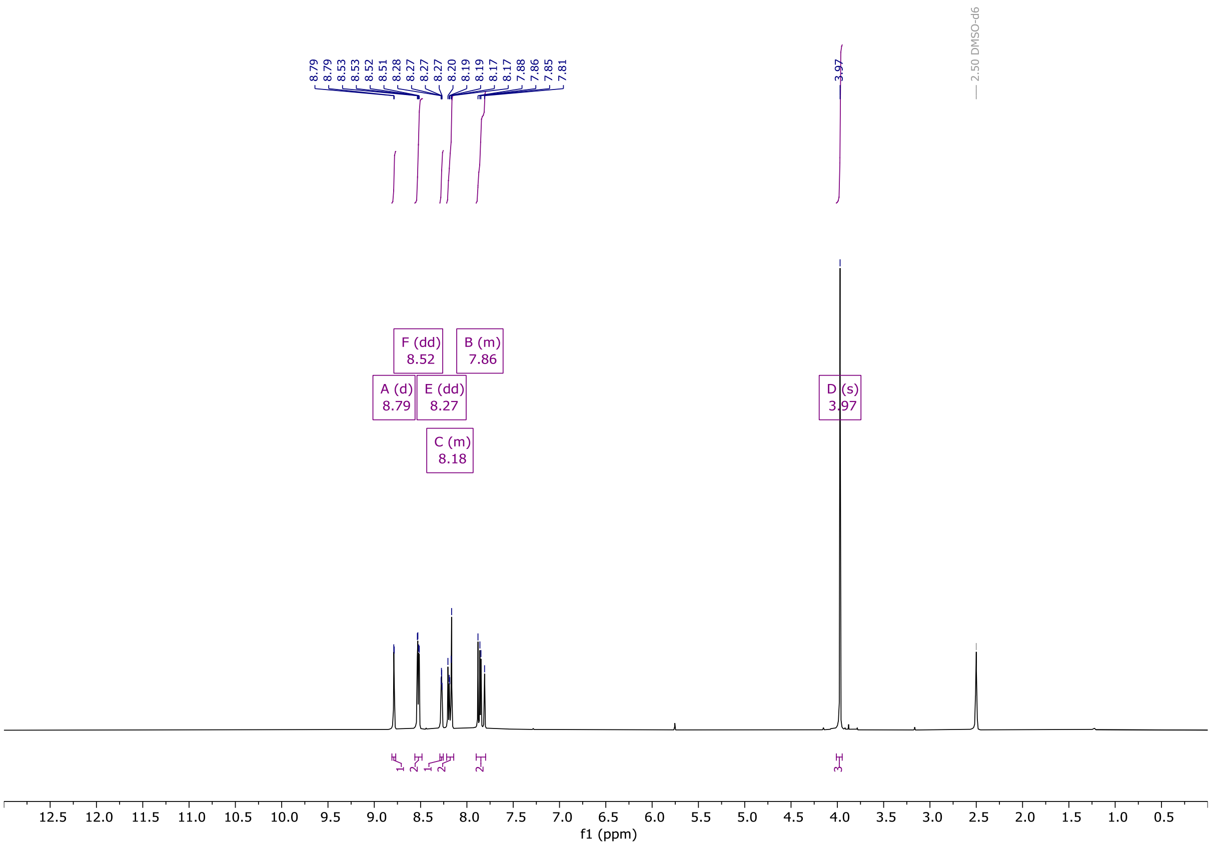


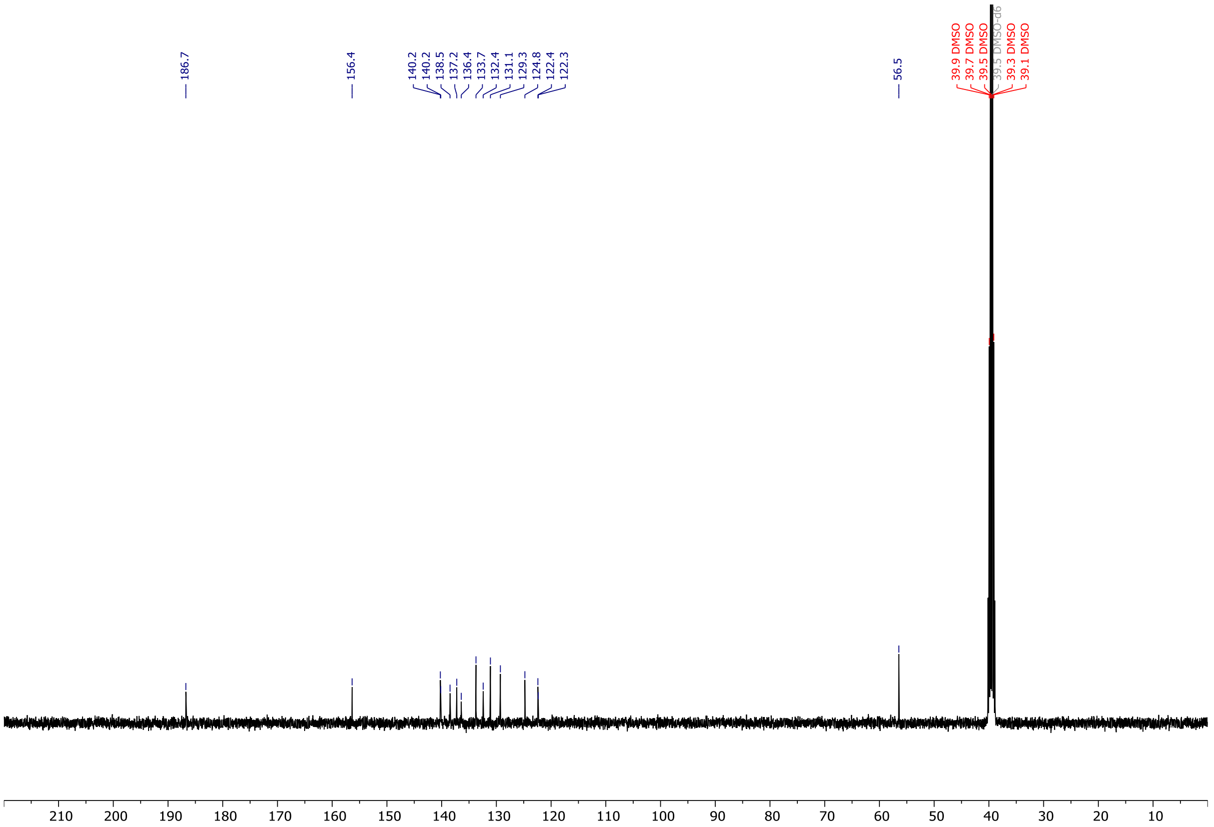

# OX04956b

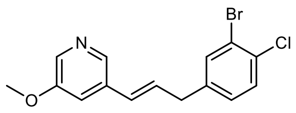

OX04956b

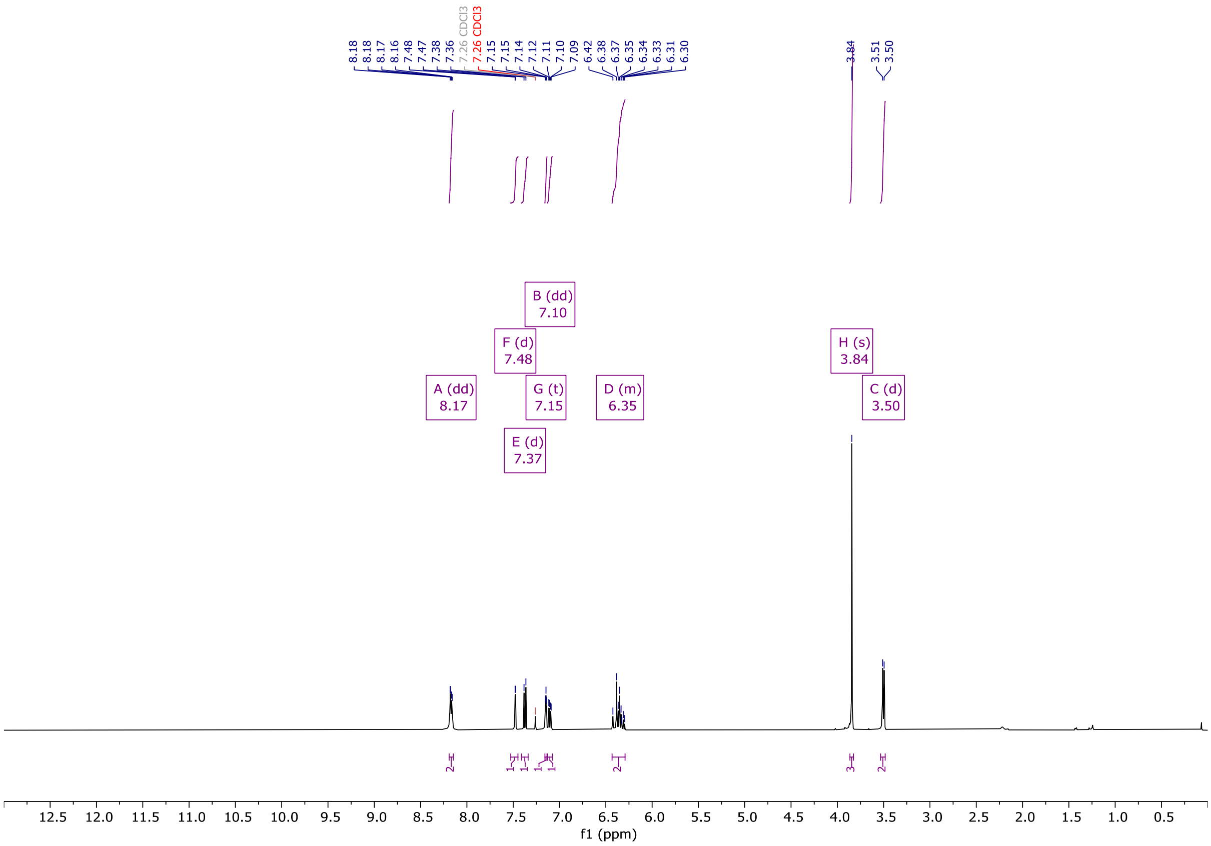


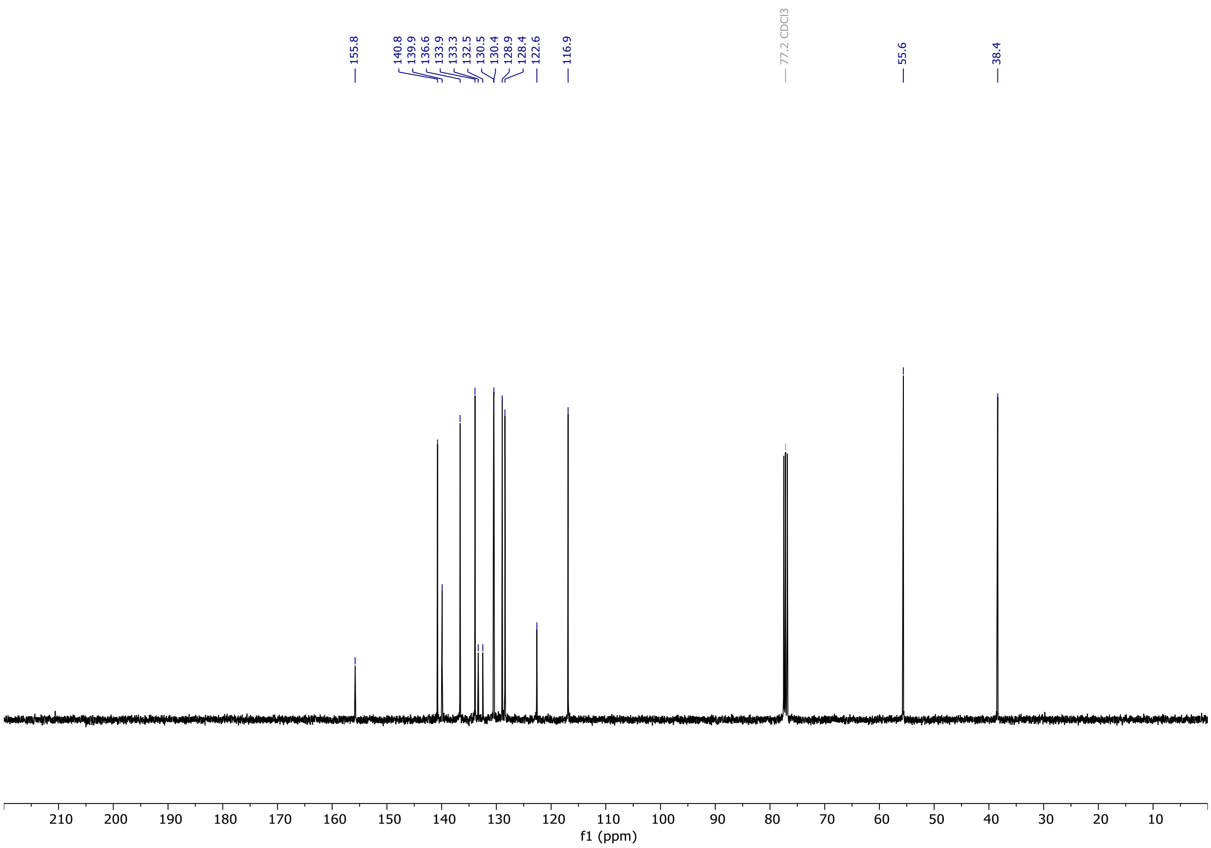

# OX04956c

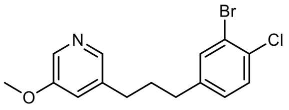


OX04956c

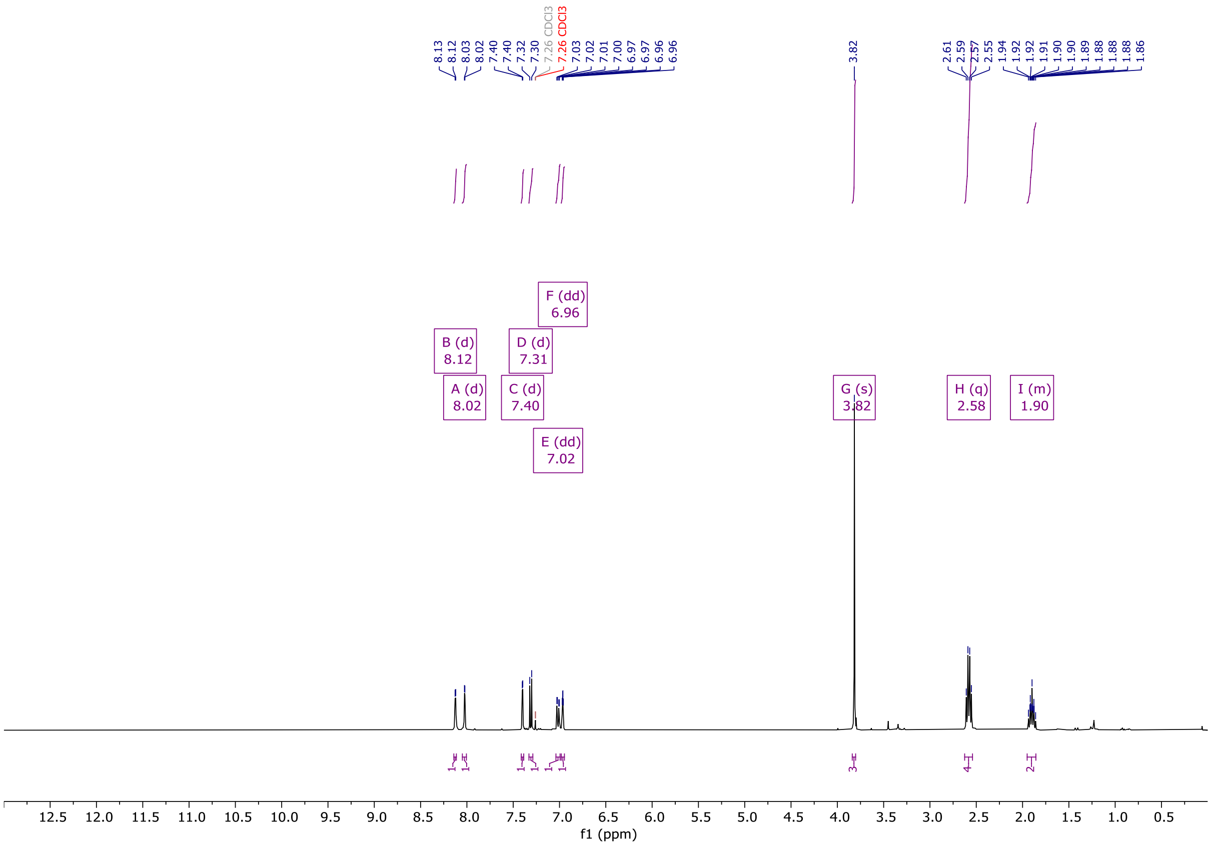


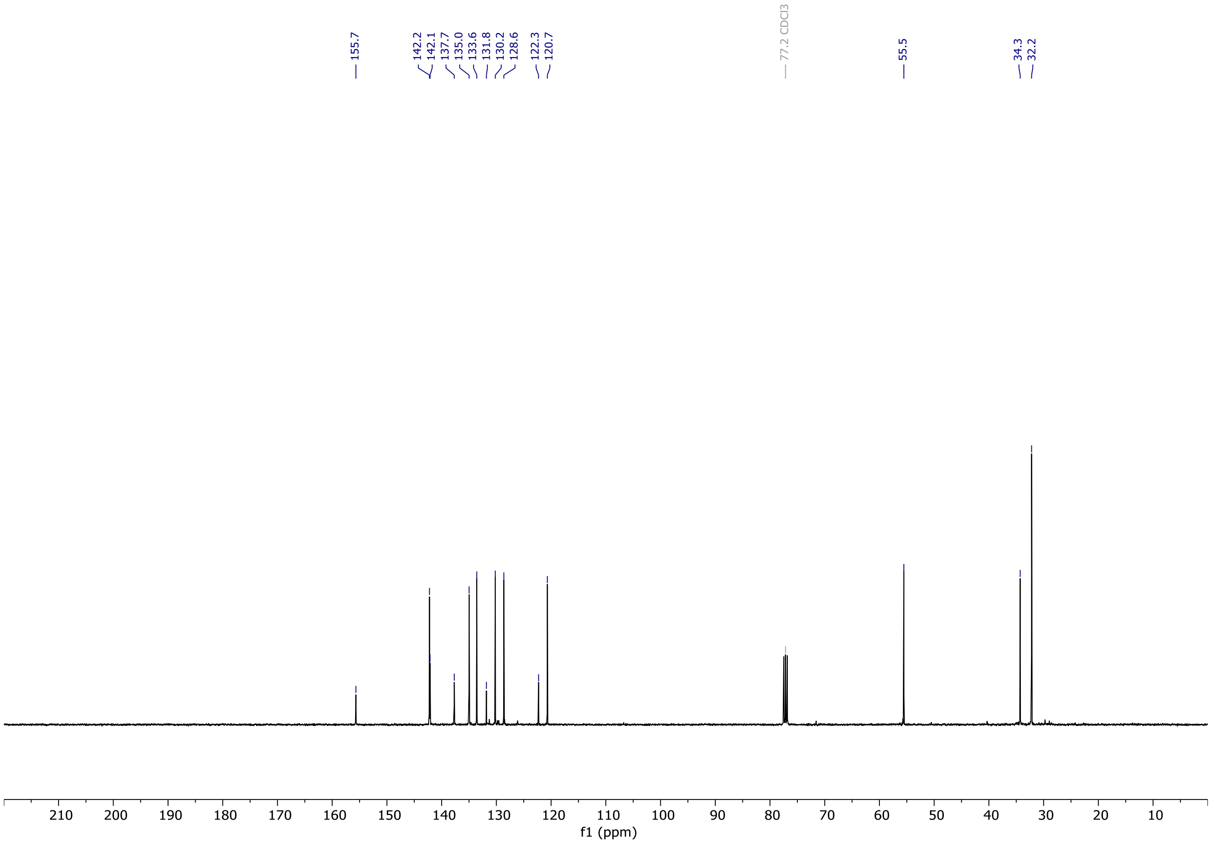

# OX04956d

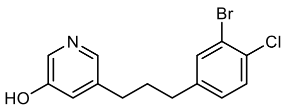


OX04956d

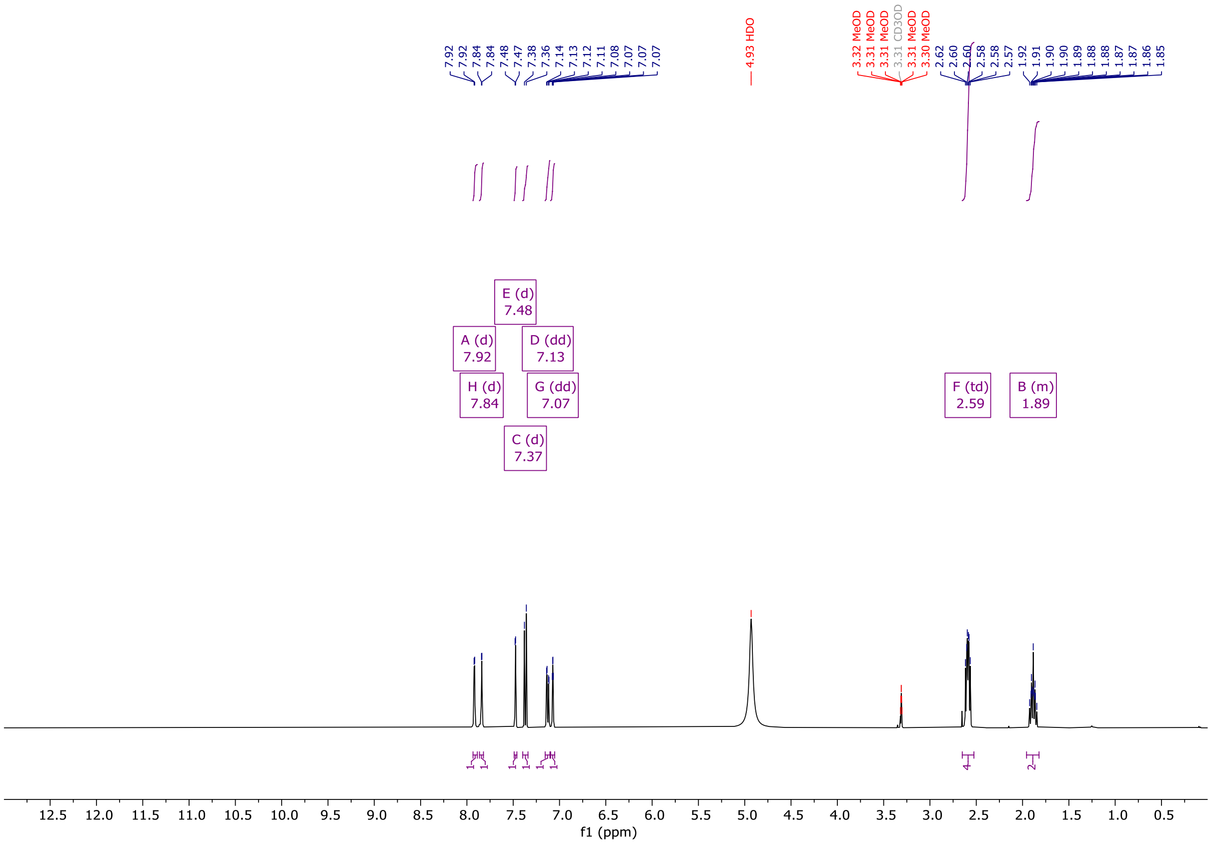


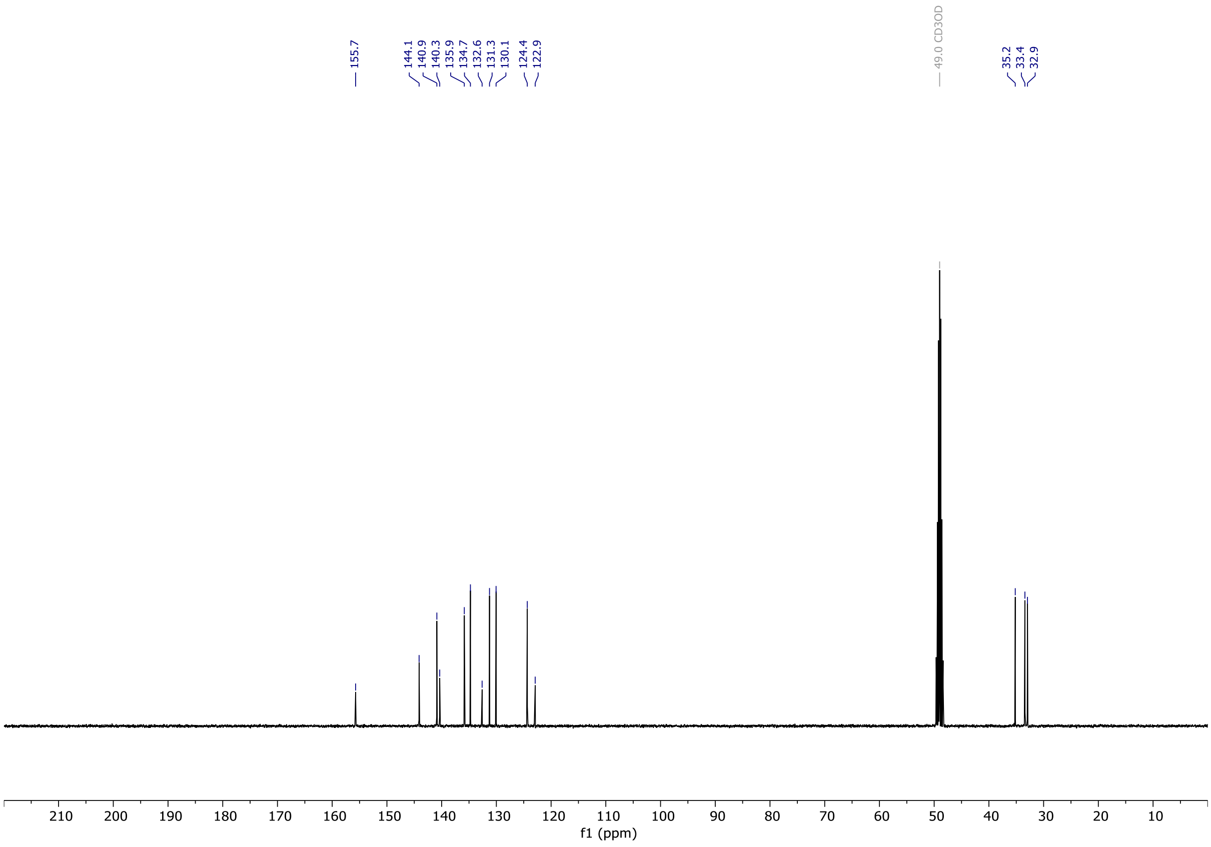

# OX04956

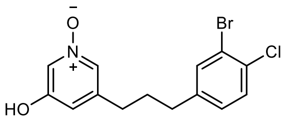


OX04956

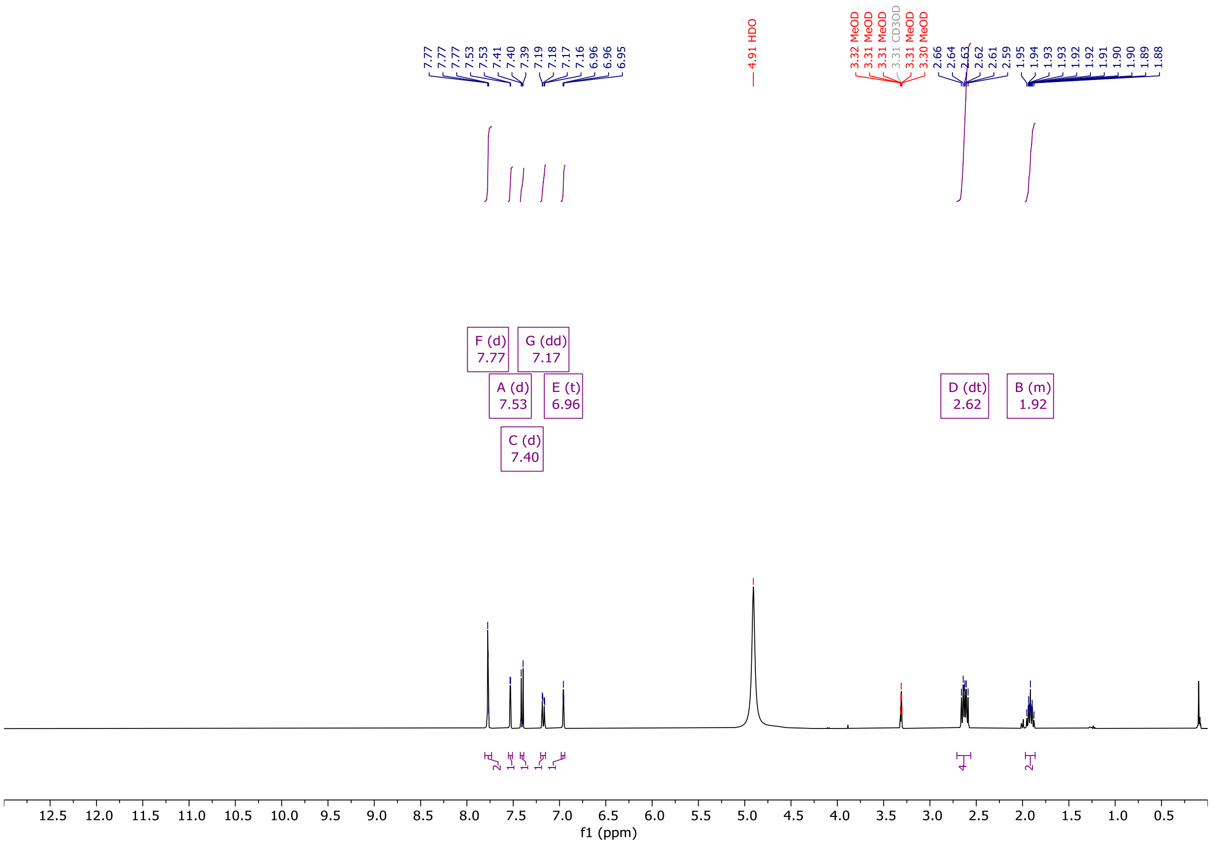


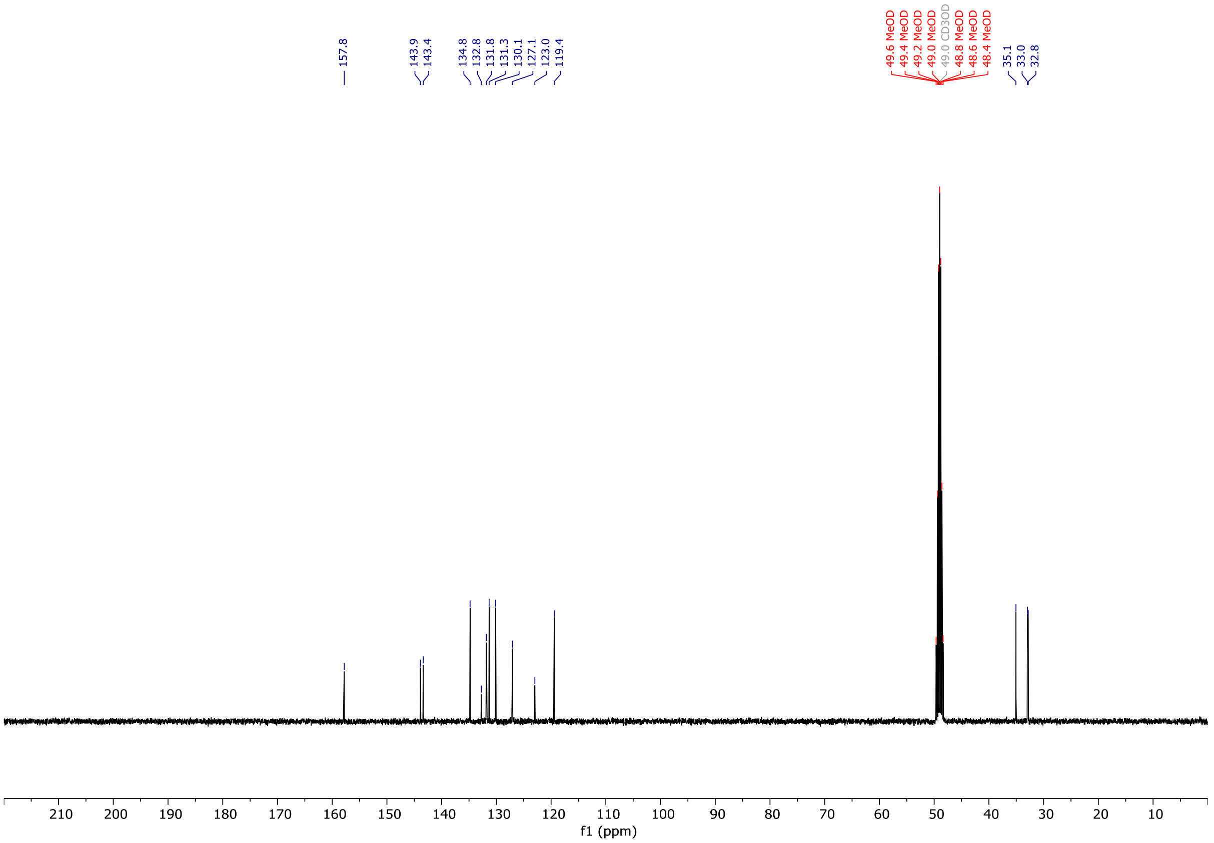

# OX04955a

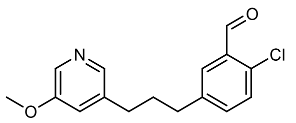


OX04955a

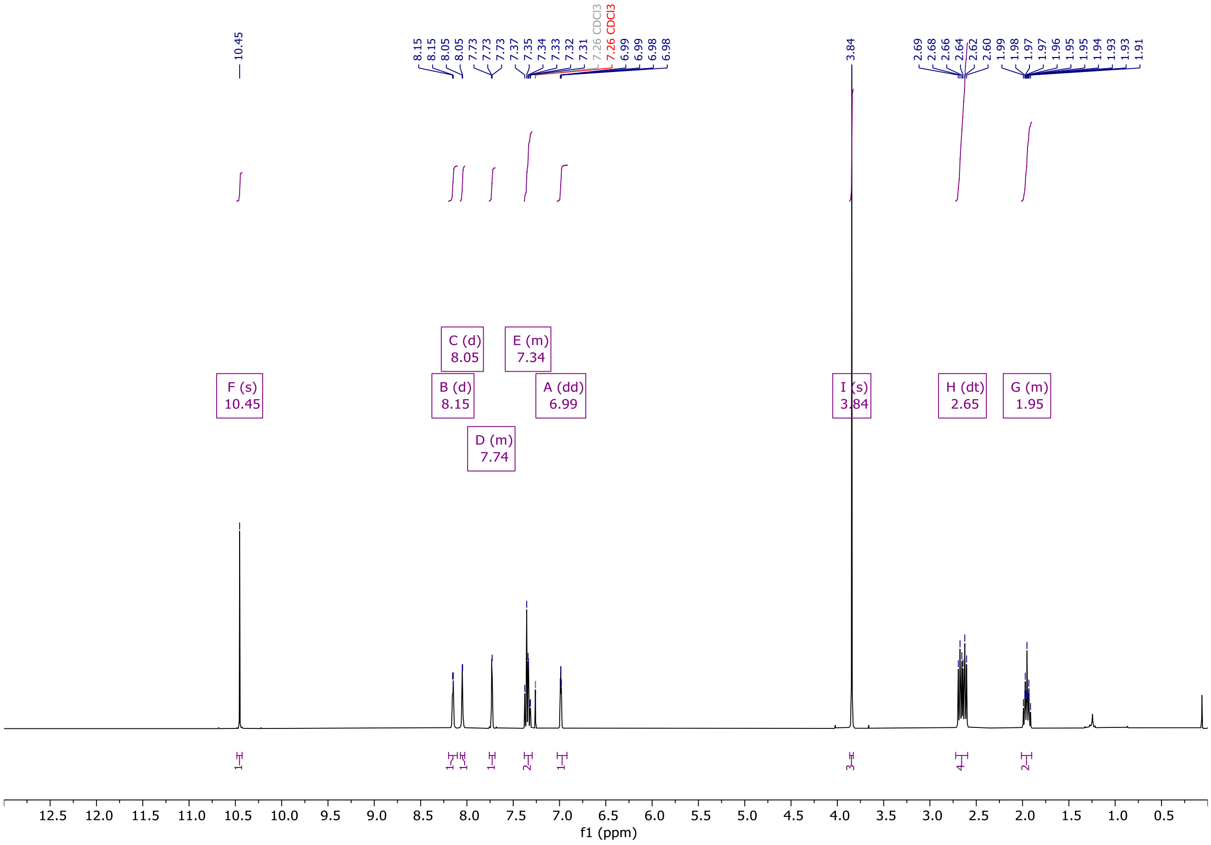


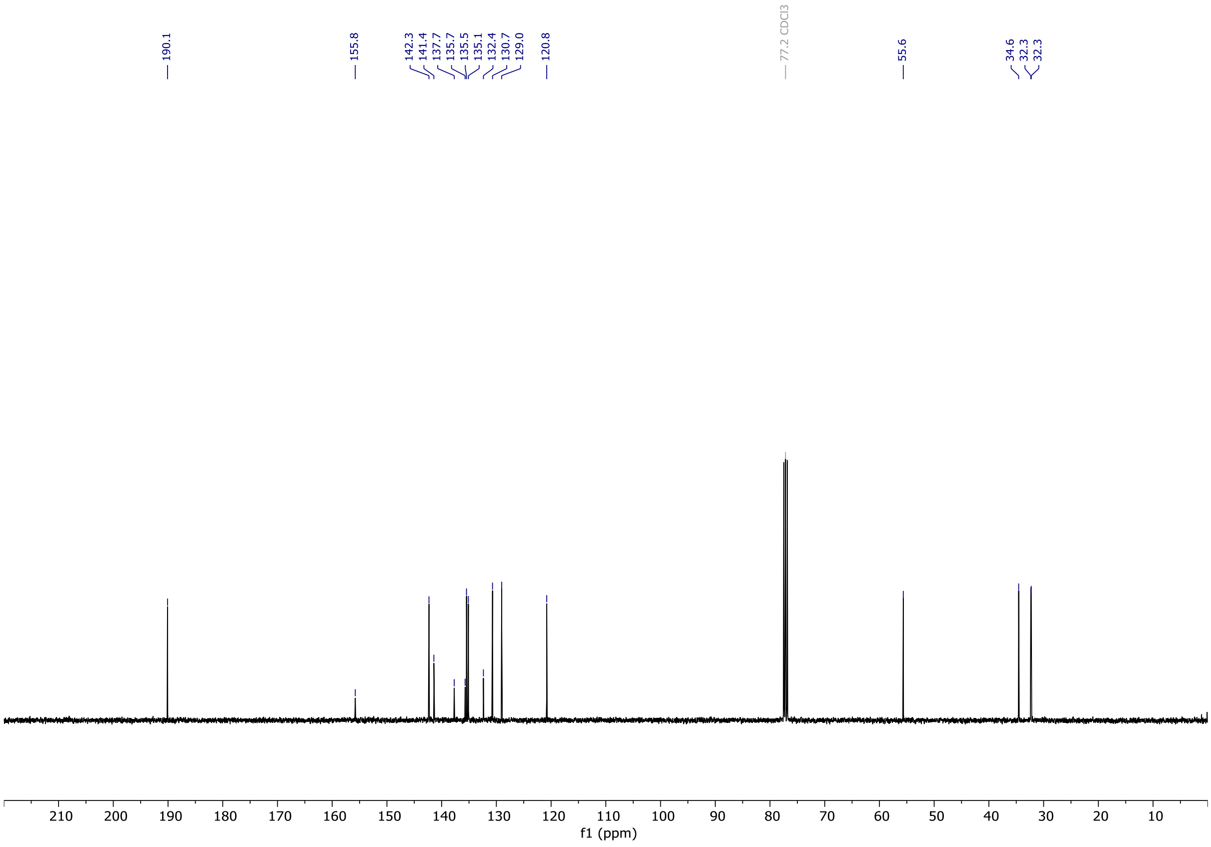

# OX04955b

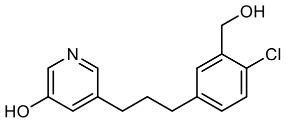


OX04955b

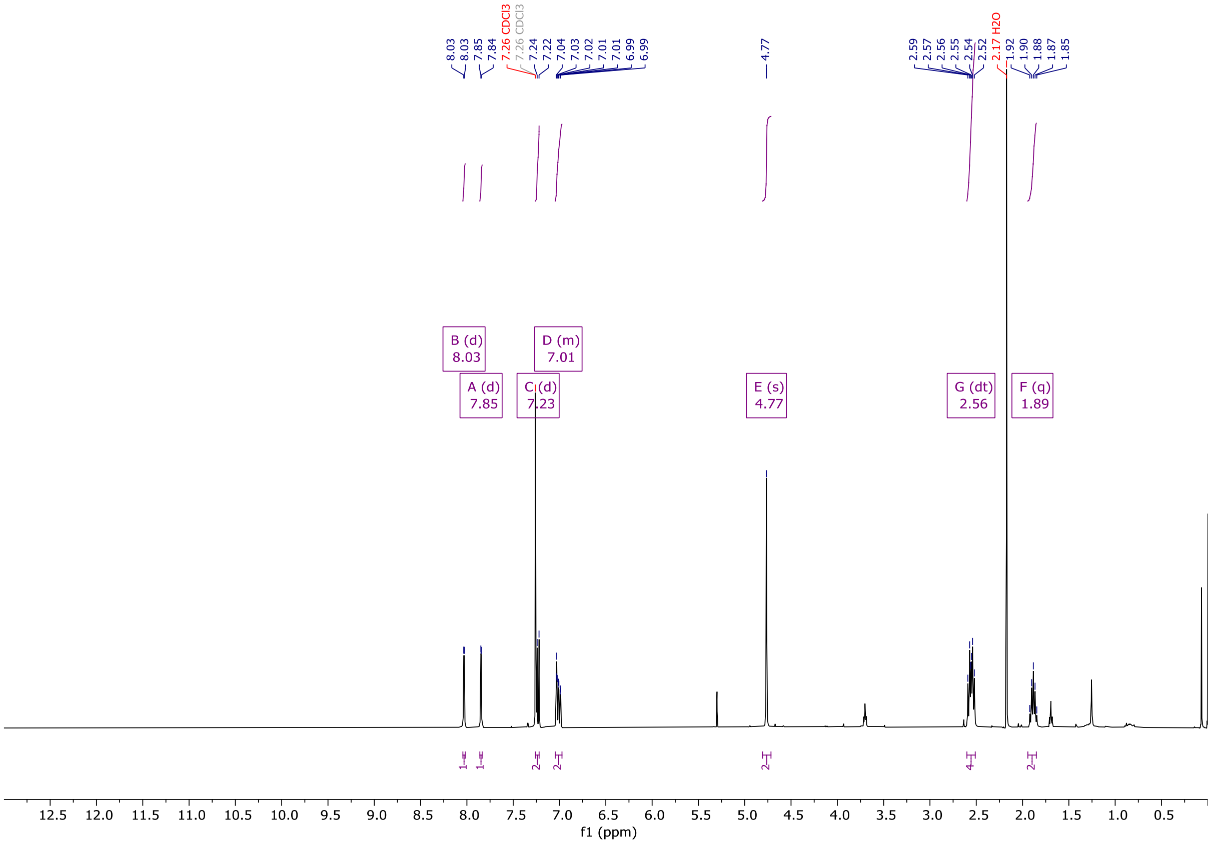


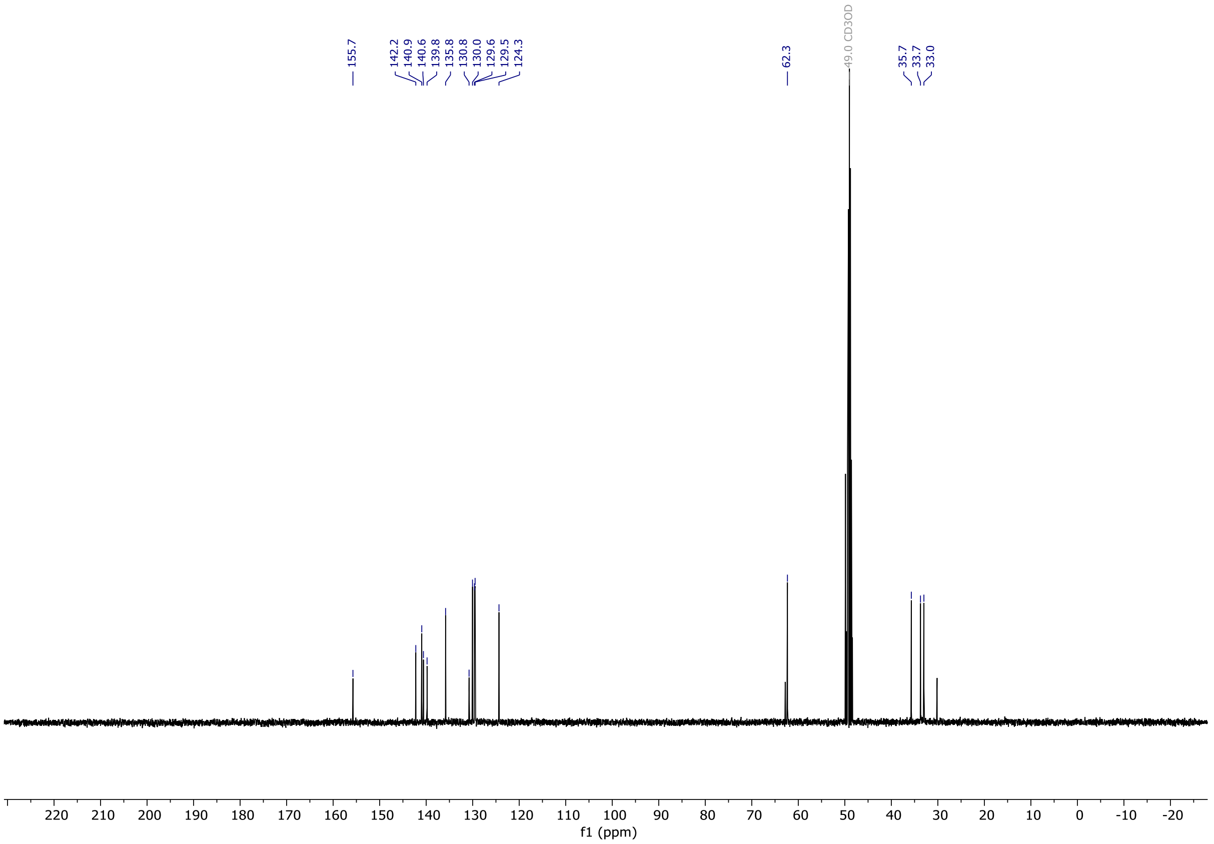

# OX04955

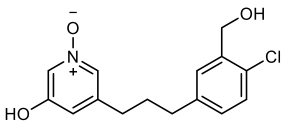

OX04955

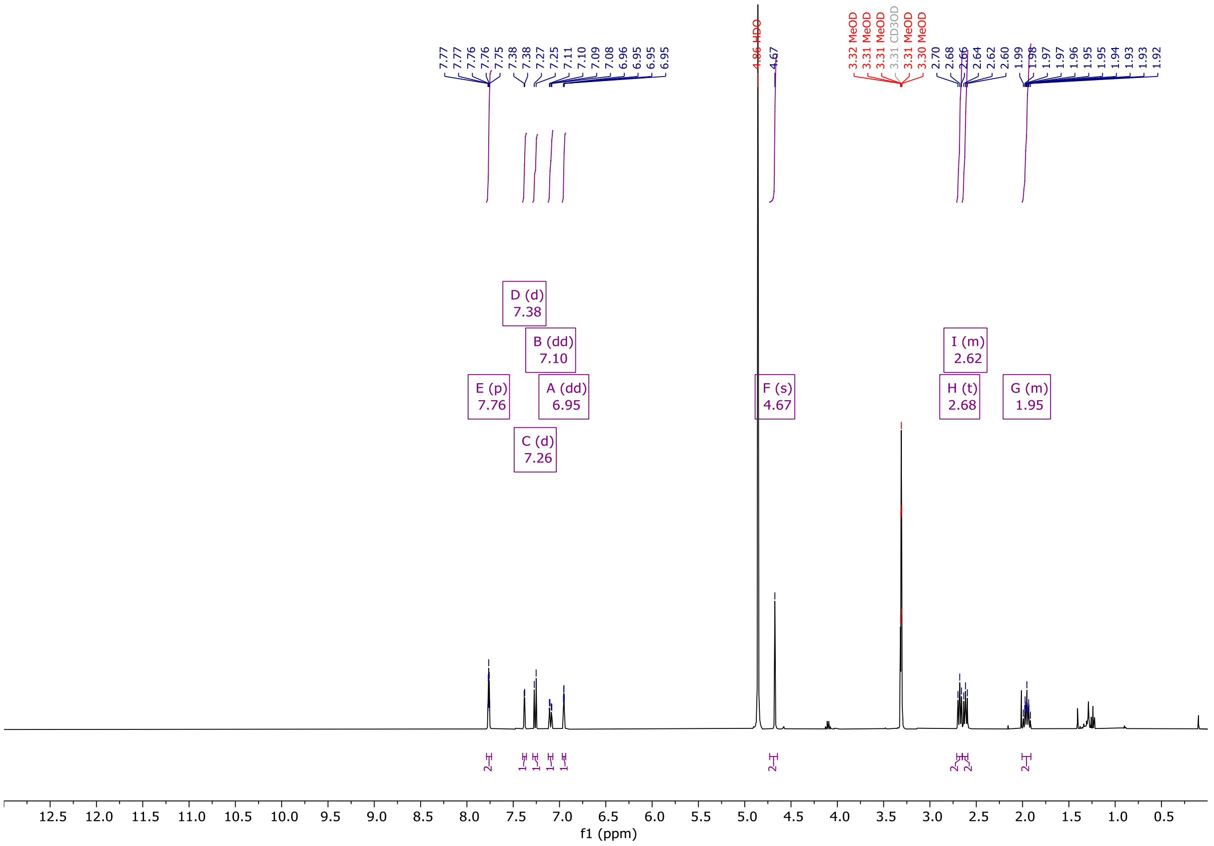


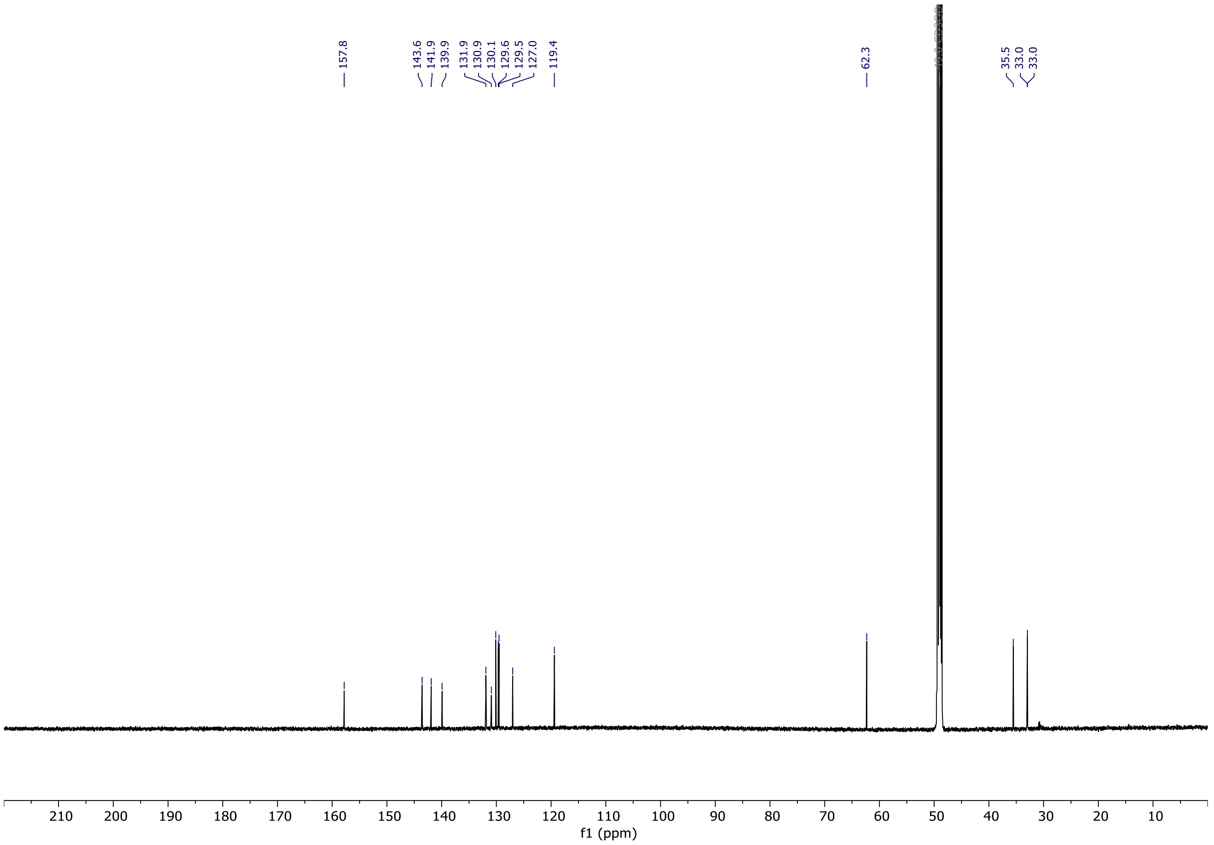

# OX04957a

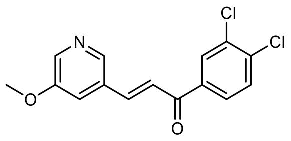


OX04957a

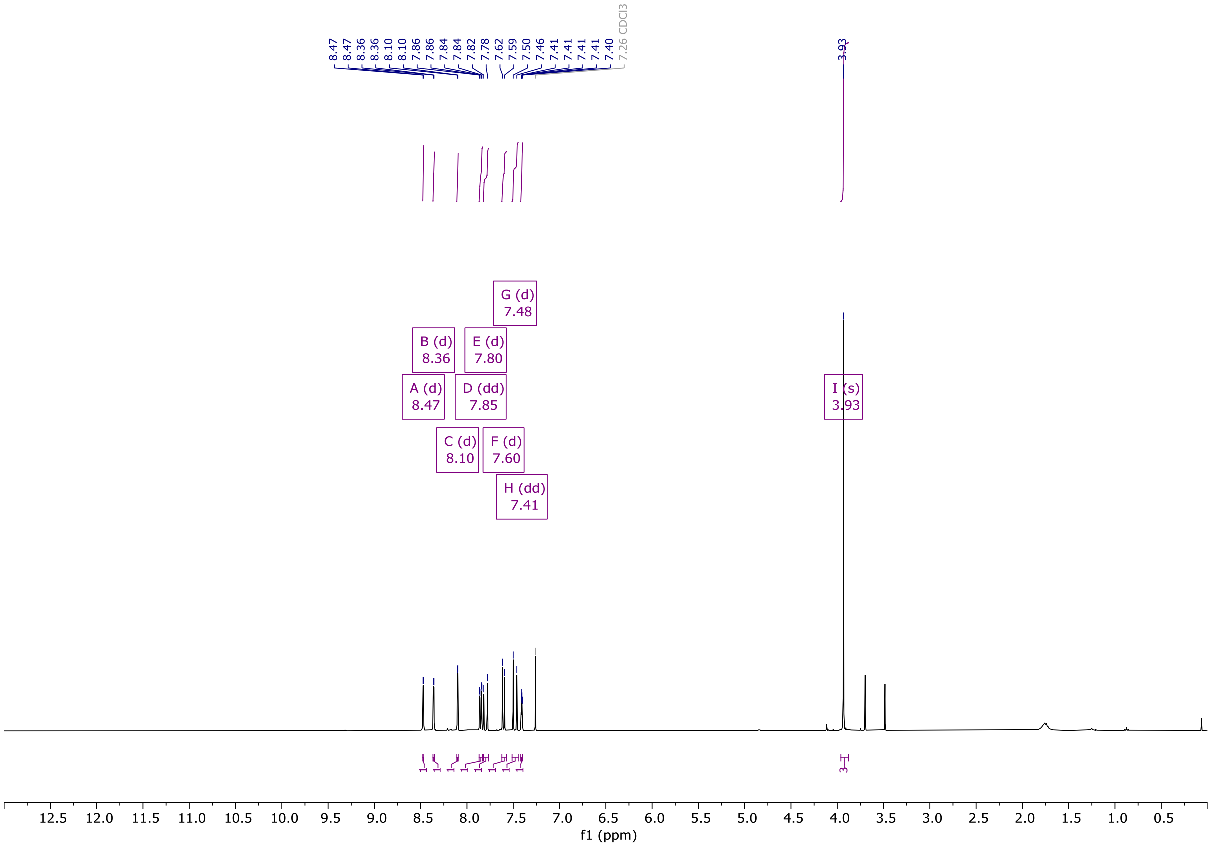

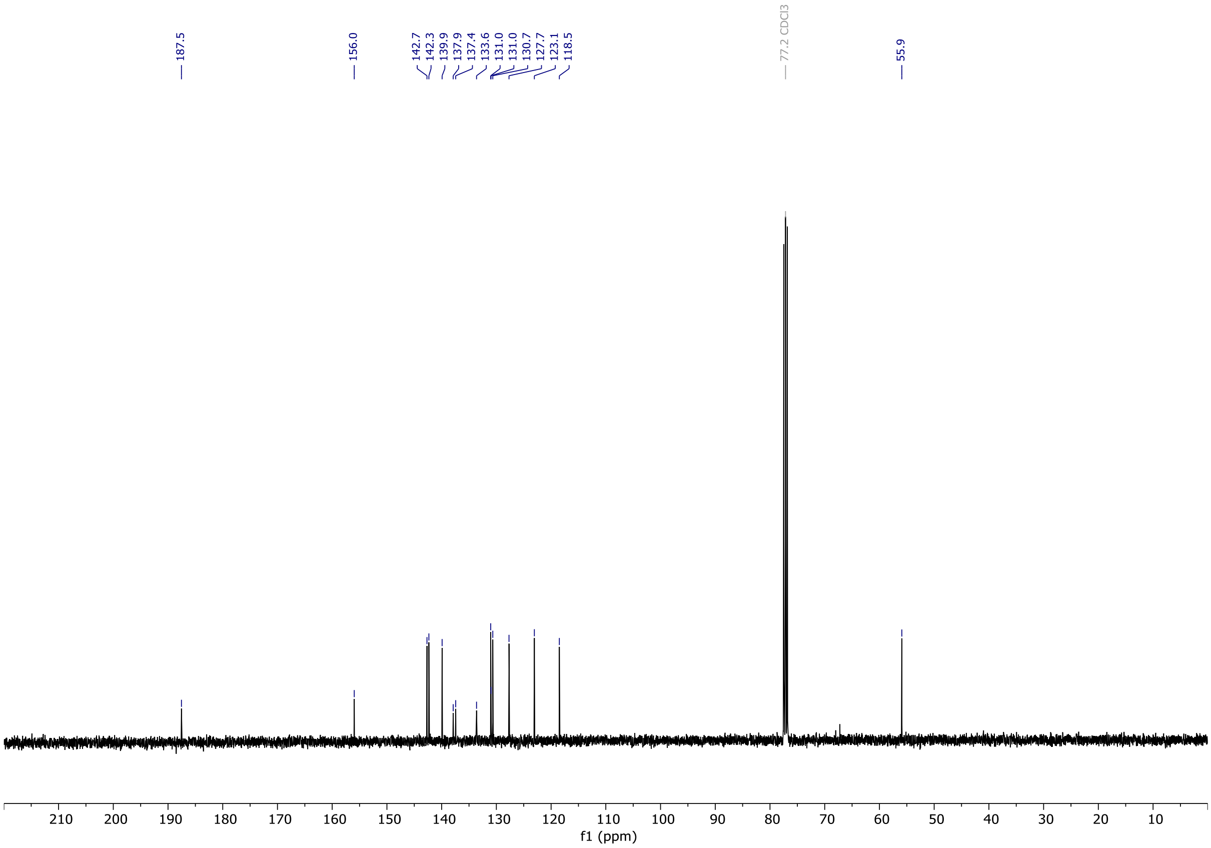

# OX04957b

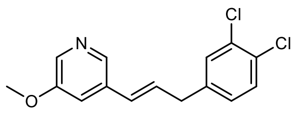


OX04957b

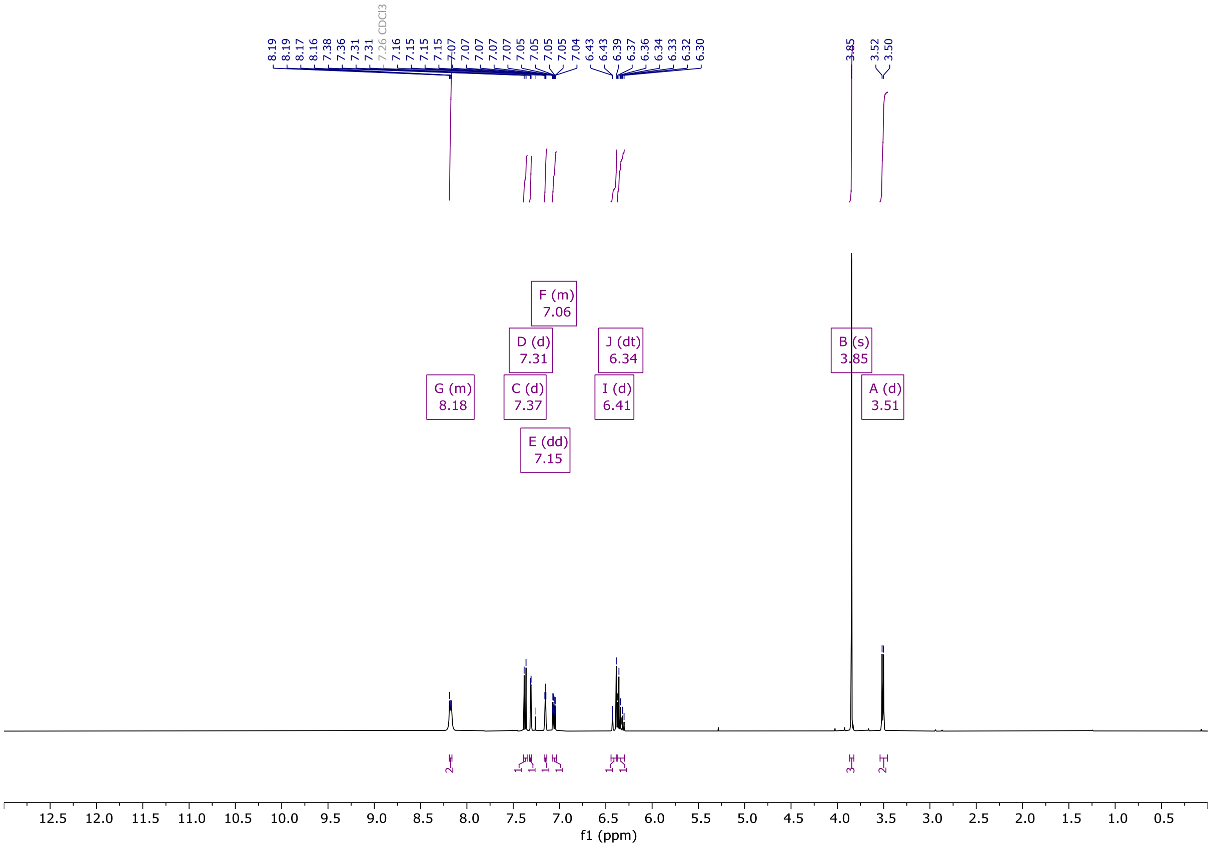


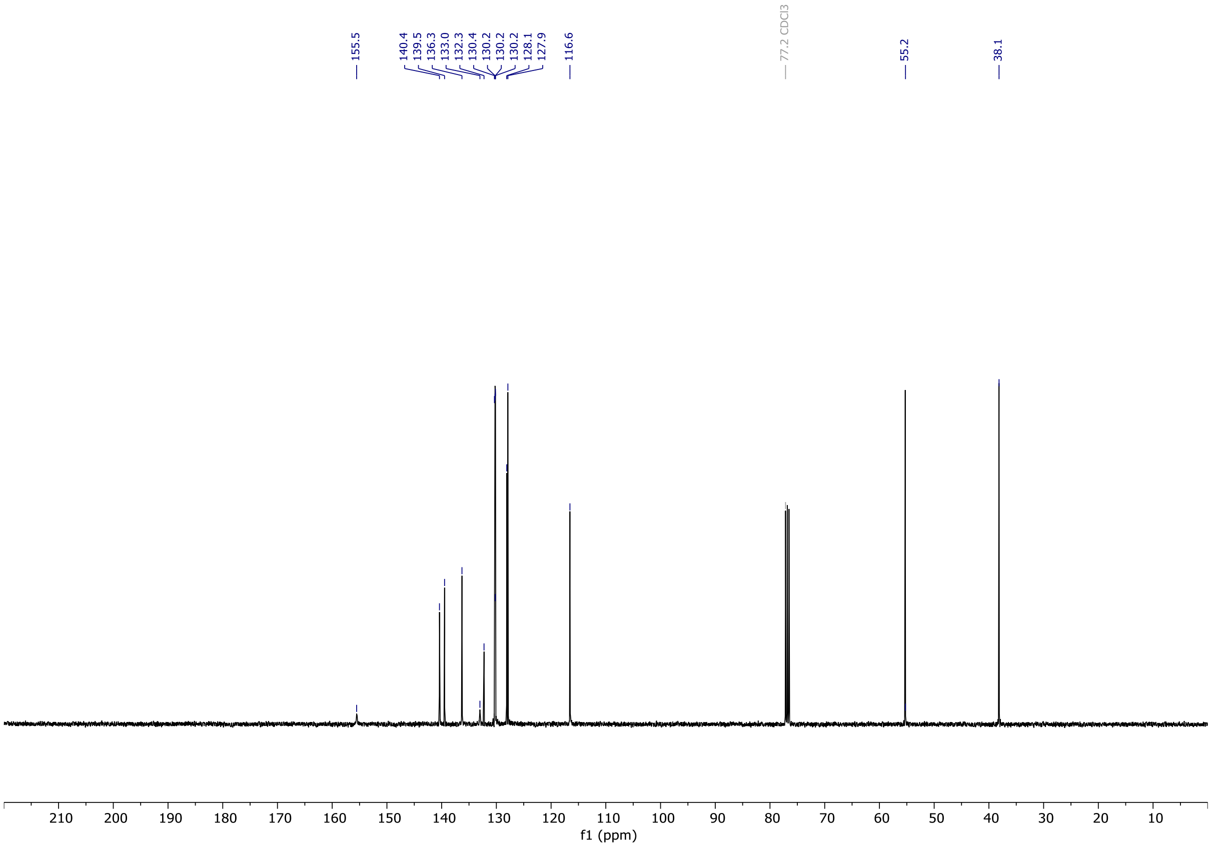

# OX04957c

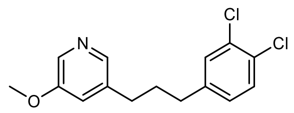


OX04957c

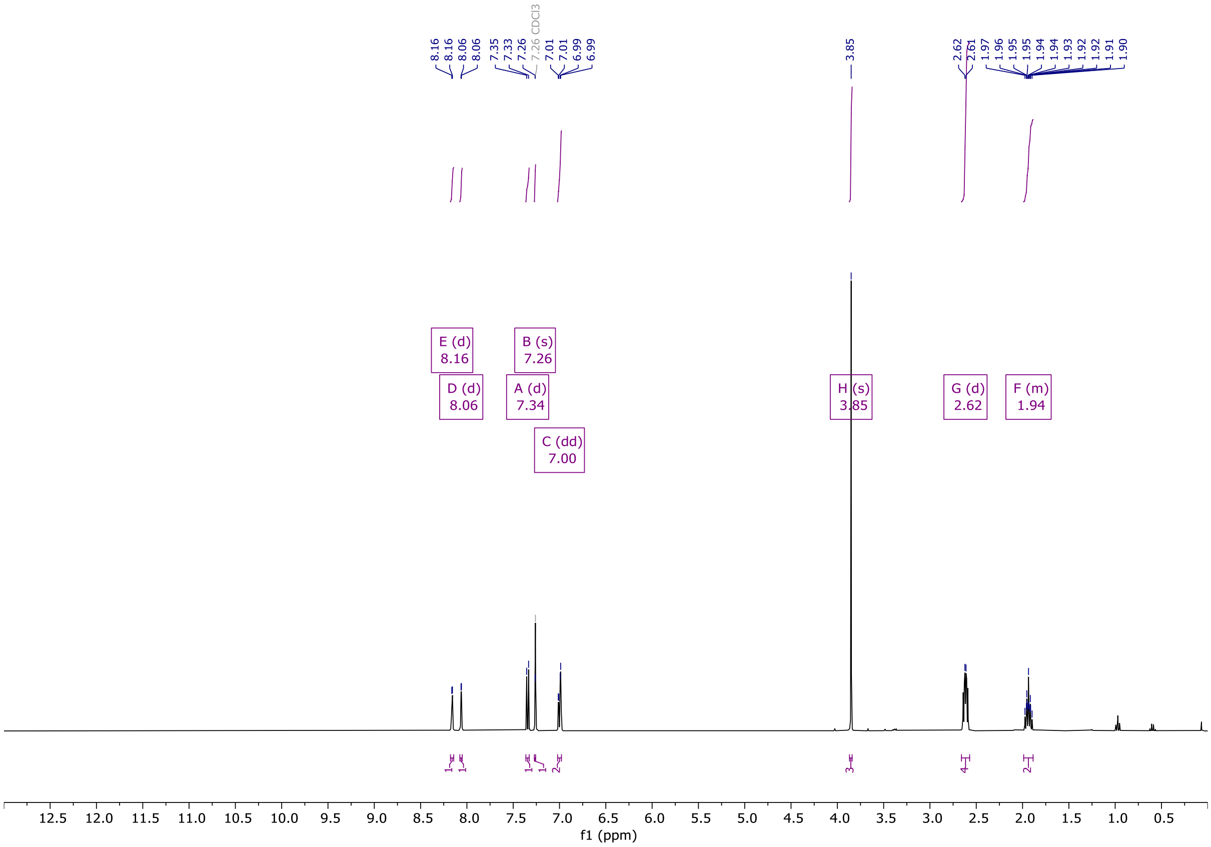


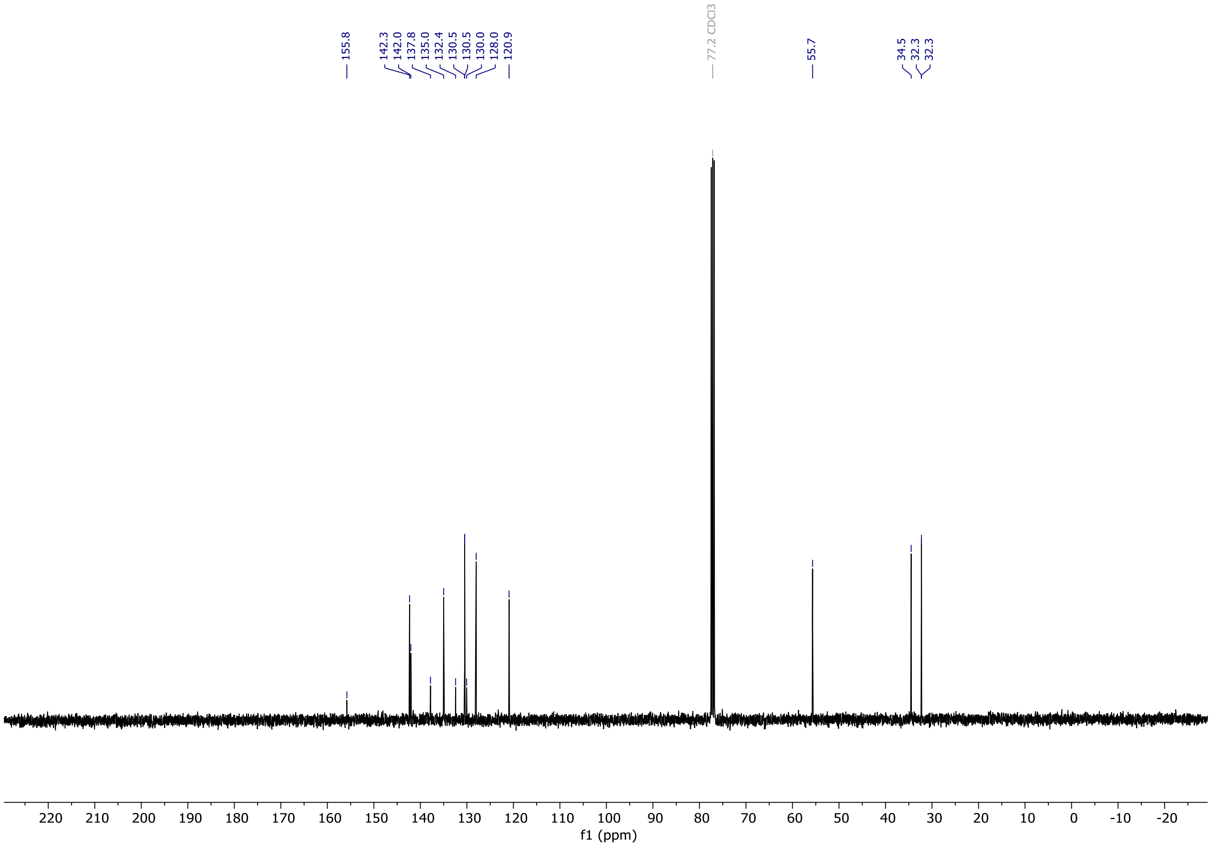

# OX04957d

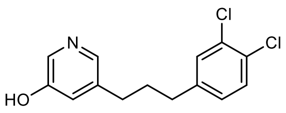

OX04957d

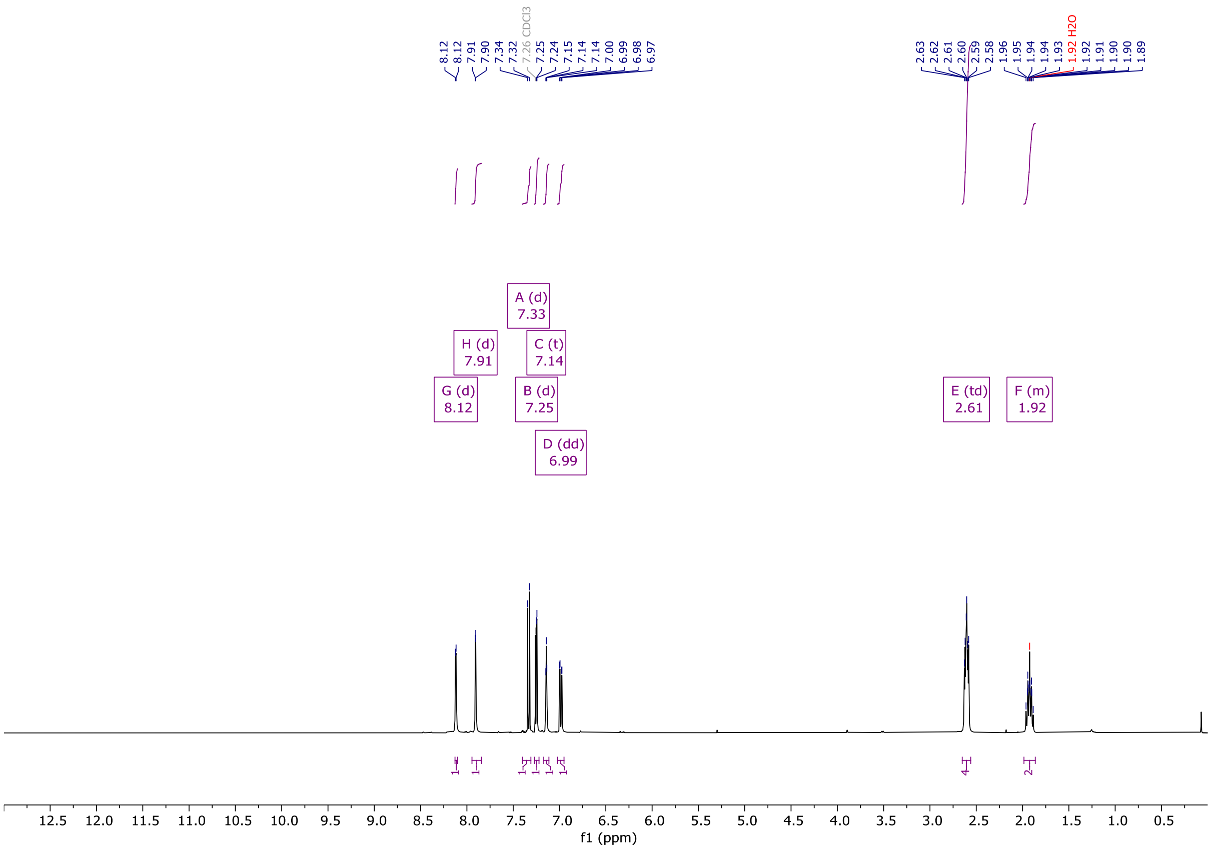


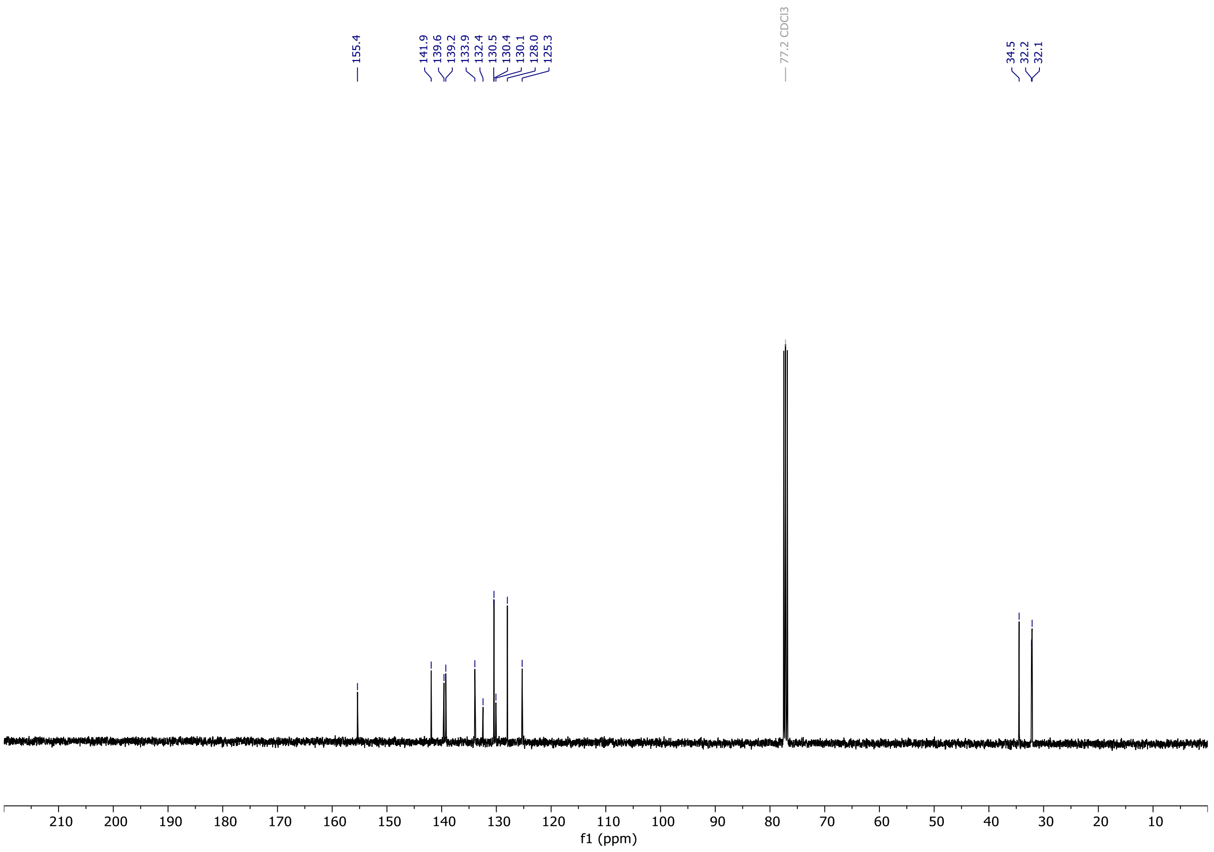

# OX04957

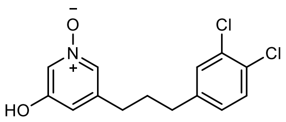


OX04957

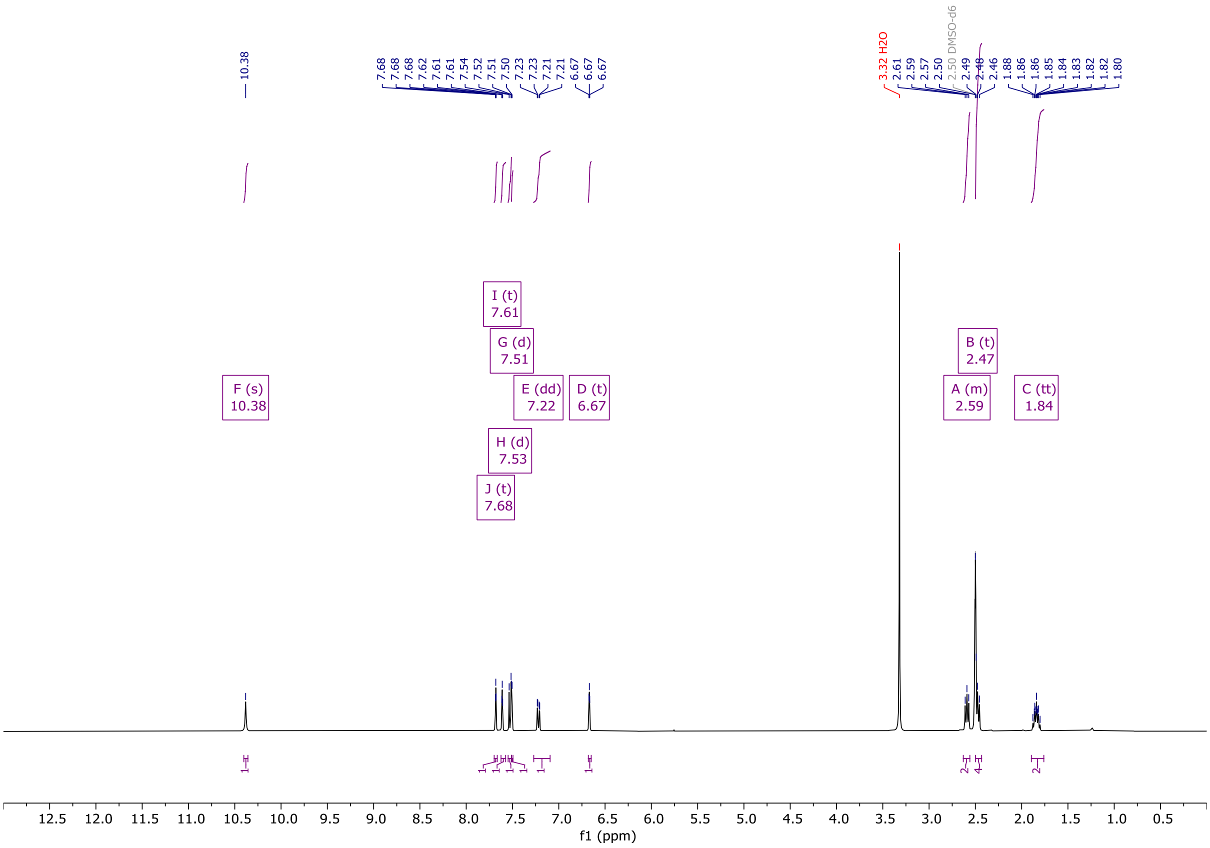


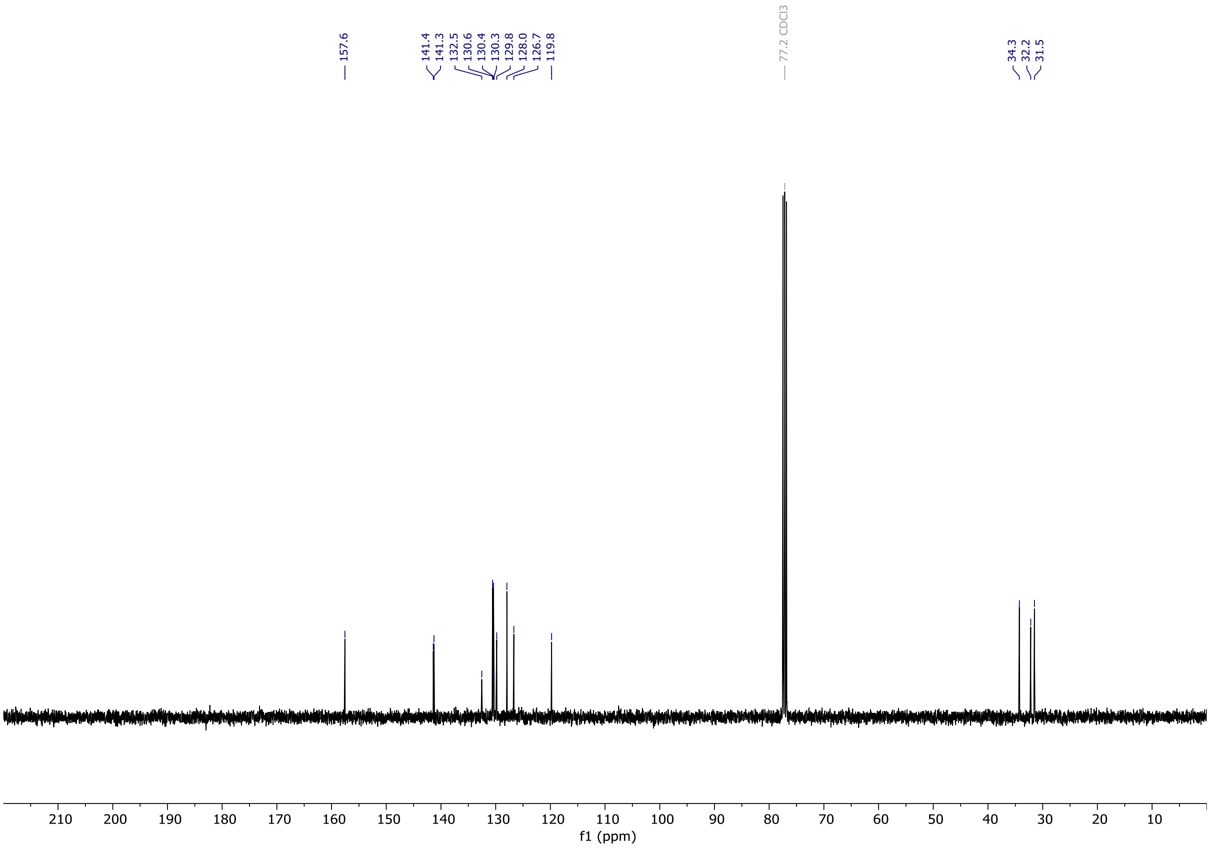

# OX04958a

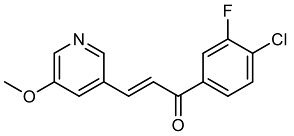


OX04958a

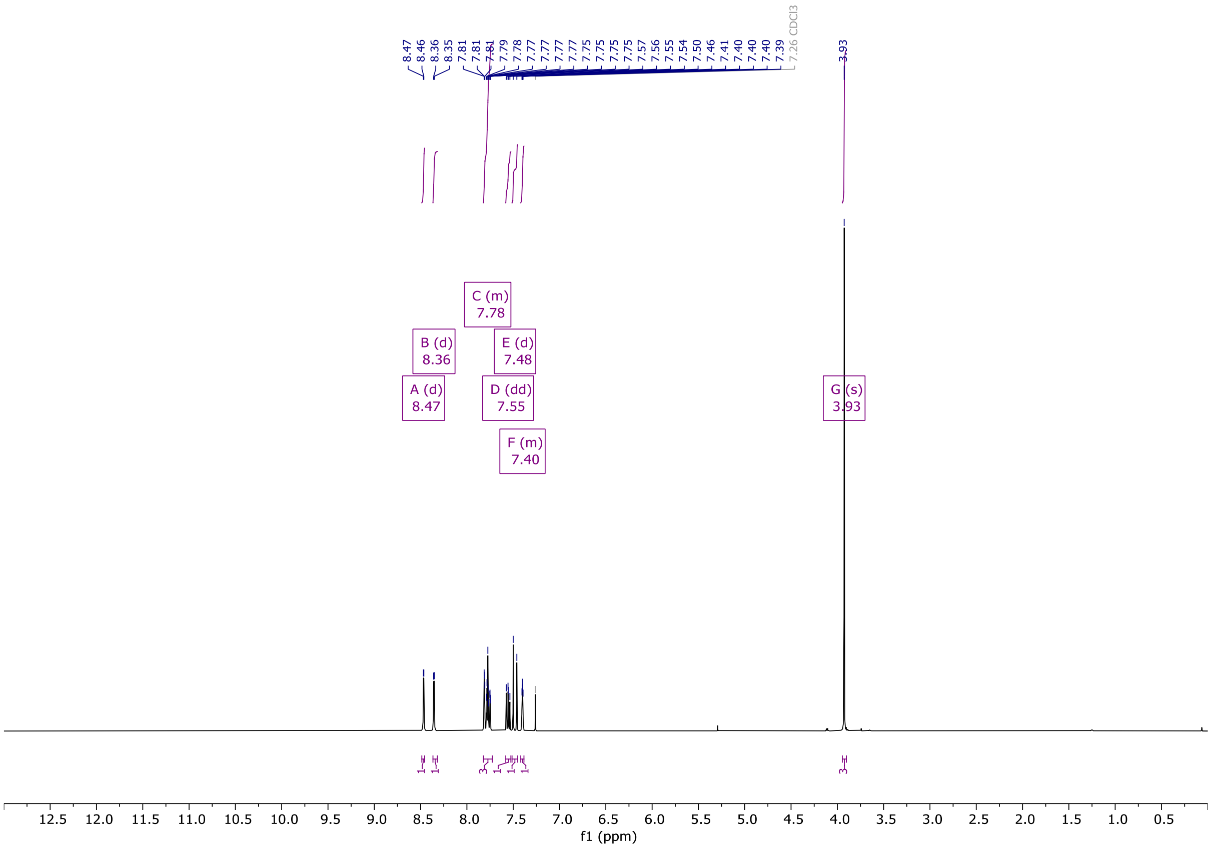


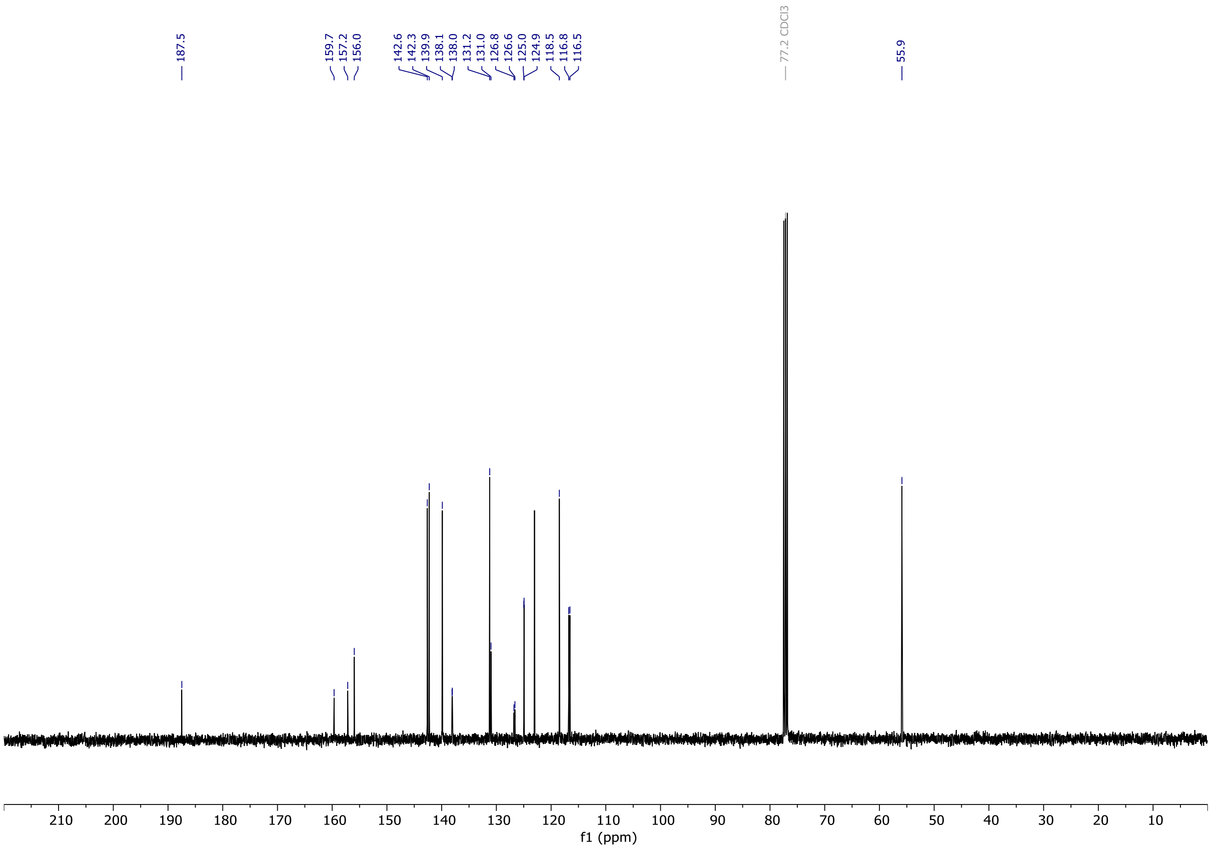

# OX04958b

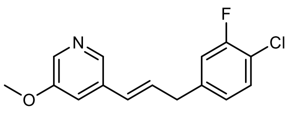


OX04958b

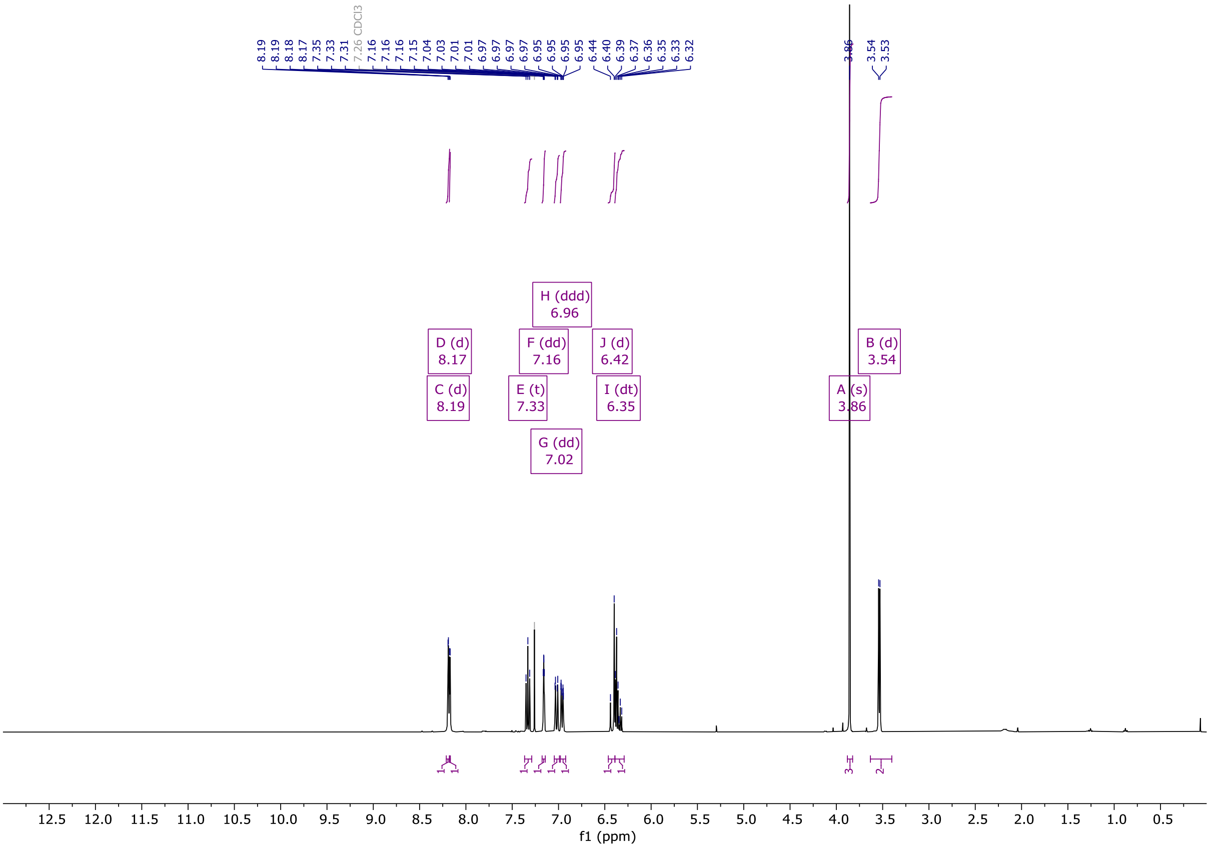


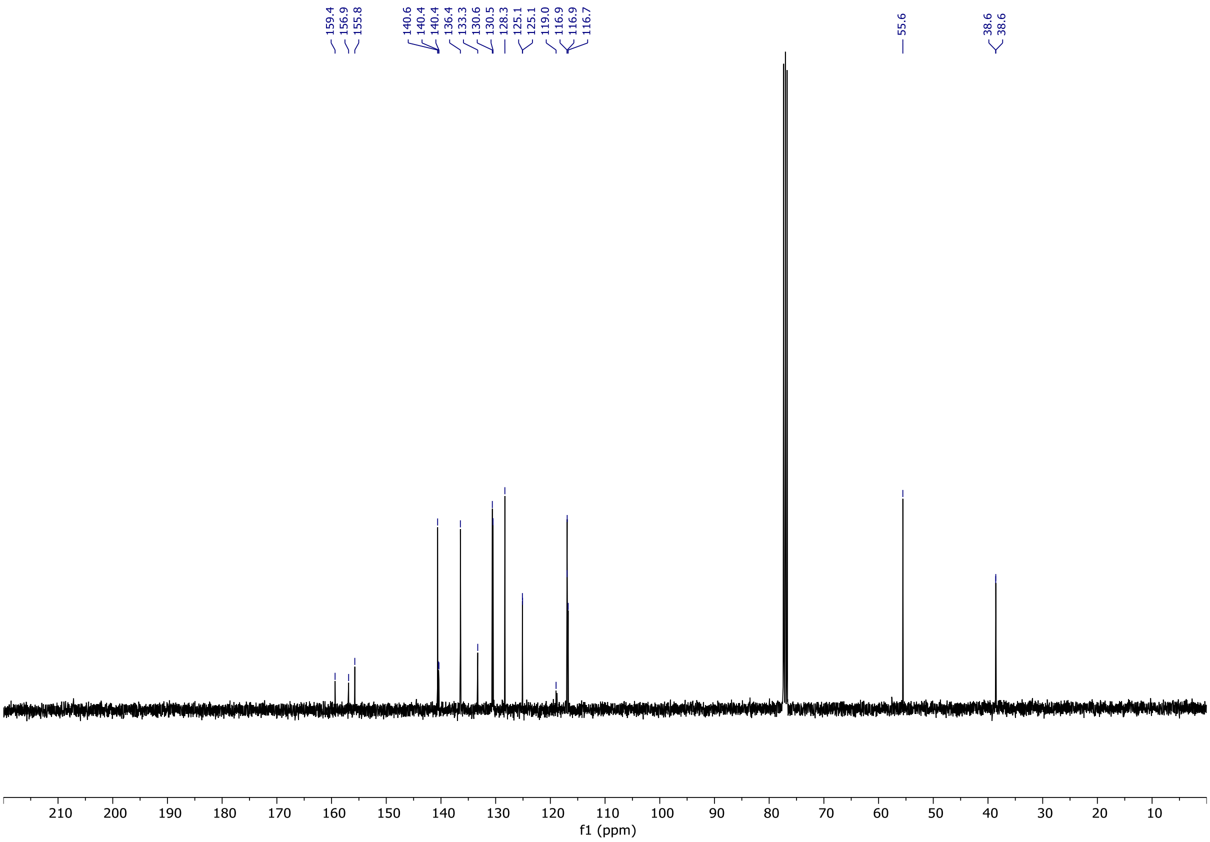

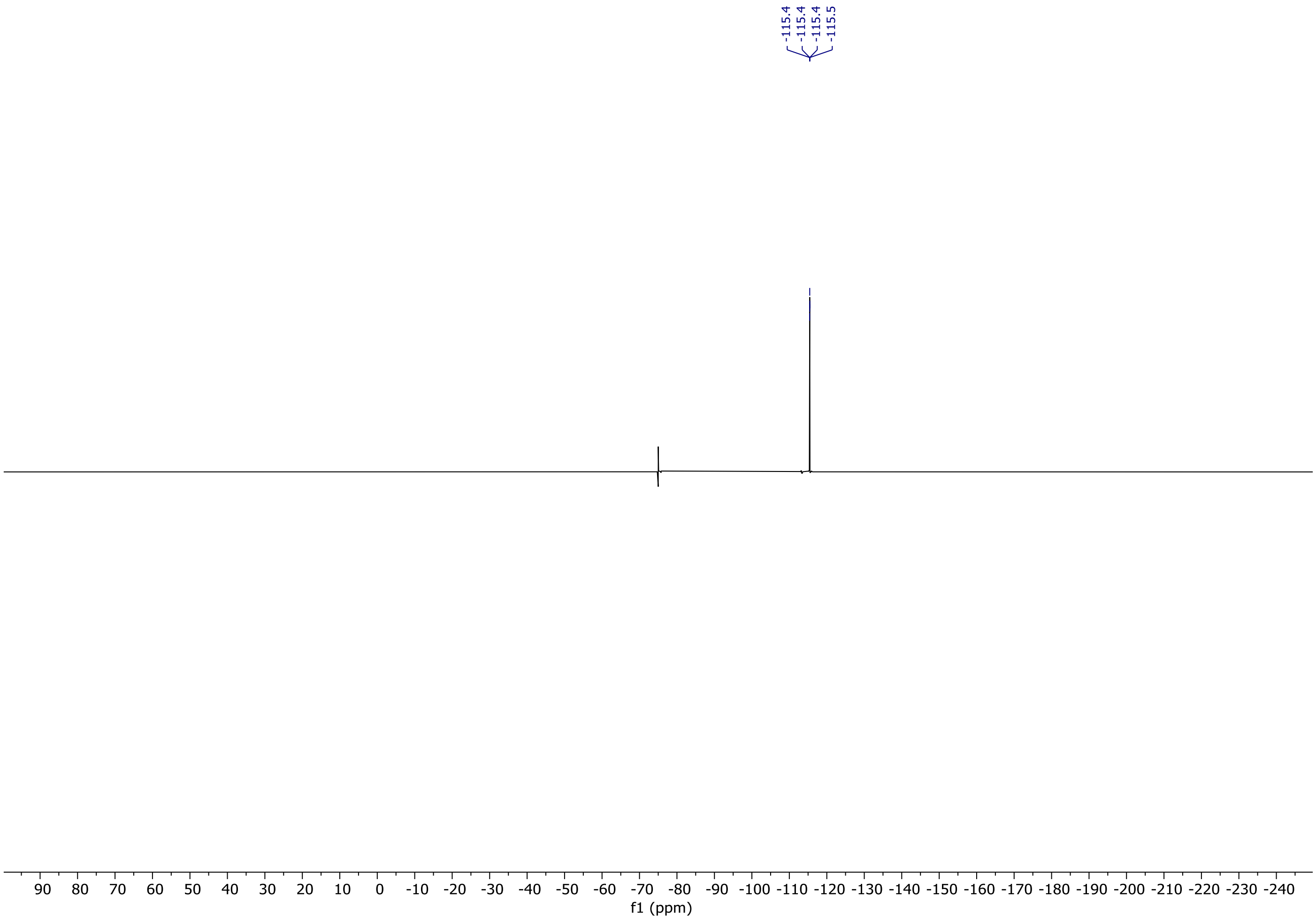
-115.4
-115.4
-115.4
-115.5
90 80 70 60 50 40 30 20 10 0 -10 -20 -30 -40 -50 -60 -70 -80 -90 -100 -110 -120 -130 -140 -150 -160 -170 -180 -190 -200 -210 -220 -230 -240
f1 (ppm)

# OX04958c

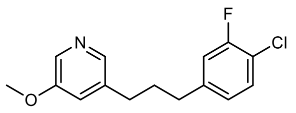

OX04958c

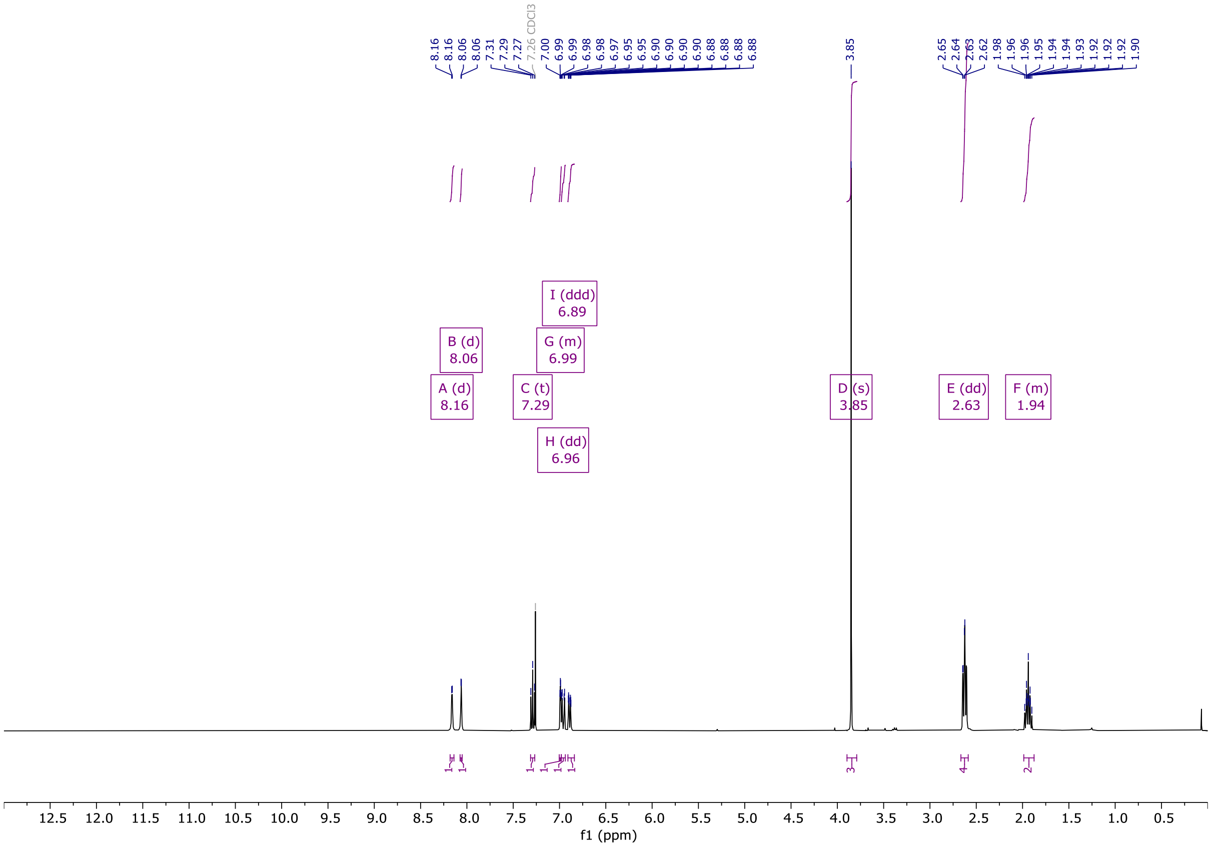

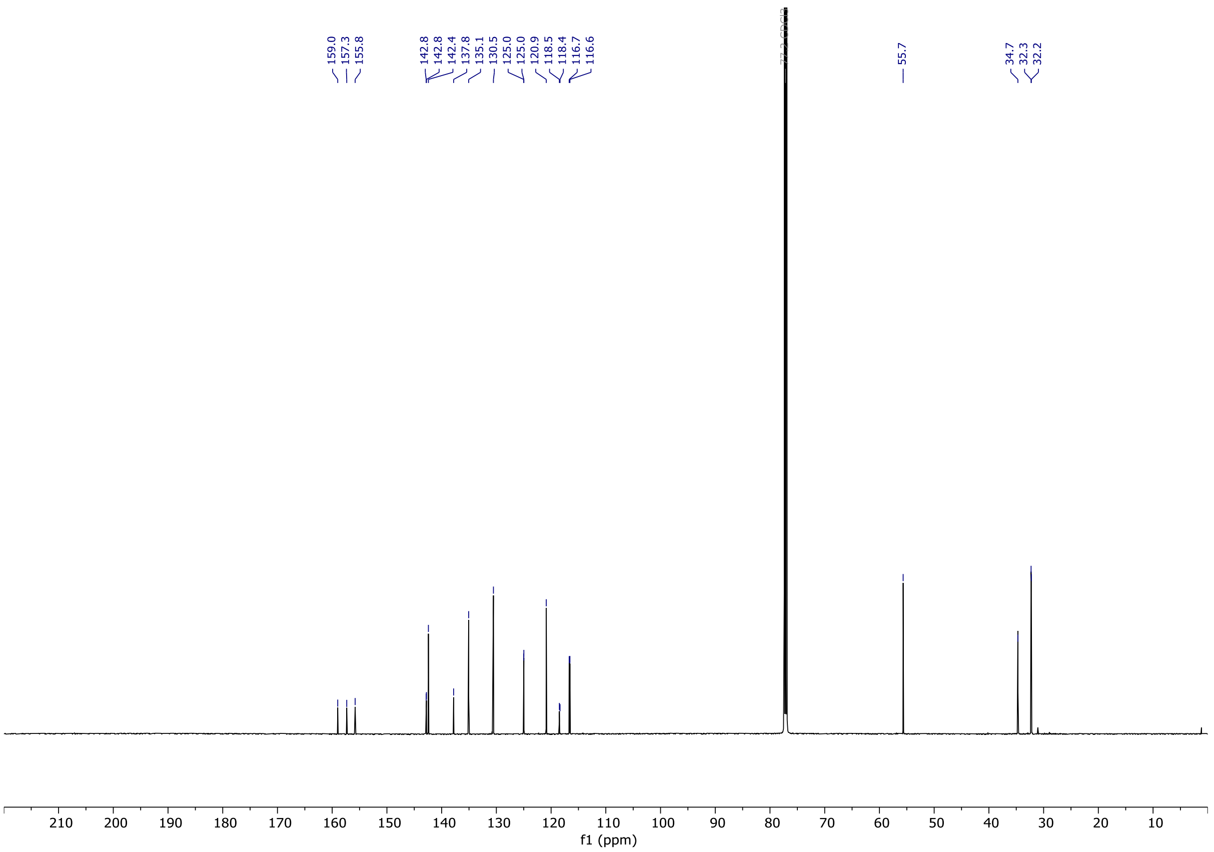

# OX04958d

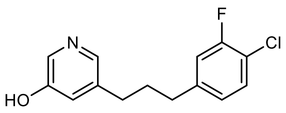

OX04958d

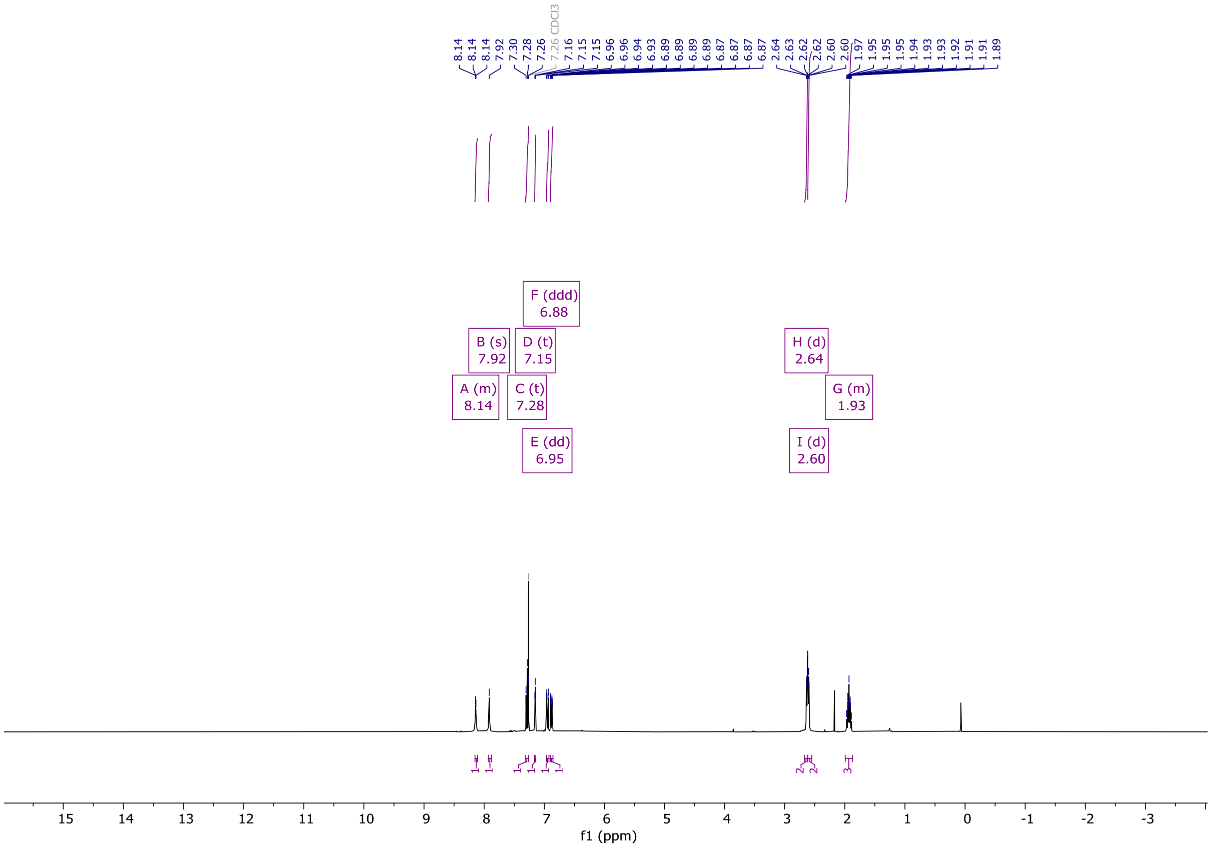

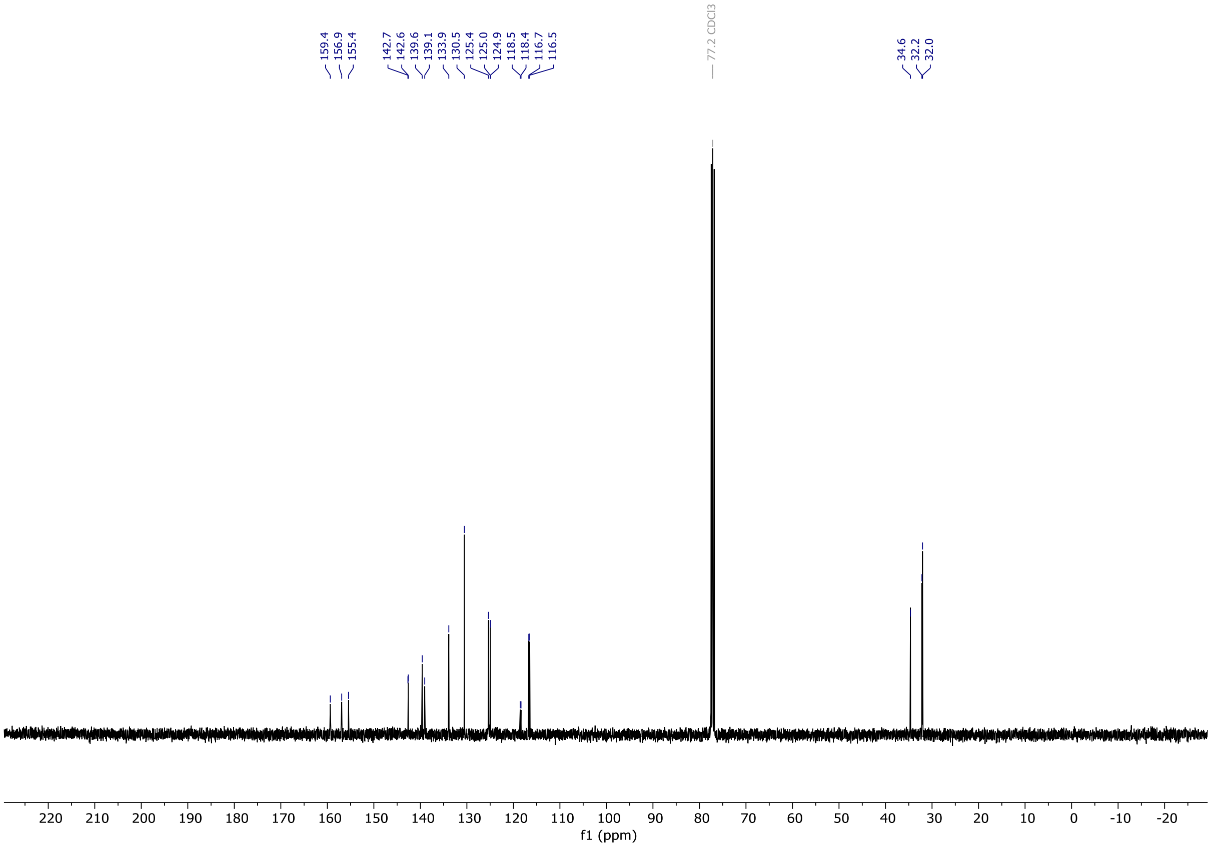

# OX04958

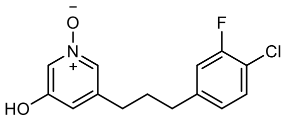


OX04958

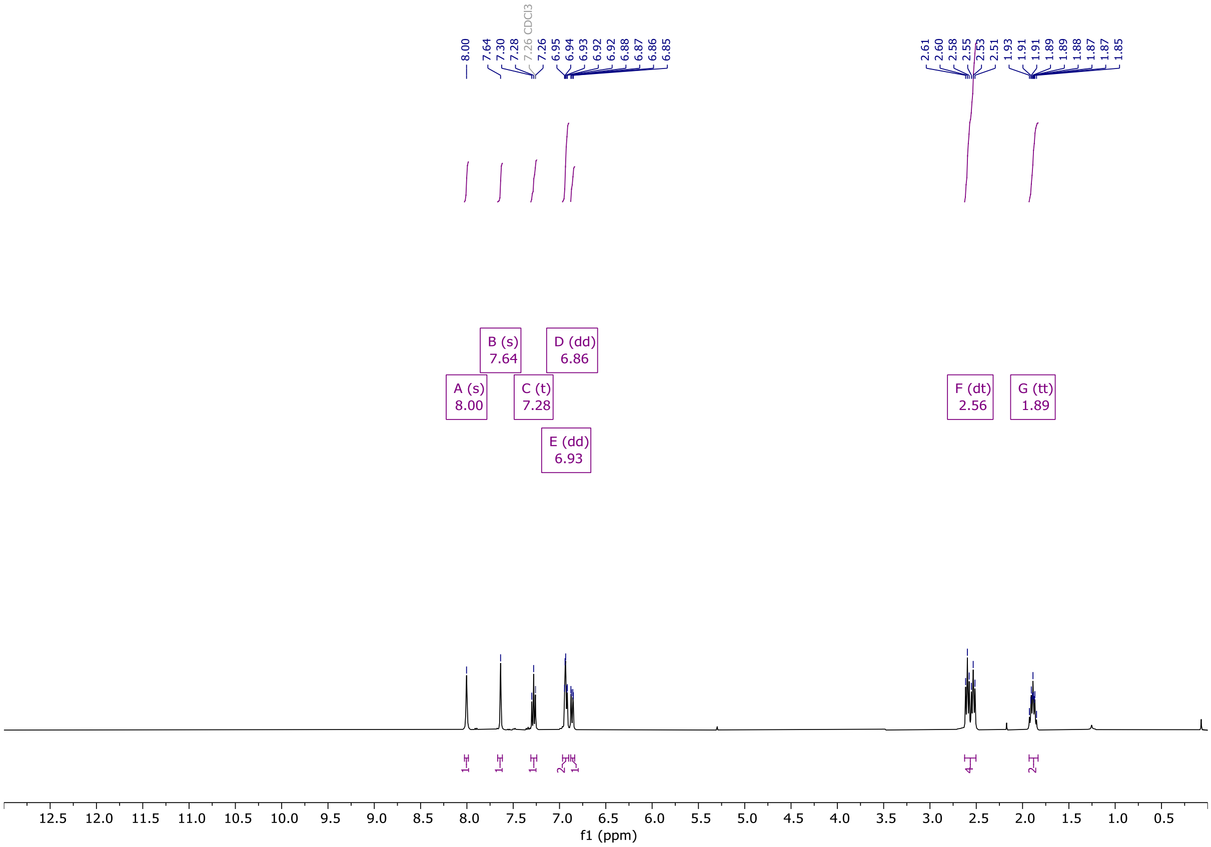


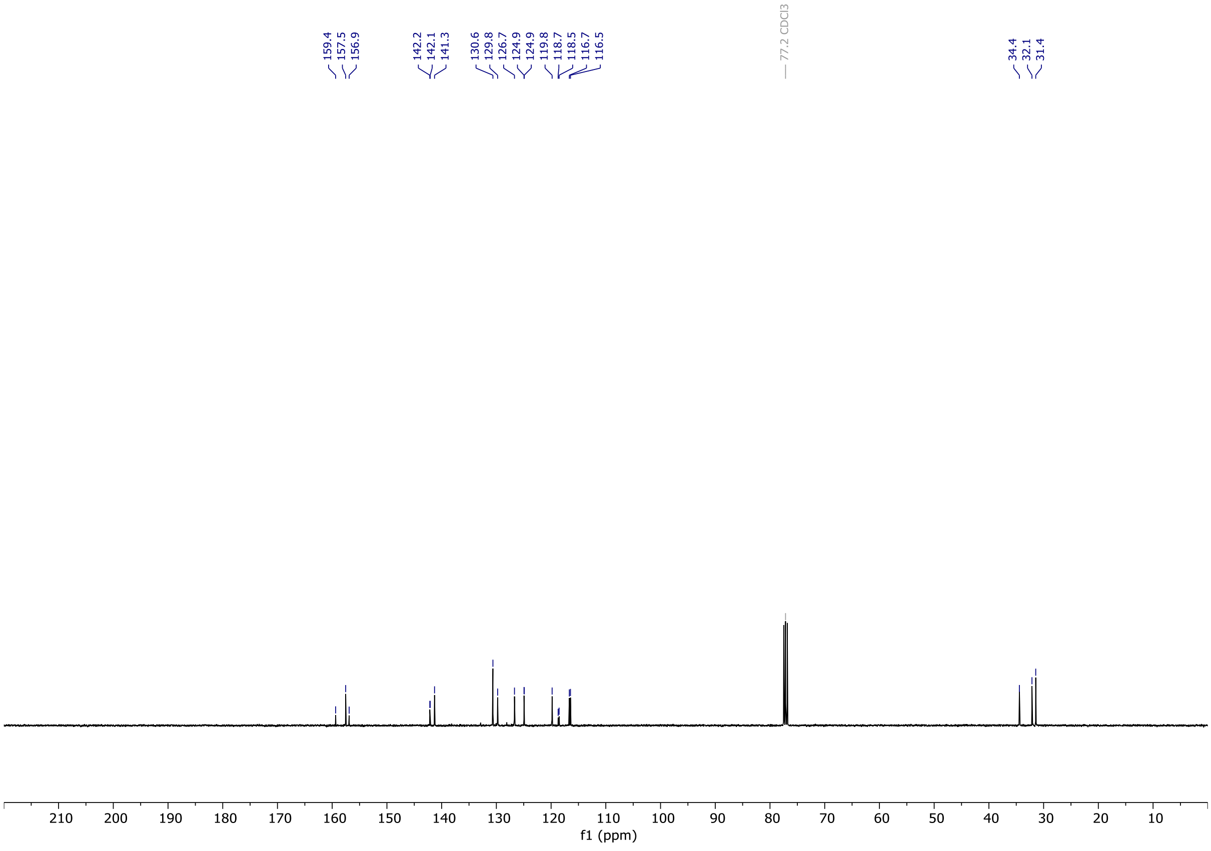

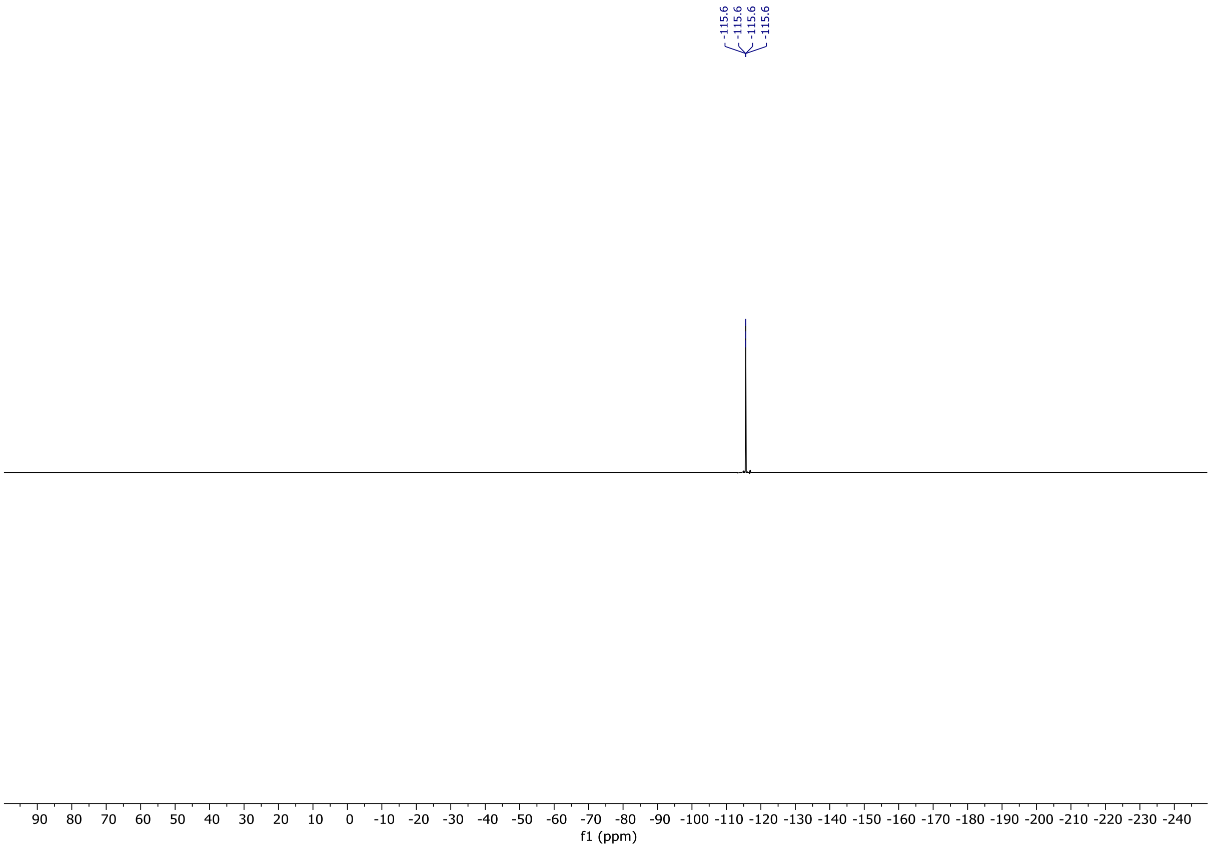

-115.6
-115.6
-115.6
-115.6
90 80 70 60 50 40 30 20 10 0 -10 -20 -30 -40 -50 -60 -70 -80 -90 -100 -110 -120 -130 -140 -150 -160 -170 -180 -190 -200 -210 -220 -230 -240
f1 (ppm)

# OX04959a

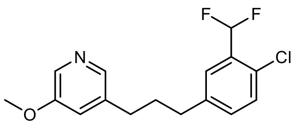

OX04959a

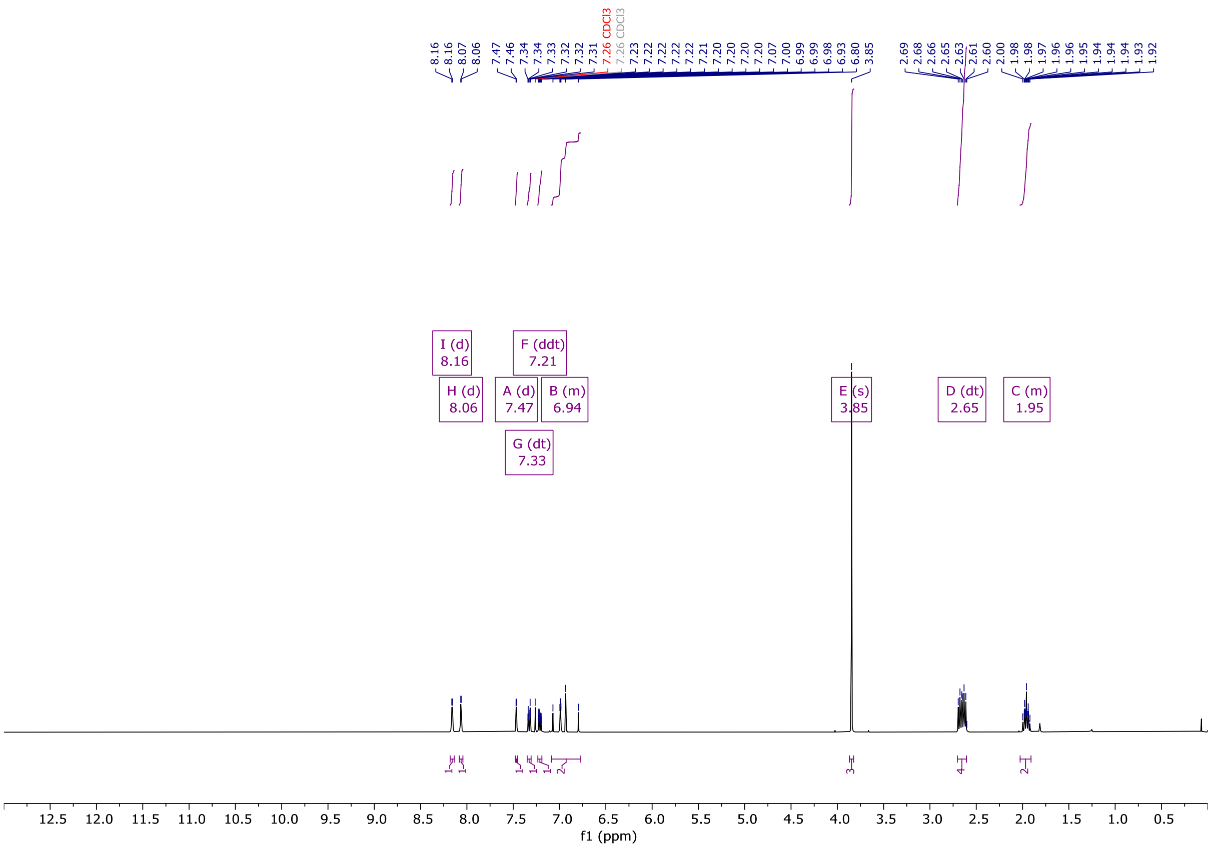

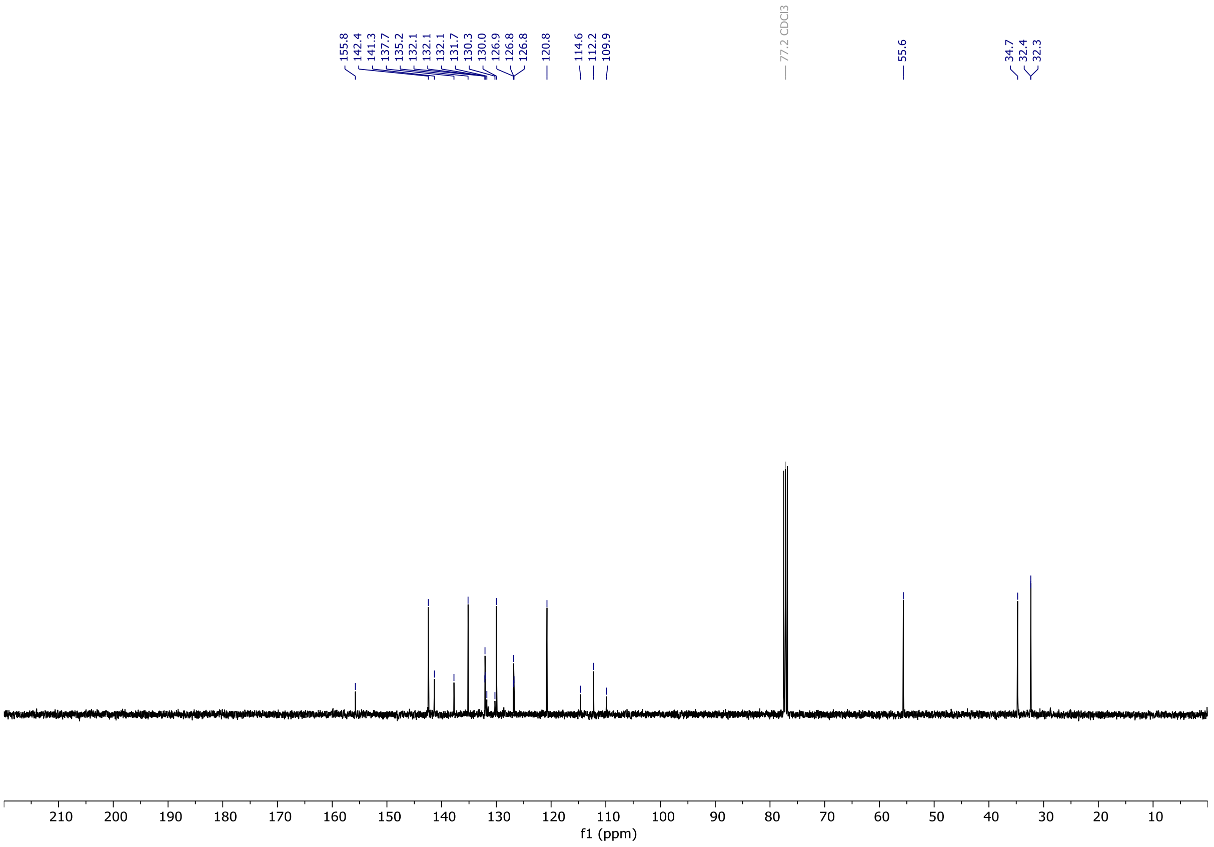

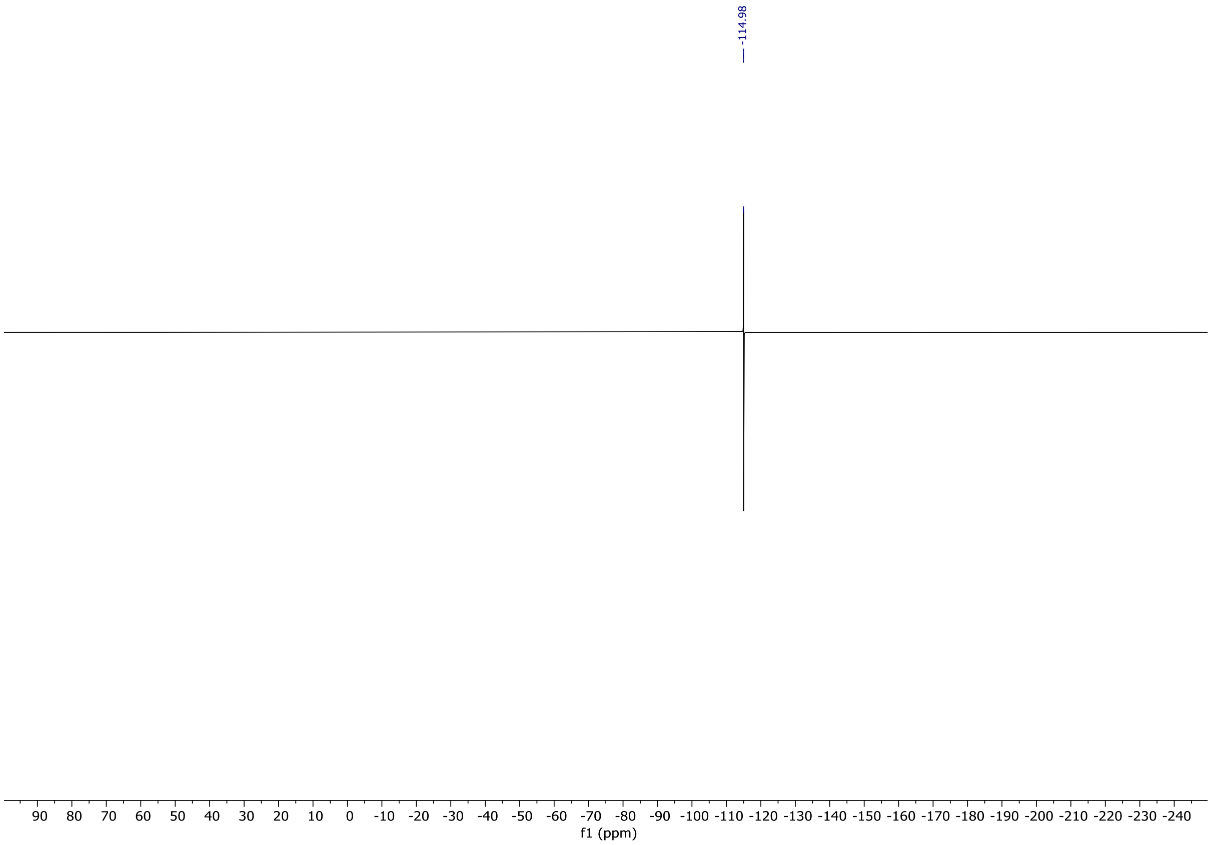
-114.98
90 80 70 60 50 40 30 20 10 0 -10 -20 -30 -40 -50 -60 -70 -80 -90 -100 -110 -120 -130 -140 -150 -160 -170 -180 -190 -200 -210 -220 -230 -240
f1 (ppm)

# OX04959b

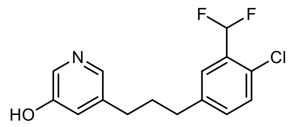


OX04959b

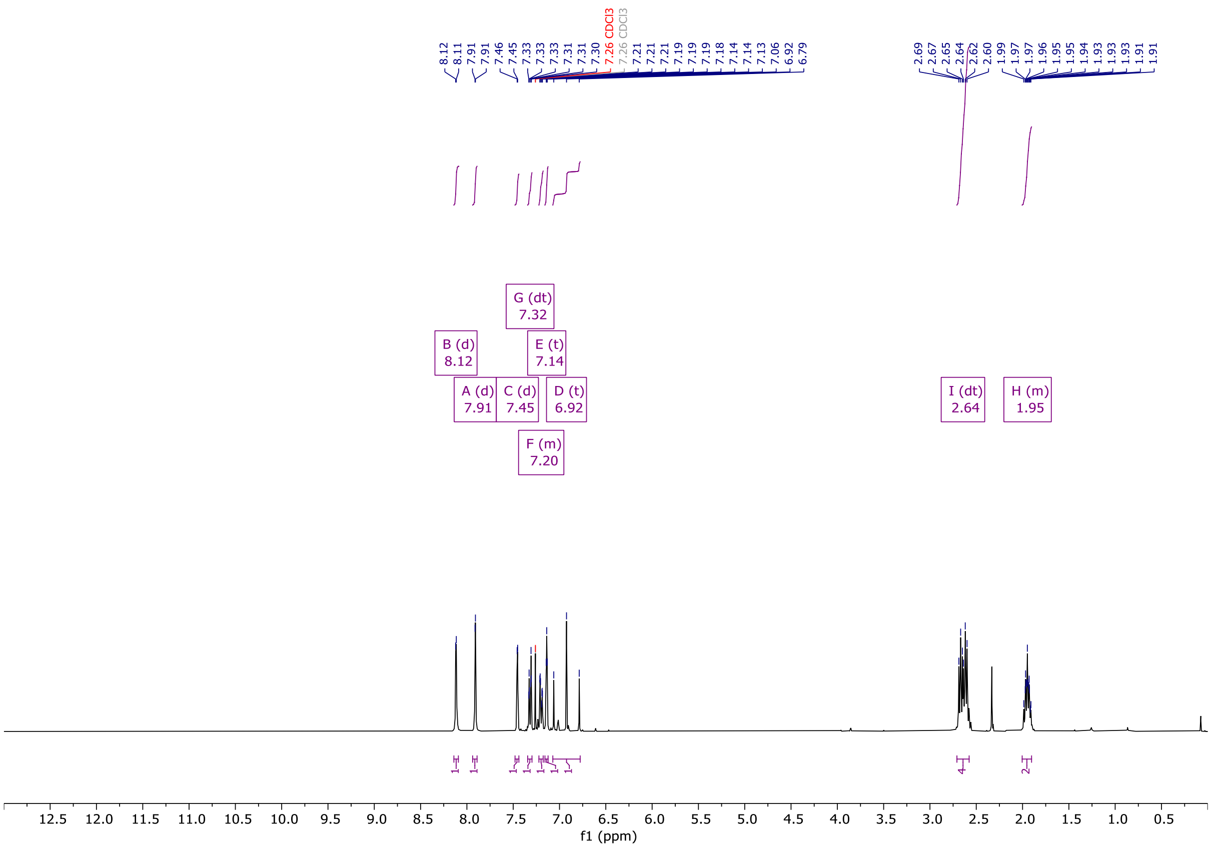


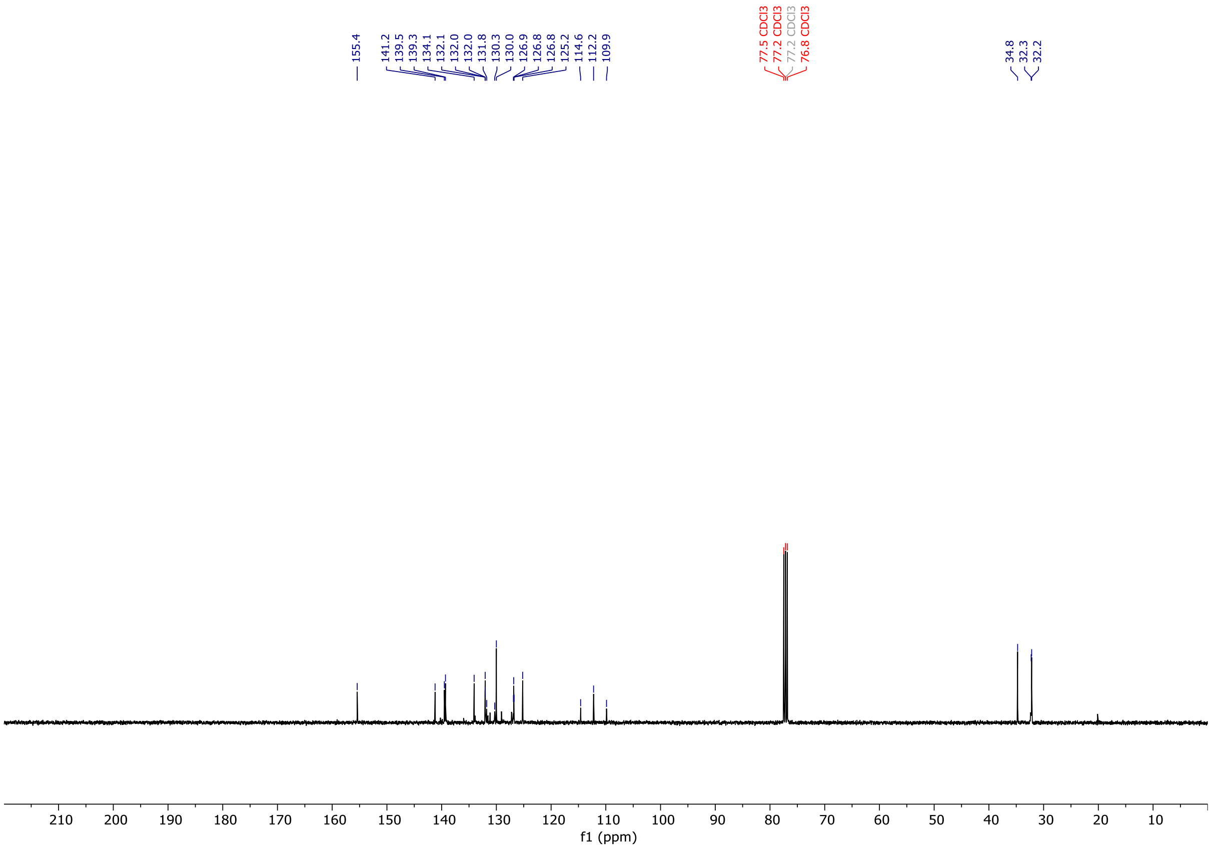

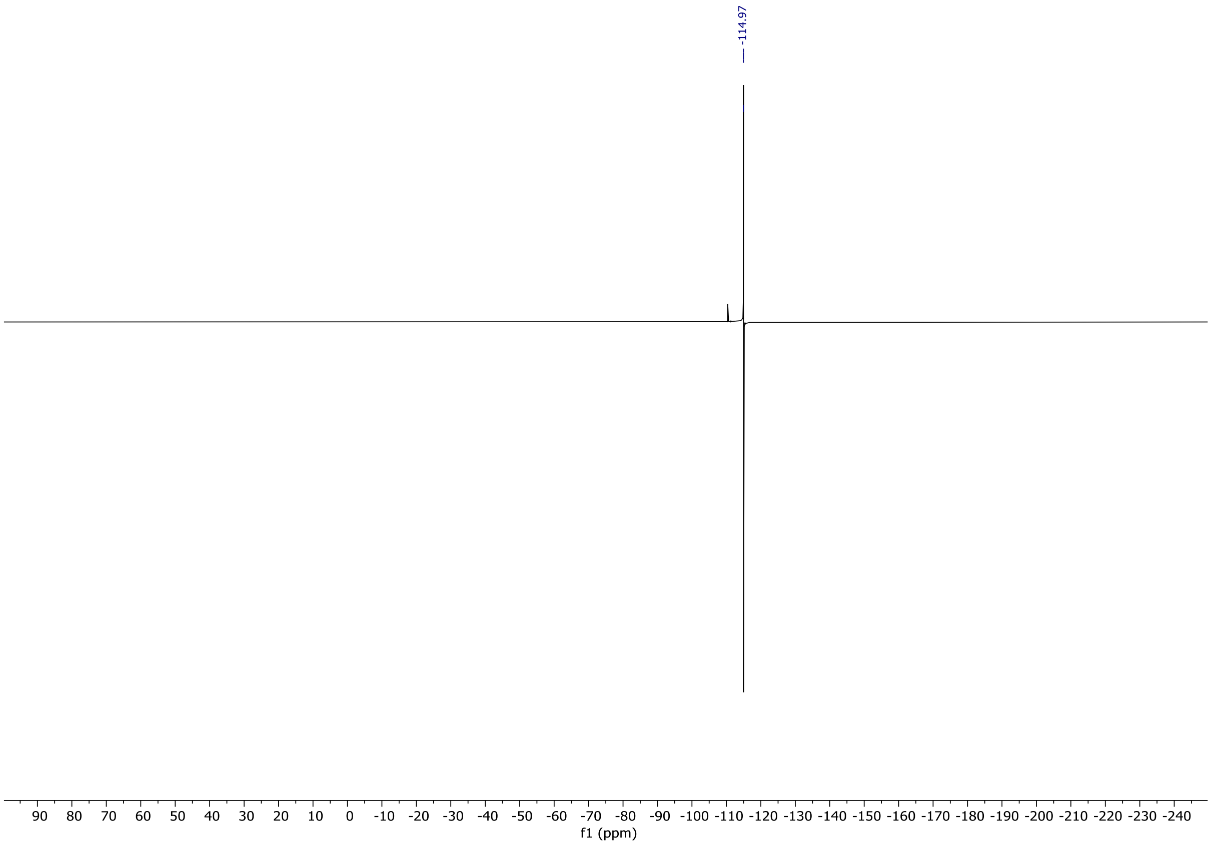
-114.97
90 80 70 60 50 40 30 20 10 0 -10 -20 -30 -40 -50 -60 -70 -80 -90 -100 -110 -120 -130 -140 -150 -160 -170 -180 -190 -200 -210 -220 -230 -240
f1 (ppm)

# OX04959

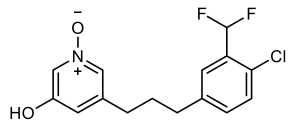


OX04959

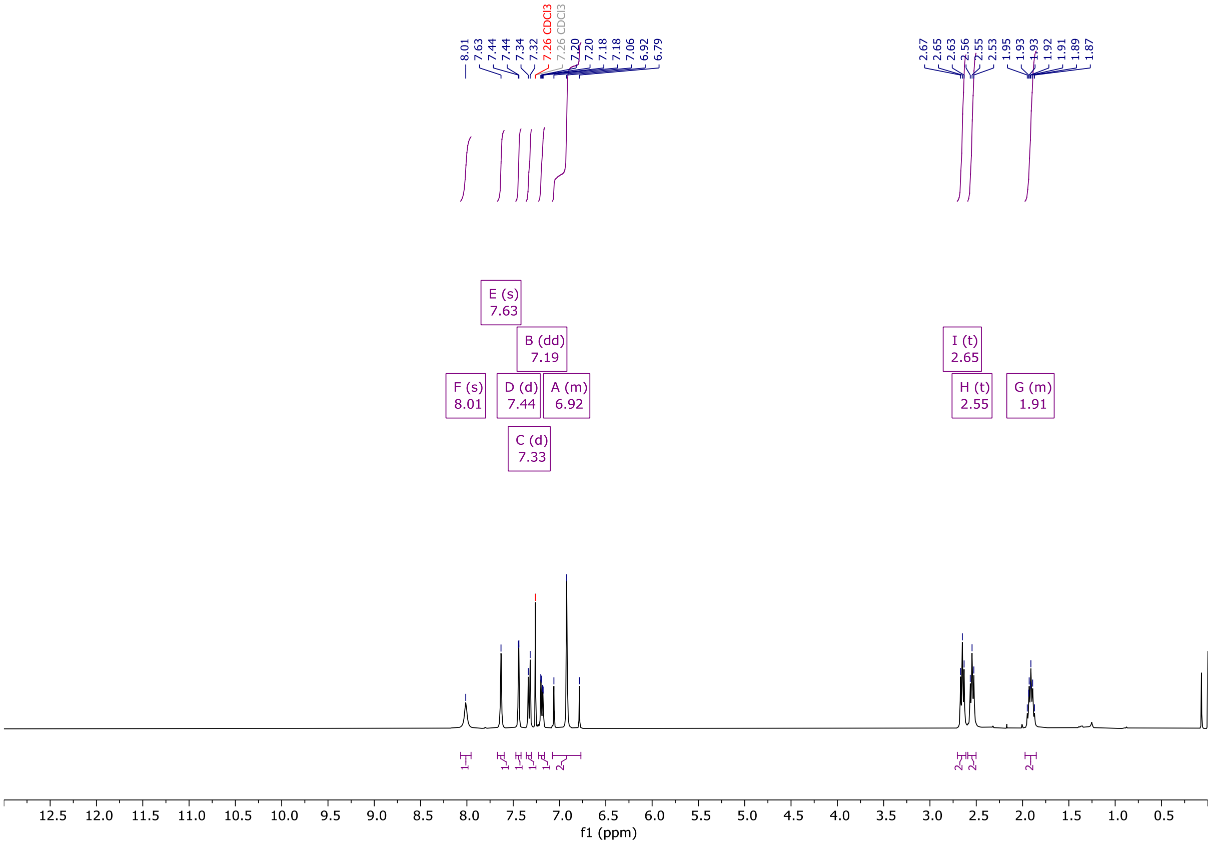


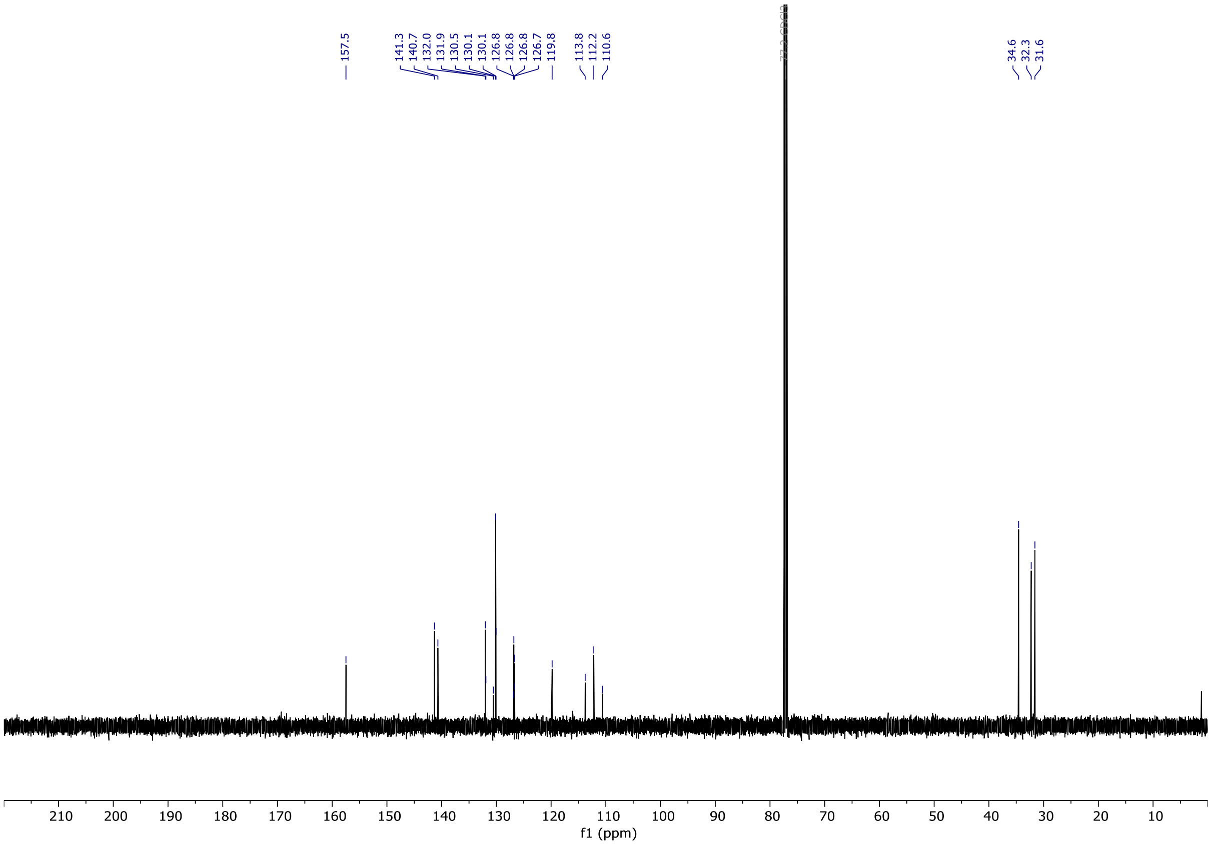

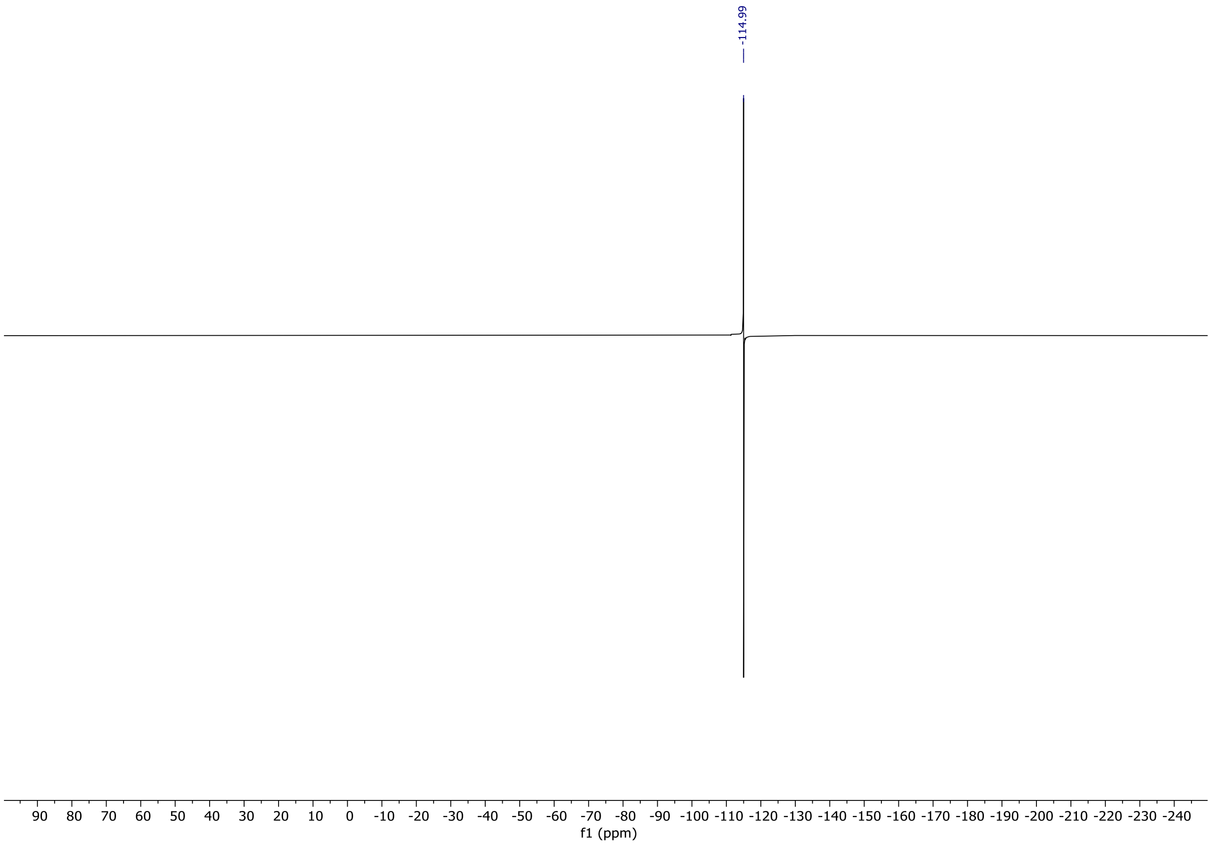
-114.99
90 80 70 60 50 40 30 20 10 0 -10 -20 -30 -40 -50 -60 -70 -80 -90 -100 -110 -120 -130 -140 -150 -160 -170 -180 -190 -200 -210 -220 -230 -240
f1 (ppm)

# OX04960a

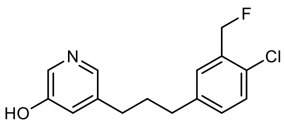


OX04960a

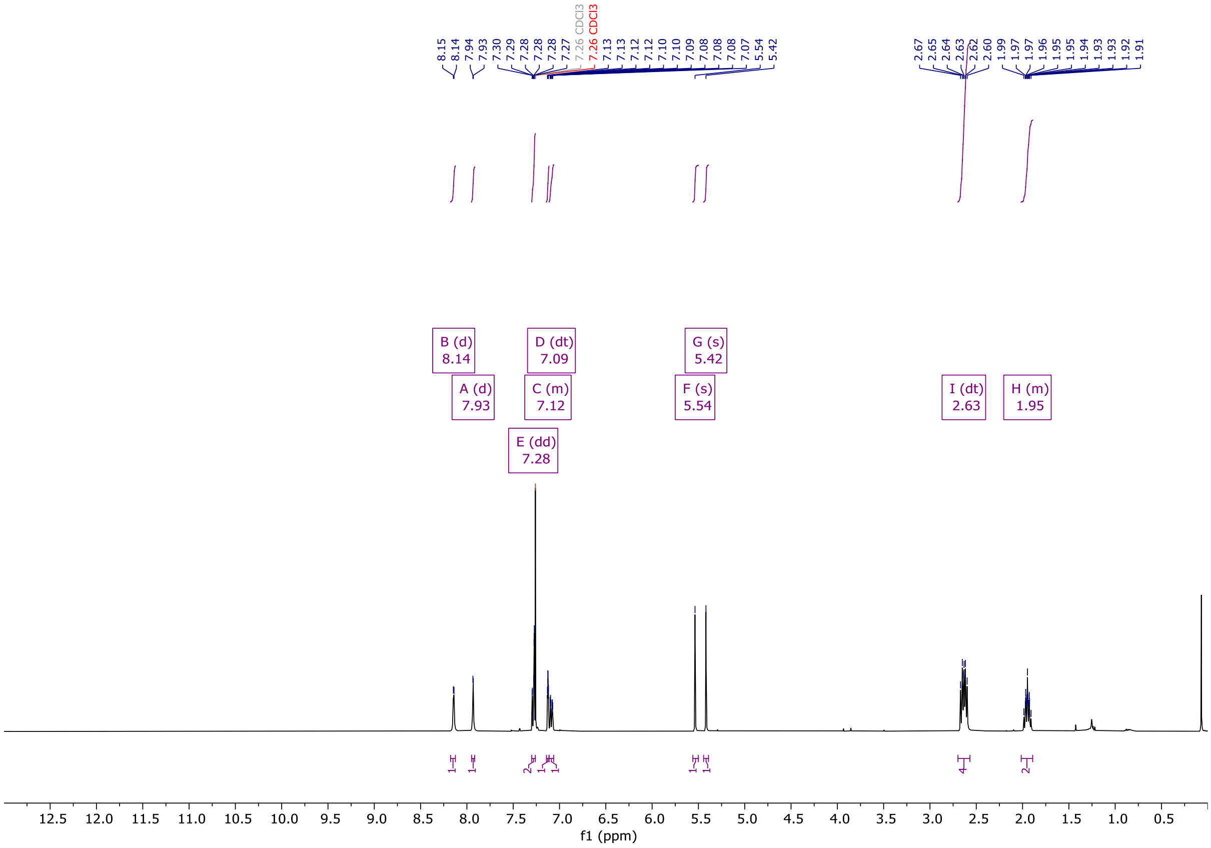


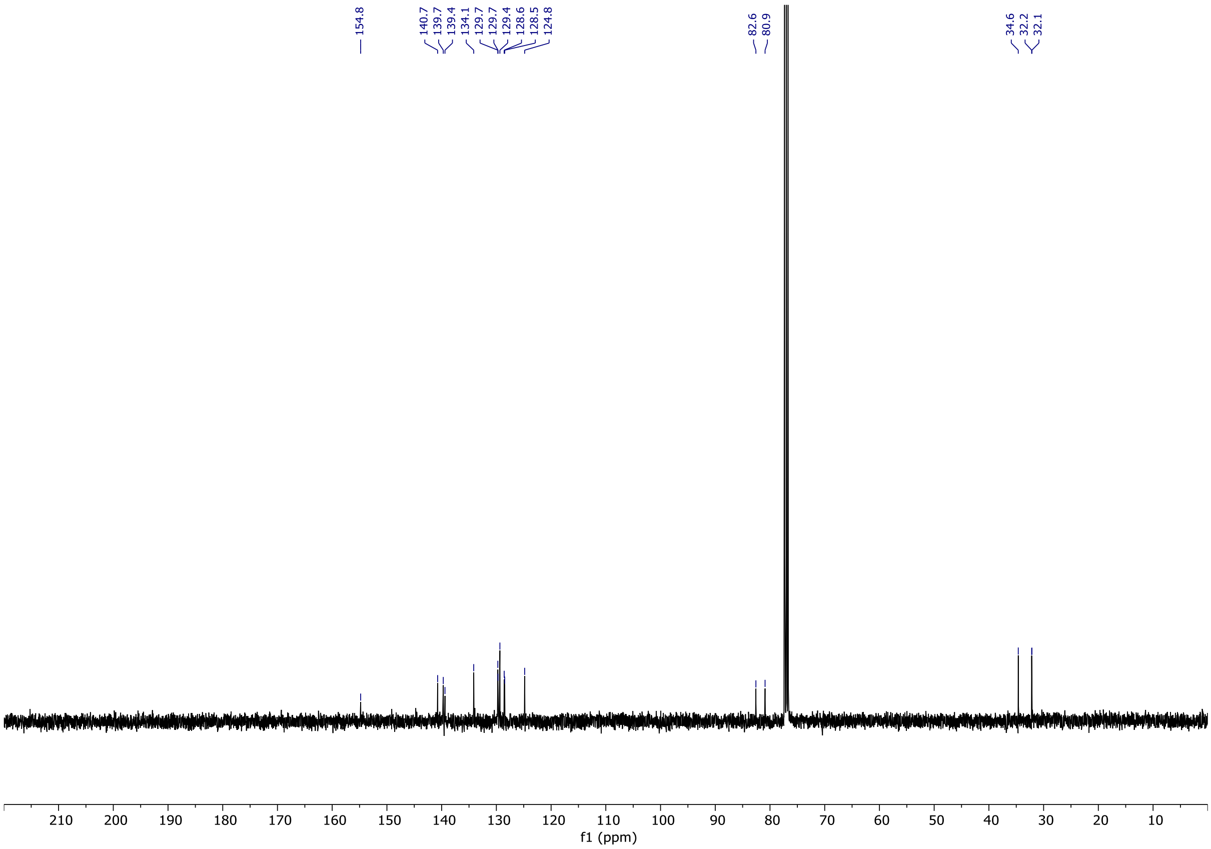

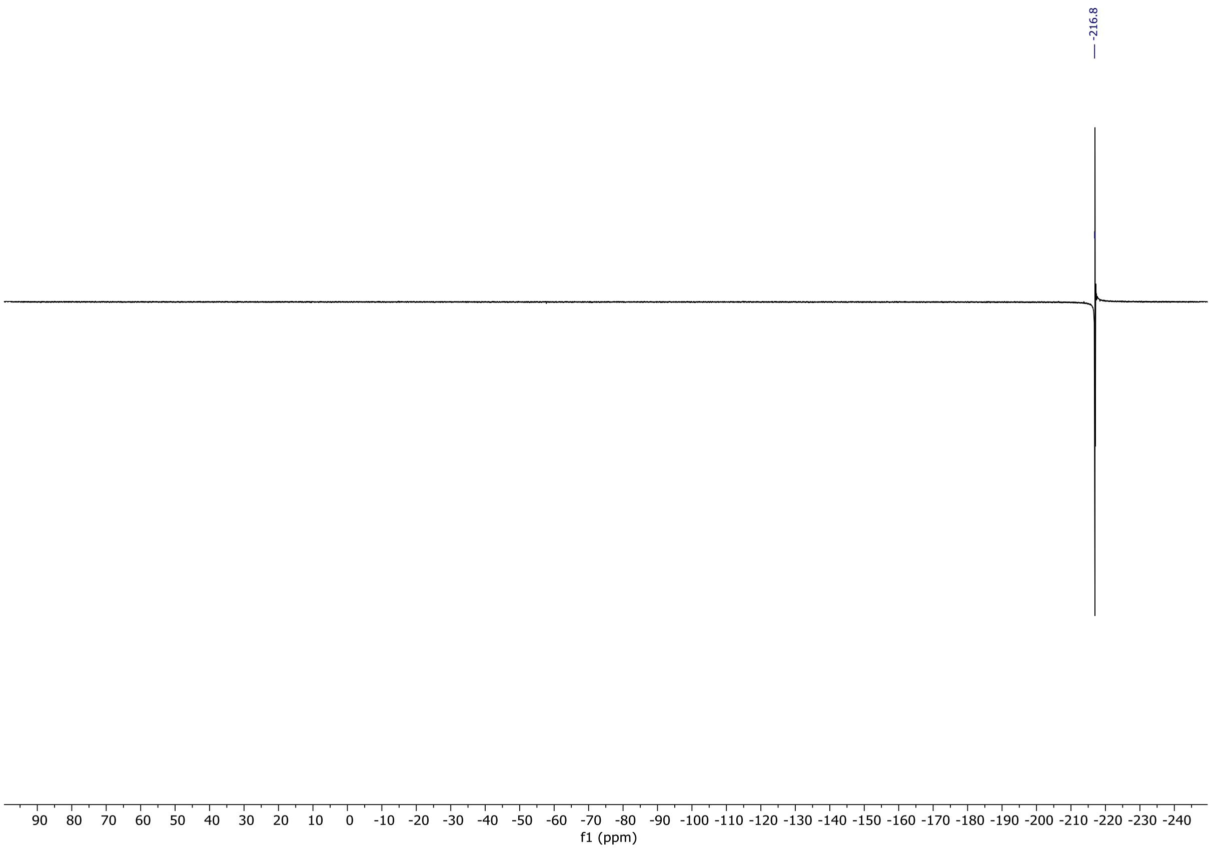
-216.8
90 80 70 60 50 40 30 20 10 0 -10 -20 -30 -40 -50 -60 -70 -80 -90 -100 -110 -120 -130 -140 -150 -160 -170 -180 -190 -200 -210 -220 -230 -240
f1 (ppm)

# OX04960

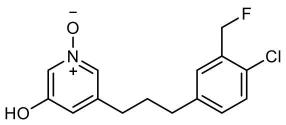

OX04960

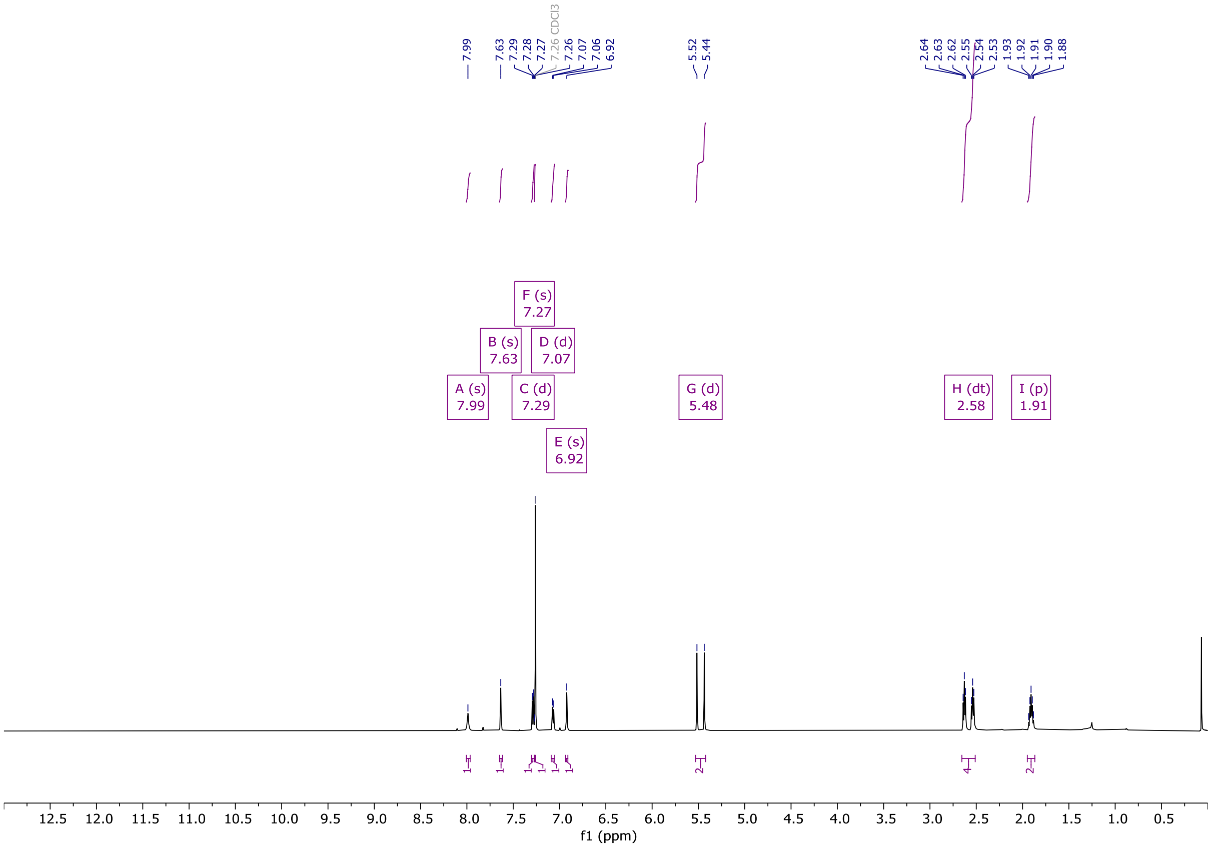


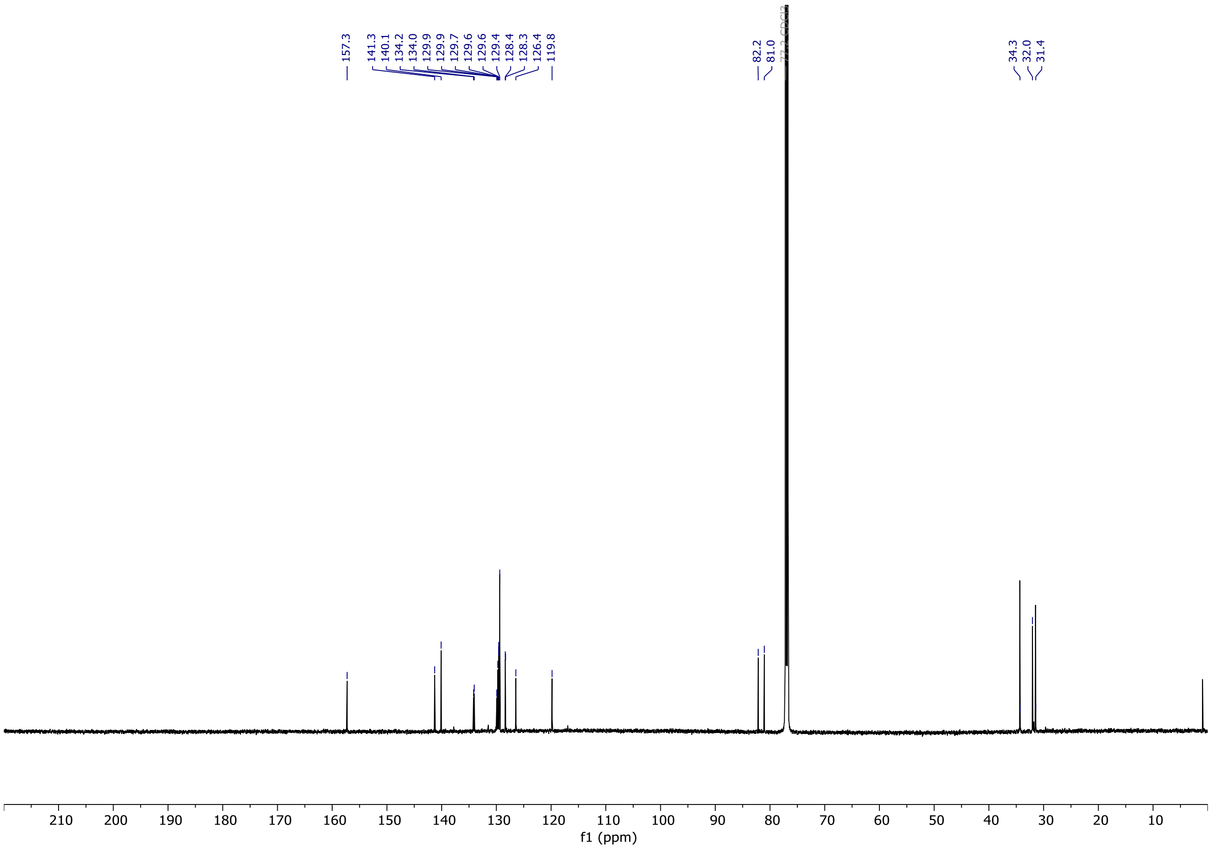

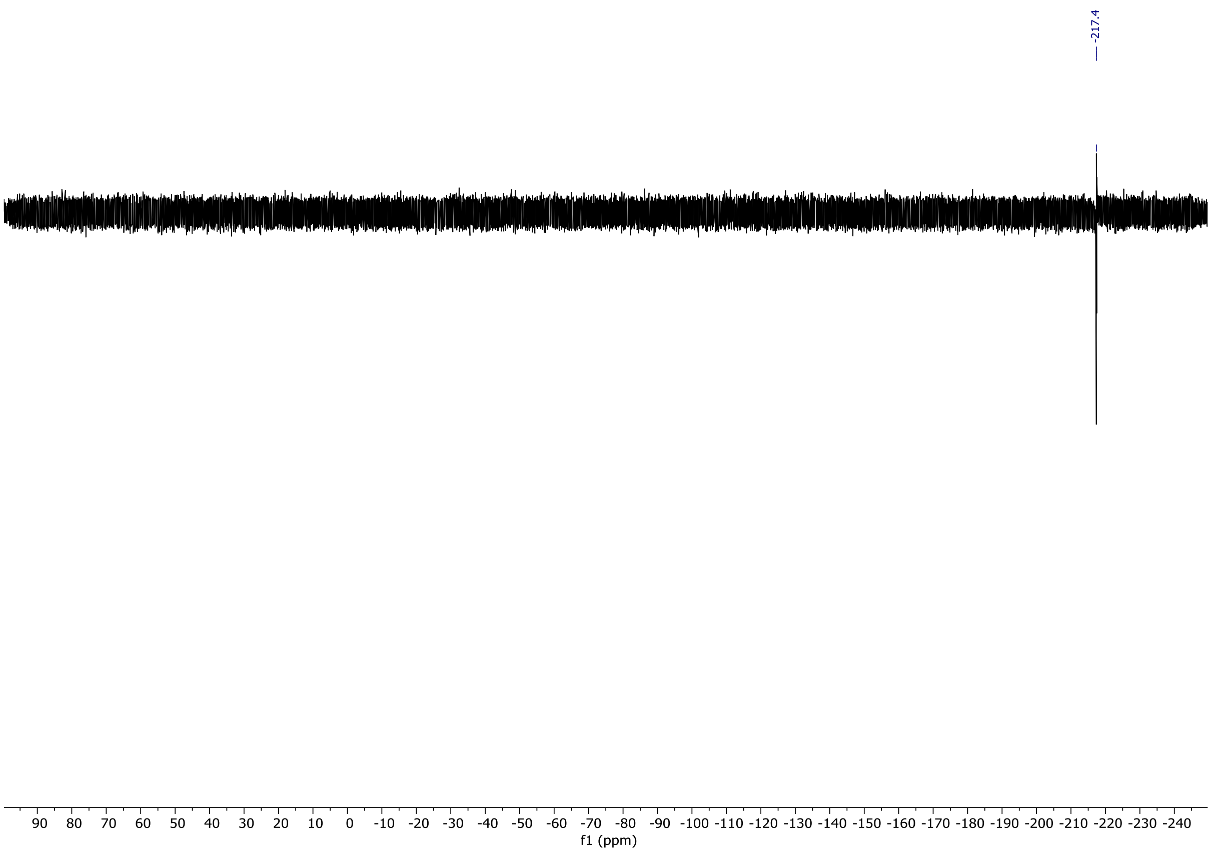

-217.4
90 80 70 60 50 40 30 20 10 0 -10 -20 -30 -40 -50 -60 -70 -80 -90 -100 -110 -120 -130 -140 -150 -160 -170 -180 -190 -200 -210 -220 -230 -240
f1 (ppm)

# OX04961a

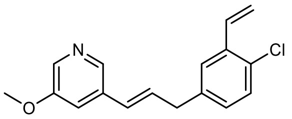


OX04961a

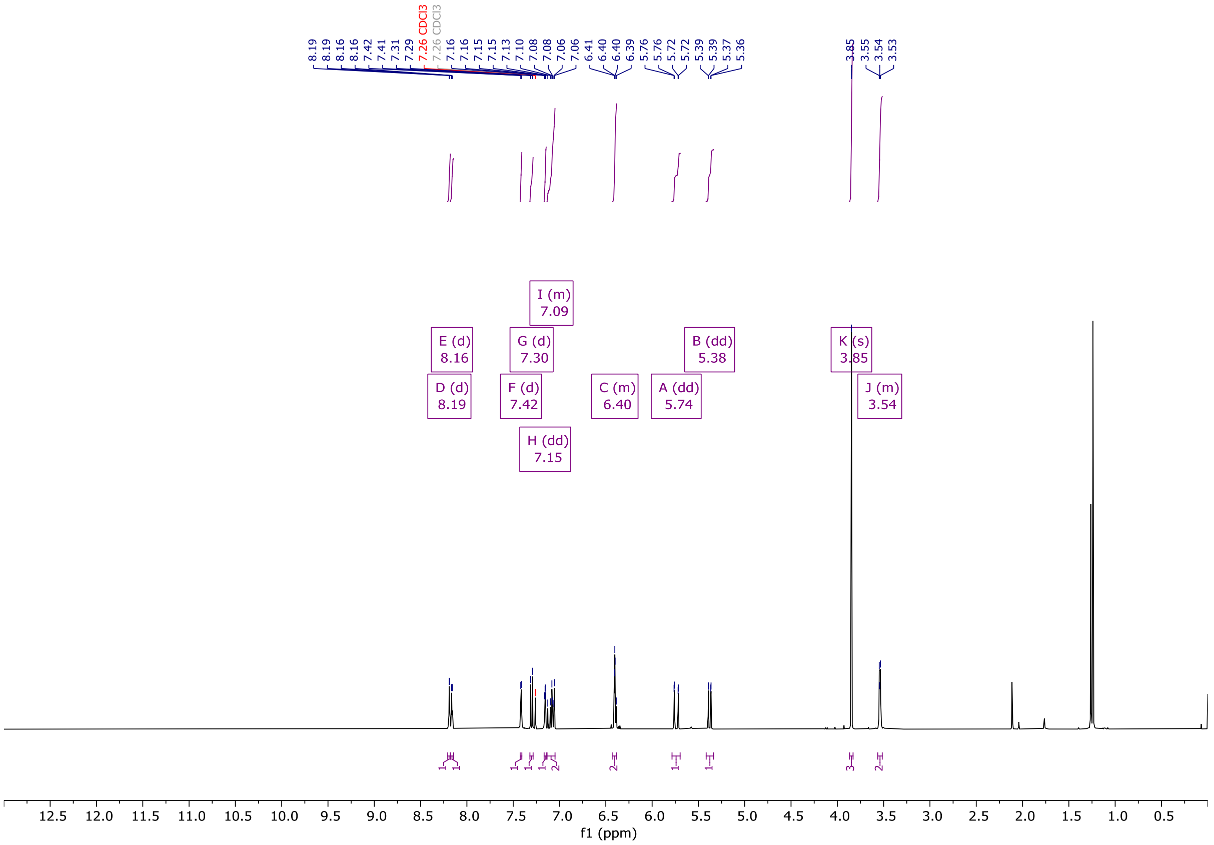


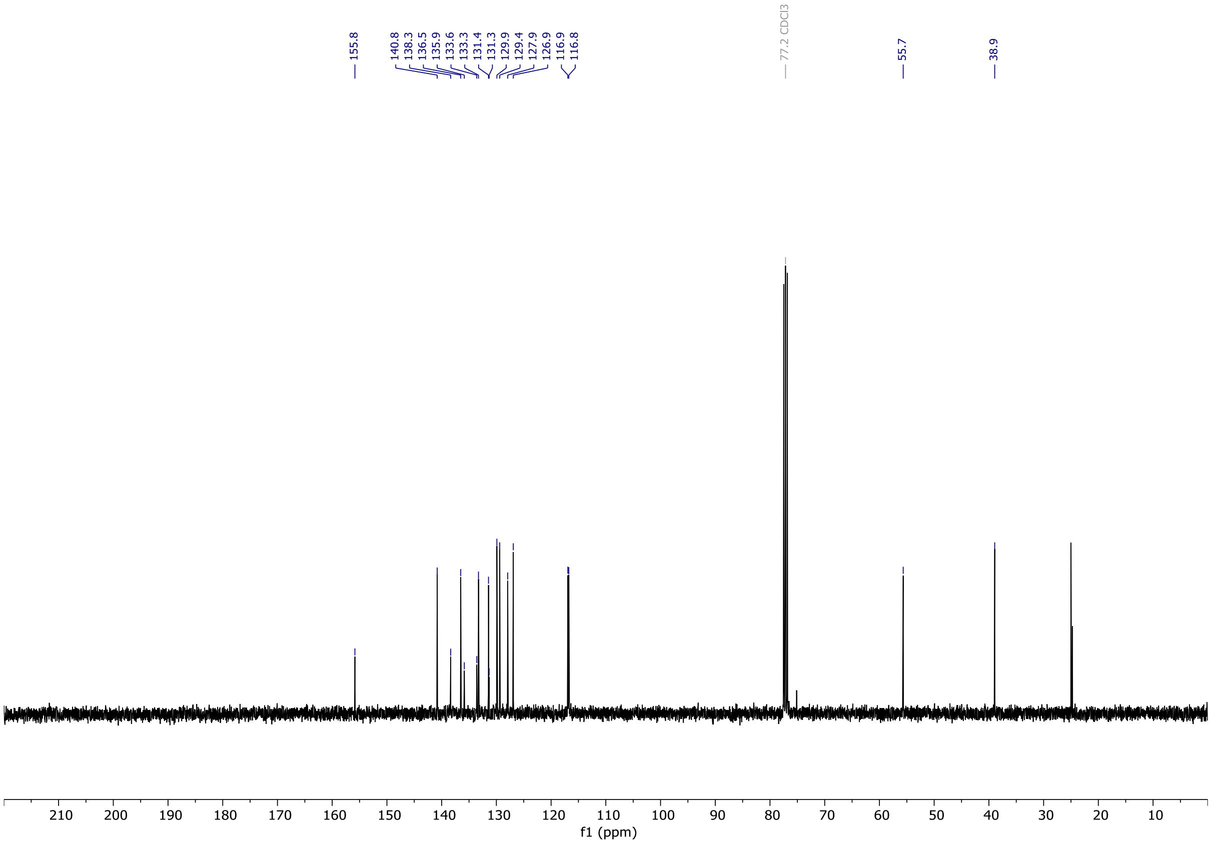

# OX04961b

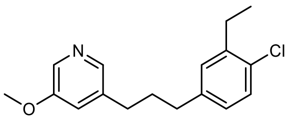

OX04961b

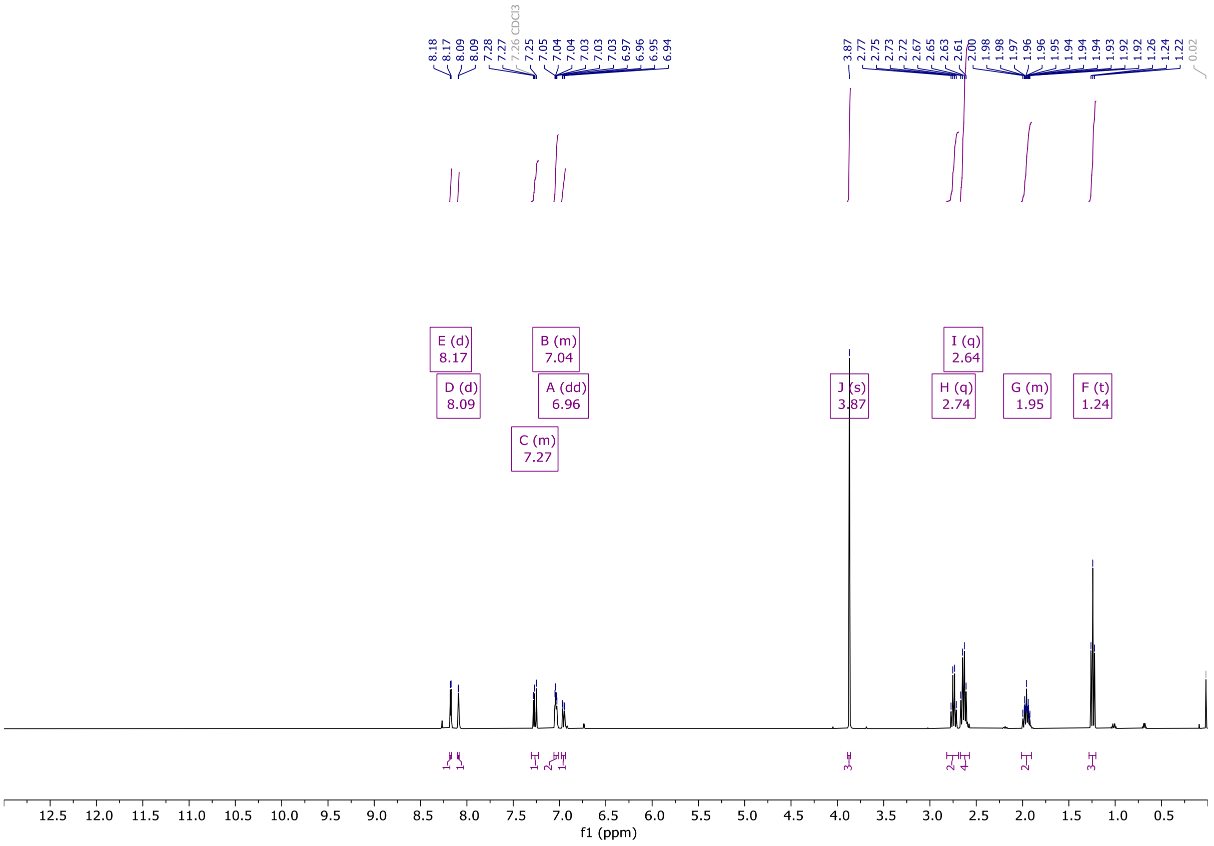

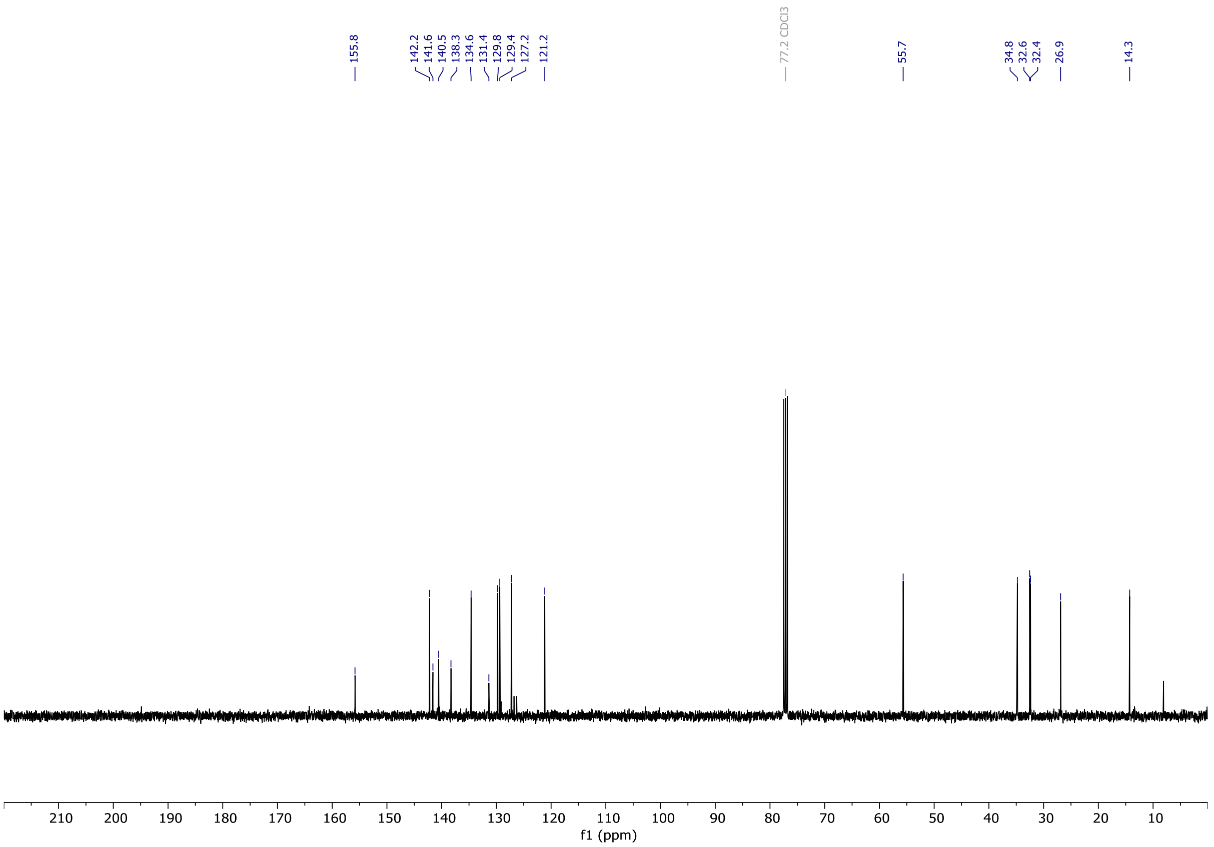

# OX04961c

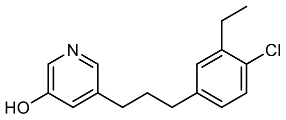


OX04961c

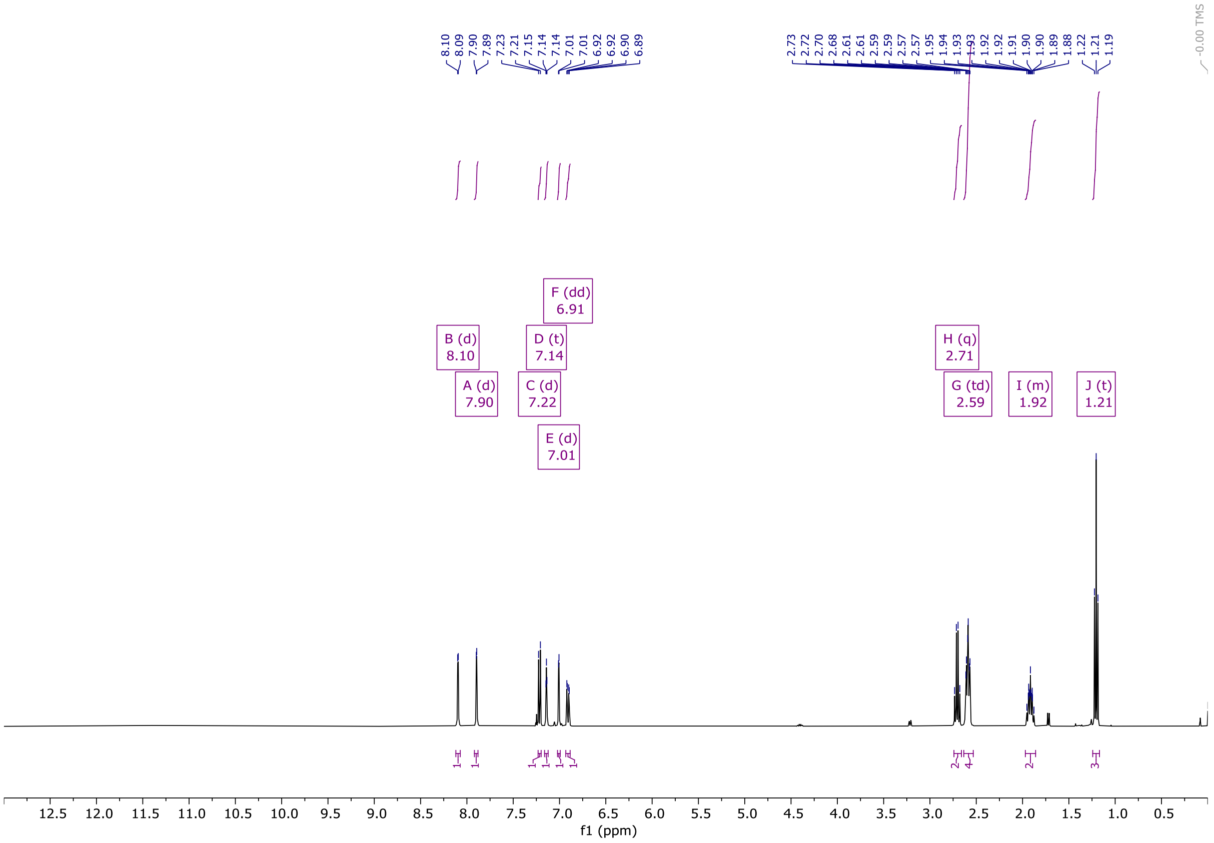


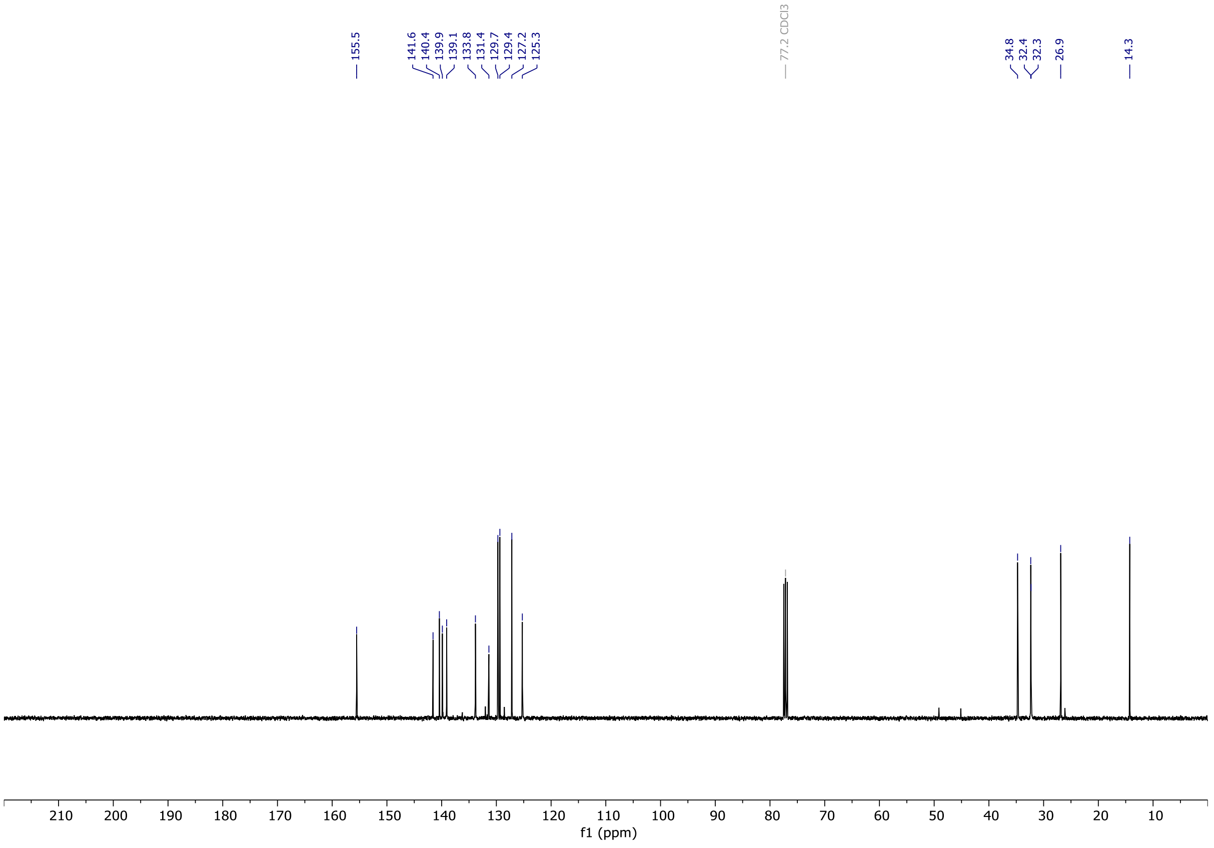

# OX04961

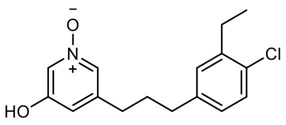

OX04961

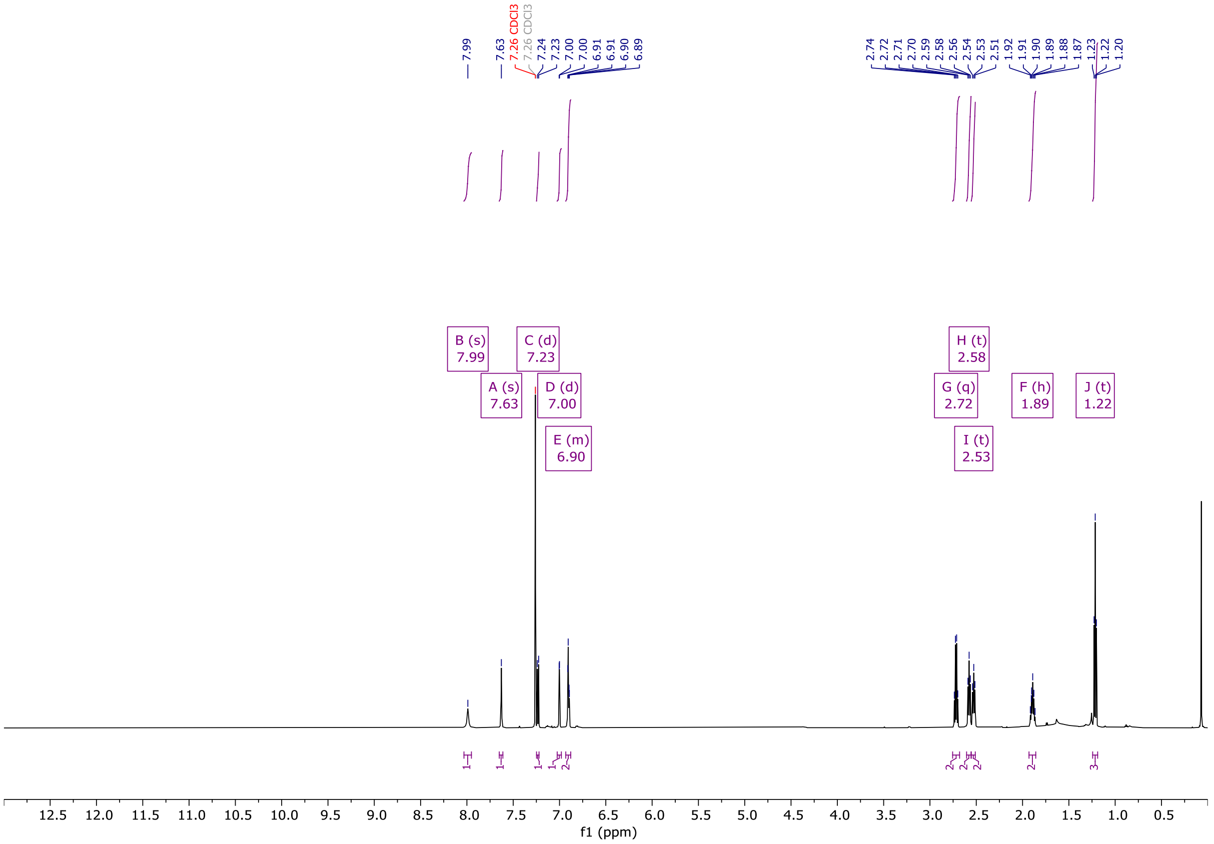


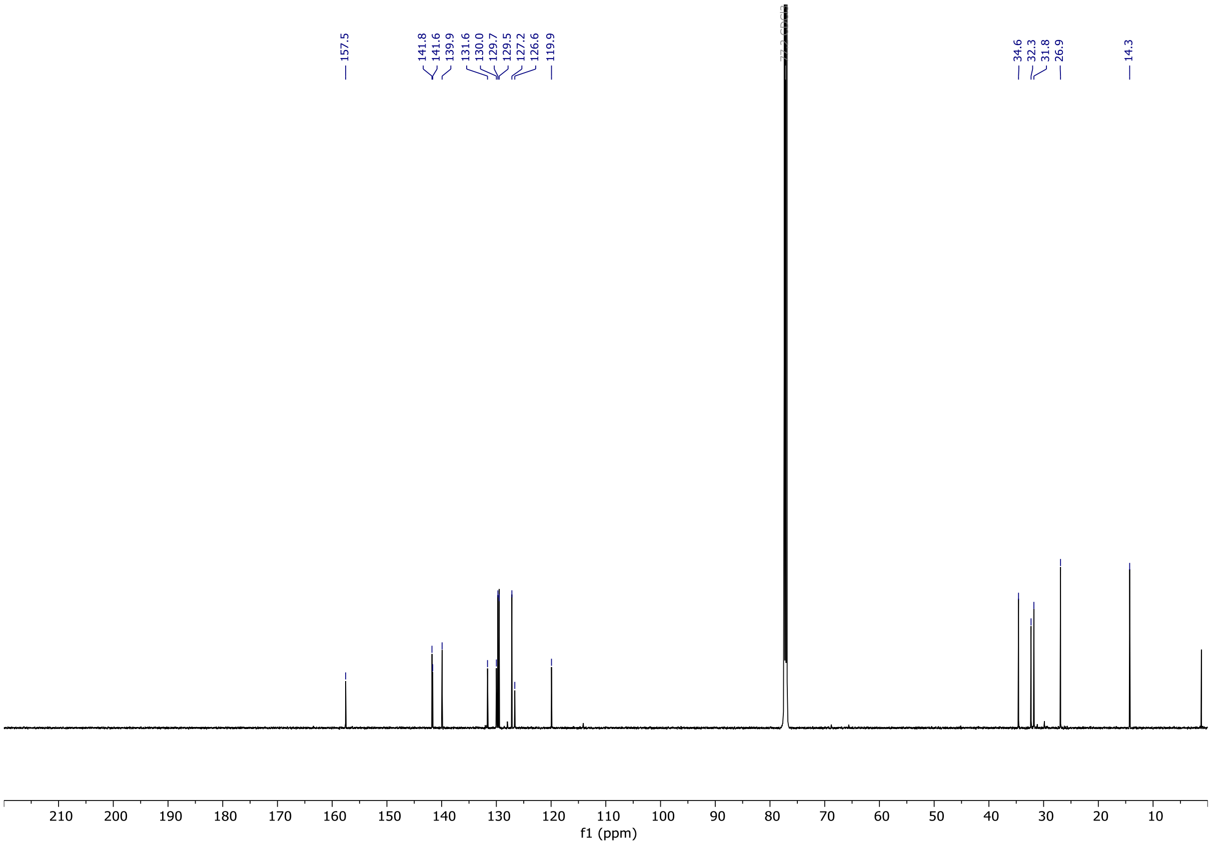

# OX04962a

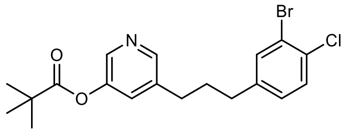


OX04962a

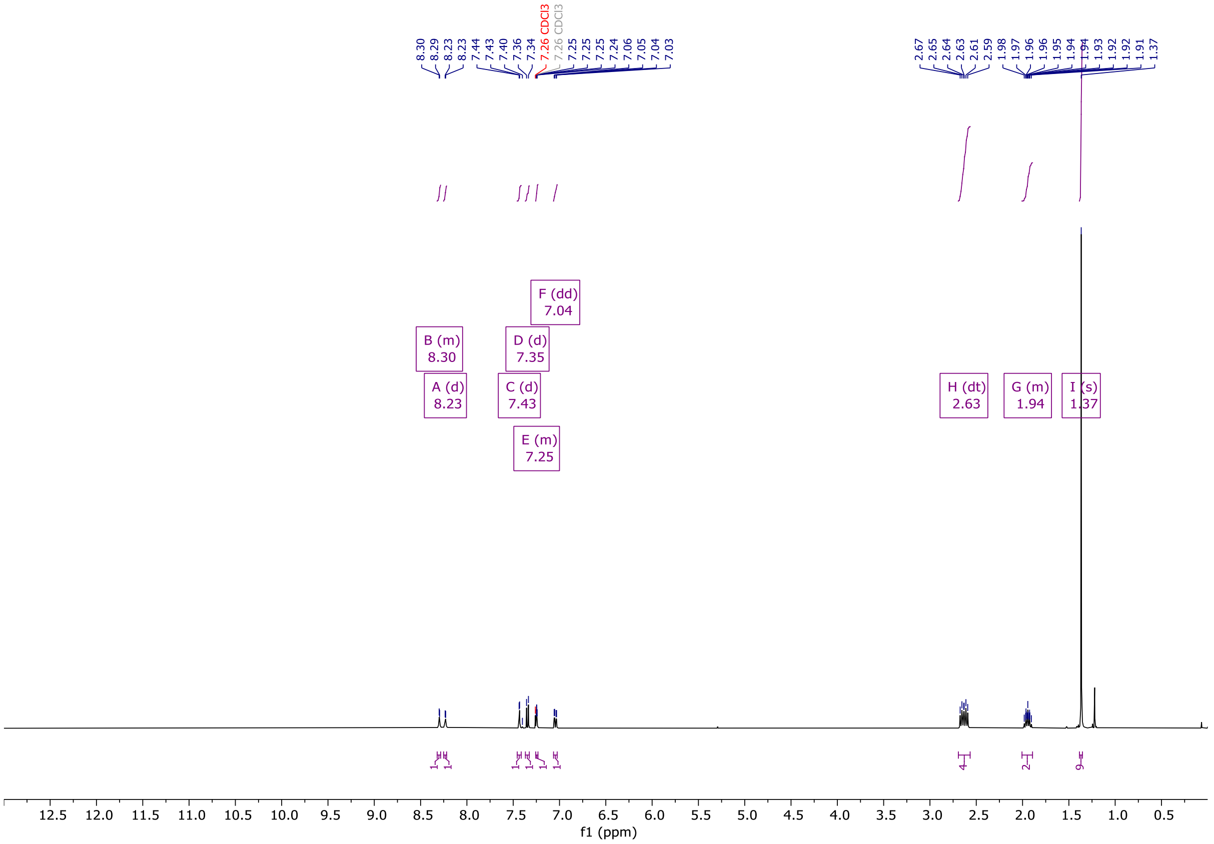


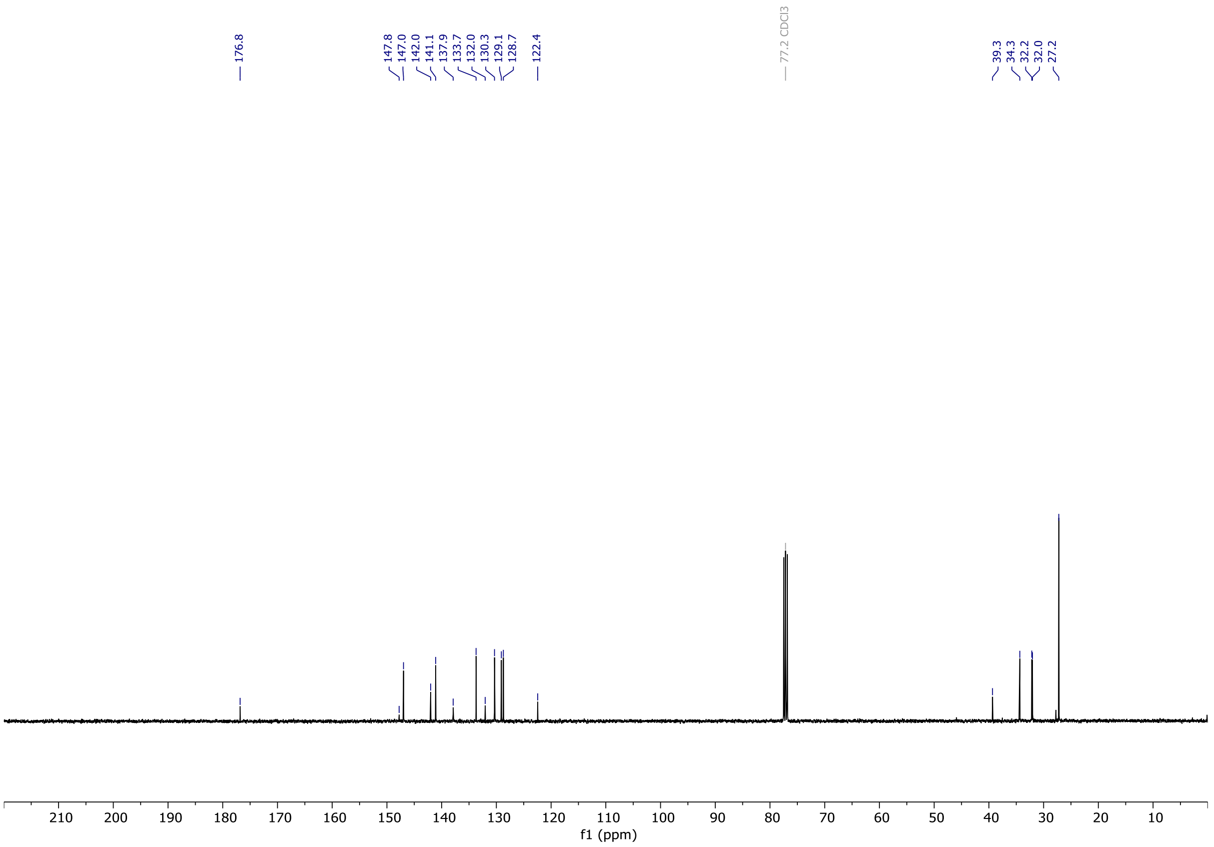

# OX04962b

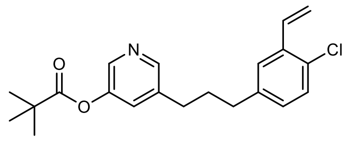

OX04962b

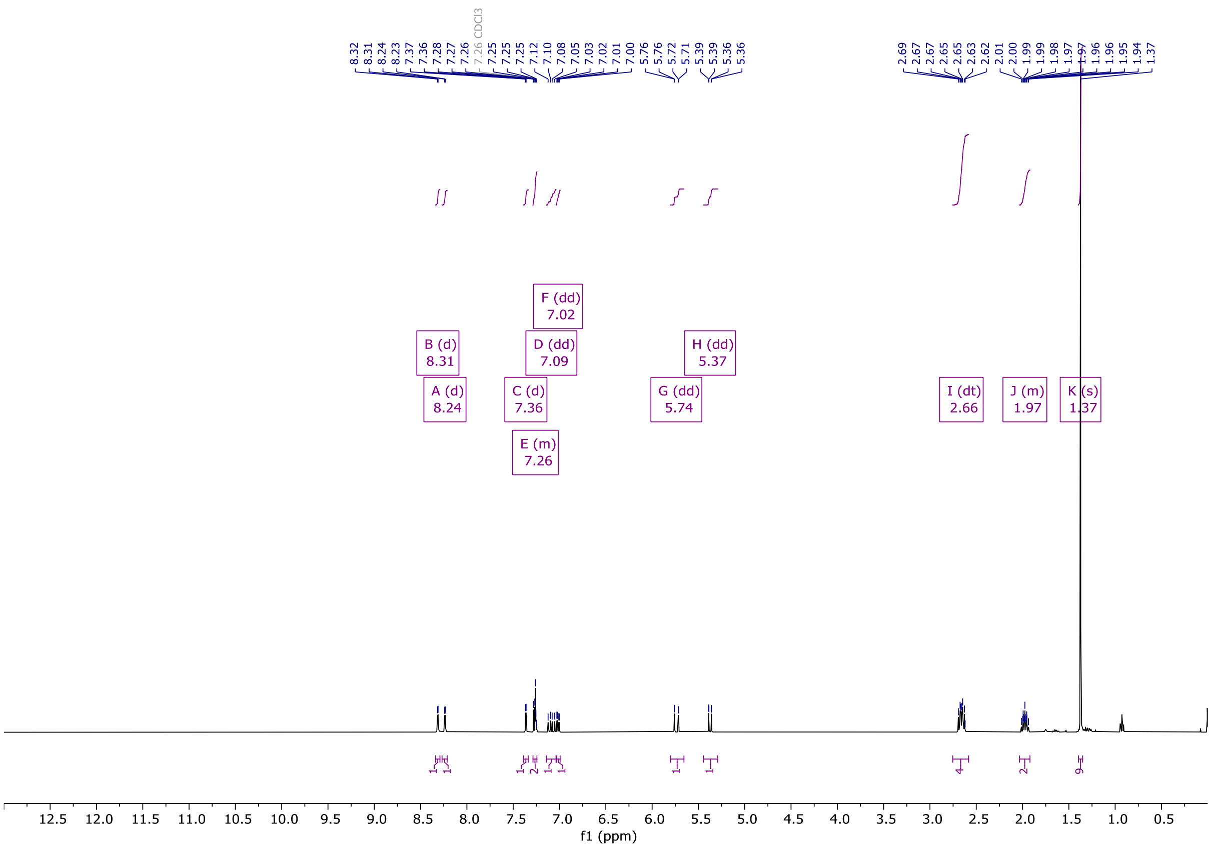


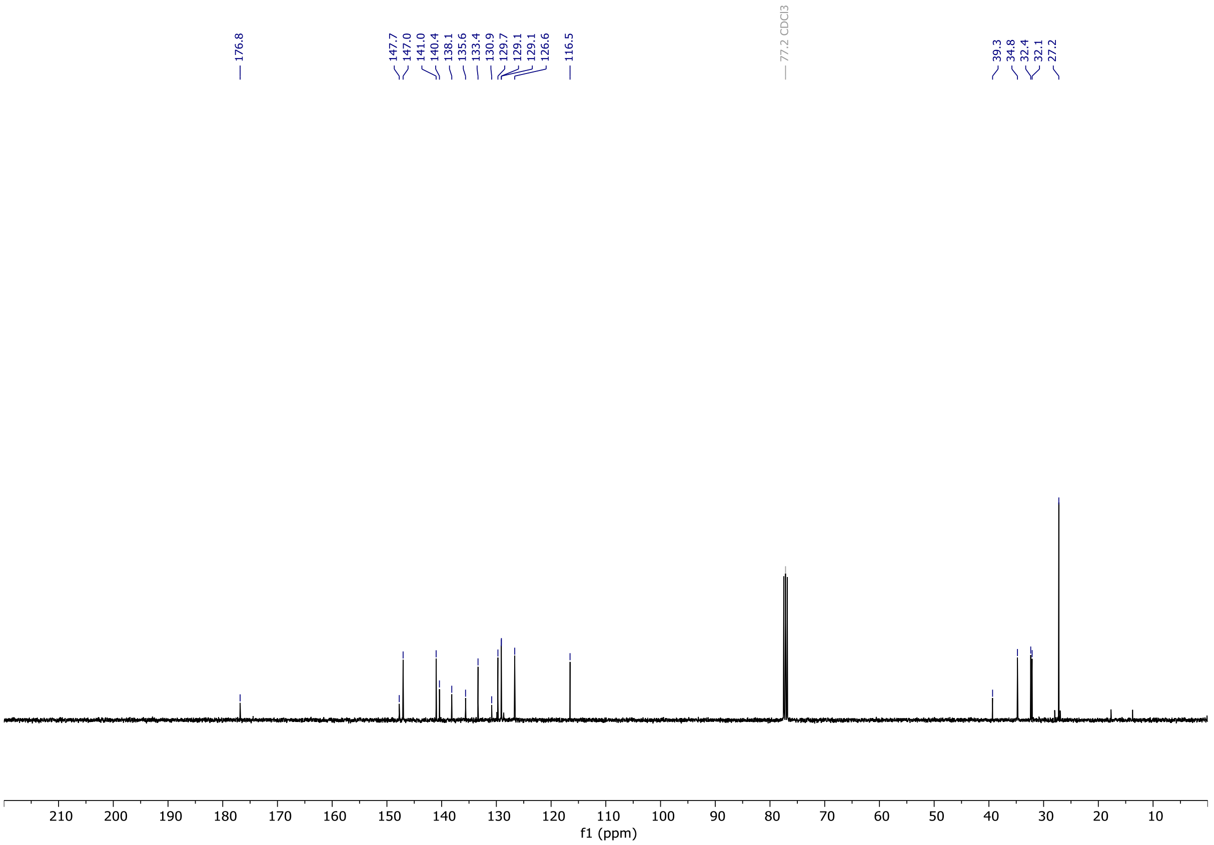

# OX04962

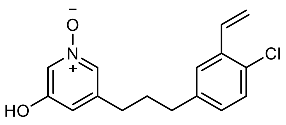

OX04962

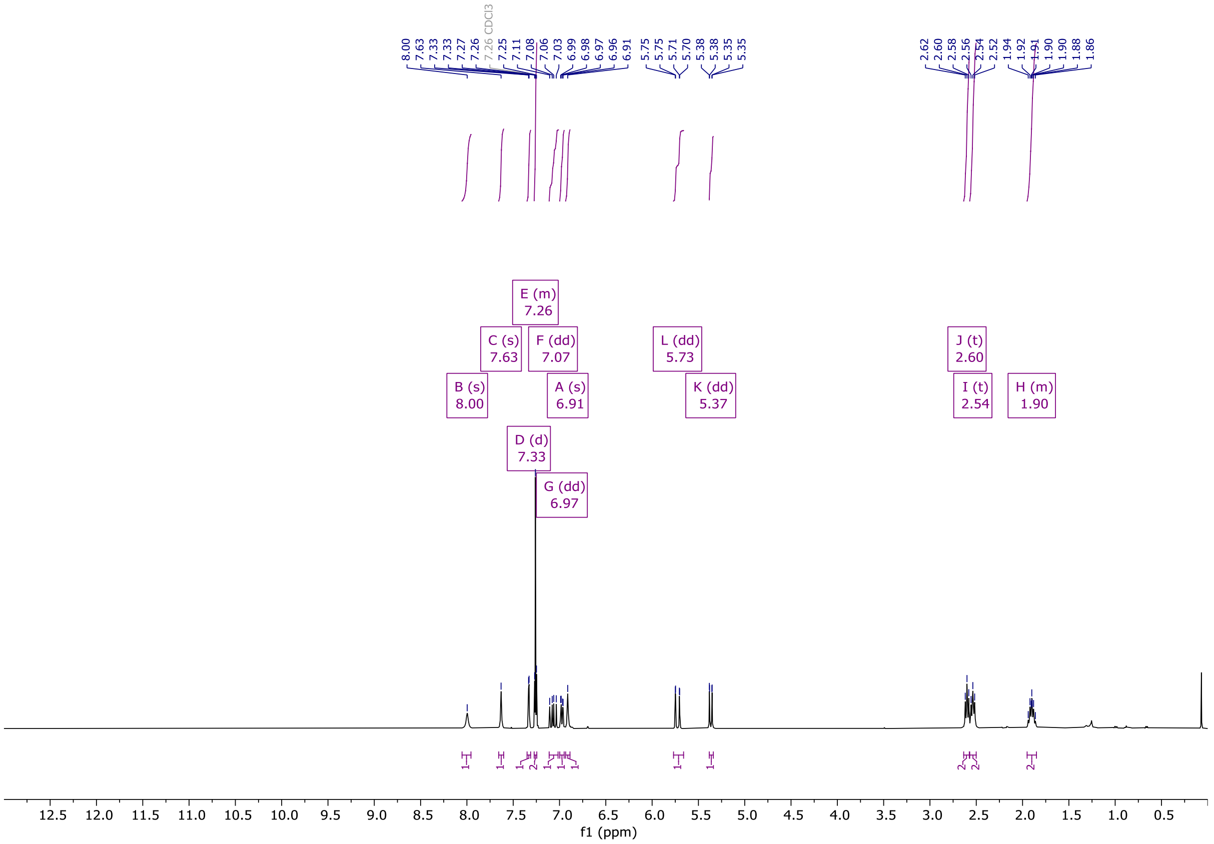

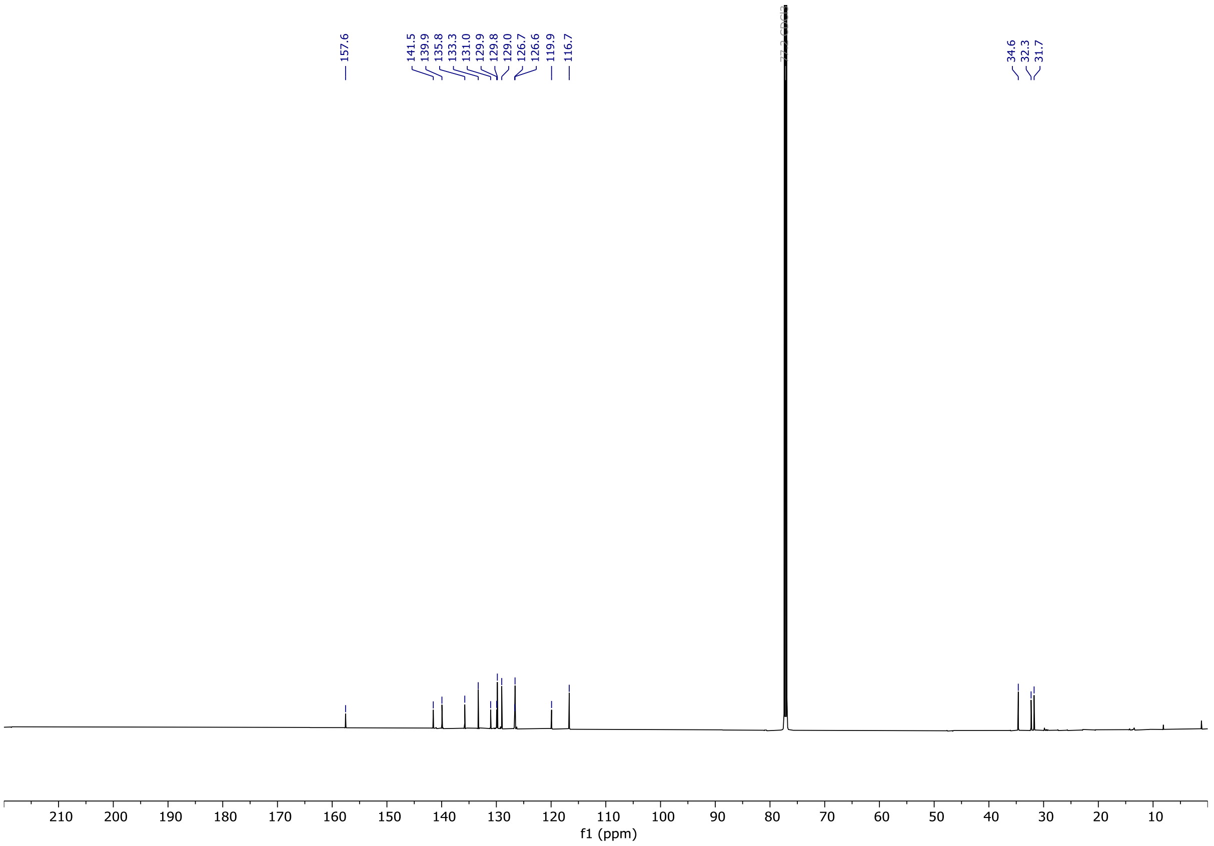

# OX04963a

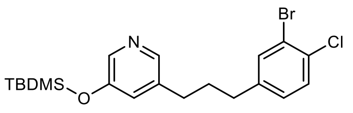

OX04963a

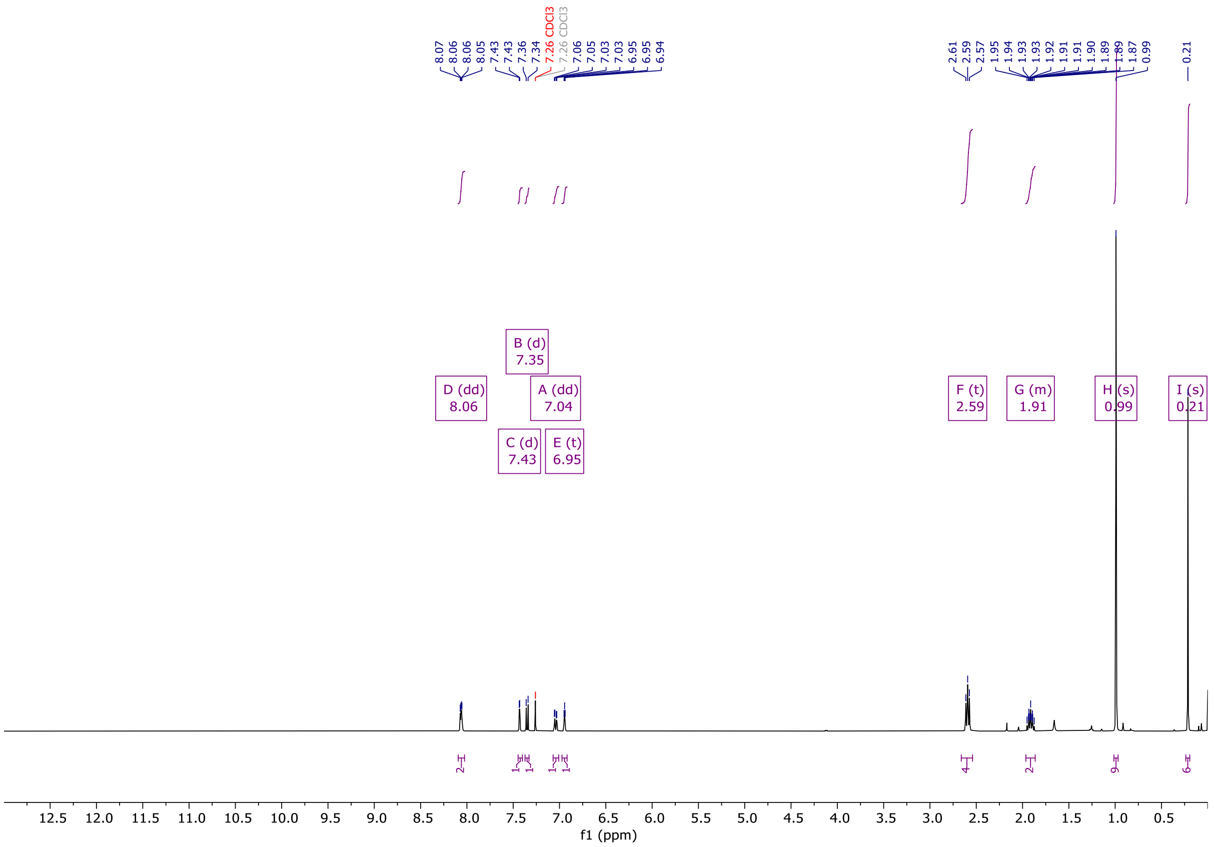

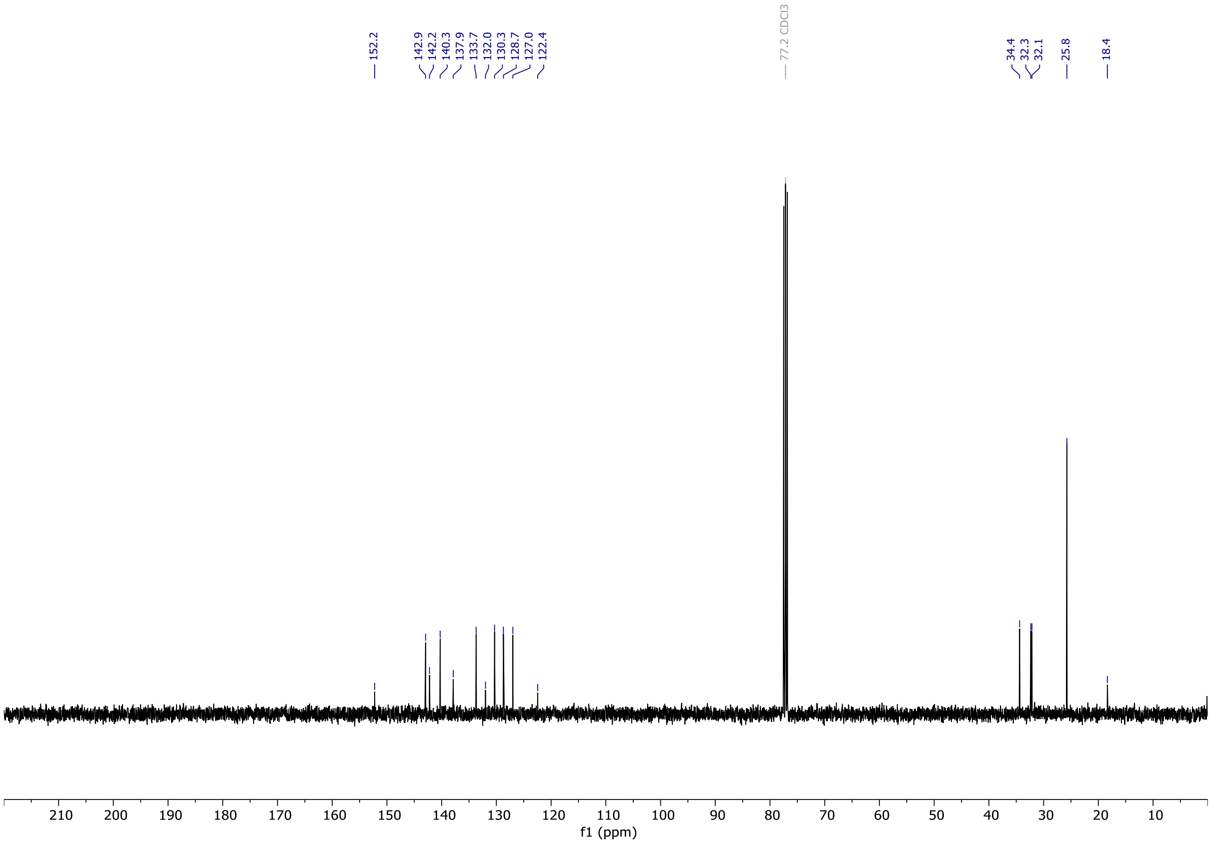

# OX04963b

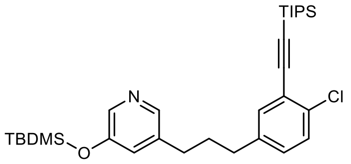


OX04963b

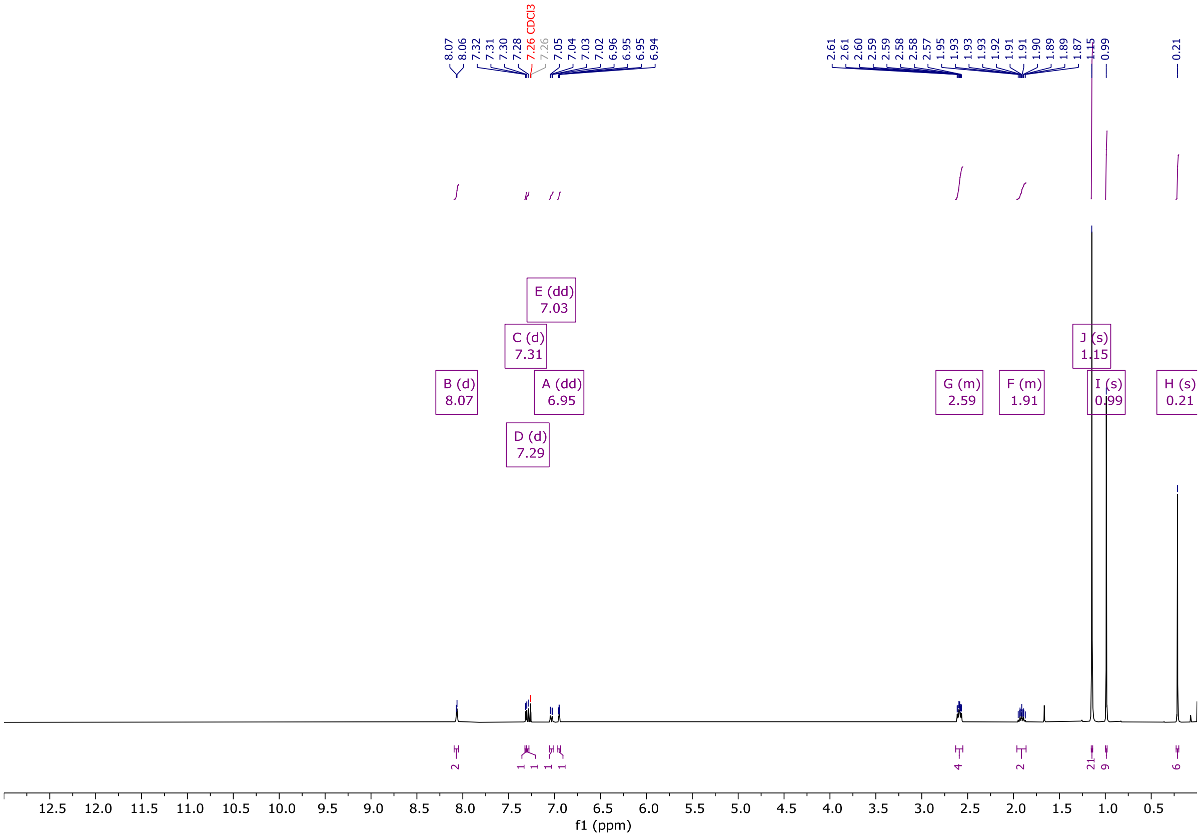


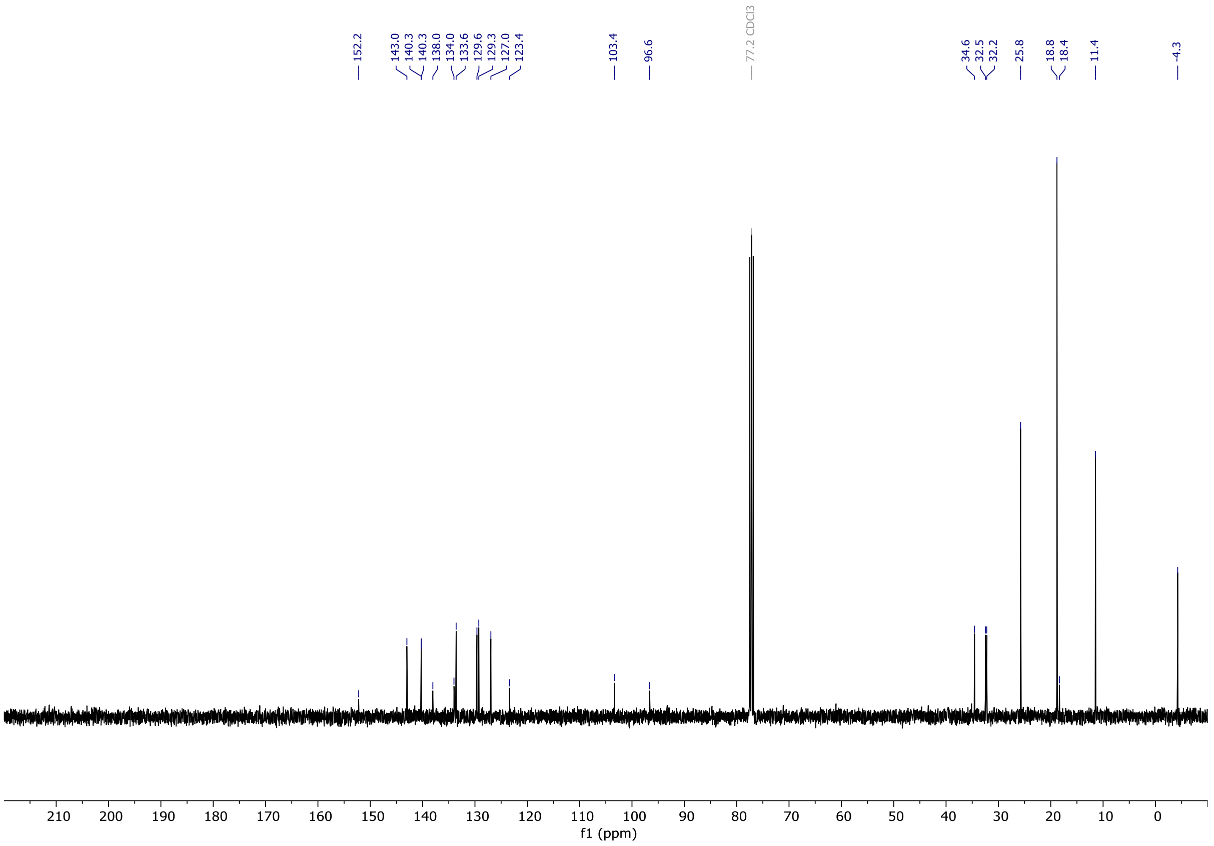

# OX04963

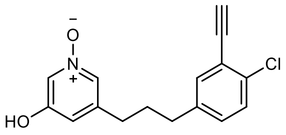


OX04963

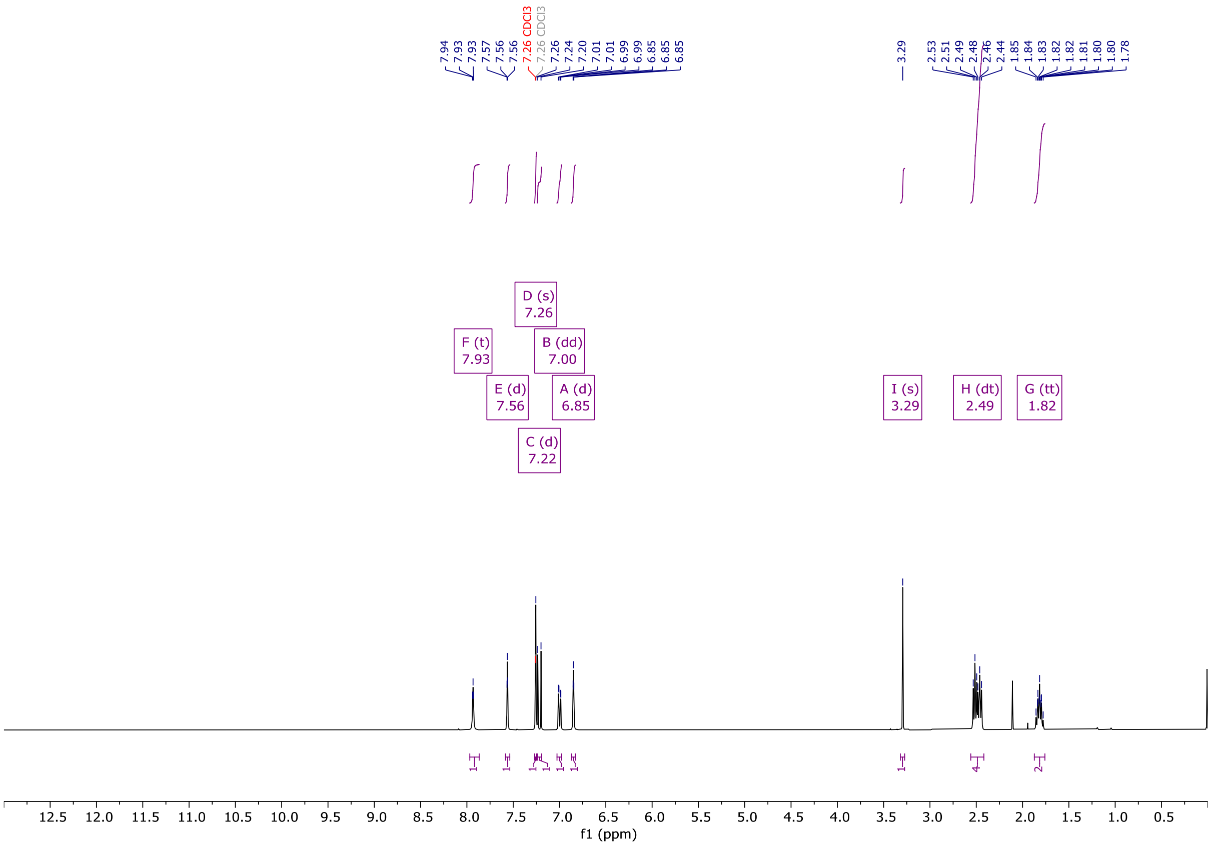


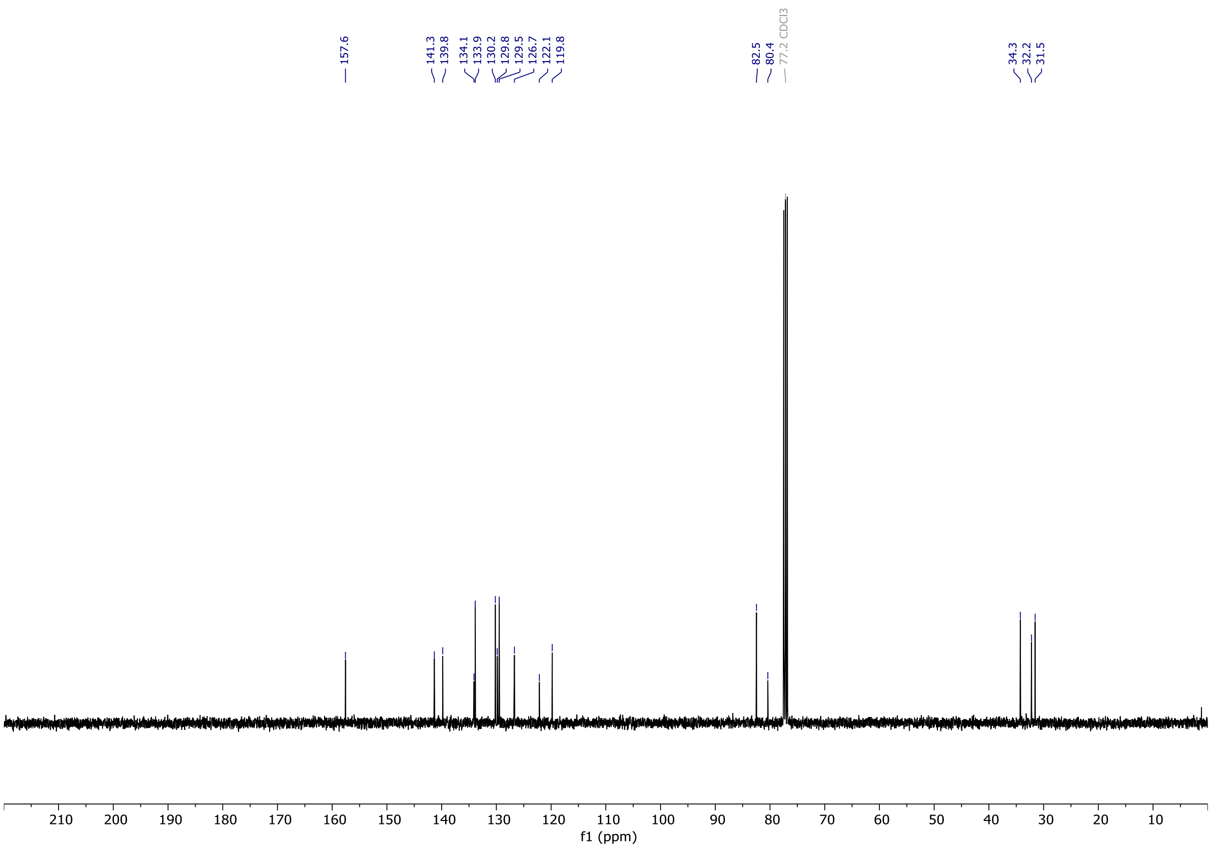

# OX04964a

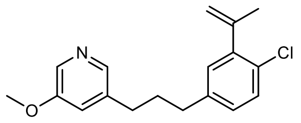

OX04964a

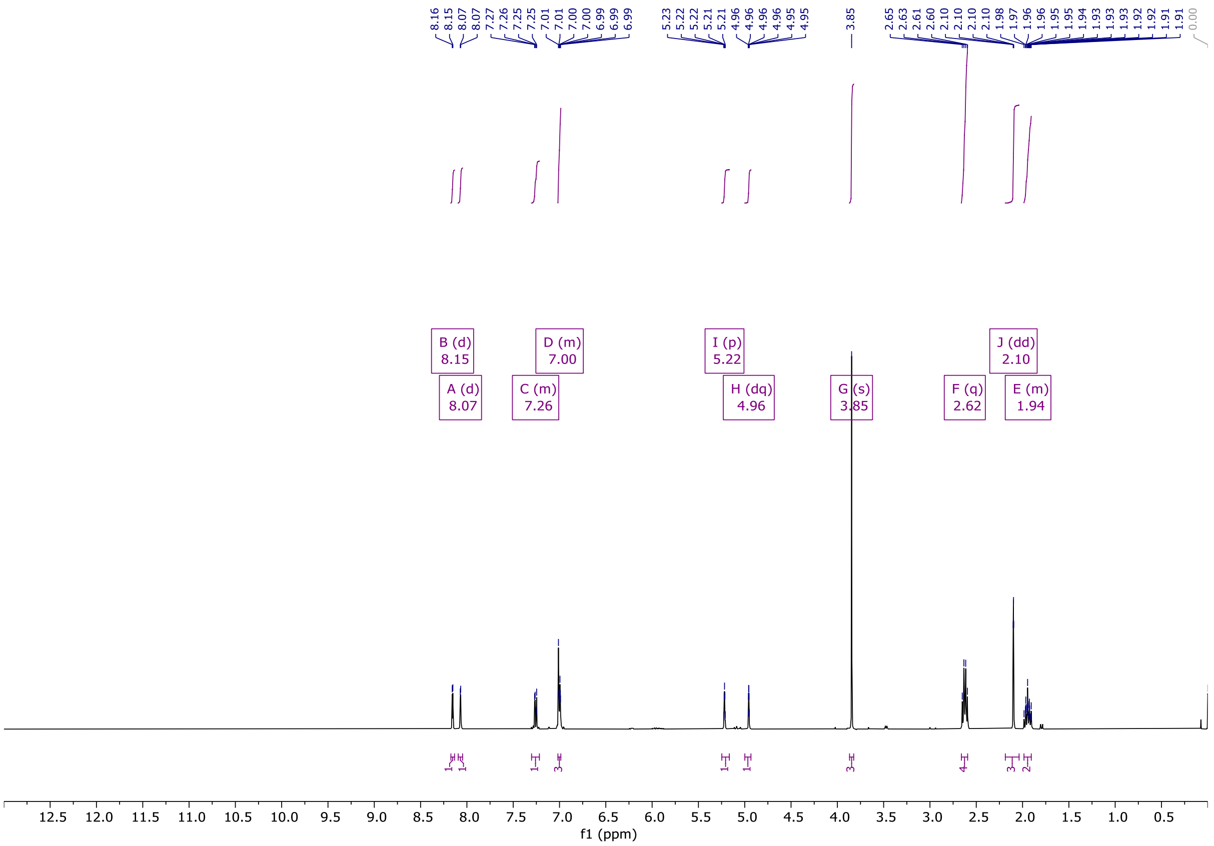

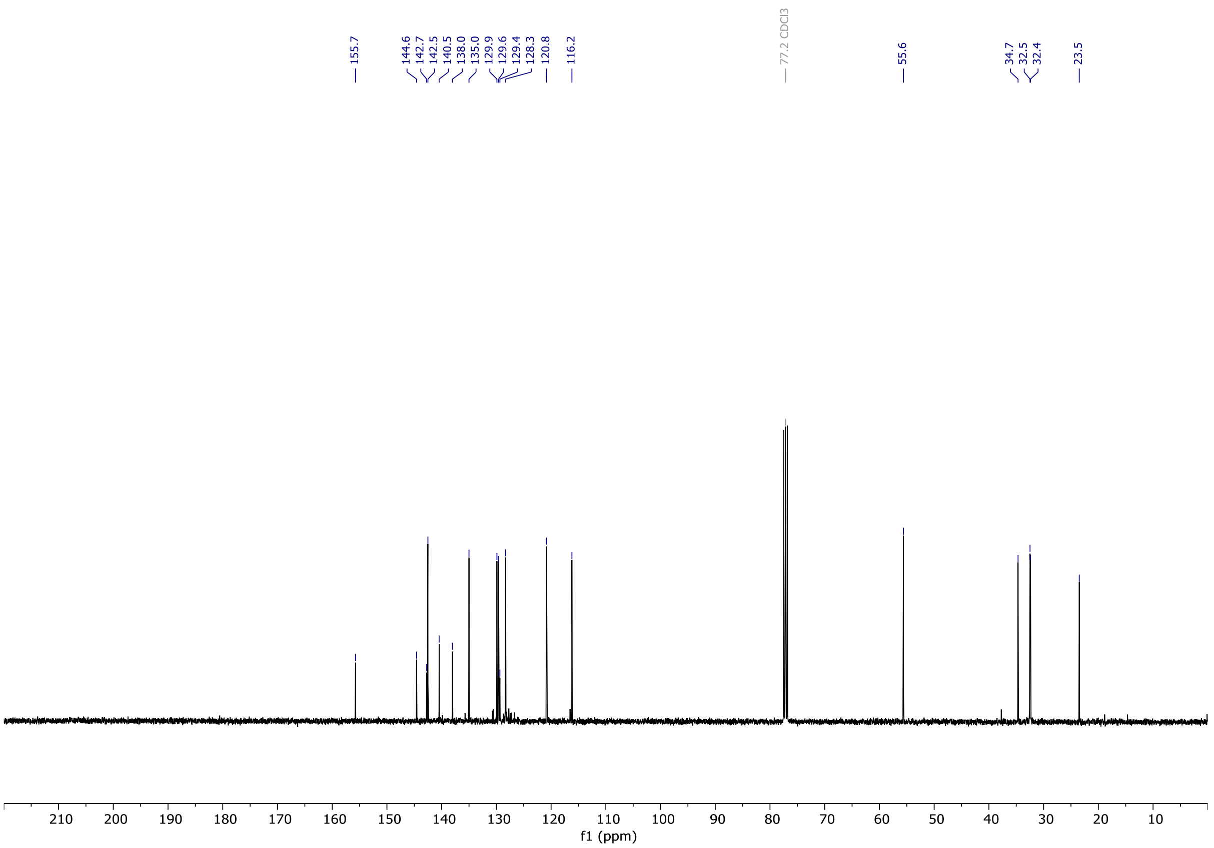

# OX04964b

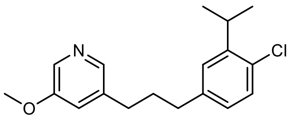


OX04964b

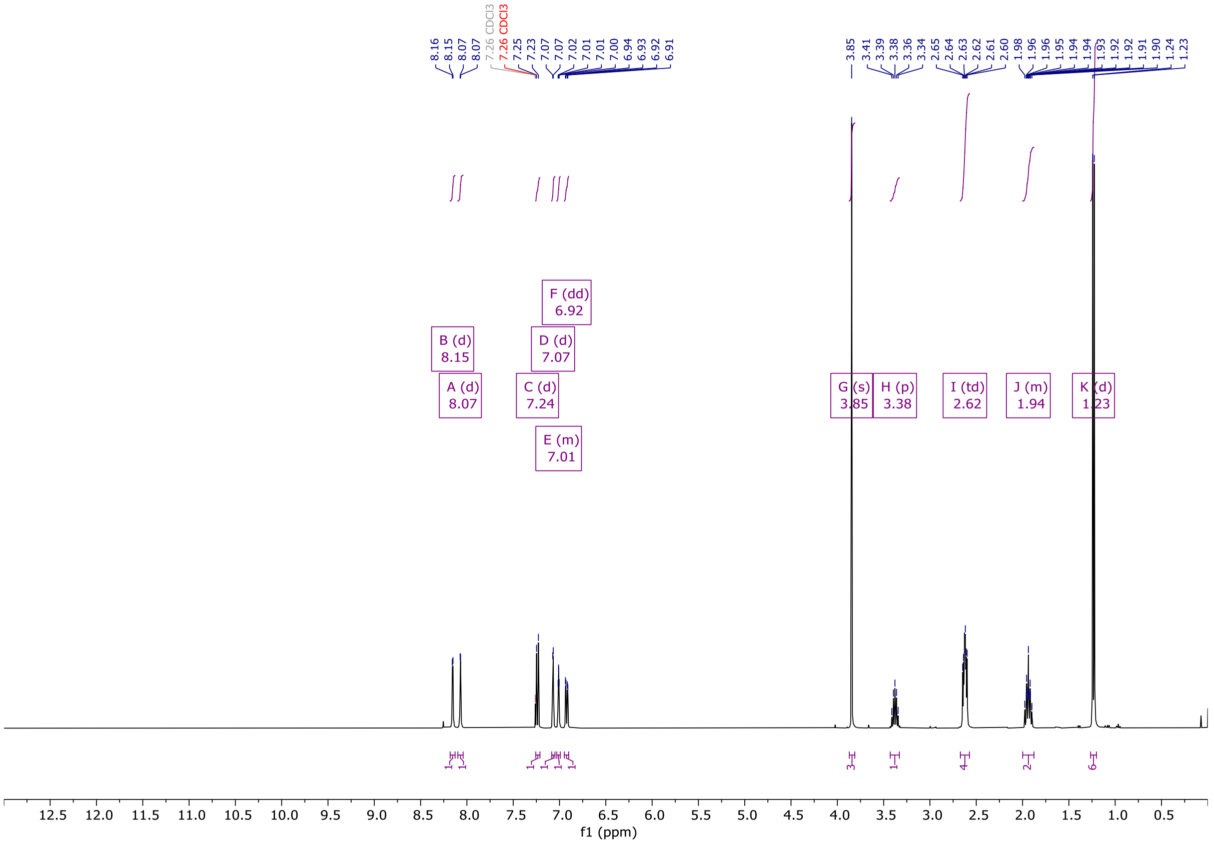


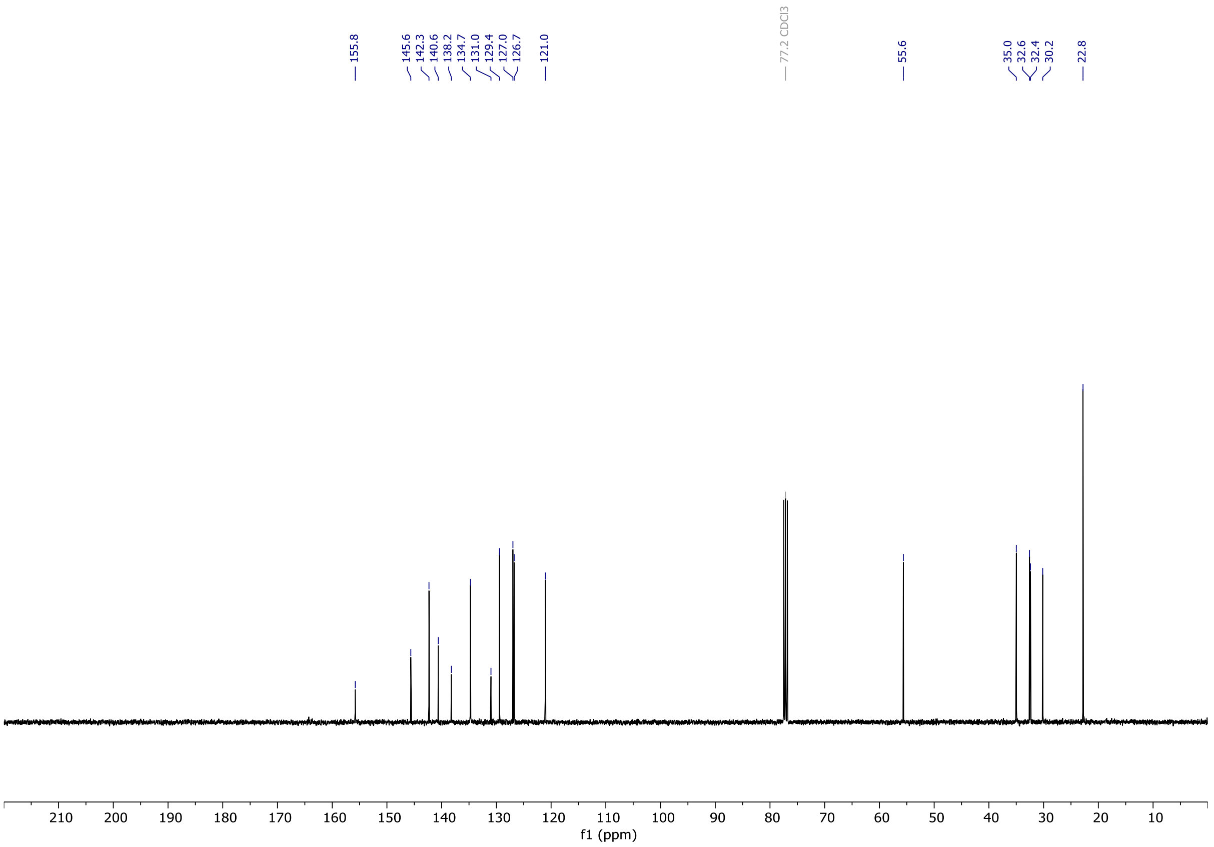

# OX04964c

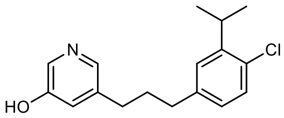


OX04964c

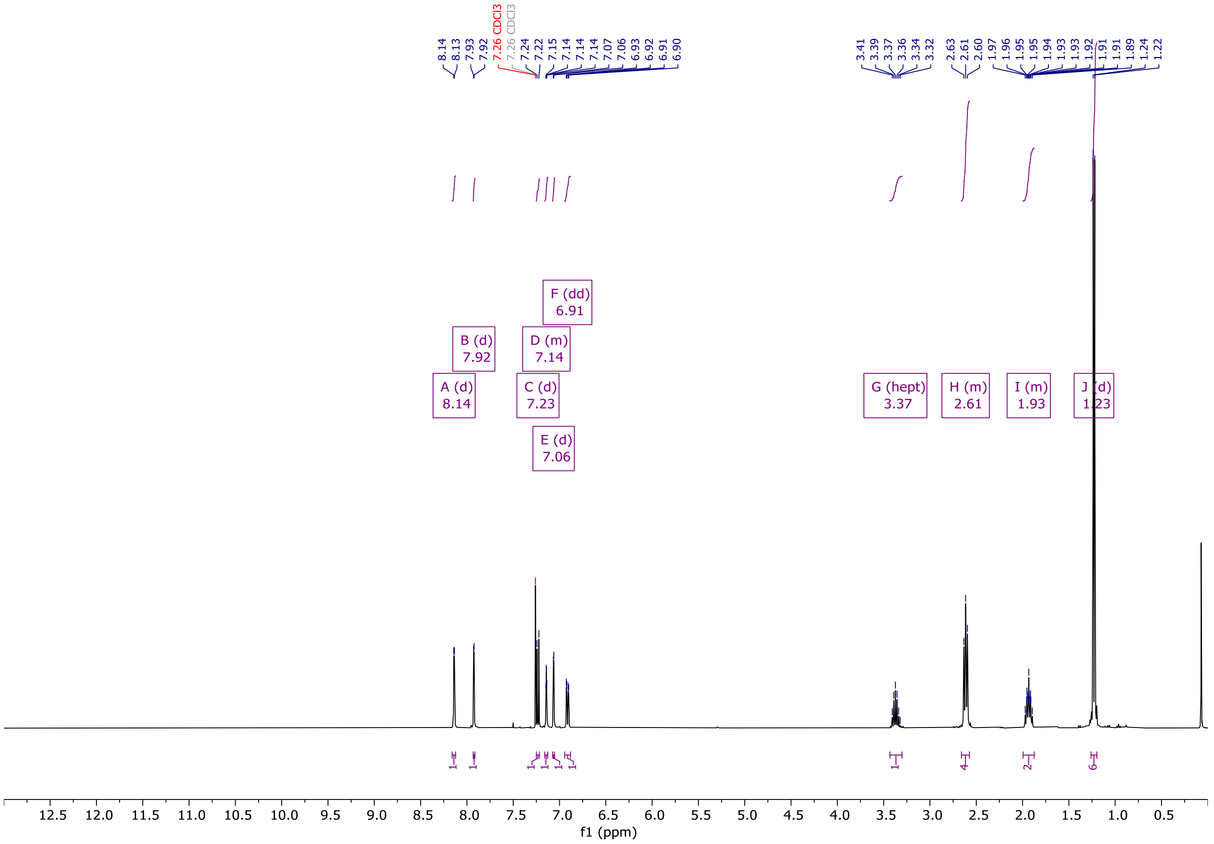

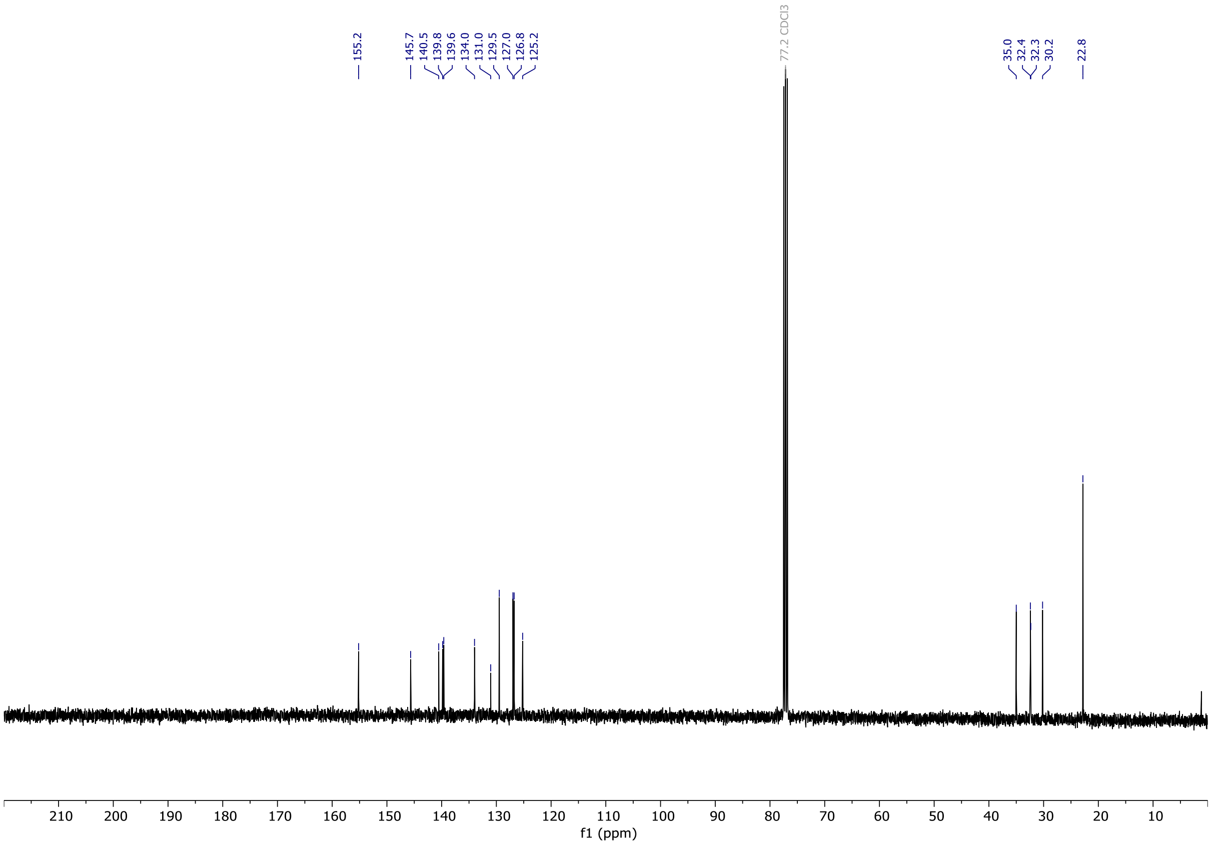

# OX04964

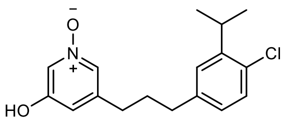


OX04964

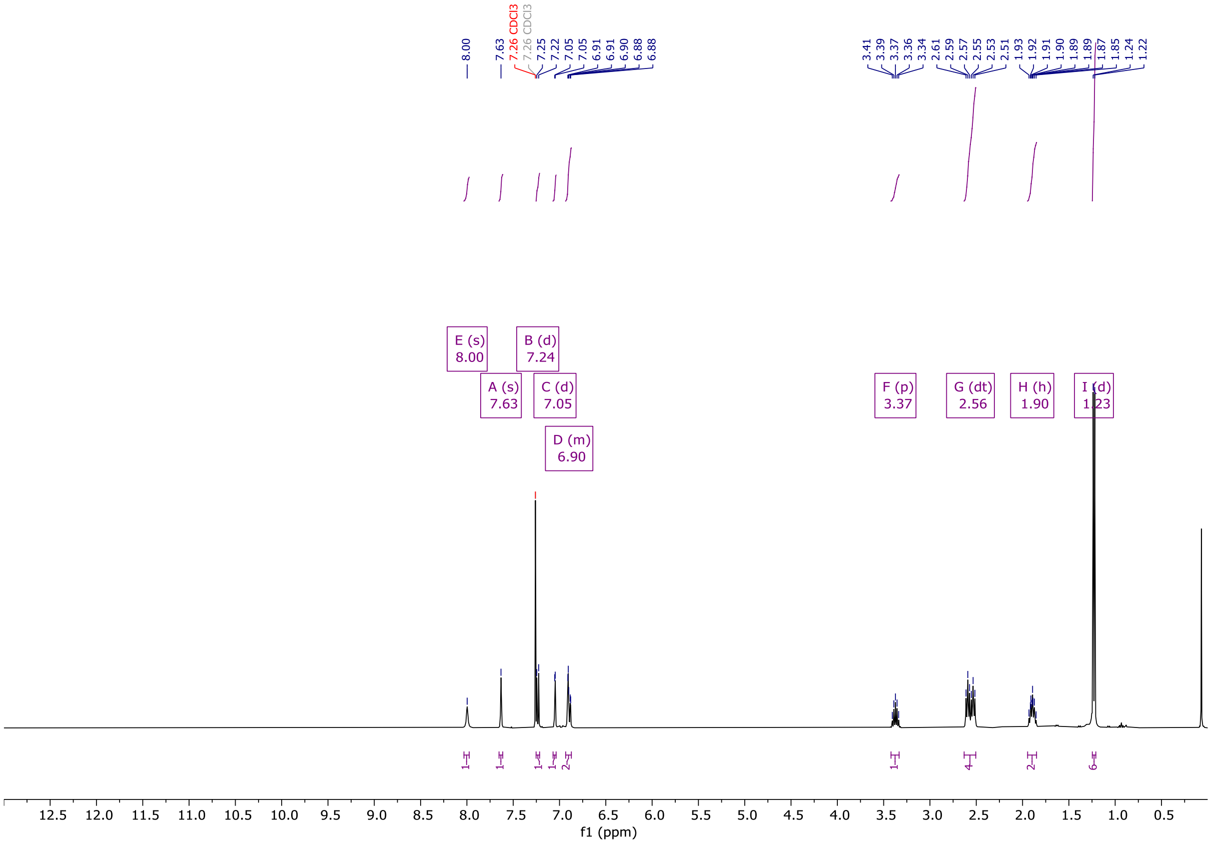


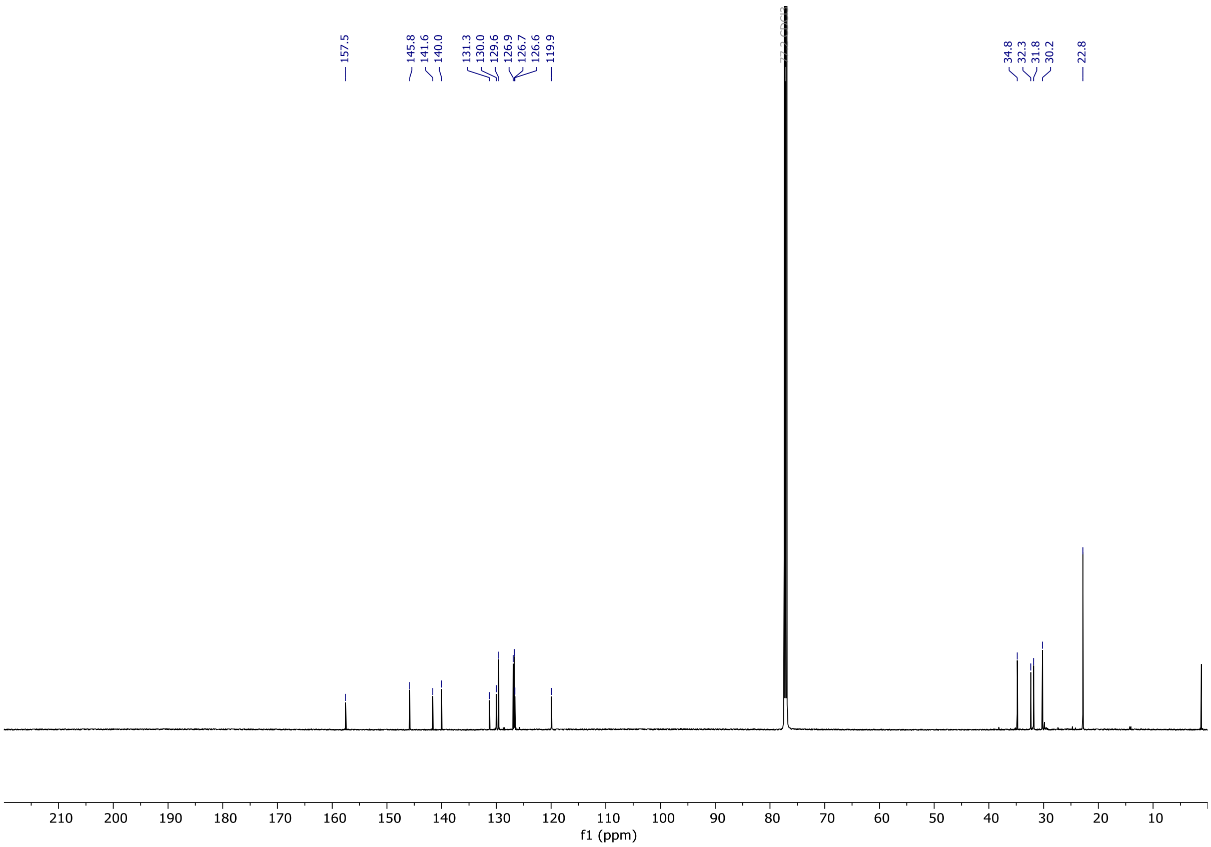

# OX04965a

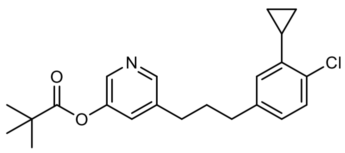

OX04965a

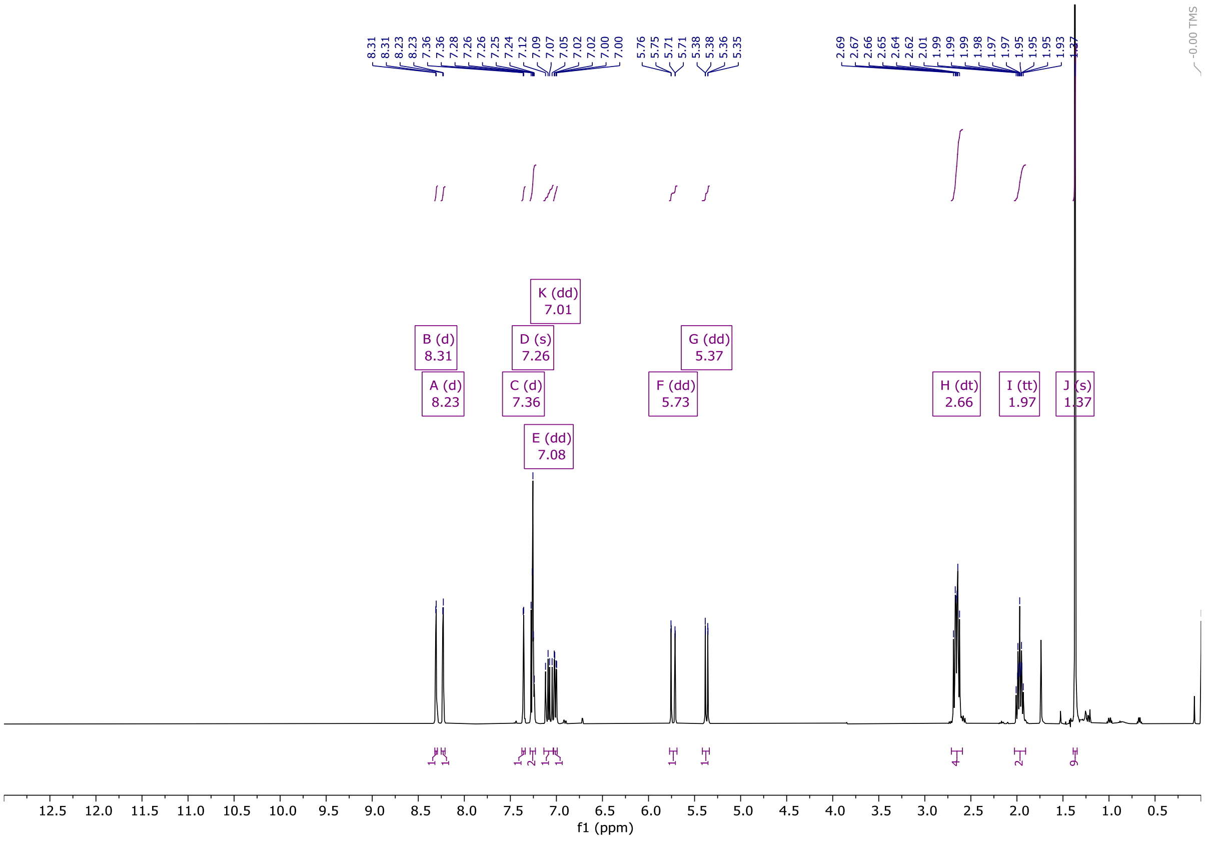

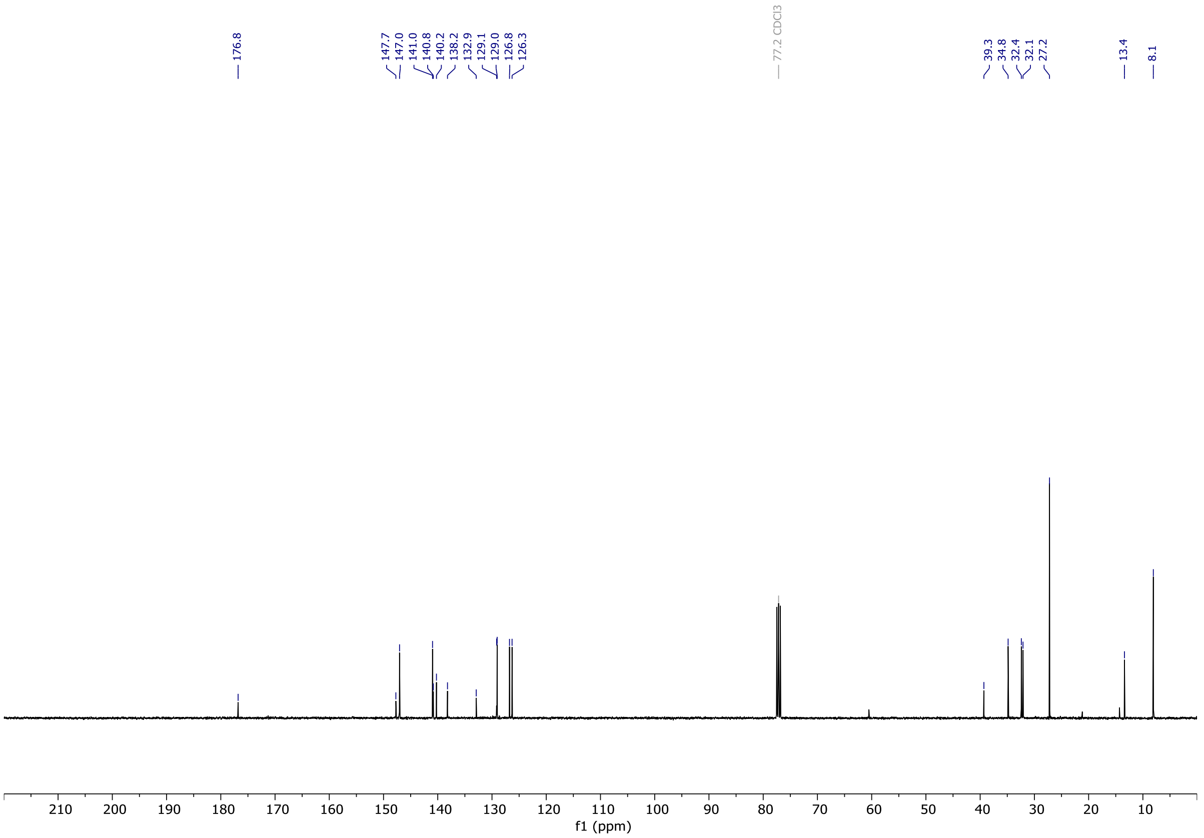

# OX04965

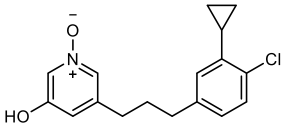


OX04965

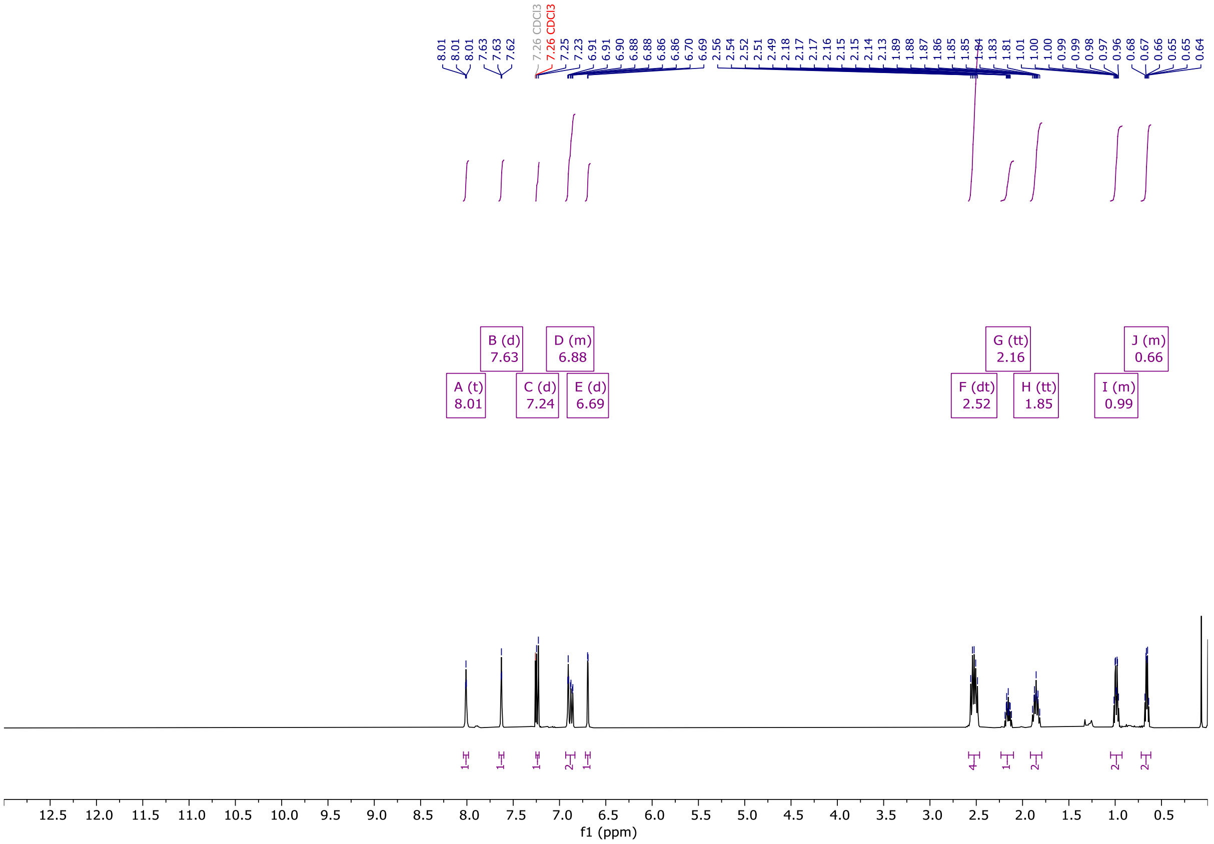


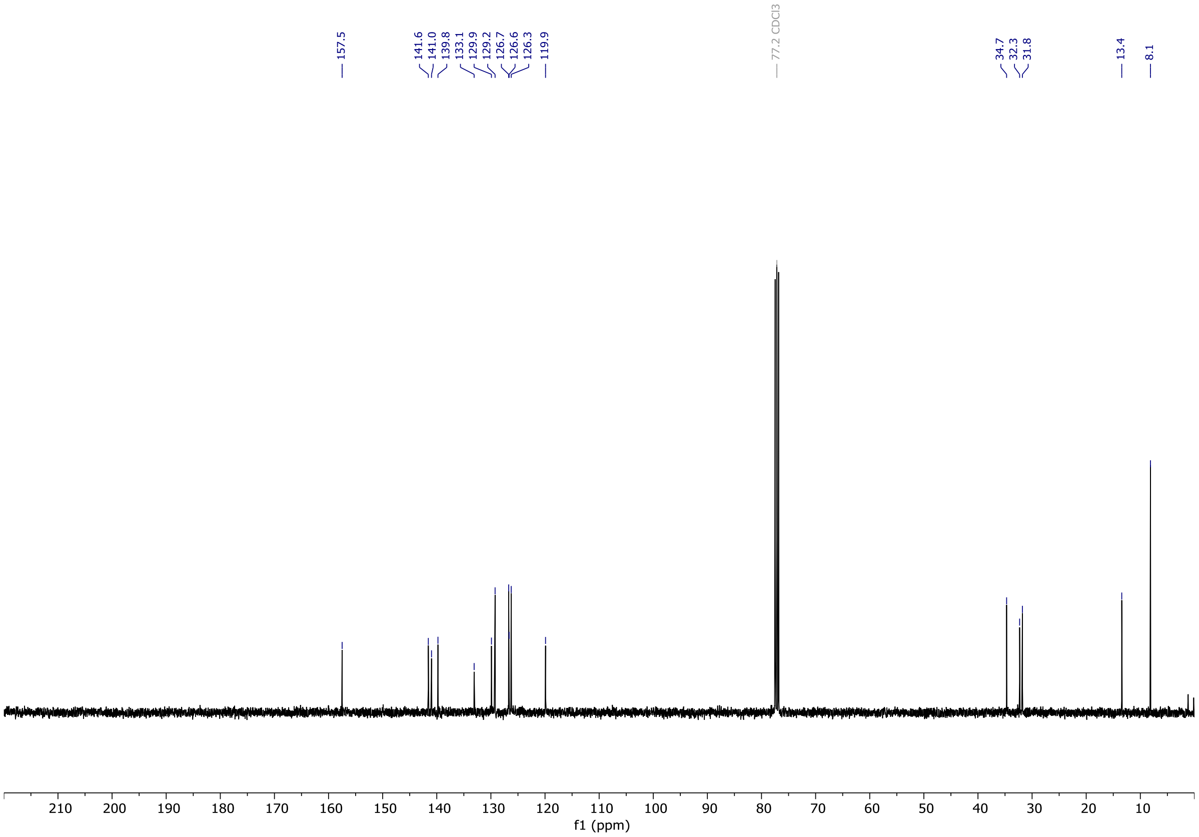

# OX04966a

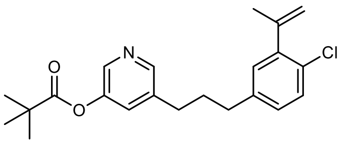


OX04966a

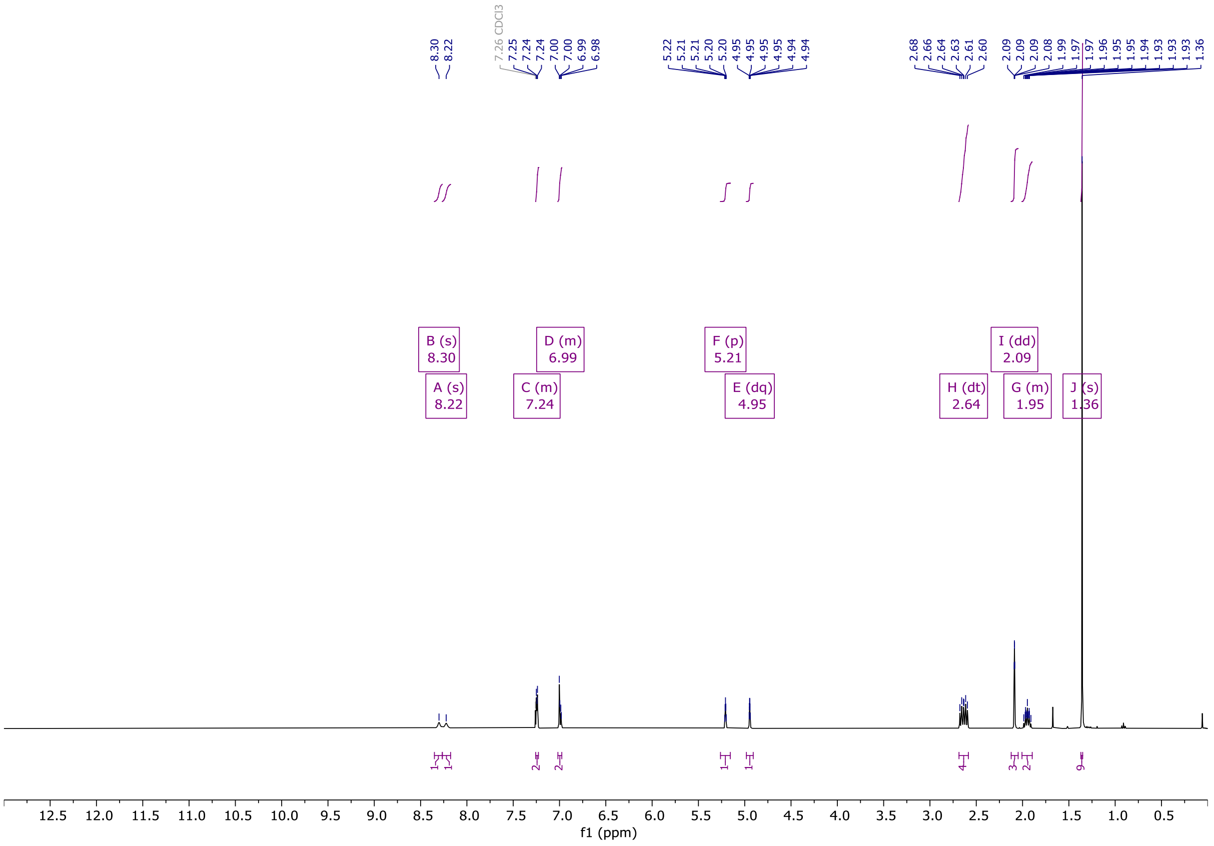


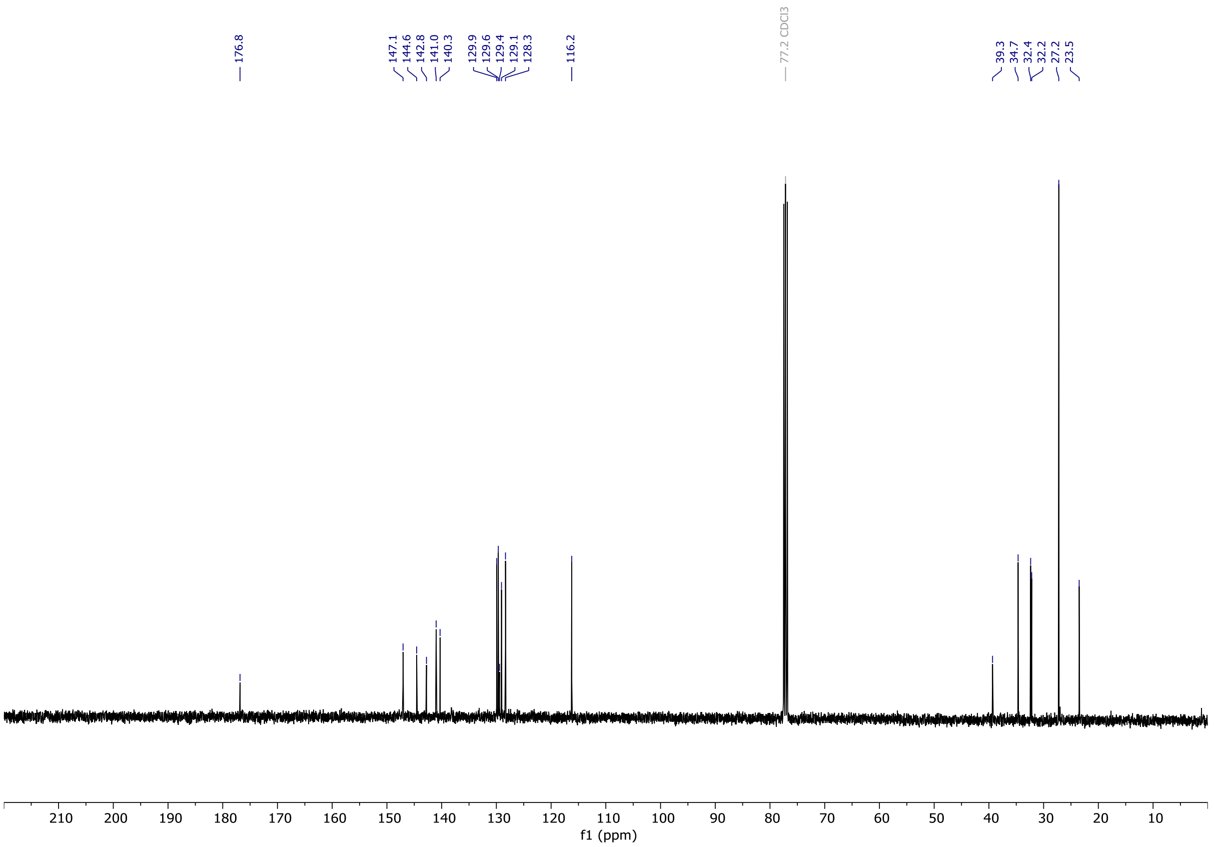

# OX04966b

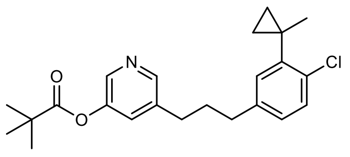


OX04966b

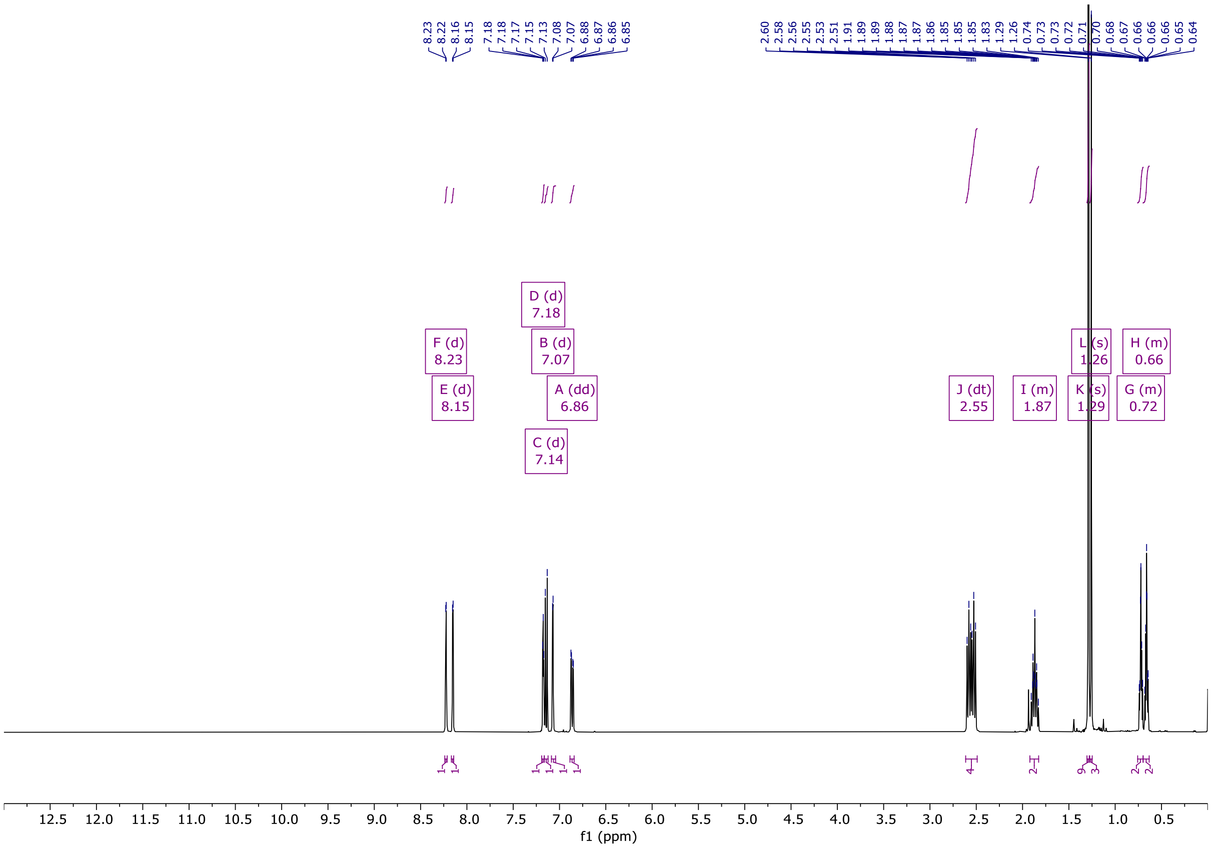


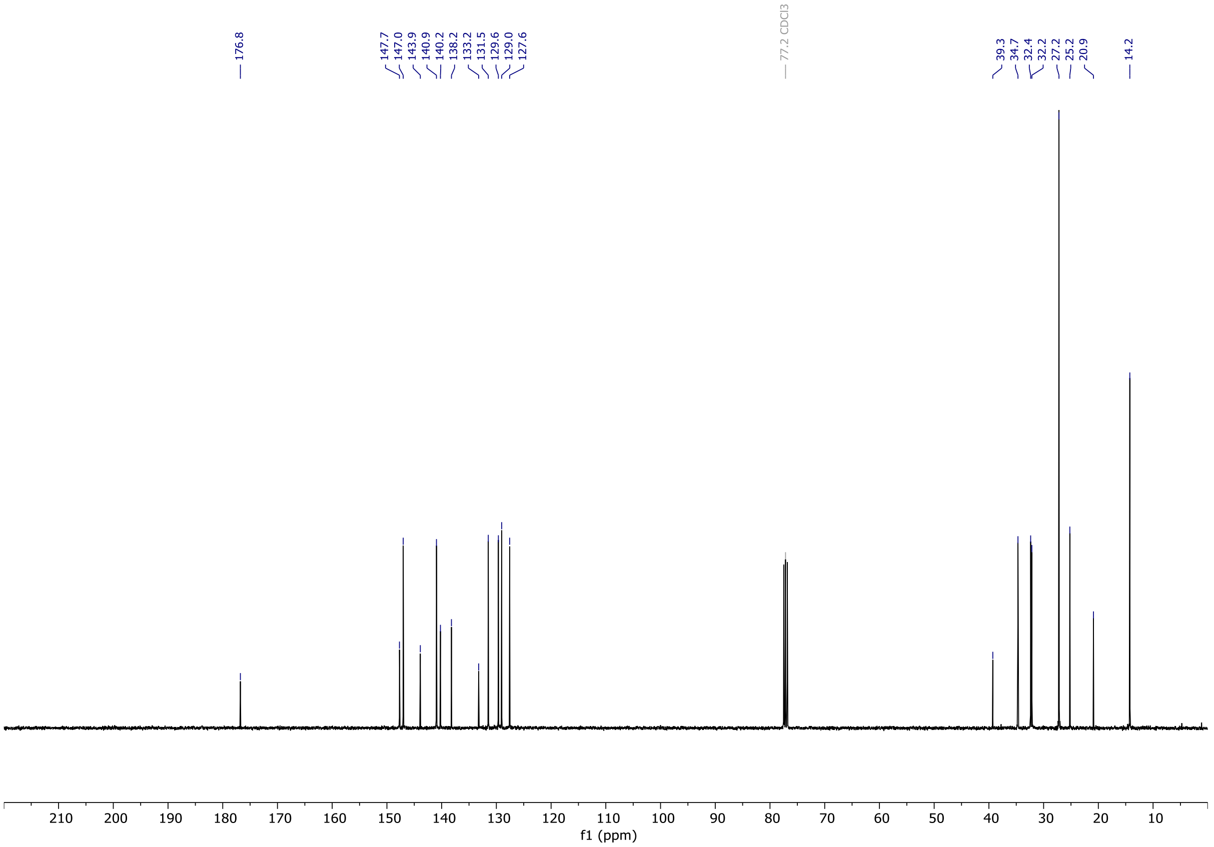

# OX04966

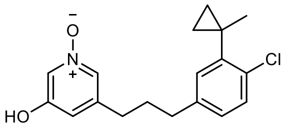

OX04966

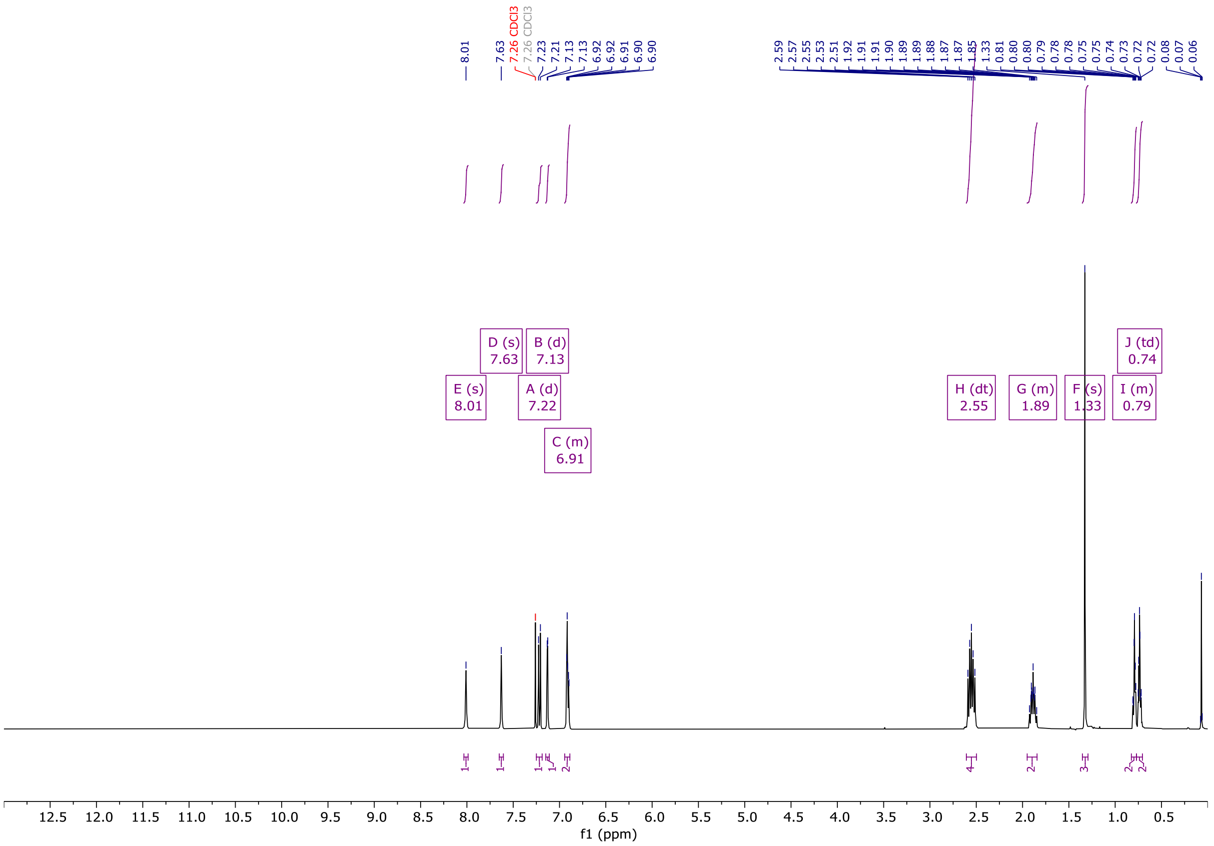

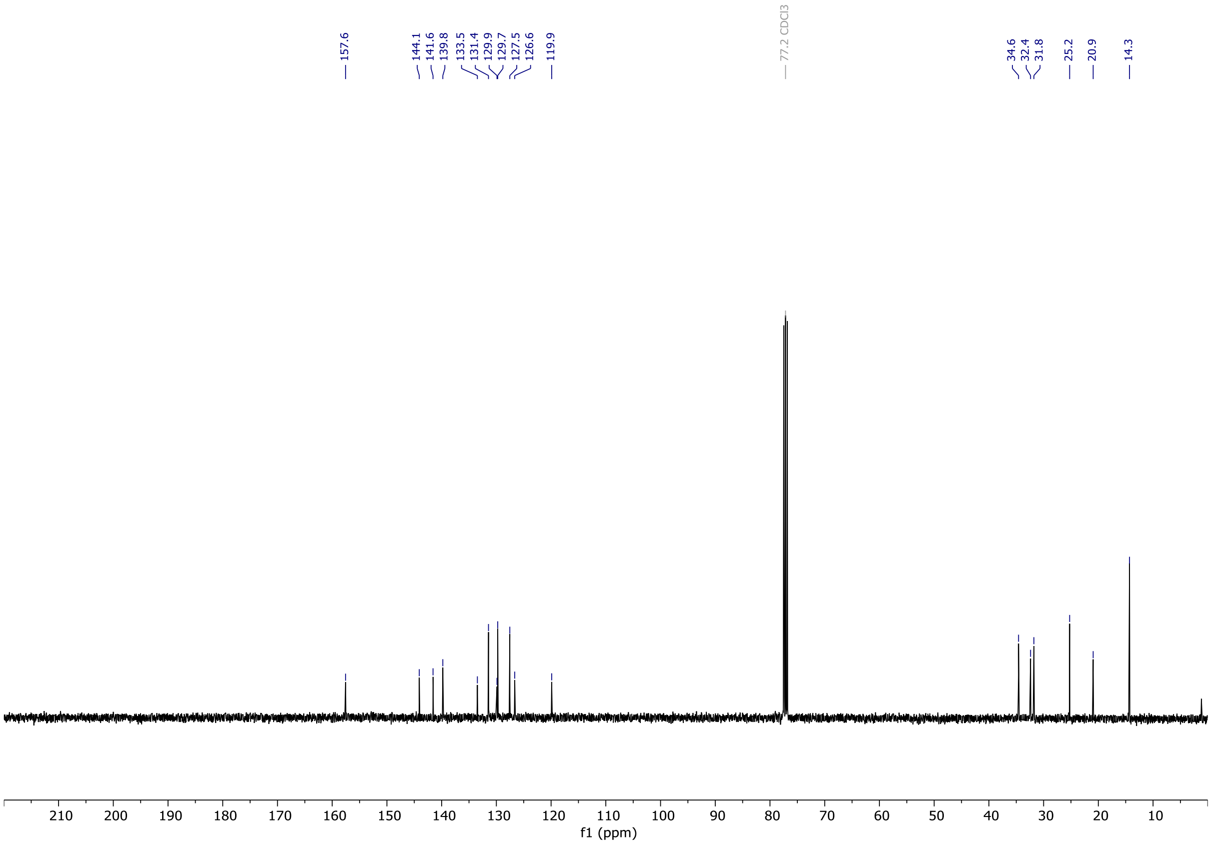

**HPLC**

**OX04539**

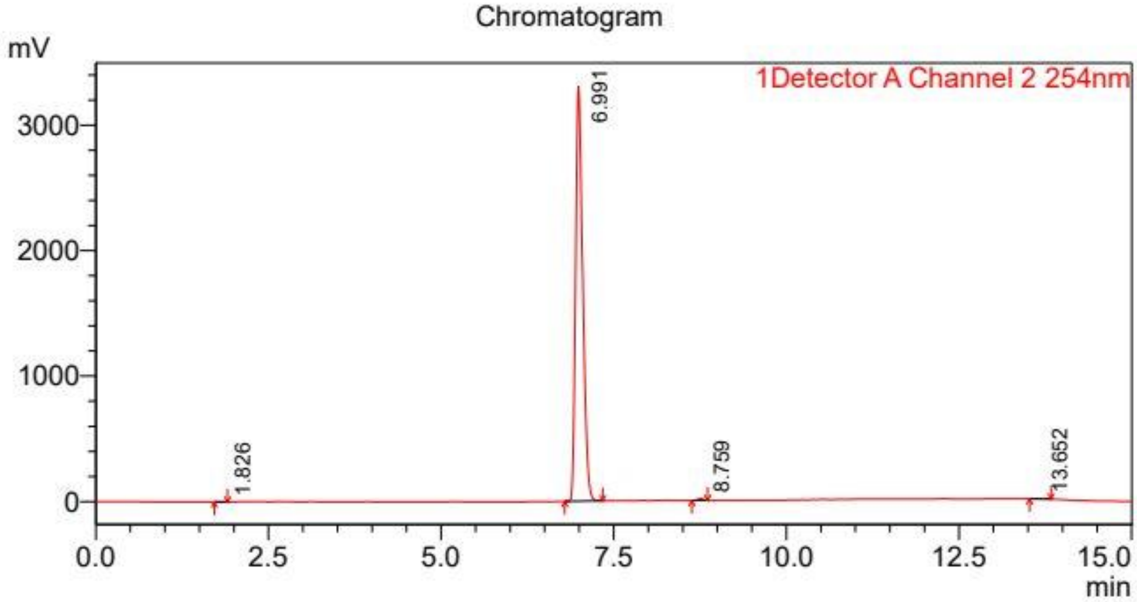


Detector A Channel 2 254nm

| Peak# | Ret. Time (min) | Area% |
|---|---|---|
| 1 | 1.826 | 0.096 |
| 2 | 6.991 | 99.377 |
| 3 | 8.759 | 0.384 |
| 4 | 13.652 | 0.143 |
| Total | | 100.000 |

**OX04540**

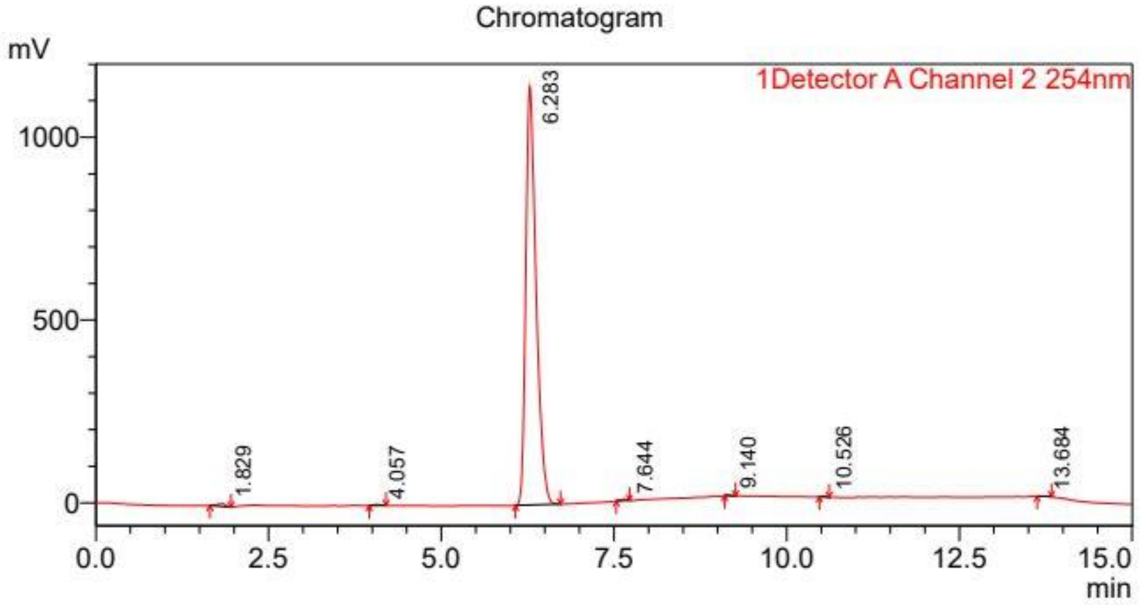


Detector A Channel 2 254nm

| Peak# | Ret. Time (min) | Area% |
|---|---|---|
| 1 | 1.829 | 0.506 |
| 2 | 4.057 | 0.189 |
| 3 | 6.283 | 98.932 |
| 4 | 7.644 | 0.178 |
| 5 | 9.140 | 0.053 |
| 6 | 10.526 | 0.074 |
| 7 | 13.684 | 0.067 |
| Total | | 100.000 |

**OX04954**

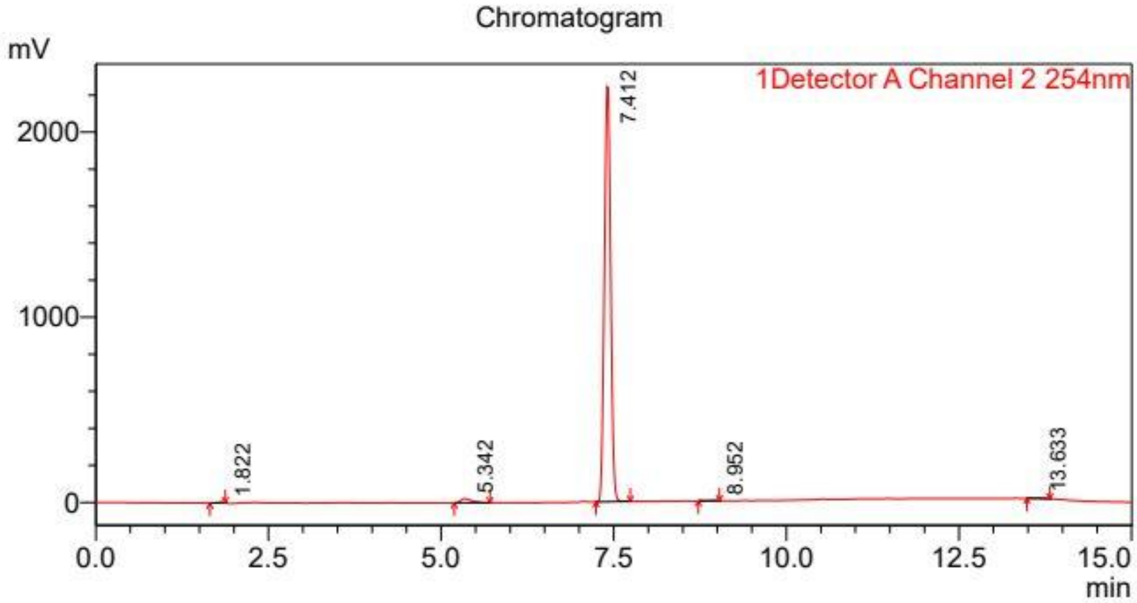


Detector A Channel 2 254nm

| Peak# | Ret. Time (min) | Area% |
|---|---|---|
| 1 | 1.822 | 0.185 |
| 2 | 5.342 | 1.771 |
| 3 | 7.412 | 97.654 |
| 4 | 8.952 | 0.150 |
| 5 | 13.633 | 0.239 |
| Total | | 100.000 |

**OX04955**

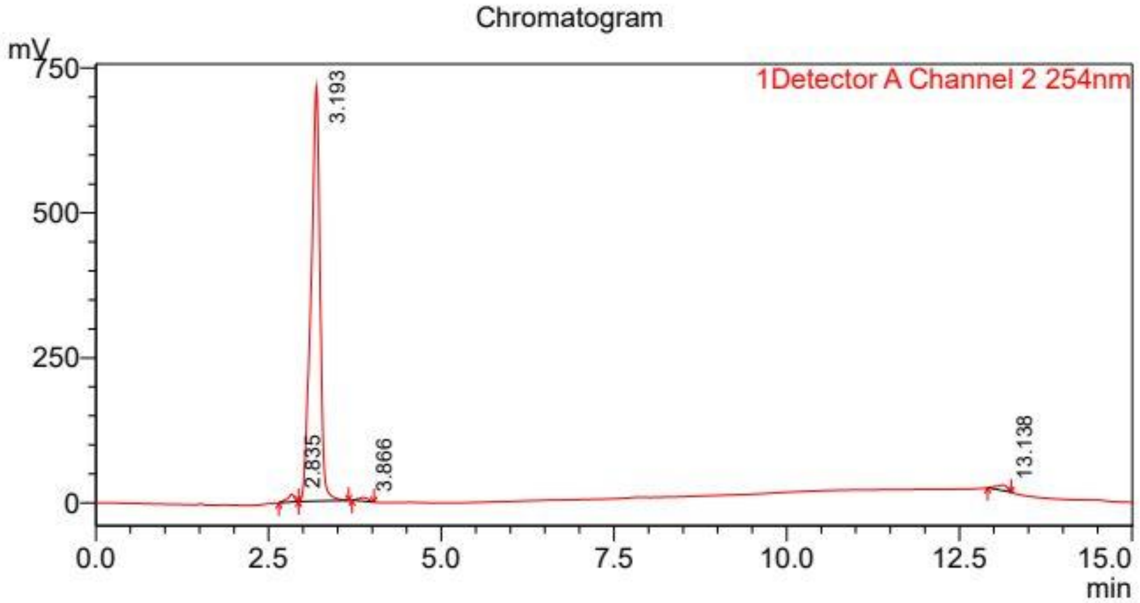


Detector A Channel 2 254nm

| Peak# | Ret. Time (min) | Area% |
|---|---|---|
| 1 | 2.835 | 1.399 |
| 2 | 3.193 | 96.196 |
| 3 | 3.866 | 0.916 |
| 4 | 13.138 | 1.489 |
| Total | | 100.000 |

**OX04956**

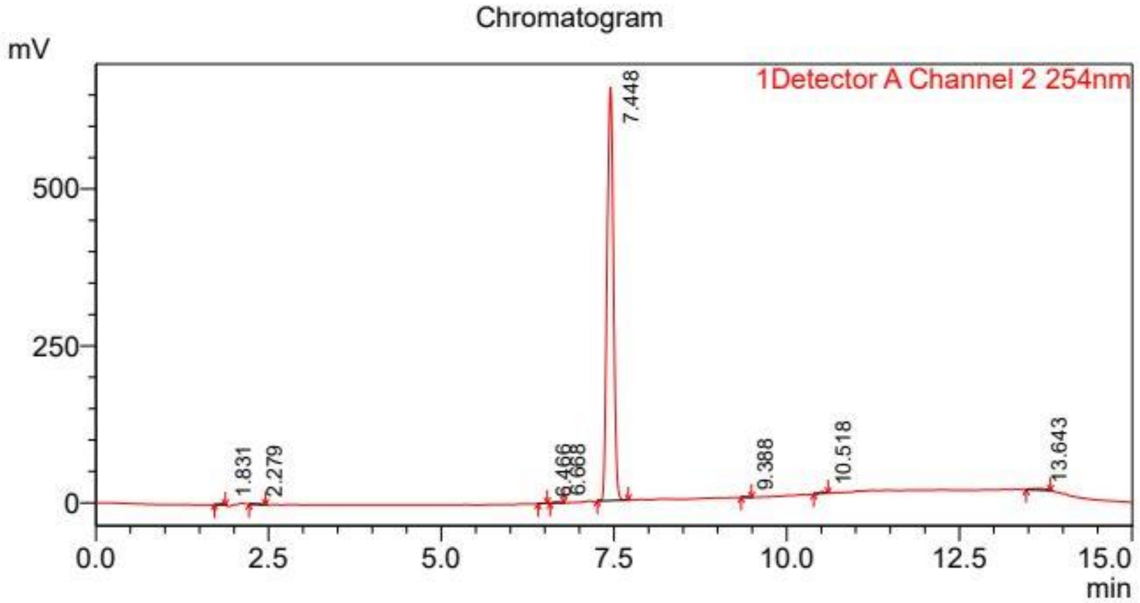


Detector A Channel 2 254nm

| Peak# | Ret. Time (min) | Area% |
|---|---|---|
| 1 | 1.831 | 0.230 |
| 2 | 2.279 | 0.205 |
| 3 | 6.466 | 0.069 |
| 4 | 6.668 | 0.234 |
| 5 | 7.448 | 98.029 |
| 6 | 9.388 | 0.059 |
| 7 | 10.518 | 0.368 |
| 8 | 13.643 | 0.807 |
| Total | | 100.000 |

**OX04957**

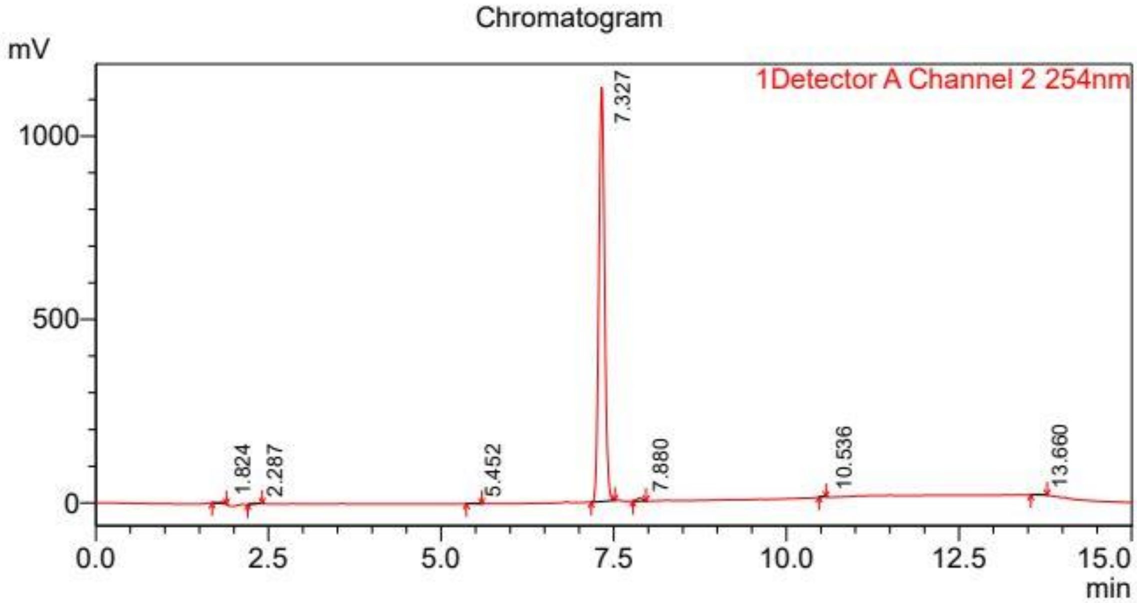


Detector A Channel 2 254nm

| Peak# | Ret. Time (min) | Area% |
|---|---|---|
| 1 | 1.824 | 0.410 |
| 2 | 2.287 | 0.381 |
| 3 | 5.452 | 0.164 |
| 4 | 7.327 | 98.053 |
| 5 | 7.880 | 0.693 |
| 6 | 10.536 | 0.082 |
| 7 | 13.660 | 0.217 |
| Total | | 100.000 |

**OX04958**

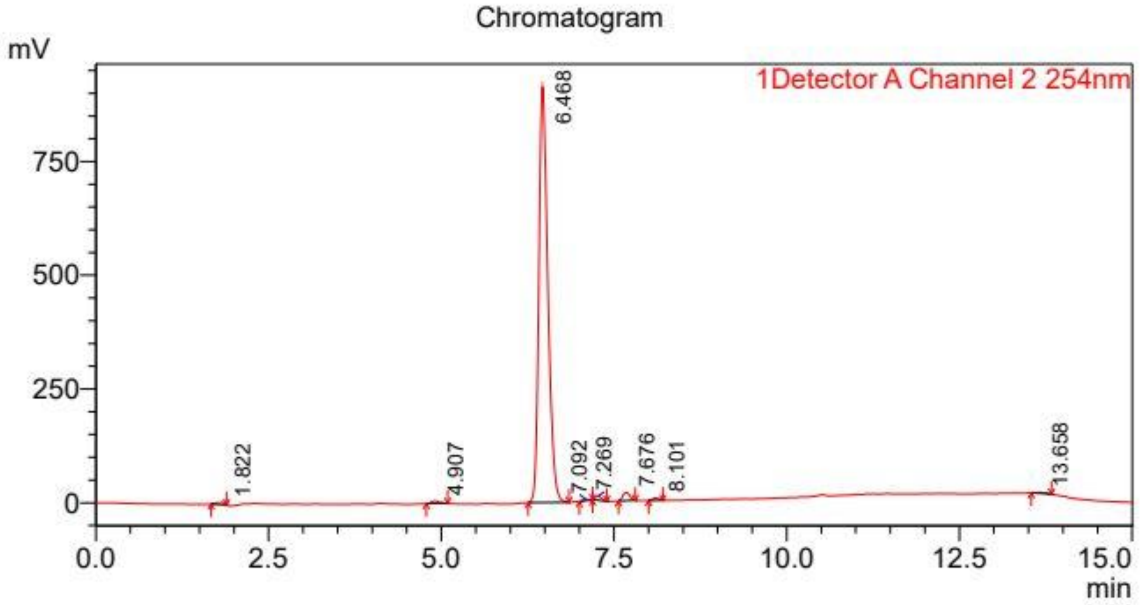


Detector A Channel 2 254nm

| Peak# | Ret. Time (min) | Area% |
|---|---|---|
| 1 | 1.822 | 0.444 |
| 2 | 4.907 | 0.558 |
| 3 | 6.468 | 96.016 |
| 4 | 7.092 | 0.192 |
| 5 | 7.269 | 0.638 |
| 6 | 7.676 | 1.503 |
| 7 | 8.101 | 0.419 |
| 8 | 13.658 | 0.231 |
| Total | | 100.000 |

**OX04959**

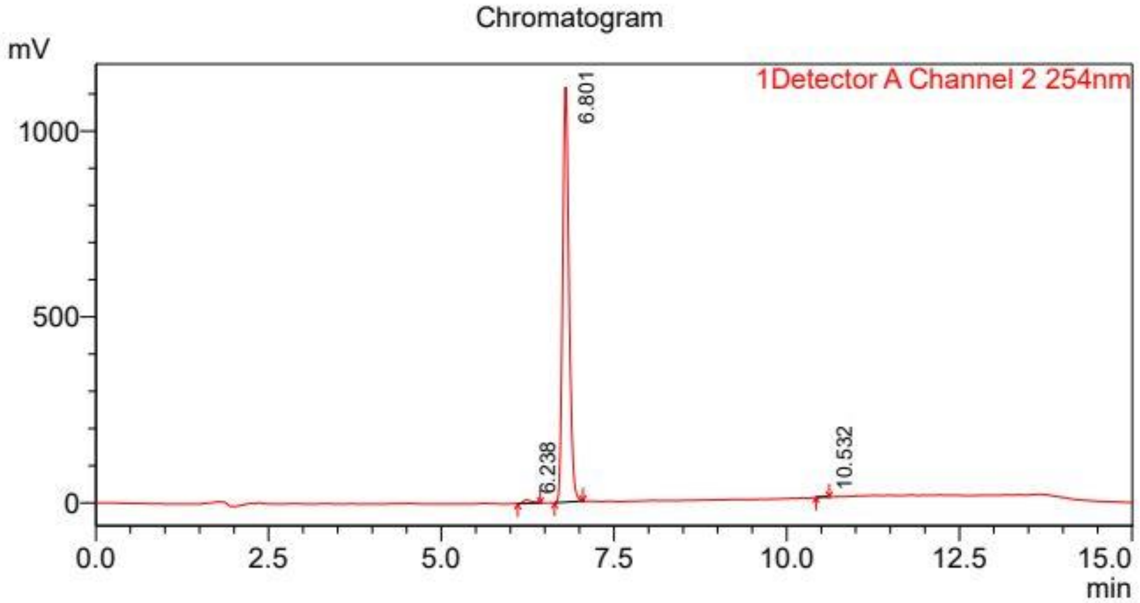


Detector A Channel 2 254nm

| Peak# | Ret. Time (min) | Area% |
|---|---|---|
| 1 | 6.238 | 1.055 |
| 2 | 6.801 | 98.719 |
| 3 | 10.532 | 0.226 |
| Total | | 100.000 |

**OX04960**

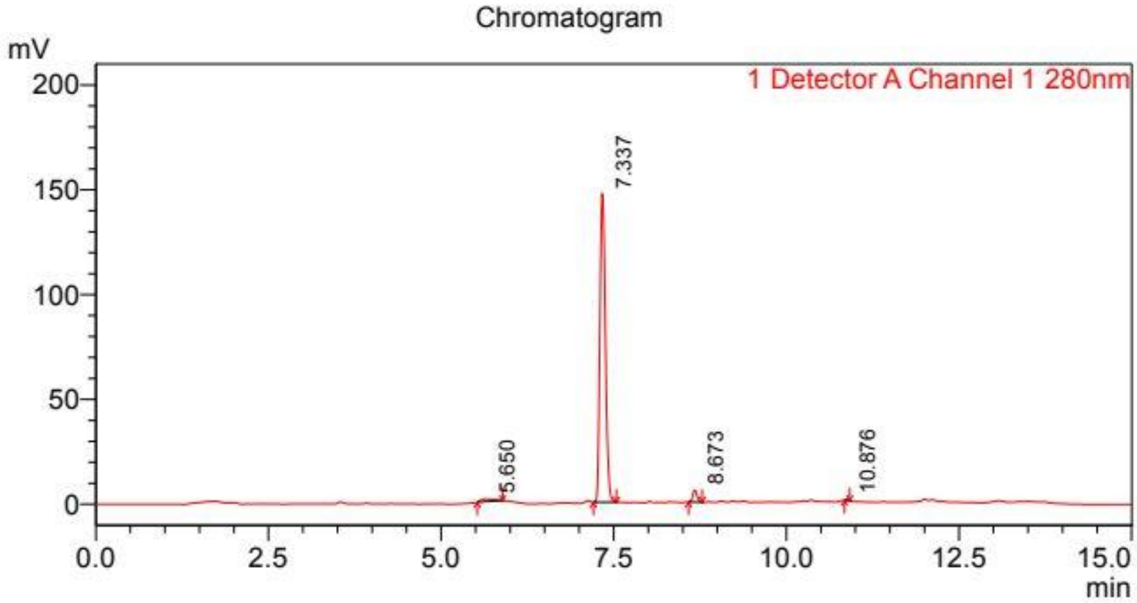


Detector A Channel 1 280nm

| Peak# | Ret. Time (min) | Area% |
|---|---|---|
| 1 | 5.650 | 1.857 |
| 2 | 7.337 | 95.238 |
| 3 | 8.673 | 2.756 |
| 4 | 10.876 | 0.148 |
| Total | | 100.000 |

**OX04961**

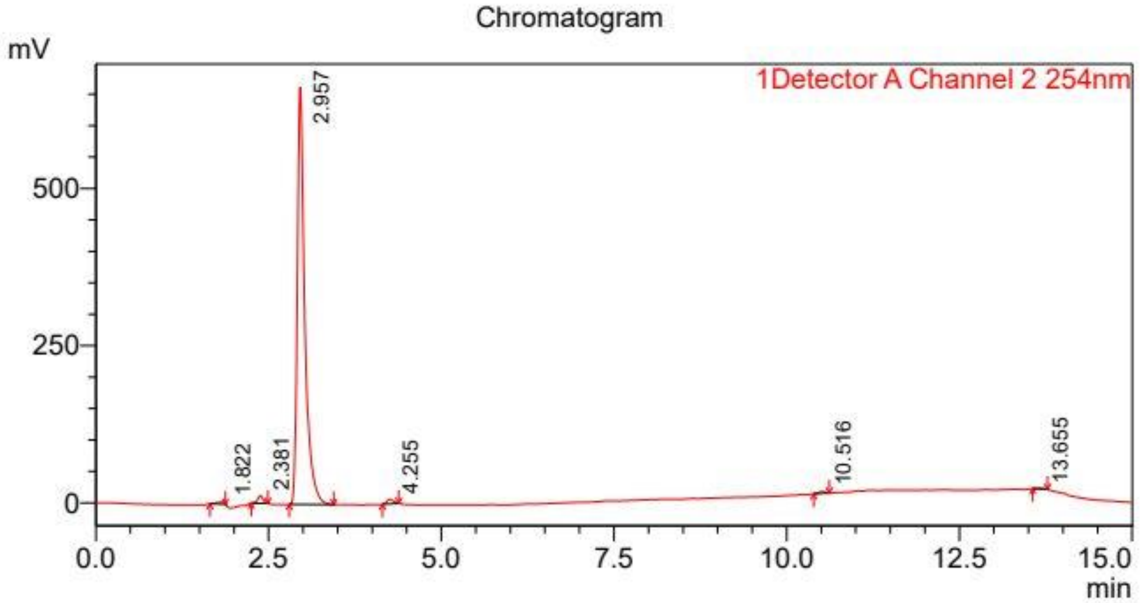


Detector A Channel 2 254nm

| Peak# | Ret. Time (min) | Area% |
|---|---|---|
| 1 | 1.822 | 0.467 |
| 2 | 2.381 | 1.245 |
| 3 | 2.957 | 96.753 |
| 4 | 4.255 | 0.961 |
| 5 | 10.516 | 0.347 |
| 6 | 13.655 | 0.227 |
| Total | | 100.000 |

**OX04962**

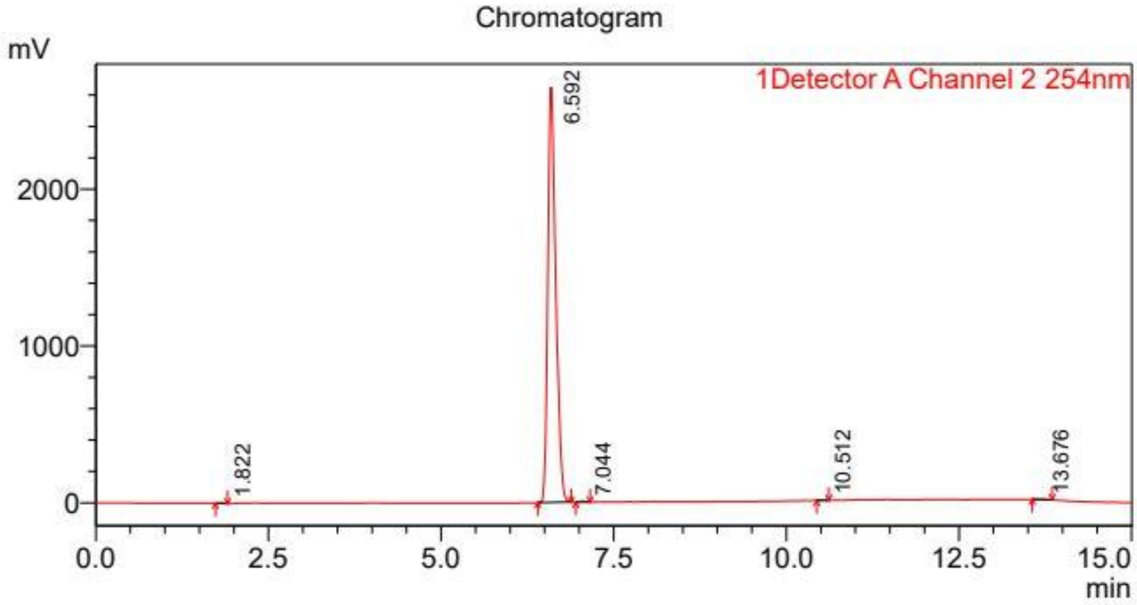


Detector A Channel 2 254nm

| Peak# | Ret. Time (min) | Area% |
|---|---|---|
| 1 | 1.822 | 0.112 |
| 2 | 6.592 | 99.524 |
| 3 | 7.044 | 0.176 |
| 4 | 10.512 | 0.056 |
| 5 | 13.676 | 0.131 |
| Total | | 100.000 |

**OX04963**

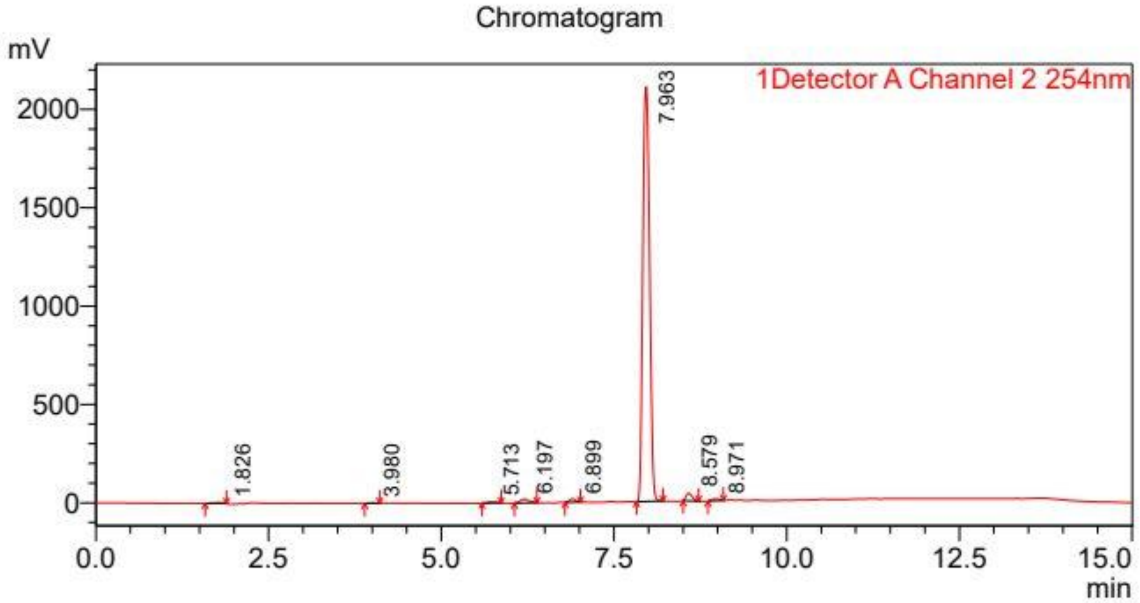


Detector A Channel 2 254nm

| Peak# | Ret. Time (min) | Area% |
|---|---|---|
| 1 | 1.826 | 0.364 |
| 2 | 3.980 | 0.109 |
| 3 | 5.713 | 0.349 |
| 4 | 6.197 | 1.096 |
| 5 | 6.899 | 0.691 |
| 6 | 7.963 | 95.281 |
| 7 | 8.579 | 1.595 |
| 8 | 8.971 | 0.515 |
| Total | | 100.000 |

**OX04964**

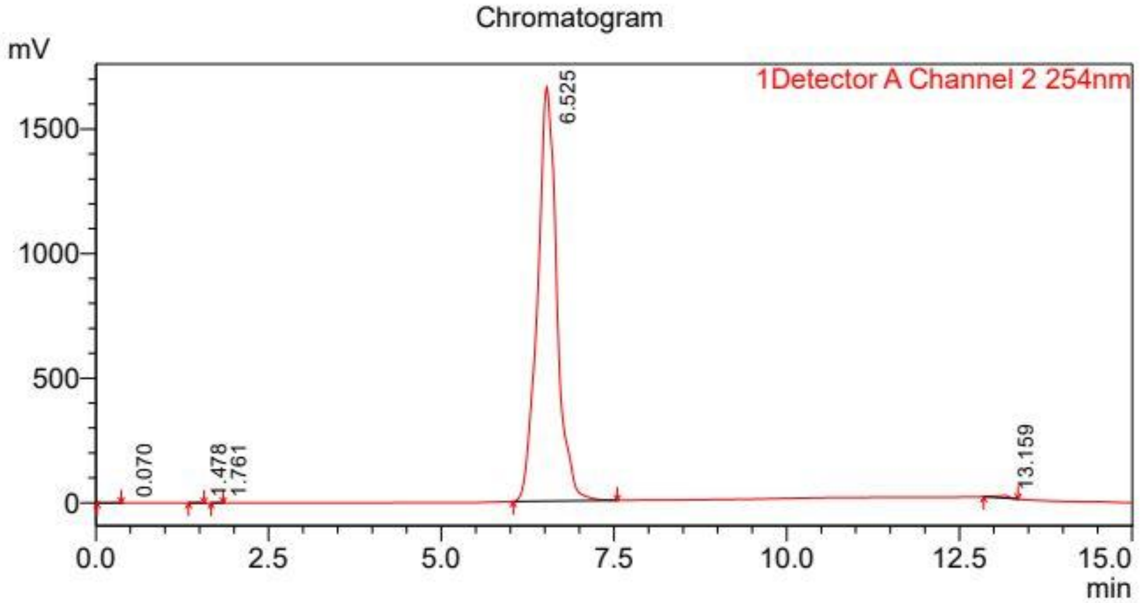


Detector A Channel 2 254nm

| Peak# | Ret. Time (min) | Area% |
|---|---|---|
| 1 | 0.070 | 0.005 |
| 2 | 1.478 | 0.067 |
| 3 | 1.761 | 0.012 |
| 4 | 6.525 | 99.482 |
| 5 | 13.159 | 0.434 |
| Total | | 100.000 |

**OX04965**

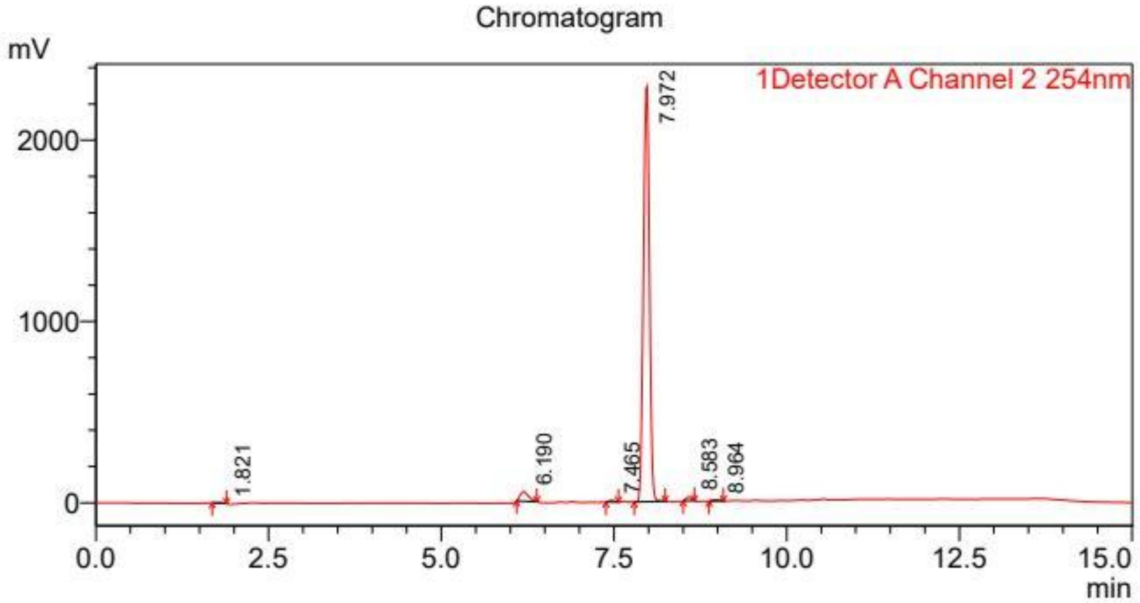


Detector A Channel 2 254nm

| Peak# | Ret. Time (min) | Area% |
|---|---|---|
| 1 | 1.821 | 0.237 |
| 2 | 6.190 | 3.121 |
| 3 | 7.465 | 0.355 |
| 4 | 7.972 | 95.148 |
| 5 | 8.583 | 0.790 |
| 6 | 8.964 | 0.349 |
| Total | | 100.000 |

**OX04966**

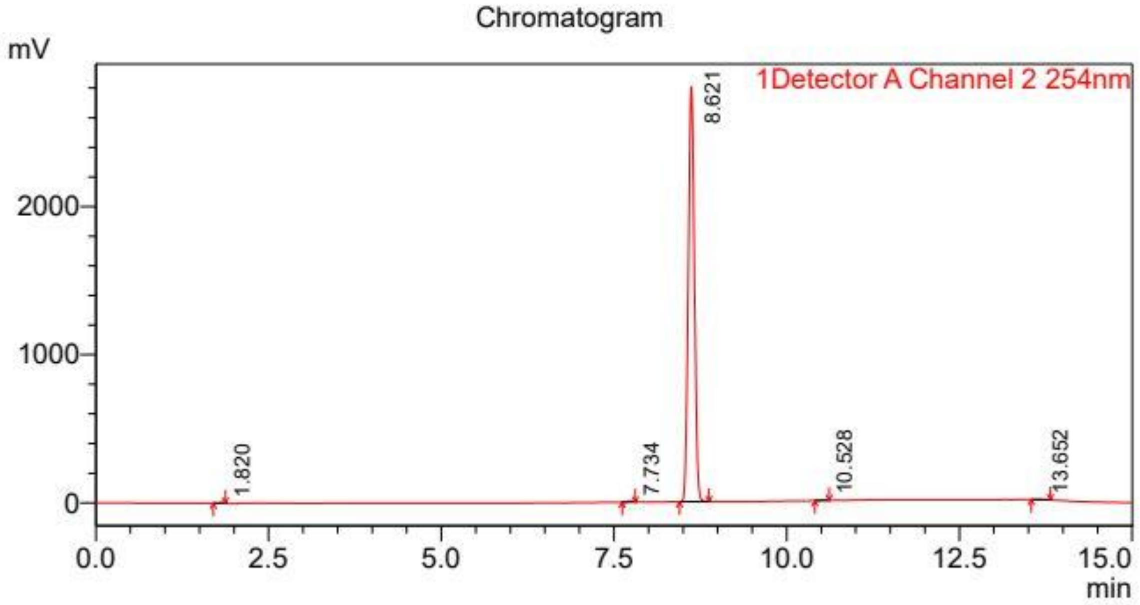


Detector A Channel 2 254nm

| Peak# | Ret. Time (min) | Area% |
|---|---|---|
| 1 | 1.820 | 0.118 |
| 2 | 7.734 | 0.234 |
| 3 | 8.621 | 99.426 |
| 4 | 10.528 | 0.094 |
| 5 | 13.652 | 0.128 |
| Total | | 100.000 |
